# Damaged glomeruli in proliferative pediatric lupus nephritis exhibit a C5a-C5aR1 induced fibrotic transcriptional program

**DOI:** 10.64898/2026.03.12.711136

**Authors:** Sarah McCuaig, Easton Elliott, Seth Anderson, David Smith, Julia Rood, Jael Gaines, Portia A Kreiger, Edward M Behrens

## Abstract

Lupus nephritis (LN) is a leading cause of morbidity in pediatric systemic lupus erythematosus (pSLE) due to suboptimal kidney remission rates and the sequelae of prolonged intensive immunosuppressive therapy. LN is patchy, with some glomeruli severely damaged while others remain histologically unaffected in the same kidney. Using spatial transcriptomic technology, we interrogated microanatomic transcriptional differences between histologically damaged and unaffected glomeruli in pSLE LN to understand local drivers of renal injury. Despite SLE being a disease of Type I interferon (IFN), IFN gene response does not associate with local glomerular damage. Rather, damage associates with a transcriptional module of higher expression of myeloid cell markers, *C5AR1* (encoding the receptor for complement component 5a [C5a]), early complement components, and fibrosis genes. Bulk RNA-sequencing of C5a stimulated human monocyte-derived-macrophages revealed upregulation of tissue-remodeling and fibrosis-related pathways reversible by the C5aR1 inhibiting drug avacopan. These same C5a-inducible fibrosis genes were significantly upregulated in histologically damaged versus unaffected LN glomeruli providing a mechanistic link between C5a-C5aR1 signaling and early fibrosis in proliferative lupus nephritis. Our data provide insight into an understudied connection between complement activation and fibrosis relevant in SLE and likely other inflammatory diseases of complement activation.

## Introduction

Lupus nephritis is a frequent complication of systemic lupus erythematosus (SLE) and is a leading cause of morbidity and mortality in this multiorgan system autoimmune disease. Among patients with pediatric SLE (pSLE), rates of lupus nephritis are 10-30% higher than their adult counterparts, renal disease arises earlier in the disease course and can be more aggressive at its onset (1). It is estimated that over 50% of patients with pSLE will develop lupus nephritis (LN) (2). Despite broad, intensive immunosuppressive therapy with systemic steroids, mycophenolate mofetil, B cell depleting agents, and/or cyclophosphamide, rates of complete remission remain suboptimal (3). Without complete remission, patients progress to develop chronic kidney disease requiring dialysis or renal transplant. Moreover, the morbidity from intensive systemic immunosuppressive therapy regimens remains high with infection being the greatest risk. Therefore, there is an urgent need to understand the mechanisms driving immune-mediated pathology in pediatric lupus nephritis to develop targeted therapeutic modalities and achieve high rates of complete remissions.

It is well understood that most cases of lupus nephritis are initiated by glomerular deposition of immunoglobulin and complement. This leads to the local production of a variety of inflammatory mediators that ultimately drive tissue damage. Type I interferons (IFNs) are thought to be central lynchpins in SLE end organ pathology, including in lupus nephritis. Type I IFNs and interferon stimulated genes (ISGs) are elevated in the blood of patients with lupus nephritis and their levels correlate with disease activity in some cohorts (4, 5). Furthermore, lupus nephritis renal biopsies show increased ISG expression (6, 7). Evidence from murine models also points toward a mechanistic role of type I IFNs in lupus nephritis. For example, IFN alpha receptor knockout mice were protected from nephritis induced with anti-GBM serum (8). Furthermore, in lupus predisposed BXSB mice, IFN alpha receptor blockade reduced glomerulonephritis scores when initiated prior to clinical evidence of renal disease, however had no effect on glomerulonephritis scores when initiated after the onset of proteinuria, suggesting that type I IFNs may be important early in the disease process (9).

Given that much of our knowledge of deranged inflammatory pathways in SLE comes from analysis of the peripheral blood, homogenized whole kidney tissue, or murine models that do not always faithfully recapitulate human disease, we do not currently know whether type I IFNs are proximal drivers of glomerular damage in human pediatric lupus nephritis. Therefore, it is not clear that type I IFNs will be effective therapeutic targets in LN. Human trials of type I IFN blockade in lupus nephritis are ongoing but until now have been inconclusive. Anifrolumab, a type I interferon receptor monoclonal antibody, did not significantly improve renal function in patients with lupus nephritis in a Phase II study, though at higher doses it induced clinically significant improvements in exploratory endpoints including complete renal response and glucocorticoid reductions (10).

Here, using spatial transcriptomic technology, we interrogate microanatomic transcriptional differences between histologically damaged and unaffected glomeruli and adjacent tubules in diagnostic renal biopsies from pediatric patients with Class III pSLE LN to understand local drivers of renal injury. We hypothesized that the local Type I interferon (IFN) response would be higher in histologically damaged glomeruli and adjacent tubules, however this was not the case. Instead, damaged glomeruli and adjacent tubules are marked by a program of local complement and fibrosis-related gene expression. *C5AR1* is more highly expressed in histologically damaged glomeruli predominately by myeloid cells. Direct complement factor 5a (C5a) stimulation of human monocyte-derived macrophages upregulates tissue-remodeling and fibrosis related pathways based on bulk RNA-sequencing analysis. These C5a-inducible fibrosis genes are more highly expressed in histologically damaged Class III LN glomeruli. These results demonstrate that fibrosis-related programs are active at the time of pSLE diagnosis and highlight an important link between complement signaling and fibrosis pathways in lupus nephritis poised for therapeutic targeting with existing small molecule inhibitors of C5aR1.

## Results

### Spatial transcriptomics accurately identifies microanatomic renal structures in biopsies from pediatric patients with lupus nephritis, non-lupus glomerulonephritis, and healthy controls

To investigate whether Type I IFNs are proximal drivers of renal injury in pSLE LN, we utilized the Visium 10X platform to perform spatial transcriptomics on formalin-fixed paraffin-embedded (FFPE) pediatric renal biopsies after H&E staining. We focused our analysis on biopsy specimens from patients diagnosed with ISN/RPS Class III proliferative LN (N=3). By definition, fewer than 50% of glomeruli show evidence of histologic damage in Class III LN (11, 12), thus providing an intrapatient or “isogenic” control to specifically make comparisons between histologically damaged and unaffected glomeruli. Biopsies from patients with pure Class V LN (N=3), which is both clinically and histologically distinct from proliferative LN, were analyzed for comparison. To assess whether transcriptional changes were lupus-specific or a feature of immune complex glomerulonephritis in general, biopsies from patients with post-streptococcal glomerulonephritis (N=1) and C3 glomerulopathy (N=1) were used as disease controls. Renal resection tissue from pediatric patients who underwent nephrectomy for non-immune indications were used as “healthy” controls (N=3). Patient demographic data is detailed in **Supplemental Table S1**. The tissues were obtained from archived specimens from clinical pathology and importantly all but one of the lupus biopsies (a Class V LN patient) were obtained at the time of lupus diagnosis. Patients had no or minimal exposure to immune suppressive medications at the time of biopsy. RNA was successfully obtained from these archived specimens and spatially barcoded gene expression libraries were mapped to 55μM diameter spots and used for probe-based gene expression analysis of ∼18,000 human genes. A pediatric pathologist then identified and annotated histologically damaged and unaffected glomeruli in the Class III LN biopsies based on H&E staining (**Figure 1A,F**). Unsupervised graph-based clustering of 55μM spots generated clusters that reflect the underlying renal microanatomy, demonstrating success of the tissue and our processing of it in generating biologically meaningful information (**Figure 1B-E)**. Myeloid cell glomerular infiltration is a feature of damaged glomeruli in Class III LN (11). Increased expression of the myeloid cell marker *CD68* was seen in histologically damaged compared to unaffected glomeruli, confirming concordance between pathologist annotation of damaged glomeruli and expected transcriptional changes (**Figure 1G**).

**Figure 1.**
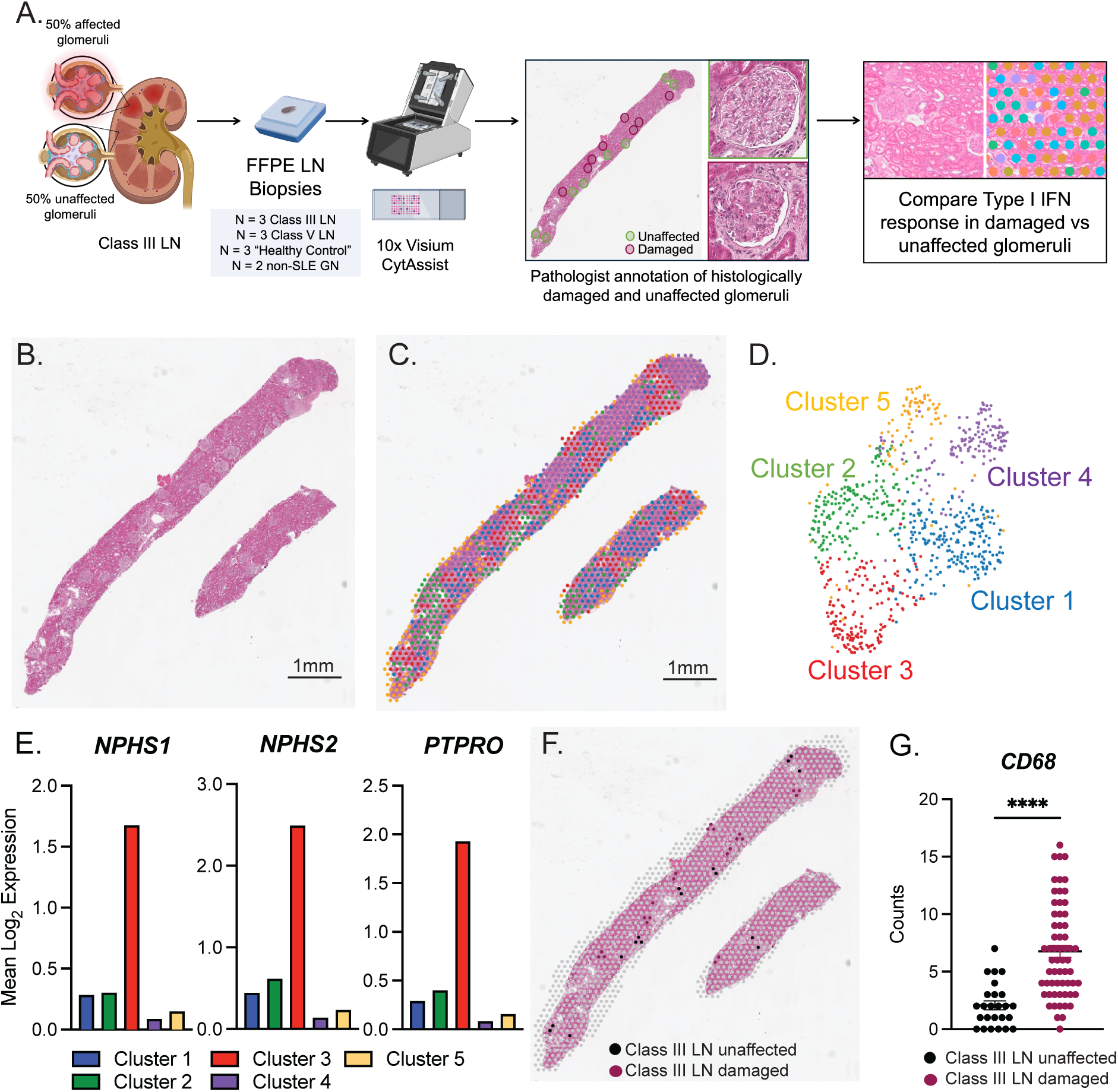
Unbiased graph-based clustering identifies glomeruli transcriptionally that corresponds to microanatomic location in renal biopsies. **(A)** Spatial transcriptomics using the Visium 10x platform was performed on FFPE renal biopsies from pediatric patients with Class III LN (N=3), Class V LN (N=3), C3 GN (N=1), PSGN (N=1), and “healthy” controls (N=3). Pathologist annotated histologically damaged versus unaffected glomeruli in Class III LN and then compared the Type I IFN response in damaged versus unaffected glomeruli. **(B)** Representative H&E section from LN biopsy. **(C)** Overlay of 55uM spots colored coded based on unbiased graph-based clustering. **(D)** UMAP projection of graph-based clusters. **(E)** Cluster 3 (red) is defined by high expression of glomerular genes *NPHS1, NPHS2, PTPRO* and spatially correlates with glomeruli as seen in **(C)**. Values are mean-normalized UMI counts in the cluster. **(F)** Representative H&E section from Class III LN biopsy showing pathologist annotation of histologically damaged (pink) and unaffected (black) glomeruli. **(G)** Expression of the myeloid marker *CD68* in histologically damaged (pink) and unaffected (black) glomeruli. Each dot is expression from a 55uM spot. N=3 independent Class III LN patients. **** p<0.0001, unpaired T-test.

### Interferon-stimulated gene expression is unchanged comparing histologically damaged and unaffected Class III LN glomeruli

To investigate whether Type I IFNs are proximal drivers of glomerular damage in Class III LN we compared expression of interferon stimulated genes (ISGs) in histologically damaged and unaffected Class III LN glomeruli. We computed an ISG score by determining the average expression of five ISGs (*IFI44, IFI44L, MX1, IFIT1, HERC6*) in each glomerular spot. This set of ISGs was selected based on a literature review of ISGs commonly associated with SLE and/or LN (4, 13–15). Amongst patients with Class III LN, there was heterogeneity in the glomerular ISG score overall (**Figure 2A**). However, when histologically damaged glomeruli were compared to unaffected glomeruli for each patient, there was no difference in ISG expression for two of the three patients (Class III LN 2, Class III LN 3) and a statistically significant but small magnitude difference in the ISG score for one of the three patients (Class III LN 1) (**Figure 2A**). These data do not support our hypothesis that there is increased Type I interferon signaling in histologically damaged glomeruli in Class III LN. Of note, glomeruli from control biopsies had low ISG scores and there was heterogeneity in ISG score amongst the Class V LN and non-SLE GN biopsies (**Figure 2A**). To determine whether this lack of difference in ISG score between damaged and unaffected Class III LN glomeruli was generalizable to other ISGs, we looked for individual genes that correlated with the ISG score across all glomeruli. Genes were filtered for those with a Pearson Correlation >0.5 and Benjamini-Hochberg corrected p-value <0.05. As expected, the five ISGs included in the ISG score were significantly correlated and the other significantly correlated genes were also ISGs (*XAF1, IFI6, ISG15, CMPK2, OAS3, RSAD2, IFI27*) (**Figure 2B**). We used this entire ISG list to generate an expanded ISG score using the average expression of these twelve ISGs computed for each glomerular spot. Using this expanded ISG score, there was also no difference in ISG expression comparing histologically damaged and unaffected Class III LN glomeruli (**Figure S1A**). Moreover, when we rank-ordered genes that correlated with the 5 gene ISG score, removed the genes that are components of the score, and then performed GSEA analysis on the significantly correlated genes, the Hallmark Interferon Alpha response and Hallmark Interferon Gamma Response pathways were significantly enriched (**Figure S1B**) suggesting that our 5 gene score correlates with ISG expression more broadly. Taken together these data confirm that there is no difference in global ISG expression between histologically damaged and unaffected Class III LN glomeruli.

**Figure 2.**
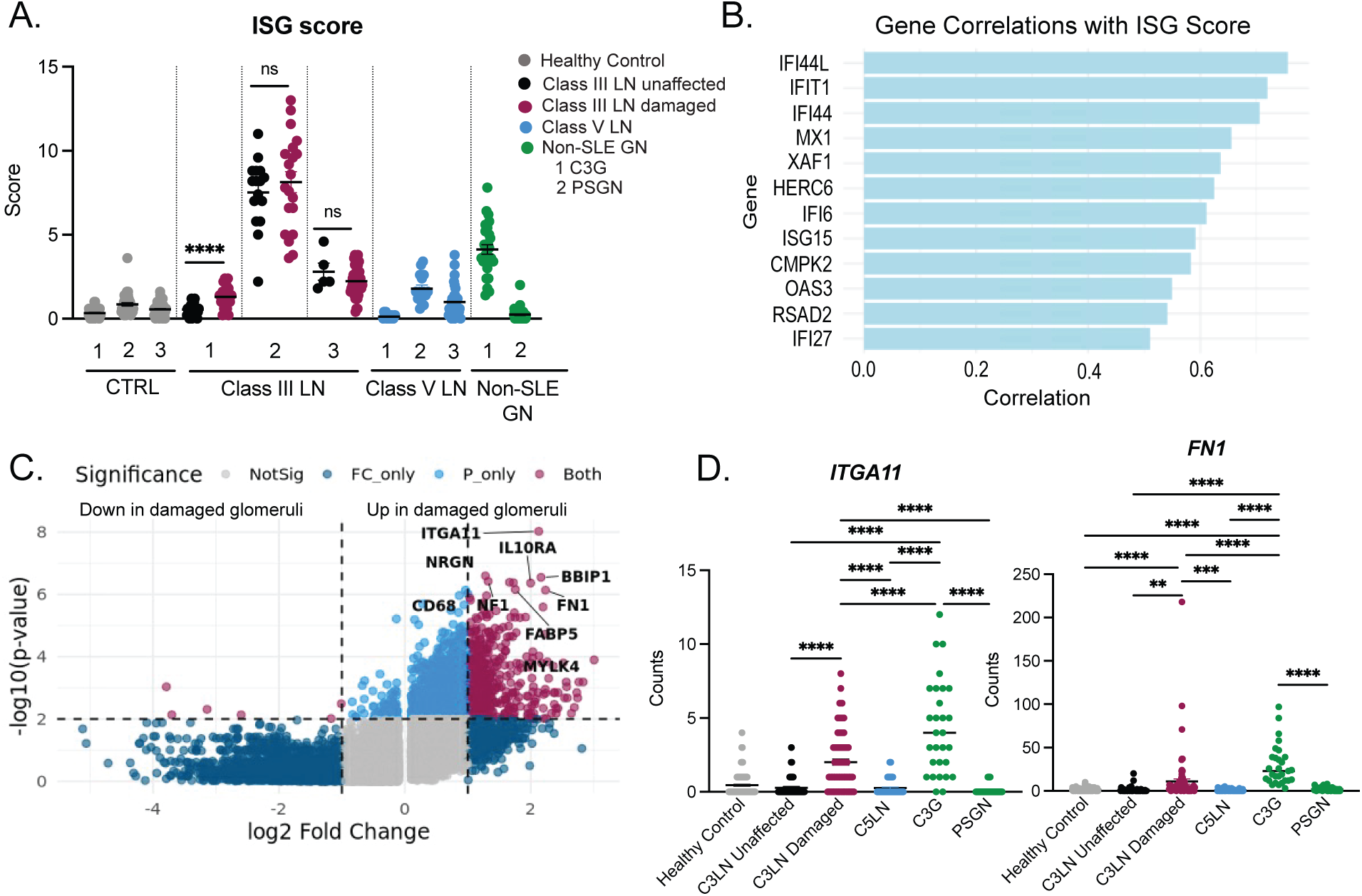
Interferon-stimulated gene (ISG) score is unchanged in damaged versus unaffected glomeruli in Class III LN but damaged glomeruli are transcriptionally distinct from unaffected glomeruli and show upregulation of fibrosis genes at the time of lupus diagnosis. **(A)** ISG score computed as the average of the expression of *IFI44, IFI44L, MX1, IFIT1, HERC6*. Each dot is expression from a 55uM spot. X axis numbers represent individual patients in each disease subset. Non-SLE patient 1 carries a diagnosis of C3 glomerulopathy (C3G). Non-SLE patient 2 carries a diagnosis of post-strep glomerulonephritis (PSGN) and separate histologic sections of renal biopsy from this patient were run on independent Visium 10X runs. **** p< 0.0001 unpaired T-test for comparison between damaged and unaffected Class III LN glomeruli. **(B)** Pearson correlation for genes that correlate with the ISG score across all glomeruli. Filtered for correlation > 0.5 and Benjamini-Hochberg corrected p-value <0.05. **(C)** Volcano plot showing differentially expressed genes comparing histologically damaged to unaffected glomeruli Class III LN glomeruli. Positive log2 fold changes are genes with increased expression in damaged compared to unaffected glomeruli. Significantly differentially expressed genes are those with a log2 fold change >1.0 and p-value < 0.01. **(D)** Highest differentially expressed genes comparing damaged to unaffected glomeruli are fibrosis-related genes *ITGA11, FN1*. Each dot is expression from a 55μM spot. N=3 healthy control. N=3 Class III LN. N=3 Class V LN. N=1 C3G. N= 1 PSGN. One way ANOVA with Tukey’s post-test for multiple comparisons. **p<0.01, ***p<0.001, ****p<0.0001.

### Histologically damaged and unaffected Class III LN glomeruli have distinct transcriptional programs

Given lack of difference in ISG expression between histologically damaged and unaffected Class III LN glomeruli, we next took an unbiased approach and assessed transcriptional differences in damaged and unaffected glomeruli in Class III LN using differential gene expression with FindMarkers (Seurat) for exploratory analysis. Damaged Class III LN glomeruli have a unique set of upregulated transcripts compared to unaffected Class III LN glomeruli (**Figure 2C**, **Supplemental Table S2**). Among the highest differentially expressed genes in histologically damaged glomeruli were the fibrosis-related genes *ITGA11* (encoding integrin alpha 11) and *FN1* (encoding fibronectin 1) (**Figure 2D**) even though these were early stage biopsies taken at the time of lupus diagnosis without significant histologic fibrosis. When controlling for patient-level differences using a linear mixed effects model with patient as a random effect and glomerular disease state (damaged/unaffected) as a fixed effect in the model, the increased expression of *ITGA11* and *FN1* in damaged compared to unaffected glomeruli remained statistically significant (**Figure S1C**). Interestingly, this fibrosis-gene expression was a unique feature of proliferative Class III LN as *ITGA11* and *FN1* expression was low in Class V LN (**Figure 2D**). However, glomerular fibrosis gene expression was not lupus-specific as glomeruli from the C3 glomerulopathy biopsy also expressed high amounts of these fibrosis genes (**Figure 2D**).

### Complement component 5a receptor 1 expression is elevated on histologically damaged versus unaffected Class III LN glomeruli

Given that data sparsity is inherent to spatial transcriptomic analyses, we liberalized our definition of differentially expressed genes (log2 FC > 0.5, p-value < 0.05) and assessed this expanded list for genes of known immunologic function and/or genes targetable with existing therapeutics. Interestingly, *C5AR1* which encodes Complement component 5a receptor 1 was more highly expressed on histologically damaged compared to unaffected glomeruli **(Figure 3A**) and this difference remained statistically significant when analyzed using the linear mixed effects model described above (**Figure S1C**). Elevated *C5AR1* expression on histologically damaged glomeruli was a unique feature of Class III compared to Class V LN, but again was not lupus-specific as the C3 glomerulopathy biopsy also expressed high amounts of *C5AR1* (**Figure 3A**). C5aR1 is the target of the small molecule inhibitor avacopan, which is FDA-approved for use in ANCA-associated vasculitis, however its efficacy has not been explored in lupus nephritis. Given C5aR1 is a potentially druggable target in LN, we next sought to better understand which intrarenal cell types most abundantly express *C5AR1*. A limitation to the Visium platform used is that we could not infer single cell resolution transcripts from the 55uM spots. Therefore, we analyzed publicly available single cell sequencing data from adult LN biopsies from the Accelerating Medicines Partnership (AMP) (16) and found that *C5AR1* is most highly expressed on the myeloid clusters (**Figure 3B**). To confirm these observations at the protein level, we performed immunofluorescence co-staining on frozen sections of pediatric LN and non-SLE LN biopsies and control kidney resection specimens for C5aR1 and other cell-type specific markers: CD68 (myeloid), PODXL (podocyte), CD31 (endothelial), aSMA (mesangial). There was no significant C5aR1 staining in the control or Class V LN biopsies. However, C5aR1 was clearly expressed at the protein level in Class III LN and non-SLE GN (C3G and PSGN) biopsies. Representative images from a glomerulus from a Class III LN biopsy are shown (**Figure 3C**). Next, to understand which cell types predominately expressed C5aR1 in the Class III LN (N=2 patients) and non-SLE GN biopsies (N=1 PSGN, N=1 C3G), we performed pixel-intensity based colocalization analysis of C5aR1 with CD68, PODXL, CD31, and aSMA in glomeruli. A Pearson Correlation of 1 reflects perfect overlay of the fluorescence signals. C5aR1 fluorescence correlated most closely with CD68 suggesting predominant myeloid expression (**Figure 3D**). This is concordant with the transcriptional data from the AMP dataset. There was minimal C5aR1 fluorescence overlay with PODXL, however overlay with CD31 and aSMA was heterogeneous, suggesting some possible endothelial and mesangial C5aR1 expression (**Figure 3D**).

**Figure 3.**
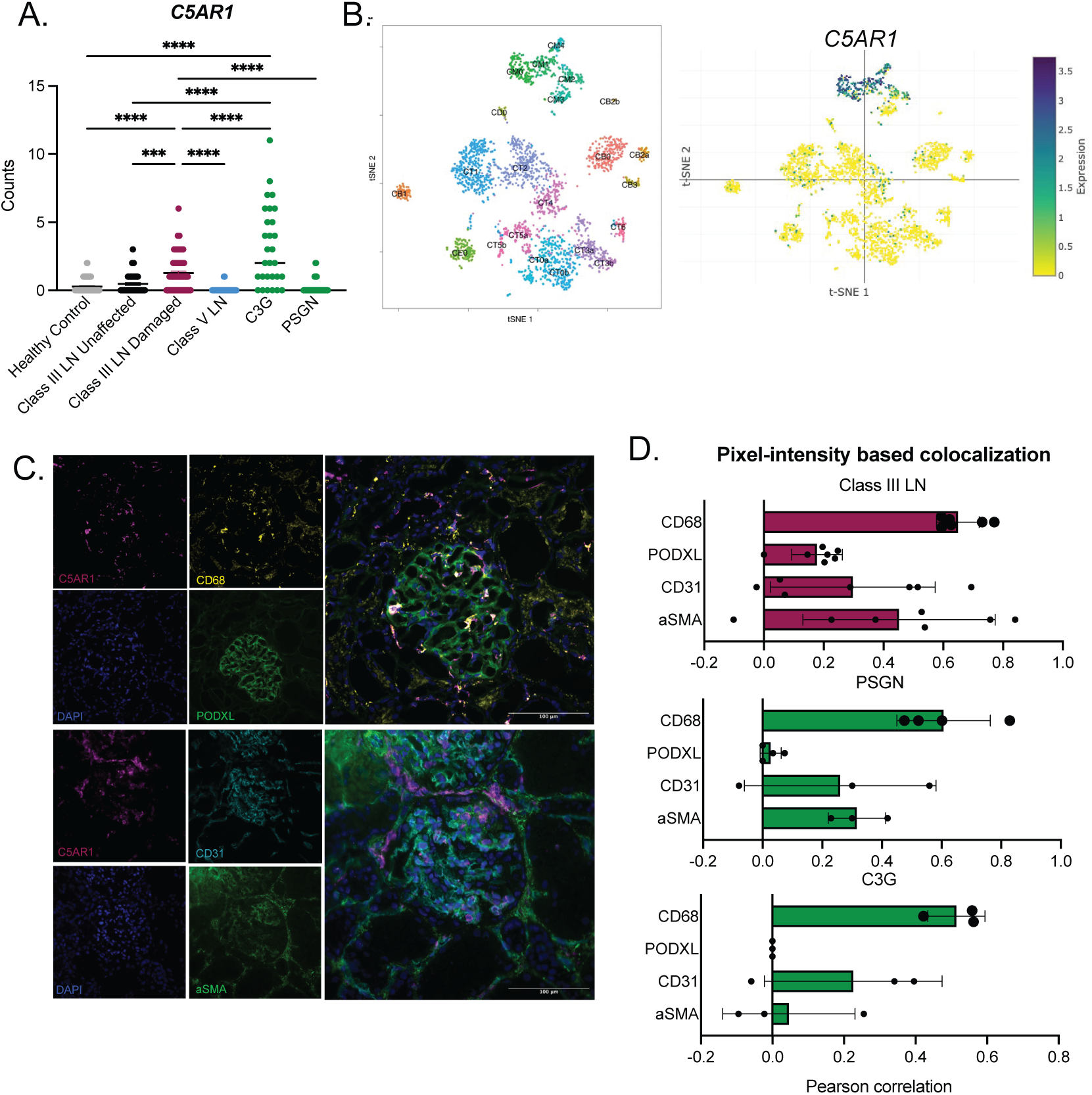
Histologically damaged glomeruli have increased expression of C5aR1. **(A)** *C5AR1*, the target of avacopan, is significantly elevated in damaged compared to unaffected Class III LN glomeruli. Each dot is expression from a 55μM spot. N=3 healthy control. N=3 Class III LN. N=3 Class V LN. N=1 C3G. N=1 PSGN. One way ANOVA with Tukey’s post-test for multiple comparisons. **(B)** Overlay of *C5AR1* expression on UMAP from AMP Dataset (16). **(C)** Representative immunofluorescence (IF) co-staining of C5aR1 with cell-type specific markers CD68 (myeloid), PODXL (podocyte), CD31 (endothelial), aSMA (mesangial) in a glomerulus from a biopsy of a patient with Class III LN. **(D)** IF pixel-intensity based colocalization analysis of C5aR1 with CD68, PODXL, CD31, aSMA in Class III LN glomeruli and glomeruli from patients with non-SLE GN where C5aR1 was detectable at the protein level. Pearson correlation of 1 is perfect overlay. Dots are single glomeruli. Class III LN N=2 patients, C3G N=1 patient, PSGN N=1 patient.

### Upregulation of myeloid and complement gene modules is a unique feature of Class III LN glomeruli compared to Class V LN glomeruli

Notably, glomeruli from patients with Class V LN do not express high amounts of *C5AR1* compared to histologically damaged Class III LN glomeruli. Pure Class V LN has a clinically distinct phenotype from Class III LN. Therefore, we next sought to explore transcriptional differences between Class III and Class V LN glomeruli. We performed differential gene expression analysis comparing all Class III LN glomeruli to Class V LN glomeruli and found a unique gene module that is upregulated in Class III LN glomeruli (**Figure 4A**, **Supplemental Table S3**). Reactome pathway analysis of the top differentially expressed genes revealed myeloid cell and complement pathways among the top 10 upregulated pathways (**Figure 4B**). Myeloid cell markers *CD163* and *CD68* were significantly upregulated in damaged Class III LN glomeruli compared to Class V glomeruli and interestingly were also significantly upregulated in unaffected Class III LN compared to Class V glomeruli (**Figure 4C**). Local glomerular expression of early complement components (*C1QA*, *C1QB*, *C1QC*) was a unique feature of Class III compared to Class V LN (**Figure 4D, E**). To understand which intraglomerular cell type was expressing these early complement components, we analyzed the AMP single cell dataset for C1Q component expression and found that myeloid cells most predominately expressed C1Q components (**Figure 4F**). Given overlapping cell type expression of *C5AR1* and C1Q component genes, we analyzed the myeloid cell subsets defined in the AMP dataset (16) for *C5AR1* and *C1QA, B, C* positivity. All *C5AR1*^+^ CM4 M2-like CD16^+^ macrophages also expressed *C1Q* component genes (**Figure 4G**). While only approximately half of CM2 Tissue Resident Macrophages were *C5AR1*^+^, among the *C5AR1*^+^ cells almost all were also *C1Q* component positive (**Figure 4G**). This contrasts with CM0 infiltrating inflammatory monocytes that were almost universally *C5AR1^+^* but a smaller proportion of the *C5AR1*^+^ cells were *C1Q* component positive (**Figure 4G**). This suggests that factors in the tissue microenvironment of the kidney are driving *C1Q* component expression in *C5AR1^+^* myeloid cells.

**Figure 4.**
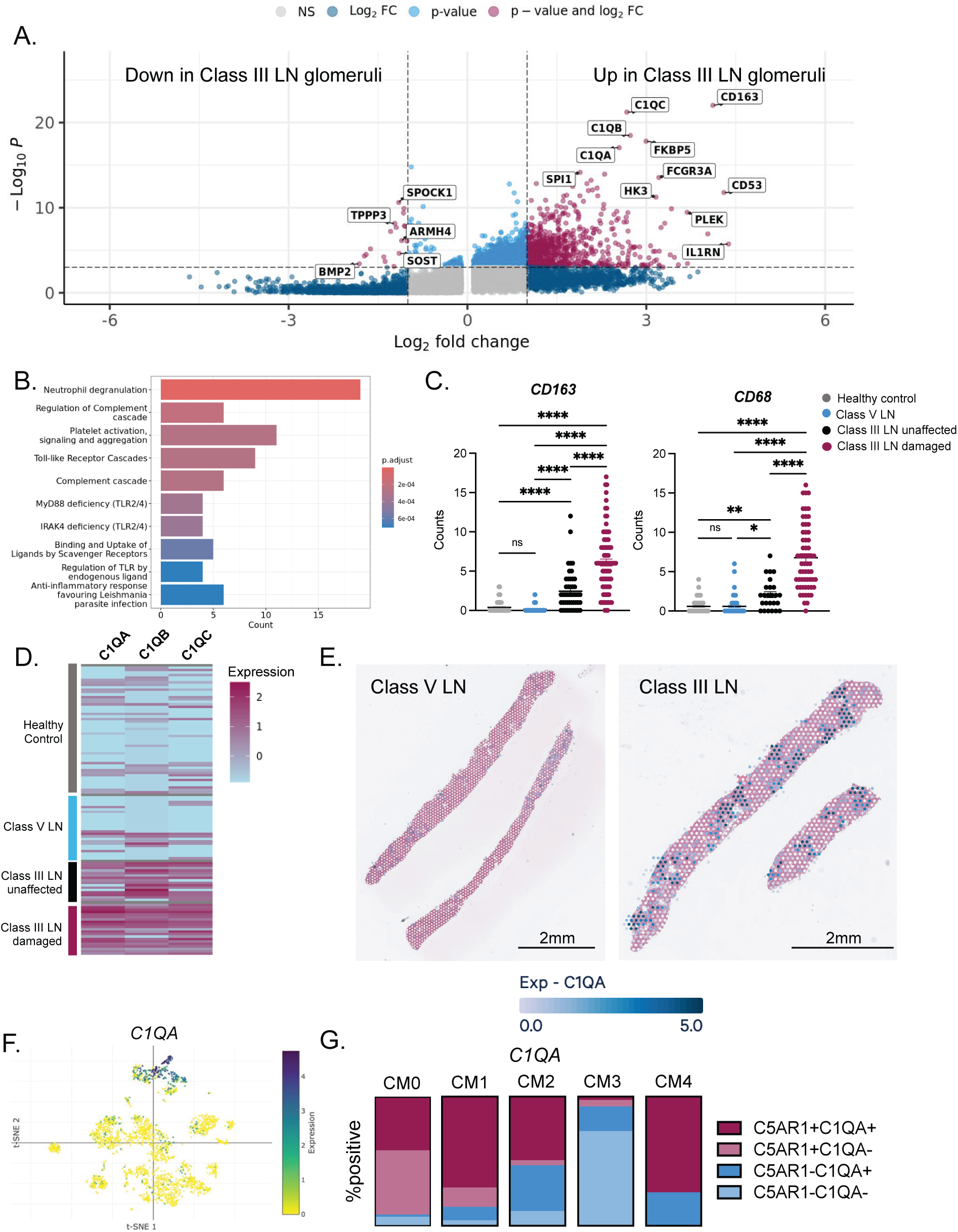
Myeloid and complement gene modules are upregulated in Class III versus Class V LN glomeruli. **(A)** Enhanced volcano plot showing differentially expressed genes comparing all Class III to Class V LN glomeruli. Positive log2 fold changes are genes with increased expression in Class III compared to Class V LN glomeruli. Significantly differentially expressed genes are those with a log2 fold change > 1 and p-value < 0.01. **(B)** Reactome pathway analysis of differentially expressed genes in Class III versus Class V LN glomeruli (Top 10 upregulated pathways). **(C)** Myeloid marker *CD163* and *CD68* expression in glomerular spots. Each dot is expression from a 55μM spot. N=3 healthy control. N=3 Class III LN. N=3 Class V LN. One way ANOVA with Tukey’s post-test for multiple comparisons. **(D)** Heatmap showing expression of *C1QA, C1QB, C1QC.* Reflects expression from 55μM glomerular spots. N=3 healthy control. N=3 Class III LN. N=3 Class V LN. **(E)** Representative H&E stained histologic sections from a Class V and Class III LN biopsy overlaid with log2 normalized expression of *C1QA* from each 55μM spot. **(F)** C1Q components are most highly expressed on myeloid cells in the AMP Dataset. **(G)** Proportion of cells of each myeloid subtype that are *C5AR1^+^*and/or *C1QA*^+^ in the AMP Dataset.

### Periglomerular tubules next to damaged Class III glomeruli show increased early complement and fibrosis gene expression

Our initial analysis focused exclusively on the glomerular compartment of the kidney. The rationale for this was twofold: first, it allowed for discrete isolation and analysis of histologically damaged and unaffected glomeruli and second, it is the microanatomic unit of the kidney that is damaged by immune complex deposition. However, there is a growing body of evidence supporting a key role of the tubulointerstitial compartment in LN pathogenesis and progression to end stage renal disease (17–19). As such, we next sought to understand transcriptional changes in the tubulointerstitial compartment based on spatial proximity to histologically damaged or unaffected glomeruli in Class III LN. 55μM spots overlaying the tubulointerstitial compartment were manually annotated as “close” to damaged or unaffected glomeruli if they immediately bordered a histologically damaged or unaffected glomerulus respectively (**Figure 5A**). We also annotated 55μM spots in the tubulointerstitial compartment that were “far” from glomeruli, defined as 2 layers of 55μM spots away from a glomerulus (**Figure 5A**). We first interrogated whether ISG expression differs between periglomerular tubules close to damaged versus unaffected Class III LN glomeruli. Like the glomeruli, there was interpatient heterogeneity in the amount of ISG expression for each patient and there was no difference in ISG expression between tubules close to damaged versus unaffected glomeruli (**Figure 5B**). Of note, ISG expression was significantly higher in tubules close to glomeruli versus far from glomeruli, regardless of their histologic damage state (**Figure 5B**). We next performed differential gene expression analysis comparing periglomerular tubules next to damaged versus unaffected glomeruli and then performed pathway analysis on the differentially expressed genes. While there were no upregulated pathways that reached statistical significance, early complement pathways and extracellular matrix organization were over-represented. Indeed, several early complement genes were more highly expressed in periglomerular tubules next to damaged glomeruli (**Figure 5C**). Of note, there was no difference in *C5AR1* expression in these two periglomerular tubules populations (**Figure S2A**). Interestingly, *POSTN* which encodes periostin, a protein with a described role in renal fibrosis (20–22) was more highly expressed in periglomerular tubules next to damaged glomeruli (**Figure 5D**). We next performed Pearson Correlation analysis to assess for genes correlated with *POSTN* in the tubulointerstitial compartment and the most highly correlated gene was *LGALS1* (**Figure 5E**). *LGALS1* encodes galectin, a protein that is upregulated in diabetic nephropathy (23) and was upregulated in podocytes in pediatric lupus nephritis as part of a module potentially linked with tubulointerstitial fibrosis (24). Thus, periglomerular tubules next to histologically damaged glomeruli have transcriptional distinctions from those next to unaffected glomeruli, namely increased early complement gene expression and upregulation of some fibrosis-related genes. These observations suggest crosstalk between the glomerular and tubulointerstitial compartments in disease pathogenesis.

**Figure 5.**
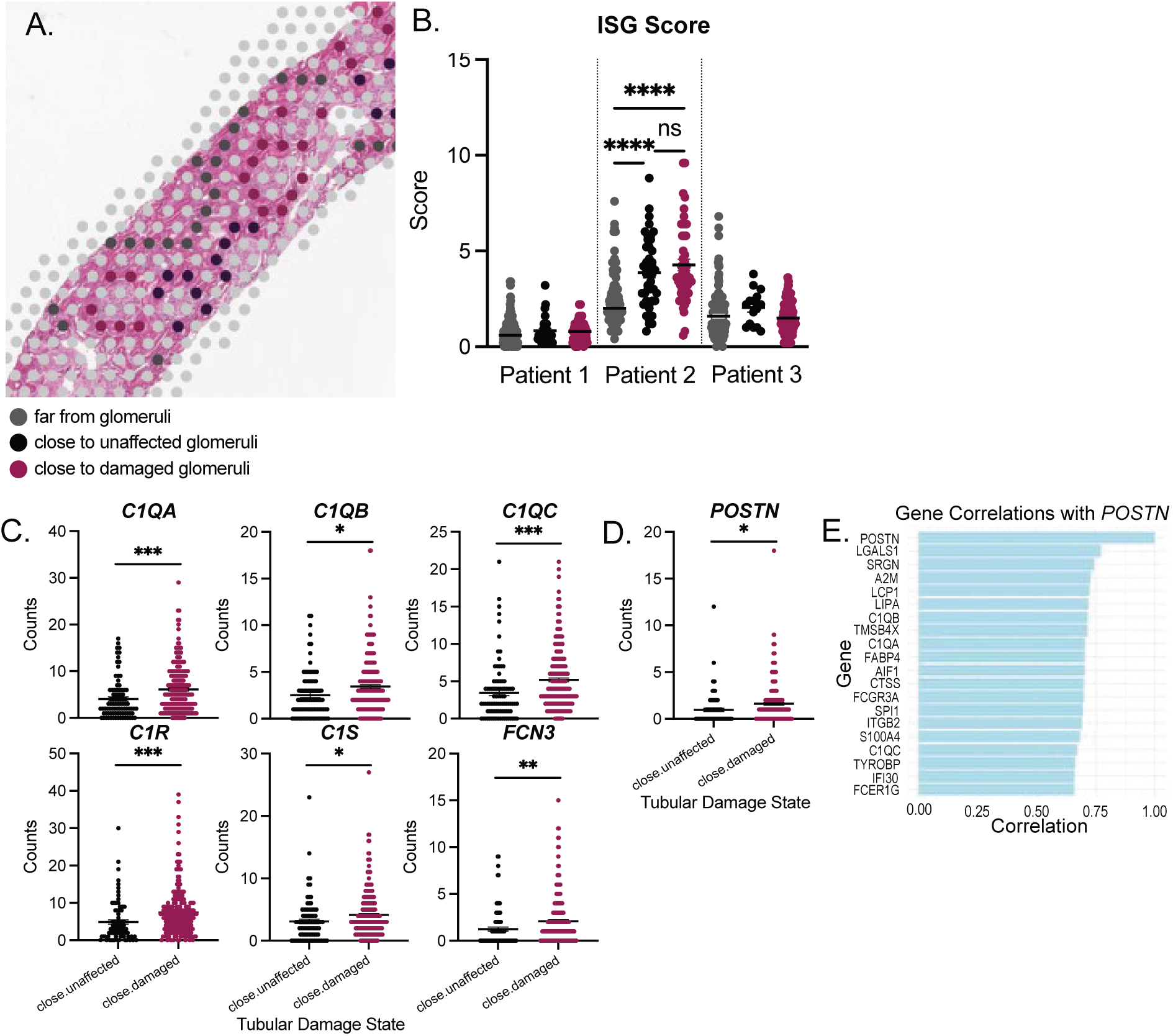
Periglomerular tubules next to damaged Class III LN glomeruli show increased early complement and fibrosis gene expression. **(A)** Representative H&E section from Class III LN biopsy with overlay of 55uM spots showing annotation of tubules that are periglomerular (black = close to unaffected glomeruli, pink = close to damaged glomeruli) or far from glomeruli (grey). **(B)** ISG score computed as the average of the expression of *IFI44, IFI44L, MX1, IFIT1, HERC6*. Each dot is expression from a 55uM spot. X axis numbers represent individual patients. **** p< 0.0001 One-way ANOVA comparing tubules close to damaged, close to unaffected, and far from glomeruli. **(C)** Complement gene expression in periglomerular tubules close to damaged and unaffected glomeruli. (D) *POSTN* expression in periglomerular tubules. Each dot represents expression from a 55μM spot. N=3 pSLE Class III LN patients. P values are unpaired T-tests. ****p<0.0001, ***p<0.001, **p<0.01, *p<0.05. **(E)** Pearson correlation analysis showing top genes correlated with *POSTN*.

### C5a signaling through C5AR1 directly induces fibrosis transcriptional pathways in human macrophages

Thus far, our spatial transcriptomic analysis has revealed microanatomic patterning of early complement and fibrosis-related gene expression in histologically damaged glomeruli and periglomerular tubules next to damaged glomeruli in Class III LN. Furthermore, the druggable target, *C5AR1* is more highly expressed in damaged glomeruli. C5a is a prototypic myeloid chemoattractant, thus a plausible mechanism by which increased *C5AR1* expression may perpetuate glomerular damage is by promoting further myeloid cell recruitment to a damaged glomerulus. However, signaling through C5aR1 likely has other transcriptional effects. Given C5aR1 is most abundantly expressed on myeloid cells in Class III LN kidneys, we next explored the transcriptional changes induced by direct C5a stimulation in human macrophages. Primary human monocytes were obtained from 3 independent adult healthy donors (see **Methods**) and differentiated into macrophages for 6 days with M-CSF. On day 6, macrophages were almost universally C5aR1 positive (**Figure 6A**). Day 6 macrophages were stimulated for 24h with C5a with or without avacopan and RNA was extracted. qPCR analysis of *CXCL5*, a known-C5a-inducible gene, confirmed the C5a stimulation and avacopan blockade was successful for each donor (**Figure 6B**). qPCR for several other genes of interest was also performed. C5a induces *C5AR1* expression and robustly induces the fibrosis-related gene *FN1* that was upregulated in histologically damaged Class III LN glomeruli (**Figure 6B**). Interestingly, C5a downregulates early complement component gene expression (**Figure 6B**). Given this, we tested whether *C5AR1^+^* lupus kidney myeloid cells express lower amounts of *C1Q* components than *C5AR1*^-^ lupus kidney myeloid cells from the AMP dataset (16). We focused on CM2 myeloid cells which are roughly 50% *C5AR1^+^*. In contrast to the results of our stimulation assay, we found that *C5AR1^+^*CM2 myeloid cells from lupus nephritis kidneys expressed higher amounts of *C1QA,B* than *C5AR1^-^*cells (**Figure S3A**) likely owing to complex interacting inflammatory signals in the lupus kidney microenvironment.

**Figure 6.**
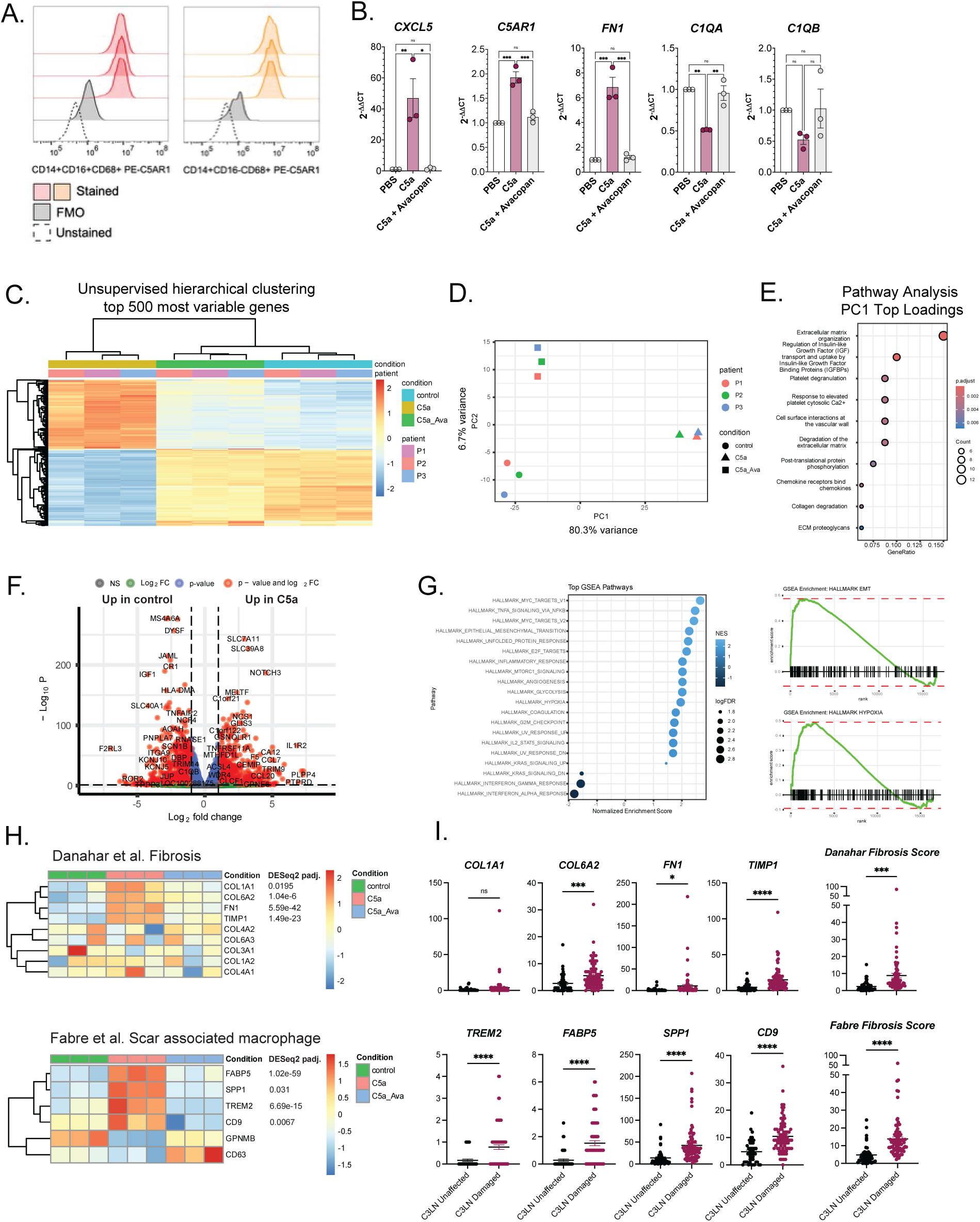
C5a signaling through C5aR1 induces fibrosis transcriptional programs in human macrophages and C5a-induced fibrosis gene expression is increased in histologically damaged LN glomeruli. **(A)** Human monocytes from the peripheral blood of healthy adult human donors (N=3) were differentiated into macrophages in culture for 6 days. Flow cytometry characterization of Day 6 (D6) monocyte-derived macrophages (MDMs) which are ∼85% CD14^+^CD16^-^ and 15% CD14^+^CD16^+^ both subsets are 100% C5aR1^+^. N=1 representative donor in technical triplicate. **(B)** D6 human MDMs were stimulated for 24h with C5a ± avacopan for 24h. RNA was extracted and qPCR and bulk RNA sequencing was performed. qPCR analysis of *CXCL5* (positive control), *C5AR1*, *FN1*, *C1QA*, *C1QB*. Dots represent the average of technical triplicates for N=3 independent donors. P values are one-way ANOVA with Tukey’s post-test for multiple comparisons. *p<0.0332, **p<0.0021, ***p<0.0002, ****p<0.0001. **(C)** Bulk RNA-sequencing unsupervised hierarchical clustering analysis of the top 500 most variable genes in the dataset following ComBat correction. **(D)** PCA analysis of ComBat corrected bulk RNA-sequencing reads from C5a ± avacopan stimulated monocyte-derived macrophages from 3 independent healthy adult human donors. The percent variance explained by Principal Component 1 (PC1) and Principal Component 2 (PC2) are indicated on the corresponding axis. **(E)** Reactome Pathway analysis of PC1 top loadings. **(F)** DESeq2 differential gene expression analysis comparing C5a to control with patient and condition included in the model. **(G)** Gene set enrichment analysis of the rank ordered list of differentially expressed genes comparing C5a to control. Dot plot of significantly differentially expressed pathways with corresponding enrichment plots for pathways associated with fibrosis (Hallmark EMT Pathway, Hallmark Hypoxia Pathway). **(H)** Heatmap showing expression of genes in the Danahar et al. Fibrosis module (24) and Fabre et al. Scar-associated macrophage module (25) in C5a stimulated versus control human MDMs (N=3 independent donors). DESeq2 adjusted p-values are indicated for the significantly upregulated genes. **(I)** Expression of C5a-induced fibrosis genes in histologically damaged versus unaffected Class III LN glomeruli. Danahar Fibrosis Score and Fabre Fibrosis Score are the average expression of *COL1A1, COL6A2, FN1, TIMP1* and *TREM2, FABP5, SPP1, CD9* respectively. P values are unpaired T-tests.

Next, to assess broad transcriptional changes induced by C5a-C5aR1 signaling in human macrophages, bulk RNA-sequencing was performed. Unsupervised hierarchical clustering of the top 500 most variable genes in the dataset showed expected clustering driven by stimulation condition (**Figure 6C**). PCA analysis revealed that C5a-induced changes drive 80% of the transcriptional variance (**Figure 6D**). Pathway analysis of the top loadings of Principal Component 1 (PC1) (**Figure S4A**) revealed expected upregulation of chemokine pathways but also of fibrosis-related pathways (**Figure 6E**). Interestingly, 6.2% of the transcriptional variance was driven by avacopan (**Figure 6D**) and the top loadings of PC2 were interferon stimulated genes and pathways (**Figure S4B**). Differential gene expression analysis comparing C5a stimulation versus control identified a large C5a-induced transcriptional program (**Figure 6F, Supplemental Table S4**). To deconvolute this list of differentially expressed genes, we performed gene set enrichment analysis (GSEA). Among the most highly upregulated pathways (GSEA Hallmark Pathways) were the EMT and Hypoxia pathways which are the Hallmark Pathways with the greatest number of fibrosis-related genes (**Figure 6G**). Interestingly, there was negative enrichment for interferon response pathways (**Figure 6G**). C5a modestly induces expression of the prototypic pro-fibrotic cytokine *TGFB1* itself (**Figure S5A**). However, to explore whether genes that have previously been associated with renal fibrosis in lupus nephritis were upregulated by C5a stimulation in macrophages, we utilized the fibrosis signature identified by Danahar et al. that was enriched in a spatial analysis of pediatric lupus nephritis kidney biopsies (24). 4/9 genes were significantly induced by C5a stimulation in human macrophages (**Figure 6H**). Fabre et al. identified a broadly fibrogenic macrophage subset in human lung and liver that they defined as scar-associated macrophages (25). We found that 4/6 of these genes were significantly upregulated by macrophage C5a-stimulation (**Figure 6H**). To confirm that the upregulation of these fibrosis genes is C5a-specific and not simply a general response to an inflammatory stimulus, we assessed a publicly available RNA-sequencing dataset of LPS-stimulated human monocyte-derived macrophages (26) for induction of the above fibrosis-genes. None of these genes were significantly induced by LPS stimulation (**Figure S5B**). Since C5aR1 is a G-protein coupled receptor (GPCR), we also wanted to ensure that the induction of fibrosis-related genes was not a general phenomenon of GPCR signaling in macrophages. We assessed a publicly available RNA-sequencing dataset of human monocyte-derived macrophages stimulated with prostaglandin E2 (PGE2) for 3h and 24h for fibrosis gene expression (27). Only TIMP1 was significantly upregulated by PGE2 after 3h (**Figure S5C**). Therefore, it seems that the induction of these fibrosis genes in macrophages is C5a-specific.

### C5a-inducible fibrosis genes are upgregulated in histologically damaged Class III LN glomeruli

We next returned to our spatial transcriptomic data and compared the expression of the C5a-inducible genes in the Danahar LN fibrosis signature and Fabre scar-associated macrophage fibrosis signature in histologically damaged versus unaffected Class III LN glomeruli. These C5a-inducible fibrosis genes were significantly upregulated in damaged Class III LN glomeruli (**Figure 6I**). Though we cannot definitively determine if these fibrosis genes were upregulated by C5a in our spatial data, we explored whether the overlap in the genes upregulated by C5a stimulation in macrophages and those upregulated in damaged Class III LN glomeruli is more than what would be expected by chance. Indeed, Fisher’s exact test revealed more significant overrepresentation of the C5a-stimulated macrophage genes in damaged Class III LN glomeruli than what would be expected by chance (p = 0.0038, OR 1.41). Taken together, these data suggest a possible mechanistic link between C5a-C5aR1 signaling and the upregulation of fibrosis-related pathways seen at the time of diagnostic biopsy in Class III LN.

## Discussion

Using spatial transcriptomic analysis, we have identified a transcriptional module associated with histologically damaged glomeruli in Class III pediatric lupus nephritis. This module is defined by higher expression of myeloid cell markers, *C5AR1*, early complement components, and fibrosis genes as compared to histologically unaffected glomeruli. C5aR1 is most abundantly expressed on myeloid cells in the LN kidneys. Direct C5a stimulation of human monocyte derived macrophages upregulates tissue-remodeling and fibrosis-related pathways. These C5a-inducible fibrosis genes are significantly upregulated in histologically damaged versus unaffected glomeruli providing a potential mechanistic link between C5a-C5aR1 signaling and early fibrosis in proliferative lupus nephritis. Taken together, these data make C5aR1 a compelling therapeutic target in proliferative LN not only to curtail myeloid chemotaxis, but also to potentially constrain fibrosis.

We utilized spatial transcriptomics to test an initial hypothesis that the Type I interferon response was elevated in histologically damaged compared to unaffected glomeruli in Class III LN. We demonstrated that ISG expression does not differ between damaged and unaffected Class III LN glomeruli. Along the same lines, it was recently shown that glomeruli from patients without LN have similar ISG expression to those with LN (28). This contrasts with what has been observed in the skin of patients with cutaneous lupus erythematosus (CLE) in which lesional skin had increased ISG expression compared to non-lesional skin (29) and may explain why interferon alpha receptor blockade with anifrolumab has not shown promise in lupus nephritis (10) but is highly efficacious for CLE (30, 31). However, similar to skin (29) histologically unaffected glomeruli had higher ISG expression than healthy control glomeruli suggesting a general state of high interferon activity. Thus, Type I IFNs are likely important upstream drivers in LN but the role of interferon signaling in causing tissue damage at the level of the glomerulus may be less direct. In our small cohort of Class III LN patients, there was heterogeneity between patients in glomerular ISG expression in general. These renal biopsies were all obtained at the time of lupus diagnosis, however we note that patients had differing lengths of clinical symptoms prior to presenting to care. Thus, disease duration prior to diagnosis may influence general interferon activity levels in the tissue as could other variables such as intercurrent viral infection.

Myeloid cell infiltration into glomeruli is a defining feature of proliferative glomerulonephritis in general and often correlates with disease activity (32). We unsurprisingly demonstrated that histologically damaged Class III LN glomeruli have increased expression of myeloid cell markers compared to unaffected glomeruli. However, even histologically unaffected Class III LN glomeruli have increased myeloid cell marker expression compared to healthy control glomeruli or glomeruli from patients with Class V LN. One of the distinctions between proliferative (Class III/IV) and Class V LN in the ISN/RPS classification system is endocapillary hypercellularity and myeloid cell infiltration (11, 12), moreover previous studies have shown by immunohistochemical analysis that among the ISN/RPS classes, CD68 staining is lowest in Class V (33). Thus, this aspect of our spatial transcriptomic analysis confirms known LN pathophysiology. Interestingly, we found that histologically damaged Class III LN glomeruli are enriched in local expression of early complement component genes (*C1QA,B,C*) relative to unaffected Class III LN glomeruli and to Class V LN glomeruli. Based on our analysis of scRNA-seq data from LN kidneys in the AMP dataset, we suspect that this signal is predominately coming from myeloid cells. When Arazi *et al.* defined myeloid cell subsets (CM0-4) from this AMP scRNA-seq data, they noted that cells in CM1 and CM4 had upregulation of *C1Q* (16). Based on trajectory analysis, they hypothesized that the CM1 and CM4 myeloid subsets are derived from inflammatory blood monocytes (CM0) that then differentiate into phagocytic (CM1) and alternatively activated (CM4) macrophages in the kidney (16). Prior spatial transcriptomic analysis of pediatric LN kidneys also revealed a spatially correlated gene module enriched in *CD68, CD163,* and *C1QA,B,C* expression (24). While the classic pathophysiologic paradigm of lupus nephritis involves immune complex deposition in the kidneys with immune complexes containing IgG, IgA, IgM, C3, and C1Q, the immune complex components were thought to be synthesized in the liver, circulate in the blood and deposit in the kidneys. The role of local complement production is less well understood and the pathophysiology of complement in lupus is complex. Deficiencies in early components of the classical complement pathway, including C1q, are some of the strongest genetic drivers of lupus (34). By comparison, deficiencies in terminal complement components are not commonly associated with lupus. Early classical pathway deficiencies likely drive a break in tolerance systemically. However, local C1q synthesis may exacerbate disease by local activation of the classical complement pathway facilitating the engulfment of apoptotic renal cells and enhancing autoantigen presentation but also by recruiting more myeloid cells into the glomerulus through C5a-C5aR1 signaling to perpetuate the local tissue inflammation and injury.

Others have previously demonstrated that *C5AR1* transcription is higher in LN compared to healthy control glomeruli in the Berthier Lupus Glom dataset (35). Interestingly, we found that *C5AR1* was more highly expressed in histologically damaged versus unaffected Class III LN glomeruli. Based on our analysis of the AMP scRNA-seq from digested LN kidneys, *C5AR1* was most abundantly expressed on myeloid cells, with the caveat that resident renal cell types were underrepresented in these data. Immunofluorescence analysis of C5aR1 in pediatric LN kidneys confirmed that expression was mostly colocalized with CD68^+^ myeloid cells, but that other renal resident cells types (CD31^+^ endothelial cells, aSMA^+^ mesangial cells) may also express C5aR1. Given its known myeloid chemotactic role, we hypothesize that the C5a-C5aR1 axis contributes to proliferative LN pathogenesis through this mechanism. Indeed, prior studies in spontaneous murine lupus models have shown that genetic deficiency of C5aR1 in the MRL/*lpr* model prolonged overall survival and markedly attenuated renal disease with reduced glomerular hypercellularity and absence of glomerular crescents (36). Similarly, treatment of MRL/*lpr* with a small molecule inhibitor of C5aR1 starting at 13 weeks of age, constrained development of renal disease and markedly reduced neutrophil and macrophage infiltration into the kidneys (37). Treatment of NZB/W F1 mice with a C5 monoclonal antibody that blocks C5 cleavage starting at 18 weeks of age ameliorated glomerulonephritis and improved survival (38). In these spontaneous murine models, systemic autoantibody production was not significantly altered by C5 or C5AR1 blockade or genetic deficiency. Others have described a C5a-induced podocyte injury mechanism in LN (35). However, the precise mechanism by which C5a-C5aR1 contributes to LN pathogenesis remains unclear. We are performing ongoing studies in murine models using cell-type specific deletions of *C5AR1* to understand the contribution of the C5a-C5aR1 axis in renal resident and immune cell types to LN pathogenesis.

Among the most striking transcriptional differences between histologically damaged and unaffected glomeruli in Class III LN was the increased expression of fibrosis related genes and pathways. Fibrosis is the final common pathway in LN that leads to end stage renal disease and need for renal replacement therapy or transplant (39). Given that the biopsies examined were taken at the time of lupus diagnosis, this suggests that fibrosis programs are already underway when we begin to treat our patients and before there is histologic evidence of fibrosis. This early upregulation of fibrosis-related pathways with recent onset disease has been observed in spatial transcriptomic analysis of an independent cohort of pediatric LN kidney biopsies (24). This is of particular importance in pediatric LN which tends to be more aggressive at onset than in adult LN and suggests that in addition to immunosuppressive therapy, targeting fibrosis pathways early may be critically important.

To our surprise, the two main signals that distinguish histologically damaged and unaffected Class III LN glomeruli, complement and fibrosis, may be mechanistically connected. We convincingly demonstrate that direct C5a stimulation of human macrophages upregulates fibrosis genes and pathways and this is reversed by C5aR1 blockade with avacopan. The C5a-inducible fibrosis genes are overexpressed in histologically damaged compared to unaffected Class III LN glomeruli suggesting a potential mechanistic link. The pro-fibrotic properties of C5a have been described in other disease contexts (idiopathic pulmonary fibrosis (40), pancreatic fibrosis (41), liver fibrosis in hepatitis B infection (42)). In the kidney, C5a inhibition reduced tubulointerstitial fibrosis in a murine model of diabetic nephropathy (43) and genetic deletion of C5aR1 in the unilateral ureteral obstruction murine model of fibrosis significantly reduced renal fibrosis (44). However, the link between C5a signaling and fibrosis is underappreciated in LN. Here, we have uncovered a potential strategy to target fibrosis by inhibiting signaling through C5a-C5aR1.

Our study has several limitations. Given the known heterogeneity in lupus nephritis, the small patient sample size in each disease group may limit the generalizability of these results. We have attempted to control for patient level differences by using linear mixed effects models to confirm gene expression differences. Moreover, we are underpowered to assess differences between LN and non-SLE GN and observations we have made between these diseases are strictly descriptive. We acknowledge that some of the transcriptional changes we have observed in histologically damaged Class III LN glomeruli may be a readout of changes in cell type abundance, of myeloid cells in particular. Given lack of single cell resolution with the spatial transcriptomic technology used for this analysis, we cannot precisely control for cell number. Nonetheless, transcriptional changes driven by changes in cell type abundance are mechanistically important in disease pathogenesis. Lack of single cell resolution also precludes dissection of cell-type contributions to transcriptional signals, for example *C5AR1*. Our ongoing work in an expanded cohort of pediatric LN patients using spatial transcriptomic technology with single cell resolution will allow us to resolve cell-type specific expression of the transcriptional module upregulated in damaged Class III LN glomeruli. We acknowledge that the in vitro system we used to broadly explore C5a-induced transcriptional changes may be imperfect in two primary ways. First, macrophages were derived from peripheral blood monocytes rather than intrarenal myeloid cells and second, the cells came from healthy donors and not lupus patients and thus we cannot account for tissue-specific or disease-specific differences in C5a-signaling in macrophages.

Despite these limitations, we have made biologically meaningful insights into local drivers of glomerular injury in pediatric lupus nephritis and have uncovered an underappreciated link between terminal complement signaling and fibrosis in lupus nephritis with opportunity for therapeutic intervention. These data pave the way for future mechanistic and interventional studies of the C5aR1-fibrosis axis in SLE and other diseases of complement activation. Furthermore, we provide the entirety of our spatial transcriptomic data in a readily accessible, interactive format that can be viewed with the freely available Loupe Browser (45) with the intention for this dataset to be a resource to the community for further interrogation.

## Methods

### Sex as a biological variable

This study was performed on archived tissue samples from renal biopsy specimens of patients with pSLE. All pSLE patients were female owing to the known female-predominance. Healthy control kidney tissue came from pediatric patients who had nephrectomies for non-immune indications were a mix of male and female patients. Disease control tissue came from patients with post-infectious glomerulonephritis and C3 glomerulopathy and incidentally were male patients. Sex was not considered as a biological variable.

### Tissue acquisition

Archived pre-treatment formalin-fixed paraffin-embedded (FFPE) and frozen renal biopsy and resection specimens were obtained for clinical diagnostic purposes at the Children’s Hospital of Philadelphia. Healthy human peripheral blood monocytes were obtained from healthy adult donors through the Human Immunology Core at the University of Pennsylvania.

### Spatial Transcriptomics

Sections were cut from archived FFPE specimens at a 5μM thickness and mounted onto Visium Gene Expression Slides (10x Genomics). Slides were deparaffinized and rehydrated and then stained with Hematoxylin and Eosin (H&E). High resolution brightfield images were obtained using an Aperio AT2 Slide Scanner (Leica Biosystems). Spatial gene expression libraries were generated using the Visium Spatial Gene Expression for FFPE v1 chemistry according to the manufacturers protocol. Briefly, following permeabilization, RNA-templated ligation probes targeting 18,530 genes in the human transcriptome (Visium Human Transcriptome Probe Set v2.0) were hybridized to the tissue. Following ligation, probes were extended to incorporate spatial barcodes and unique molecular identifiers (UMIs). cDNA was recovered and amplified by PCR. Libraries were constructed and sequenced on a NovaSeq6000 (Illumina). Raw sequencing data and high-resolution H&E images were processed using SpaceRanger (v2.1.1, 10x Genomics). The Loupe Browser (v7) in conjunction with SpaceRanger pipeline were used to perform image alignment, tissue detection, and obtain transcript counts using the GRCh38 reference genome. Seurat (v5) was used to perform downstream analysis in R (v4.4.0).

Manual annotation of histologically damaged and unaffected glomeruli in biopsy specimens was performed in Loupe Browser (v8.1.1) by a Pediatric Pathologist. Only 55μM spots that were exclusively glomerular (ie. Did not overlay both glomerular and tubular areas) were annotated. Tubules were annotated in Loupe Browser (v8.1.1) according to the following criteria: tubules “close” to damaged or unaffected glomeruli were the first layer of 55μM tubular spots surrounding a damaged or unaffected glomerulus respectively; tubules “far” from glomeruli were 55μM tubular spots two layers away from the closest glomeruli (independent of glomerular disease state). If a spot satisfied criteria for being “close” and “far” from glomeruli, it was excluded. Barcodes for manually annotated glomeruli and tubules were exported and incorporated as a meta-data layer in the Seurat object for subsequent analysis.

Differential gene expression analysis comparing histologically damaged to unaffected Class III LN glomeruli and Class III to Class V LN glomeruli was performed using the FindMarkers function in Seurat.

### Immune fluorescence staining of frozen renal biopsy specimens

Frozen renal biopsies specimens from pediatric patients with Class III LN (N=2), Class V LN (N=3), post-streptococcal glomerulonephritis (N=1), C3 glomerulopathy (N=1), and control (N=3) were co-immuostained for C5AR1, CD68, PODXL and C5AR1, CD31, αSMA. Briefly, slides were thawed and fixed in 4% paraformaldehyde, permeabilized with Triton X-100, and blocked with 5% goat serum. Slides were incubated with primary antibodies (1:250 C5AR1 clone 8D6 [MA1-70060 Invitrogen], 1:100 CD68 clone KP1 [14-0688-82 Invitrogen], 1:100 PODXL clone E8O1S [55601, Cell Signaling Technology], 1:100 CD31 clone JC/70A [ab9498, Abcam], 1:500 αSMA clone D4K9N [19245, Cell Signaling Technology]) overnight at 4°C overnight. Slides were then washed and incubated with fluorescently-conjugated secondary antibody (1:500 goat anti-rat IgG-Cy3 [112-165-167, Jackson ImmunoResearch], 1:500 goat anti-mouse IgG-Cy5 [ab6563, Abcam], 1:400 goat anti-rabbit IgG 488 [ab150077, Abcam]) for one hour at room temperature. Slides were washed and nuclear stain (DAPI) was added and incubated for 5 minutes. Images were acquired on a Leica DM6000 microscope. Pixel-intensity based colocalization analysis was performed in Fiji (46) and used to assess the overlap in fluorescence signal between C5aR1 and other cell-type markers of interest.

### Human monocyte derived macrophage differentiation, stimulation, and RNA extraction

Density purified human monocytes from the peripheral blood of healthy adult donors (N=3 separate donors processed on different days) were seeded into 48 well tissue culture plates at a density of 500,000 cells per well and cultured in RPMI 1640 with 100ng/mL M-CSF (Peprotech) for 6 days with replacement of 50% of the media with fresh M-CSF-containing media on Day 3 to differentiate into macrophages (MDMs). On Day 6, 100% of the media was replaced with fresh M-CSF-containing media and cells were left untreated or stimulated with 100nM C5a (Peprotech) or 100nM C5a and 1uM avacopan (MedChemExpress) for 24h hours. Media was removed, cells were washed with PBS, and total RNA was extracted using the Qiagen RNeasy Minikit (Qiagen) according to the manufacturers protocol.

### Human monocyte derived macrophage quantitative PCR

cDNA was synthesized cDNA was synthesized using the SuperScript III First-Strand Synthesis System (Invitrogen) according to the manufacturer’s protocol. Real-time quantitative polymerase chain reaction (qPCR) analysis was performed using QuantiTect Primer Assays (Qiagen) and Power SYBR Green Master Mix (ThermoFisher) and run on a StepOne Real Time PCR System (Applied Biosystems). Data were analyzed using the ΔΔC_T_ method. *ACTB* was used as the endogenous reference gene for all qPCR analysis.

### Human monocyte derived macrophage bulk RNA sequencing

Total RNA integrity was assessed using a Bioanalyzer 2100 (Agilent). All samples had an RIN > 9.2. Libraries were prepared using Illumina Stranded mRNA Prep (Illumina) to capture polyadenylated fragments for detection of the protein-coding transcriptome. Final libraries were pooled and sequenced on an Illumina NextSeq2000 platform using paired-end reads to target a depth of 77 million reads per sample. The quality of raw FASTQ files was analyzed using DRAGEN FastQC (Version 4.3, Illumina 2024). Adapters, low-quality reads trimming and alignment to the human reference genome (GrCh38) were all executed using DRAGEN (Version 4.3). Gene expression quantification was performed using featureCounts. Subsequent analysis was performed in R (Version 4.4.1). Counts were Combat corrected to account for patient-level differences for clustering visualization. Unsupervised hierarchical clustering analysis of the top 500 most variable genes and PCA analysis was performed on Combat-corrected counts. Reactome pathway analysis was performed on the top loadings for PC1 and PC2. Differential genes expression analysis was performed using DESeq2 with patient and condition included in the model (design = ∼ patient + condition) to compare C5a stimulation to control. Genes were considered significantly differentially expressed if log_2_fold change > 1 and Benjamini-Hochberg adjusted p value <0.05. Gene set enrichment analysis using the Hallmark pathways was performed on a ranked list of differentially expressed genes comparing C5a stimulation to control.

### Human monocyte derived macrophage flow cytometry

Day 6 human monocyte derived macrophages were surface stained for flow cytometry with the following antibodies to assess for expression of myeloid cell markers and C5aR1 (CD68 [333814, BioLegend], CD14 [562693, BD], CD16 [302010, BioLegend], C5aR1 [344303, BioLegend]). Flow cytometry was performed on a CytoFLEX (Beckman Coulter) instrument. Analysis was performed in FlowJo (BD Biosciences).

### Statistics

In the spatial transcriptomics analyses, unpaired T-tests were used to compare spot level gene expression differences between two groups. To confirm significantly differentially expressed genes between histologically damaged and unaffected Class III LN glomeruli were not due to patient-level differences, gene expression differences were also analyzed using a linear mixed effects model: Fixed effect – glom status (affected/unaffected), Random effects (patient sample). One-way ANOVA with Tukey’s post-test for multiple comparisons were used to compare spot level gene expression differences between more than two groups. Exploratory differential gene expression analysis between two groups (eg. histologically damaged and unaffected Class III LN glomeruli) was performed using FindMarkers within Seurat. Gene expression correlation analyses were performed using Pearson Correlations.

### Study Approval

This spatial transcriptomic study using archived biopsy specimens obtained for clinical diagnostic purposes was approved by the Institutional Review Board (IRB) of the Children’s Hospital of Philadelphia. A waiver of consent was obtained from the IRB for the archived biopsy specimens. Human monocytes were obtained under IRB protocol at the University of Pennsylvania School of Medicine.

## Data Availability

The cloupe files (spatial transcriptomic data overlayed on corresponding H&E image) for each kidney biopsy specimen with the corresponding glomeruli and tubule annotations will be available for download upon publication and can be opened using the publicly available Loupe Browser software. The raw and processed spatial transcriptomic data and MDM bulk RNA sequencing data will be deposited on GEO after publication.

## Author Contributions

SM, EE, SA, JG, JR conducted experiments. SM, DS, EMB performed the computational analysis. SM, EMB designed the project and experiments. PK provided histologic specimens and manually annotated damaged and unaffected glomeruli. SM and EMB drafted the manuscript. All authors reviewed the manuscript and provided comments.

## Acknowledgements

The authors thank Max Eldabbas, Emileigh Maddox, Jiayi Shu, and Tanishk Sinha [as of 7/24/2024, contact us for updates] of the Human Immunology Core at the Perelman School of Medicine at the University of Pennsylvania for assistance with purification of monocytes. The HIC is supported in part by NIH P30 AI045008 and P30 CA016520. HIC RRID: SCR_022380.

The authors thank the Histology Core, Single Cell Technology Core, and High-Throughput Sequencing Core at the Children’s Hospital of Philadelphia for H&E staining of renal biopsy specimens, library preparation, quality control, and sequencing for the renal spatial transcriptomic studies, as well as library preparation, quality control, and RNA-sequencing of C5a stimulated monocyte derived macrophages.

