## Supplemental Figures for "Damaged glomeruli in proliferative pediatric lupus nephritis exhibit a C5a-C5aR1 induced fibrotic transcriptional program"

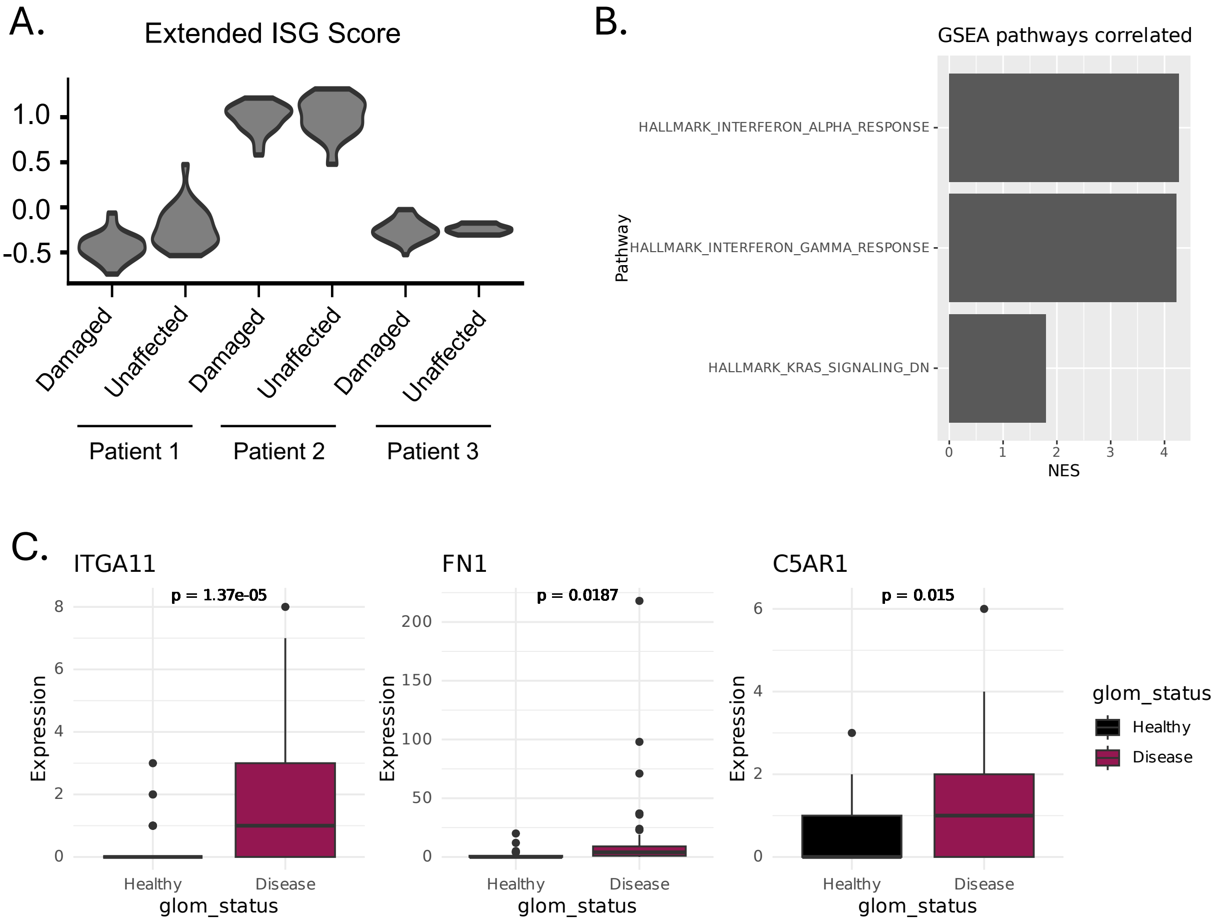


**Supplemental Figure S1 There is no difference in expression of an extended ISG score between histologically damaged and unaffected Class III LN glomeruli but ITGA11, FN1, and C5AR1 expression remain significantly different in a linear mixed effects model**

**(A)** Extended ISG score of all ISGs (Figure 2B) that correlate with core ISGs computed using "AddModuleScore" for patients with Class III LN (N=3). X axis are individual patients comparing histologically damaged (disease) and unaffected (healthy) glomeruli. **(B)** Significantly enriched pathways based on GSEA analysis of a rank-ordered list of genes that correlate with the 5 gene core ISG score (excluding the 5 genes that are components of the score). X-axis is normalized enrichment score (NES). **(C)** *ITGA11, FN1, C5AR1* expression in Class III LN (N=3) in histologically damaged (disease) versus unaffected (healthy) glomeruli assessed using linear mixed effects model: Fixed effect – glom status (affected/unaffected), Random effects (patient sample).

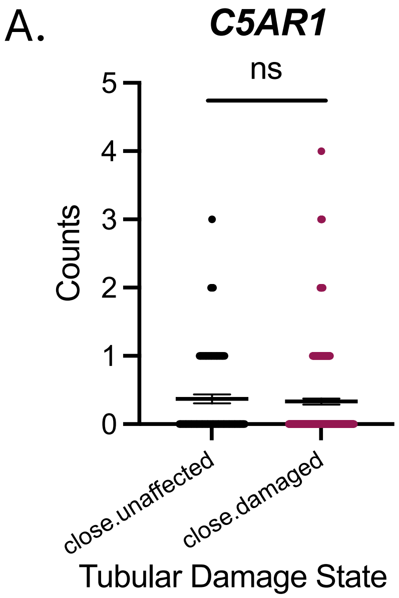


**Supplemental Figure S2 C5AR1 expression is no different between tubules close to histologically damaged versus unaffected Class III LN glomeruli**

**(A)** C5AR1 expression in tubules close to histologically damaged (close.damaged) versus unaffected (close.unaffected) glomeruli from patients with Class III LN (N=3). Each dot is expression from a 55μM spot. P values are unpaired T-tests.

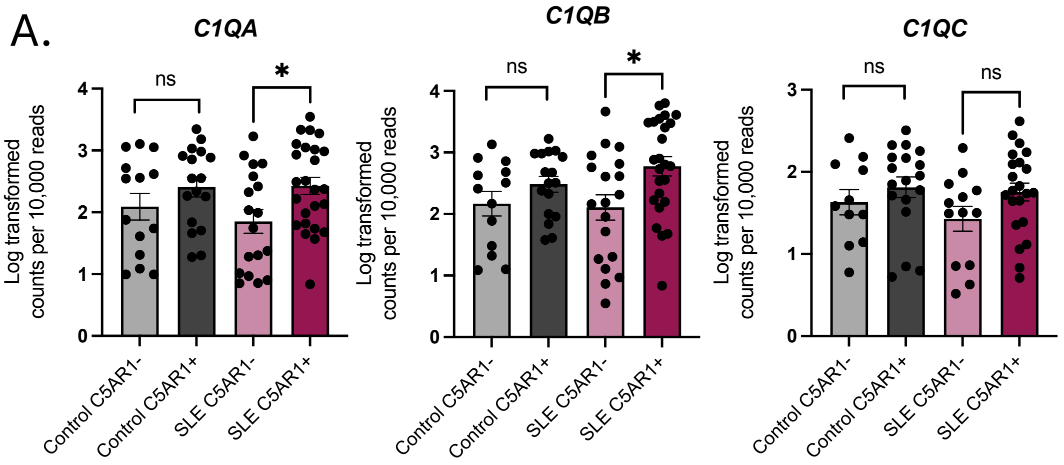


**Supplemental Figure S3 C1q component expression is higher in *C5AR1*$^{+}$ compared to *C5AR1*$^{-}$ myeloid cells in SLE kidneys**

**(A)** *C1QA,B,C* expression in control and SLE kidney CM2 myeloid cells from publicly available AMP data (Arazi et al. 2019). Dots are expression from individual cells. P values are unpaired T-tests between *C5AR1*$^{+}$ and *C5AR1*$^{-}$ cells in control and SLE. *$p<0.05$.

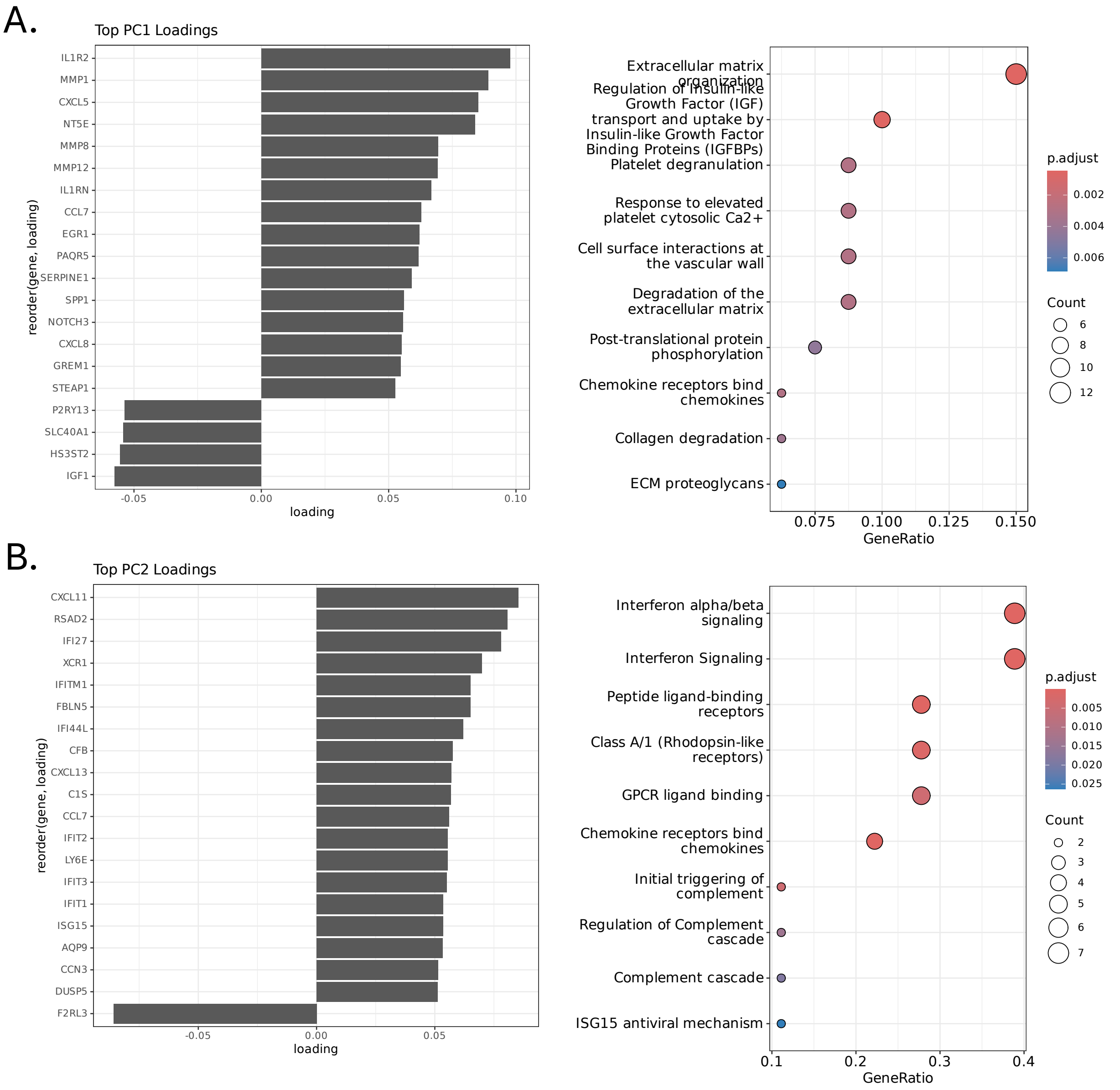


**Supplemental Figure S4 PCA analysis of bulk RNA sequencing from C5a stimulated human monocyte derived macrophages reveals PC1 is driven by chemokine and fibrosis pathways**

**(A)** PC1 top loadings (top 20 genes) [left]. Reactome pathway analysis of PC1 top loadings (top 500 genes) [right]. **(B)** PC2 top loadings (top 20 genes) [left]. Reactome pathway analysis of PC2 top loadings (top 500 genes) [right].

A.

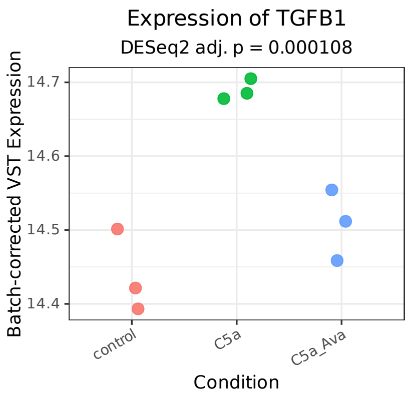


B. Log2 fold change of all LPS-responsive genes in MDMs:

**Positive Controls:**

| Gene | Log2 Fold Change |
|---|---|
| *TNF* | 4.25 |
| *IL6* | 6.81 |

| Gene | Log2 Fold Change |
|---|---|
| *TIMP1* | 0.95 |
| *TREM2* | -0.86 |
| *FABP5* | Not found |
| *SPP1* | Not found |
| *FN1* | Not found |
| *COL6A2* | Not found |
| *TGFB1* | Not found |

C.

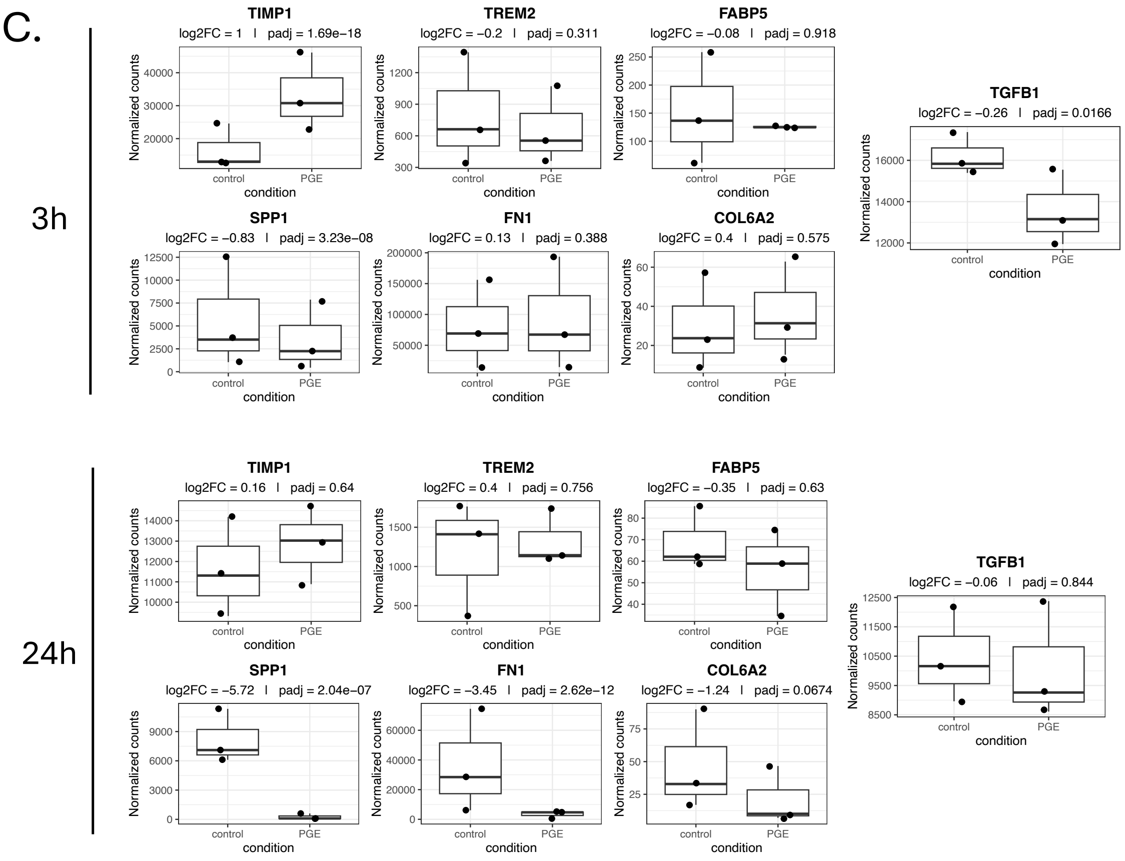

**Supplemental Figure S5 C5a induces expression of TGFB1 in human monocyte-derived macrophages, LPS- and PGE2-stimulated human monocyte-derived do not show upregulation of fibrosis genes induced by C5a.**

**(A)** *TGFB1* expression in human monocyte derived macrophages stimulated with C5a ± avacopan for 24h. N=3 independent donors. Individual dots represent expression from an independent donor. Adjusted p-value from DESeq2 with patient and condition included in the model. VST expression is batch-corrected with patient as batch. **(B)** Log2 fold change of significantly differentially expressed LPS-responsive genes in human monocyte derived macrophages stimulated for 24h with LPS. Genes known to be induced by LPS (*TNF, IL6*) are listed as positive controls [left], C5a-induced fibrosis genes are listed in table [right]. "Not found" indicates that the gene was not significantly differentially expressed. Data derived from Alasoo et al. Immunity 2015; Supplementary Table S3. **(C)** Independent analysis of RNA-sequencing from human monocyte derived macrophages stimulated for 3h or 24h with PGE2 (GSE272019) for induction of C5a-induced fibrosis genes. Raw counts analyzed with DESeq2. Y-axis are DESeq2 normalized counts. Log2FC and adjusted p value from DESeq2 analysis.
