## Supplemental Tables for "Damaged glomeruli in proliferative pediatric lupus nephritis exhibit a C5a-C5aR1 induced fibrotic transcriptional program"

### Supplemental Table S1

Patient Demographic/clinical data

| Disease State (N) | sex = female (%) | age (mean (SD)) | Pre-biopsy immune suppressivetreatment | Time between symptom onset and biopsy (months) |
|---|---|---|---|---|
| Class III LN (3) | 100 | 13.7(2.5) | 10 days of 10mg oral prednisone daily | 1 |
| | | | 1g IV methylprednisolone for 3 days | 5 |
| | | | 1g IV methylpredisolone for 3 days, then 60mg oral prednisone daily and 250mg mycophenolate mofetil twice daily for 3 days | 0.75 |
| Class V LN (3) | 100 | 12.0(4.0) | none | 3 |
| | | | daily hydroxychloroquine, weekly subcutaneous methotrexate, intermittent low dose oral prednisone [illegible] | 36 |
| | | | 60mg oral prednisone daily for 2.5 weeks immediately prior to biopsy | 1 |
| Non-lupus glomerulonephritis (2) | 0 | 10.5(2.1) | none | 0.25 |
| | | | none | 0.25 |
| Healthy Control (3) | 33 | 10 | N/A | N/A |

IV = intravenous

### Supplemental Table S2

**Differential Gene Expression Analysis Class III LN damaged glomeruli vs Class III LN unaffected glomeruli**

Differential gene expression analysis performed using FindMarkers comparing gene expression in 55uM spots from histologically damaged Class III LN glomeruli to histologically unaffected Class III LN glomeruli. For exploratory analysis log2FC>|0.5|, p <0.05 was included in the list of differentially expressed genes.

| | p_val | avg_log2FC | p_val_adj |
|---|---|---|---|
| ITGA11 | 9.41E-09 | 2.1258543 | 0.00017022 |
| NRGN | 2.54E-07 | 1.27926646 | 0.00459101 |
| BBIP1 | 2.84E-07 | 2.16174057 | 0.00513893 |
| NF1 | 3.80E-07 | 1.3227381 | 0.00686861 |
| SLC30A7 | 4.09E-07 | 1.66343679 | 0.00739836 |
| RNF166 | 4.24E-07 | 1.72448193 | 0.00766251 |
| IL10RA | 4.32E-07 | 1.99744322 | 0.00781591 |
| FABP5 | 6.95E-07 | 1.75412797 | 0.01256901 |
| CD68 | 7.26E-07 | 0.96406256 | 0.01313696 |
| FN1 | 7.30E-07 | 2.23272823 | 0.01319692 |
| ANXA6 | 9.43E-07 | 0.99854918 | 0.01705297 |
| TNIP1 | 1.09E-06 | 0.86435683 | 0.01977023 |
| PACS1 | 1.10E-06 | 1.29660538 | 0.01992349 |
| SHTN1 | 1.29E-06 | 1.03028655 | 0.02336664 |
| TCIM | 1.55E-06 | 1.04112148 | 0.02800856 |
| RASGRP3 | 1.88E-06 | 1.24106877 | 0.03395529 |
| KANK2 | 2.08E-06 | 1.23256714 | 0.03762087 |
| COPG1 | 2.18E-06 | 0.93802779 | 0.03937809 |
| MPZL1 | 2.56E-06 | 2.19430911 | 0.04631563 |
| PSMC4 | 3.25E-06 | 0.7309206 | 0.05872008 |
| CARD10 | 3.38E-06 | 1.44874366 | 0.06105282 |
| TLE4 | 3.41E-06 | 1.26127945 | 0.06169667 |
| UBXN6 | 3.82E-06 | 0.8520966 | 0.0690117 |
| NUBP2 | 3.86E-06 | 1.73834329 | 0.06983595 |
| UBE2A | 4.39E-06 | 1.29289843 | 0.07933748 |
| COPE | 4.45E-06 | 1.34333816 | 0.08048958 |
| LAMTOR2 | 4.78E-06 | 1.22323348 | 0.08640514 |
| TEAD4 | 5.37E-06 | 1.91608192 | 0.09705494 |
| COPS3 | 5.41E-06 | 1.30258497 | 0.09781782 |
| CALM1 | 5.49E-06 | 1.67837502 | 0.09930883 |
| CACNB1 | 5.49E-06 | 1.75109205 | 0.09933661 |
| BCL3 | 5.56E-06 | 1.12931722 | 0.10062913 |

| | | | |
|---|---|---|---|
| RAB22A | 5.61E-06 | 1.82825787 | 0.10152449 |
| ALPK3 | 6.28E-06 | 1.15273641 | 0.11351136 |
| KTN1 | 7.88E-06 | 1.04590531 | 0.14245873 |
| UBE2J1 | 9.12E-06 | 0.76944108 | 0.16496367 |
| SH3BP4 | 1.04E-05 | 0.91396347 | 0.18854371 |
| TFEC | 1.04E-05 | 1.64802305 | 0.18867918 |
| CERK | 1.14E-05 | 1.56092026 | 0.20670374 |
| ATP6V1C1 | 1.16E-05 | 0.87707295 | 0.20963741 |
| PPP1R21 | 1.17E-05 | 1.21846699 | 0.21106849 |
| MLH1 | 1.21E-05 | 1.4319823 | 0.21920746 |
| TBL1XR1 | 1.24E-05 | 1.12888047 | 0.22456191 |
| S100A11 | 1.35E-05 | 0.8653586 | 0.24433444 |
| ZNF639 | 1.36E-05 | 1.50156091 | 0.24539016 |
| YKT6 | 1.43E-05 | 1.12431381 | 0.2582105 |
| TGM2 | 1.50E-05 | 1.03171806 | 0.27062905 |
| STX5 | 1.56E-05 | 1.11589745 | 0.28269552 |
| RDH14 | 1.57E-05 | 1.33762594 | 0.28344981 |
| SYNPO2 | 1.59E-05 | 0.96755688 | 0.28720615 |
| MYO9A | 1.60E-05 | 1.05452837 | 0.28922268 |
| YIF1A | 1.60E-05 | 1.32680188 | 0.28938664 |
| TTYH3 | 1.63E-05 | 0.59612355 | 0.29438405 |
| EIF4G1 | 1.66E-05 | 0.63259384 | 0.30110365 |
| DNAJC3 | 1.67E-05 | 0.92592686 | 0.30148744 |
| NDUFA4L2 | 1.67E-05 | 1.76856691 | 0.30266735 |
| SCMH1 | 1.68E-05 | 1.28740048 | 0.30371066 |
| GGT5 | 1.70E-05 | 1.03758082 | 0.30785653 |
| PLBD2 | 1.73E-05 | 0.8928416 | 0.31234034 |
| TCAIM | 1.78E-05 | 0.91843699 | 0.3224912 |
| POLE3 | 1.80E-05 | 0.6946367 | 0.32621018 |
| TAF5L | 1.90E-05 | 2.22362136 | 0.34415938 |
| CXCL16 | 2.03E-05 | 0.91315594 | 0.36741655 |
| POGLUT1 | 2.10E-05 | 0.87066236 | 0.37963716 |
| CLPP | 2.11E-05 | 1.39894982 | 0.38097553 |
| C11orf95 | 2.14E-05 | 0.80348876 | 0.3864866 |
| EFNB2 | 2.16E-05 | 0.77468122 | 0.39039844 |
| GPR4 | 2.18E-05 | 1.3649341 | 0.39399299 |
| ZKSCAN3 | 2.19E-05 | 1.66810933 | 0.3961334 |
| SERTAD2 | 2.19E-05 | 1.18149996 | 0.39664345 |
| SLC35B2 | 2.25E-05 | 1.44946695 | 0.40645131 |
| RINT1 | 2.28E-05 | 1.7109581 | 0.41212751 |
| PSMB3 | 2.33E-05 | 1.62549545 | 0.42208423 |
| LSM14A | 2.61E-05 | 1.35497918 | 0.47223816 |
| SENP6 | 2.62E-05 | 0.57127055 | 0.47400763 |

| | | | |
|---|---|---|---|
| WDR18 | 2.69E-05 | 1.12369816 | 0.4865785 |
| EXOC5 | 2.71E-05 | 1.19713467 | 0.49041138 |
| USP21 | 2.72E-05 | 1.3848097 | 0.49220622 |
| HTATSF1 | 2.76E-05 | 0.88189923 | 0.49955513 |
| DESI2 | 2.81E-05 | 1.19853188 | 0.50751345 |
| ARHGEF3 | 2.95E-05 | 0.85177925 | 0.53320912 |
| USP5 | 2.97E-05 | 0.91416406 | 0.53719759 |
| CPEB2 | 3.04E-05 | 1.08528944 | 0.55008588 |
| PALD1 | 3.04E-05 | 0.83418721 | 0.55036753 |
| FTL | 3.08E-05 | 0.62574159 | 0.55704204 |
| PIEZO1 | 3.48E-05 | 0.68573776 | 0.6296562 |
| TFE3 | 3.57E-05 | 0.73205781 | 0.64482537 |
| FGF1 | 3.59E-05 | 0.70700246 | 0.64903004 |
| NT5DC2 | 3.64E-05 | 1.02147796 | 0.65864836 |
| SEPTIN2 | 3.66E-05 | 0.72077715 | 0.66121863 |
| PPP1R3C | 3.66E-05 | 1.60124038 | 0.6627187 |
| MAPK1 | 3.80E-05 | 0.81214838 | 0.68798148 |
| RECQL | 3.81E-05 | 1.34752926 | 0.68921918 |
| DERA | 3.82E-05 | 1.27437649 | 0.6912184 |
| SF1 | 3.83E-05 | 1.01114289 | 0.6933272 |
| EIF3M | 3.96E-05 | 1.14051027 | 0.71616177 |
| TMEM185A | 4.00E-05 | 0.95909206 | 0.72412325 |
| ITGB1BP1 | 4.05E-05 | 0.75068364 | 0.73190756 |
| SHISA2 | 4.10E-05 | 1.12519883 | 0.74128648 |
| SUPT6H | 4.13E-05 | 0.62556809 | 0.74707454 |
| KCTD15 | 4.14E-05 | 1.19541653 | 0.7481343 |
| MCAM | 4.22E-05 | 1.08126761 | 0.76387097 |
| FOPNL | 4.33E-05 | 1.24797046 | 0.78258598 |
| DCUN1D1 | 4.42E-05 | 1.13386538 | 0.79917608 |
| FAM91A1 | 4.47E-05 | 0.85513885 | 0.80848772 |
| TIMM9 | 4.47E-05 | 1.40785905 | 0.80867878 |
| OXR1 | 4.57E-05 | 1.0719125 | 0.82707764 |
| RIPOR1 | 4.63E-05 | 0.77330718 | 0.83718088 |
| ANXA5 | 4.78E-05 | 0.60718694 | 0.86381289 |
| RAB32 | 4.92E-05 | 1.13281059 | 0.88901759 |
| PDE3A | 4.97E-05 | 1.53079589 | 0.89900526 |
| CCDC88A | 4.99E-05 | 1.10248875 | 0.90284675 |
| ARHGAP1 | 5.18E-05 | 0.91537768 | 0.93735959 |
| MICAL2 | 5.32E-05 | 0.52580251 | 0.96294137 |
| RPP21 | 5.33E-05 | 1.08706259 | 0.96395748 |
| RAB31 | 5.43E-05 | 1.24230107 | 0.98140528 |
| FANCF | 5.51E-05 | 0.55850325 | 0.99602307 |
| CHPF2 | 5.51E-05 | 1.20142884 | 0.99690973 |

| | | | |
|---|---|---|---|
| GPR153 | 0.00012719 | 3.00526594 | 1 |
| MYLK4 | 0.0006503 | 2.83090139 | 1 |
| RPP25 | 0.0006503 | 2.76309925 | 1 |
| SPOCD1 | 0.00210712 | 2.74372787 | 1 |
| FAM71F2 | 0.00142991 | 2.72984057 | 1 |
| CXCL8 | 0.00425563 | 2.69112251 | 1 |
| ARL14EPL | 0.00659759 | 2.64408188 | 1 |
| NLRP3 | 0.00210712 | 2.5792523 | 1 |
| PARS2 | 0.00092476 | 2.56894433 | 1 |
| GDF15 | 0.0006503 | 2.564239 | 1 |
| MTCL1 | 0.00210712 | 2.53546596 | 1 |
| TTLL4 | 0.00015875 | 2.51403476 | 1 |
| LSAMP | 0.00142991 | 2.4337729 | 1 |
| EMB | 0.00452471 | 2.43098684 | 1 |
| TRPV5 | 0.00659759 | 2.40342748 | 1 |
| TRIM29 | 0.00452471 | 2.40120753 | 1 |
| SPATA17 | 0.00142991 | 2.38294789 | 1 |
| FANCG | 0.00339472 | 2.37144673 | 1 |
| RGS4 | 0.02915733 | 2.37001294 | 1 |
| TNFSF8 | 0.00142991 | 2.36505022 | 1 |
| EYA2 | 0.00020077 | 2.35668331 | 1 |
| NLRP11 | 0.00659759 | 2.32711729 | 1 |
| CADM3 | 0.01391671 | 2.30010585 | 1 |
| NAT2 | 0.02015377 | 2.29116316 | 1 |
| ACTR6 | 0.00049694 | 2.28305815 | 1 |
| MOXD1 | 0.00014599 | 2.26160978 | 1 |
| AC132217.2 | 0.00959323 | 2.24371174 | 1 |
| CCNB1 | 0.0014733 | 2.2341744 | 1 |
| NR2C2AP | 0.00111596 | 2.22909717 | 1 |
| IL1RL2 | 0.00959323 | 2.20735634 | 1 |
| YTHDF1 | 0.00163819 | 2.2029943 | 1 |
| B4GALNT1 | 0.00452471 | 2.19727812 | 1 |
| TIAM2 | 0.02015377 | 2.18105688 | 1 |
| TMEM130 | 0.0001605 | 2.17242485 | 1 |
| CCL2 | 9.48E-05 | 2.1295703 | 1 |
| TCTE3 | 0.00141552 | 2.11450834 | 1 |
| C11orf21 | 0.01022648 | 2.10455071 | 1 |
| LMNTD2 | 0.00226895 | 2.08206424 | 1 |
| SMIM17 | 0.00959323 | 2.06864021 | 1 |
| FBXO48 | 0.00339472 | 2.06402518 | 1 |
| TICAM2 | 0.00275092 | 2.05859434 | 1 |
| PCDH19 | 0.00659759 | 2.04696171 | 1 |
| PCDHA1 | 0.01391671 | 2.04389828 | 1 |

| | | | |
|---|---|---|---|
| ZFP82 | 0.00092549 | 2.04063048 | 1 |
| HOXB1 | 0.00959323 | 2.03907067 | 1 |
| ASB12 | 0.00959323 | 2.03552074 | 1 |
| SERPINE1 | 6.93E-05 | 2.03164411 | 1 |
| BRK1 | 0.00134815 | 2.03045711 | 1 |
| C2orf42 | 0.00010972 | 2.02989839 | 1 |
| AURKB | 0.02257486 | 2.02019811 | 1 |
| VTN | 0.00048765 | 2.01750429 | 1 |
| PNO1 | 0.00367855 | 2.0156027 | 1 |
| FMO5 | 0.00016225 | 2.00890702 | 1 |
| SIGLEC7 | 0.00659759 | 1.99767652 | 1 |
| ZNF480 | 0.00925089 | 1.99638759 | 1 |
| GLA | 0.00021618 | 1.97392156 | 1 |
| FMOD | 0.00959323 | 1.96201828 | 1 |
| GFPT2 | 0.00659759 | 1.96080091 | 1 |
| FCAMR | 0.00191522 | 1.95867844 | 1 |
| MAFA | 0.00452471 | 1.94885652 | 1 |
| PHOSPHO1 | 0.00766152 | 1.94711719 | 1 |
| JPH2 | 0.01391671 | 1.94589231 | 1 |
| RGS22 | 0.02015377 | 1.94291825 | 1 |
| NETO2 | 0.0018168 | 1.94036433 | 1 |
| ABHD3 | 0.00010016 | 1.93449863 | 1 |
| CSN2 | 0.02915733 | 1.91513611 | 1 |
| SPDYE1 | 0.02015377 | 1.91393776 | 1 |
| GALNT12 | 0.0421829 | 1.90814939 | 1 |
| LRG1 | 0.01391671 | 1.88792512 | 1 |
| CARD11 | 0.01391671 | 1.87979294 | 1 |
| ACTN2 | 0.0421829 | 1.87931519 | 1 |
| OR13C8 | 0.02015377 | 1.8786875 | 1 |
| PAK3 | 0.0421829 | 1.86682673 | 1 |
| FBXO8 | 0.00022669 | 1.84250531 | 1 |
| SLC51A | 0.00070484 | 1.83577081 | 1 |
| UBE2C | 0.0039487 | 1.83438495 | 1 |
| FAF2 | 0.00016814 | 1.82649519 | 1 |
| CRYBB3 | 0.01391671 | 1.82560939 | 1 |
| PRR18 | 0.02915733 | 1.82493886 | 1 |
| TMCC2 | 0.00200151 | 1.81099808 | 1 |
| PSAPL1 | 0.0421829 | 1.81085685 | 1 |
| KCNA1 | 0.02015377 | 1.80284849 | 1 |
| GLRA1 | 0.02915733 | 1.79993532 | 1 |
| TIGAR | 0.00086076 | 1.79990807 | 1 |
| SLC16A6 | 0.0421829 | 1.79937369 | 1 |
| ULBP2 | 0.0421829 | 1.78898883 | 1 |

| | | | |
|---|---|---|---|
| EXOSC5 | 0.00028662 | 1.78766234 | 1 |
| LRP5L | 0.02015377 | 1.78695886 | 1 |
| CS | 0.02015377 | 1.78687051 | 1 |
| APOC2 | 0.00268594 | 1.77497517 | 1 |
| IKBIP | 0.00016169 | 1.77185861 | 1 |
| LRIT3 | 0.01158956 | 1.76272013 | 1 |
| TAS2R16 | 0.0421829 | 1.75743064 | 1 |
| CLEC2A | 0.02015377 | 1.75511778 | 1 |
| EIF5A2 | 0.00143377 | 1.75357622 | 1 |
| NEURL3 | 0.00618694 | 1.74018111 | 1 |
| FDXACB1 | 0.02015377 | 1.73998801 | 1 |
| STIL | 0.0078646 | 1.73462121 | 1 |
| ENPP6 | 0.00442406 | 1.72995315 | 1 |
| FUNDC1 | 0.00959323 | 1.72328324 | 1 |
| DNMT3B | 0.02015377 | 1.72316643 | 1 |
| TTPA | 0.00450063 | 1.72312045 | 1 |
| KCNAB3 | 0.02915733 | 1.71711886 | 1 |
| ENO3 | 0.02915733 | 1.71618427 | 1 |
| P2RY8 | 0.02915733 | 1.71490954 | 1 |
| MGAM2 | 0.0421829 | 1.71479697 | 1 |
| DOLK | 0.00146143 | 1.71366191 | 1 |
| ST3GAL5 | 0.00140054 | 1.71163152 | 1 |
| SPOCK3 | 0.0421829 | 1.71093906 | 1 |
| SLC10A4 | 0.01391671 | 1.70711825 | 1 |
| TECTA | 0.02915733 | 1.7063249 | 1 |
| OR14K1 | 0.0421829 | 1.70568099 | 1 |
| DDX11 | 0.0096129 | 1.70329517 | 1 |
| MYH7B | 0.01391671 | 1.69401825 | 1 |
| GPR157 | 0.02015377 | 1.69253999 | 1 |
| C21orf58 | 0.02428257 | 1.69171927 | 1 |
| GPR176 | 0.02015377 | 1.68988762 | 1 |
| PTHLH | 0.01391671 | 1.68473825 | 1 |
| NLRP12 | 0.02015377 | 1.68240853 | 1 |
| METTL1 | 0.00068584 | 1.68056896 | 1 |
| SIGLEC8 | 0.0421829 | 1.67728637 | 1 |
| HSPB7 | 0.01085187 | 1.67571194 | 1 |
| ADGRA3 | 0.00390266 | 1.67564211 | 1 |
| YRDC | 0.00036129 | 1.67362118 | 1 |
| NPPB | 0.0421829 | 1.67220386 | 1 |
| SGTB | 0.00018855 | 1.67210856 | 1 |
| HSD17B13 | 0.02915733 | 1.66975271 | 1 |
| SEC14L2 | 0.0186475 | 1.66791447 | 1 |
| VWF | 0.01379913 | 1.66694313 | 1 |

| | | | |
|---|---|---|---|
| CACNA1S | 0.02015377 | 1.66650019 | 1 |
| SPIB | 0.02015377 | 1.6654543 | 1 |
| RPRML | 0.02915733 | 1.66262639 | 1 |
| OR11A1 | 0.02015377 | 1.65928968 | 1 |
| C4orf54 | 0.0421829 | 1.65578071 | 1 |
| LCE4A | 0.0421829 | 1.65508175 | 1 |
| PGK2 | 0.00913715 | 1.65317478 | 1 |
| PMAIP1 | 0.00515936 | 1.64858264 | 1 |
| ASTL | 0.00438804 | 1.64857818 | 1 |
| SLC25A33 | 0.00197869 | 1.64574791 | 1 |
| FBXO40 | 0.02015377 | 1.63618468 | 1 |
| TRPC5OS | 0.02915733 | 1.63571386 | 1 |
| P4HA3 | 0.02015377 | 1.62887435 | 1 |
| SLC12A8 | 0.02195932 | 1.62132115 | 1 |
| C3orf14 | 0.0032634 | 1.61999588 | 1 |
| KCNK3 | 0.00041999 | 1.61668449 | 1 |
| MTNR1A | 0.02015377 | 1.61659775 | 1 |
| OR6C70 | 0.02915733 | 1.61495656 | 1 |
| PRKCQ | 0.04453301 | 1.61125855 | 1 |
| ARNTL | 0.00129845 | 1.60950546 | 1 |
| ACYP1 | 0.00229846 | 1.60534595 | 1 |
| KRTAP19-6 | 0.02915733 | 1.60476745 | 1 |
| MORC4 | 0.00189062 | 1.60248235 | 1 |
| CNTROB | 6.39E-05 | 1.600008 | 1 |
| SLAIN1 | 0.00650526 | 1.5977509 | 1 |
| PSD2 | 0.02015377 | 1.59771801 | 1 |
| ZNF329 | 6.68E-05 | 1.59699151 | 1 |
| CAPN13 | 0.00236672 | 1.59437286 | 1 |
| SPC24 | 0.0421829 | 1.59197783 | 1 |
| ATXN7L2 | 0.00647746 | 1.59084076 | 1 |
| GBX1 | 0.0421829 | 1.58962382 | 1 |
| TEX30 | 0.01824667 | 1.58704559 | 1 |
| ANXA13 | 0.00959323 | 1.58689045 | 1 |
| APH1B | 0.00018847 | 1.58666297 | 1 |
| FCGR2B | 0.02015377 | 1.58286557 | 1 |
| FSIP2 | 0.02915733 | 1.58228729 | 1 |
| UBD | 0.01376814 | 1.58216559 | 1 |
| STAC | 0.0421829 | 1.57933929 | 1 |
| CENPU | 0.01599646 | 1.5792419 | 1 |
| PADI2 | 0.00709552 | 1.57794086 | 1 |
| PF4 | 0.0421829 | 1.57734068 | 1 |
| LBX2 | 0.03625371 | 1.57427577 | 1 |
| OSM | 0.01188077 | 1.5719131 | 1 |

| | | | |
|---|---|---|---|
| NOCT | 0.00895425 | 1.57040836 | 1 |
| CD6 | 0.01391671 | 1.56784464 | 1 |
| MLPH | 0.03625371 | 1.55687527 | 1 |
| RGS16 | 9.86E-05 | 1.55163463 | 1 |
| ALKBH3 | 0.00245863 | 1.5511781 | 1 |
| HTR7 | 0.03484726 | 1.54570144 | 1 |
| STAG3 | 0.04683441 | 1.54305582 | 1 |
| SLC25A43 | 0.00091555 | 1.53627387 | 1 |
| SELP | 0.01241539 | 1.53617532 | 1 |
| OR8H1 | 0.02915733 | 1.53508563 | 1 |
| COX6A2 | 0.02915733 | 1.53064929 | 1 |
| ZNF540 | 0.01077616 | 1.52971468 | 1 |
| RNASE8 | 0.02015377 | 1.51931048 | 1 |
| GSTM1 | 0.02541516 | 1.51662594 | 1 |
| QPCTL | 0.00322048 | 1.5133088 | 1 |
| ECEL1 | 0.0421829 | 1.51196488 | 1 |
| SLC9A1 | 0.00011644 | 1.51141732 | 1 |
| TRPC3 | 0.0421829 | 1.51051378 | 1 |
| FAHD2B | 0.02470451 | 1.50777982 | 1 |
| SLC39A12 | 0.02915733 | 1.50740071 | 1 |
| OR14J1 | 0.02915733 | 1.50700253 | 1 |
| DEFB4A | 0.0421829 | 1.50629959 | 1 |
| FDX1 | 0.00037732 | 1.50283421 | 1 |
| OR6A2 | 0.0421829 | 1.50211137 | 1 |
| SAPCD1 | 0.00098613 | 1.50144635 | 1 |
| TMEM200B | 0.00053809 | 1.50122163 | 1 |
| MALL | 0.03375212 | 1.49991054 | 1 |
| SLAMF1 | 0.01875089 | 1.49843264 | 1 |
| FOXP2 | 0.02915733 | 1.49694606 | 1 |
| CPS1 | 0.01920946 | 1.49637737 | 1 |
| RAG2 | 0.02915733 | 1.48968296 | 1 |
| ATP6AP1L | 0.0096129 | 1.48954046 | 1 |
| CD3E | 0.0421829 | 1.48804202 | 1 |
| POU3F1 | 0.03216767 | 1.48770888 | 1 |
| PDZD7 | 0.01967779 | 1.48754545 | 1 |
| PLK4 | 0.02428257 | 1.48051885 | 1 |
| PRUNE1 | 0.00014049 | 1.47965886 | 1 |
| ORM2 | 0.0421829 | 1.47627042 | 1 |
| 1-Mar | 0.00070233 | 1.47040056 | 1 |
| IFT52 | 0.00297844 | 1.4669121 | 1 |
| LRP12 | 0.00840681 | 1.46644861 | 1 |
| ARL6IP6 | 0.00020151 | 1.46187905 | 1 |
| DDX51 | 0.00096399 | 1.46064643 | 1 |

| | | | |
|---|---|---|---|
| CCL17 | 0.0421829 | 1.45614668 | 1 |
| ITGB7 | 0.02167523 | 1.45117701 | 1 |
| OAZ3 | 0.0421829 | 1.44789527 | 1 |
| MS4A8 | 0.0421829 | 1.44580572 | 1 |
| MAGEB16 | 0.0421829 | 1.4432917 | 1 |
| NUDT14 | 0.00015111 | 1.4409631 | 1 |
| AL162596.1 | 0.0421829 | 1.4404884 | 1 |
| RCOR2 | 0.03800784 | 1.43826127 | 1 |
| TENM3 | 0.00294703 | 1.43806375 | 1 |
| SPDYE6 | 0.0421829 | 1.43764137 | 1 |
| MAGEB5 | 0.0421829 | 1.43464361 | 1 |
| CHCHD3 | 0.00021768 | 1.43321904 | 1 |
| STPG1 | 0.0012422 | 1.4232817 | 1 |
| DERPC | 0.02915733 | 1.42299173 | 1 |
| NAA15 | 0.00011676 | 1.41788076 | 1 |
| EN2 | 0.0421829 | 1.41346061 | 1 |
| ENPP3 | 0.00315003 | 1.4116217 | 1 |
| OR2AE1 | 0.02015377 | 1.41142518 | 1 |
| GPR162 | 0.04683441 | 1.40934307 | 1 |
| ZNF720 | 0.01343628 | 1.40590532 | 1 |
| GSDME | 0.02803628 | 1.40473828 | 1 |
| UNC79 | 0.02015377 | 1.40241802 | 1 |
| TRIM55 | 0.02915733 | 1.40152589 | 1 |
| PSMG1 | 0.00066832 | 1.40063868 | 1 |
| XKR4 | 0.03954676 | 1.40005641 | 1 |
| THAP6 | 0.00118377 | 1.39987195 | 1 |
| OR6C2 | 0.0421829 | 1.39985327 | 1 |
| SSX2IP | 0.00584933 | 1.39980331 | 1 |
| HIST4H4 | 0.0096129 | 1.39873411 | 1 |
| ZFP69 | 0.04683441 | 1.39135016 | 1 |
| ZNF471 | 0.02673295 | 1.38924861 | 1 |
| MCRIP2 | 0.00231179 | 1.38558278 | 1 |
| SRRM5 | 0.0421829 | 1.3851078 | 1 |
| GCA | 0.00132716 | 1.38472013 | 1 |
| EEF1AKMT1 | 0.00030425 | 1.38419505 | 1 |
| CAPN6 | 0.02782939 | 1.38316152 | 1 |
| UTF1 | 0.0421829 | 1.383106 | 1 |
| ADORA2B | 0.0421829 | 1.38286486 | 1 |
| PHLDB3 | 0.00602748 | 1.38279923 | 1 |
| COX8C | 0.0421829 | 1.38252345 | 1 |
| ASIC3 | 0.03800784 | 1.38210454 | 1 |
| DCP1B | 0.00206624 | 1.37814212 | 1 |
| ADAL | 0.00092957 | 1.37530666 | 1 |

| | | | |
|---|---|---|---|
| CEBPA | 0.00037 | 1.37407902 | 1 |
| OR5T3 | 0.02915733 | 1.37298219 | 1 |
| TREM1 | 0.01417525 | 1.36986587 | 1 |
| MND1 | 0.03983363 | 1.36657684 | 1 |
| GCFC2 | 0.00279203 | 1.36522497 | 1 |
| PCGF6 | 0.01121998 | 1.36482182 | 1 |
| NRIP3 | 0.0421829 | 1.3634646 | 1 |
| PDE4B | 0.00033526 | 1.36224778 | 1 |
| IL34 | 0.04245209 | 1.35705922 | 1 |
| AL032819.3 | 0.0421829 | 1.35688776 | 1 |
| C1orf112 | 0.00269607 | 1.35686235 | 1 |
| NSUN5 | 0.04600387 | 1.35683722 | 1 |
| SYT7 | 0.03712195 | 1.35492882 | 1 |
| CFAP44 | 0.00049884 | 1.35266972 | 1 |
| CXCL1 | 0.01445366 | 1.35241091 | 1 |
| THBS2 | 0.00425867 | 1.35230866 | 1 |
| KPNA2 | 0.00045697 | 1.35193564 | 1 |
| ZNF808 | 0.00254018 | 1.35168944 | 1 |
| THAP3 | 0.00051393 | 1.35162821 | 1 |
| SEMA4F | 0.00953331 | 1.35068215 | 1 |
| IL1RAP | 0.00035964 | 1.34979713 | 1 |
| ZNF678 | 0.01147083 | 1.34974121 | 1 |
| ZNF700 | 0.00450728 | 1.34614273 | 1 |
| ATL1 | 0.00168635 | 1.34595329 | 1 |
| GNG8 | 0.02915733 | 1.34332946 | 1 |
| PDE4C | 0.01376814 | 1.34312151 | 1 |
| CCDC113 | 0.0421829 | 1.34283001 | 1 |
| FARS2 | 0.00615078 | 1.34260979 | 1 |
| ZNF382 | 0.02673295 | 1.34259115 | 1 |
| MYH15 | 0.0421829 | 1.34130742 | 1 |
| OCEL1 | 0.00017165 | 1.34065867 | 1 |
| TARS2 | 0.00155468 | 1.33930084 | 1 |
| RHNO1 | 0.00109328 | 1.33831234 | 1 |
| PLAC9 | 0.00455271 | 1.33773621 | 1 |
| CHST7 | 0.0020689 | 1.3352361 | 1 |
| THPO | 0.0421829 | 1.33359892 | 1 |
| TMEM265 | 0.00473369 | 1.33359127 | 1 |
| ZNF331 | 0.00290995 | 1.33296298 | 1 |
| TAAR8 | 0.0421829 | 1.33152735 | 1 |
| ATRNL1 | 0.00132168 | 1.33015546 | 1 |
| MAP2 | 0.00236384 | 1.32988249 | 1 |
| C9orf40 | 0.00898548 | 1.32950691 | 1 |
| PTPN11 | 0.00513592 | 1.32949885 | 1 |

| | | | |
|---|---|---|---|
| DOK3 | 0.00485836 | 1.32915081 | 1 |
| FIGNL2 | 0.00794227 | 1.32857508 | 1 |
| FAM136A | 0.0009068 | 1.32768904 | 1 |
| RNMT | 7.12E-05 | 1.32727729 | 1 |
| C8orf48 | 0.02737792 | 1.32647527 | 1 |
| ADAMTS6 | 0.00203031 | 1.32353846 | 1 |
| PGR | 0.01176128 | 1.31663701 | 1 |
| NFKBID | 0.0007815 | 1.31647753 | 1 |
| STAT4 | 0.04934623 | 1.31612474 | 1 |
| CHD7 | 0.00012623 | 1.31438817 | 1 |
| MCM7 | 5.77E-05 | 1.31194502 | 1 |
| SYNDIG1 | 0.01947232 | 1.30989009 | 1 |
| CCDC89 | 0.00247798 | 1.30746366 | 1 |
| RNF157 | 0.00958501 | 1.30640979 | 1 |
| ZNF679 | 0.0421829 | 1.30570128 | 1 |
| TNNC2 | 0.02618619 | 1.30488119 | 1 |
| ATP8B4 | 0.00550413 | 1.30413542 | 1 |
| HCST | 0.00027063 | 1.30351993 | 1 |
| GUCY1B1 | 0.00017402 | 1.30165907 | 1 |
| SLC30A4 | 0.00034821 | 1.30150715 | 1 |
| CT62 | 0.0421829 | 1.30139602 | 1 |
| IL20 | 0.0421829 | 1.29960379 | 1 |
| RAPSN | 0.03800784 | 1.29911446 | 1 |
| URB2 | 0.02951577 | 1.29874466 | 1 |
| SLC26A3 | 0.0421829 | 1.29848357 | 1 |
| CPN2 | 0.03068822 | 1.29751977 | 1 |
| MOGAT3 | 0.0421829 | 1.29709332 | 1 |
| PSMD11 | 0.00075774 | 1.29557315 | 1 |
| FZD3 | 0.02608777 | 1.29536621 | 1 |
| NDUFAF7 | 7.09E-05 | 1.29348586 | 1 |
| SUB1 | 0.01324841 | 1.29318661 | 1 |
| PSCA | 0.02608777 | 1.29272441 | 1 |
| PYGL | 5.89E-05 | 1.29205124 | 1 |
| SDK1 | 0.01335136 | 1.29153098 | 1 |
| ADAM7 | 0.02915733 | 1.29103899 | 1 |
| CLCN2 | 0.02669996 | 1.28835342 | 1 |
| FMO1 | 0.00553339 | 1.28599213 | 1 |
| FP565260.1 | 0.0421829 | 1.28589195 | 1 |
| DGKD | 0.00138874 | 1.28284494 | 1 |
| RPP40 | 0.01546747 | 1.2819175 | 1 |
| CCNA2 | 0.00753566 | 1.28067331 | 1 |
| ZFYVE19 | 0.00038144 | 1.28052568 | 1 |
| MFNG | 0.00083996 | 1.28002423 | 1 |

| | | | |
|---|---|---|---|
| CCDC192 | 0.0421829 | 1.27785826 | 1 |
| PLGLB2 | 0.00588768 | 1.277453 | 1 |
| PRRG4 | 0.00040907 | 1.27698866 | 1 |
| GIPR | 0.02915733 | 1.27686372 | 1 |
| DCAF8L2 | 0.02915733 | 1.27670492 | 1 |
| CCDC188 | 0.0421829 | 1.27588194 | 1 |
| NUP93 | 0.00034833 | 1.27460801 | 1 |
| DEF6 | 0.00044865 | 1.27401773 | 1 |
| SELL | 0.00216315 | 1.27395952 | 1 |
| LRP8 | 0.02737792 | 1.27373855 | 1 |
| HAUS8 | 0.0421829 | 1.27370906 | 1 |
| RNF122 | 0.00824377 | 1.27279176 | 1 |
| TMEM88B | 0.00341958 | 1.27152014 | 1 |
| RRAS2 | 0.00948742 | 1.2711655 | 1 |
| LEFTY1 | 0.01098605 | 1.27058078 | 1 |
| LMOD1 | 0.00553157 | 1.26813352 | 1 |
| CNTN3 | 0.00015589 | 1.26660498 | 1 |
| CCDC90B | 0.00155008 | 1.26602653 | 1 |
| ART1 | 0.02803628 | 1.26452423 | 1 |
| CLEC18A | 0.00320193 | 1.26427622 | 1 |
| ZNF484 | 0.00210709 | 1.26384716 | 1 |
| CENPE | 0.02673295 | 1.26201867 | 1 |
| ADCK5 | 0.00297705 | 1.26201211 | 1 |
| WDR89 | 0.00156358 | 1.26122203 | 1 |
| APOC1 | 0.04063991 | 1.25990709 | 1 |
| TRIL | 0.00242766 | 1.25945239 | 1 |
| TMEM71 | 0.00776601 | 1.25940405 | 1 |
| KIF7 | 0.00315303 | 1.25913231 | 1 |
| ZNF576 | 0.00191741 | 1.25825104 | 1 |
| FDPS | 0.0001644 | 1.2573756 | 1 |
| NGLY1 | 0.00074375 | 1.25680288 | 1 |
| ASIC5 | 0.0421829 | 1.25678845 | 1 |
| RMDN1 | 0.00062461 | 1.25550513 | 1 |
| SLC16A4 | 0.0001877 | 1.25483975 | 1 |
| LIN54 | 0.00064351 | 1.25402329 | 1 |
| DMRTA2 | 0.0421829 | 1.25370046 | 1 |
| FOXF2 | 0.0421829 | 1.25169377 | 1 |
| GXYLT1 | 0.01084088 | 1.25051825 | 1 |
| ZNF491 | 0.04077409 | 1.24859782 | 1 |
| SLITRK4 | 0.03068822 | 1.24788674 | 1 |
| RRAGB | 0.00483286 | 1.24464591 | 1 |
| STXBP4 | 0.00917696 | 1.24380156 | 1 |
| HES6 | 0.00561237 | 1.24316048 | 1 |

| | | | |
|---|---|---|---|
| RFK | 0.03983363 | 1.24248343 | 1 |
| TRIM66 | 0.01057288 | 1.24107192 | 1 |
| CPSF1 | 0.00025316 | 1.24045827 | 1 |
| RSL24D1 | 0.02064445 | 1.23925448 | 1 |
| NEDD1 | 0.00036052 | 1.23542036 | 1 |
| KCNIP1 | 0.02608777 | 1.23293418 | 1 |
| COL5A3 | 0.01420235 | 1.22779255 | 1 |
| RFTN2 | 0.00240816 | 1.22696704 | 1 |
| TIMM17B | 0.00025568 | 1.22628185 | 1 |
| ZNF425 | 0.04077409 | 1.22585843 | 1 |
| CTHRC1 | 0.00726423 | 1.22210119 | 1 |
| RGS12 | 9.81E-05 | 1.22142467 | 1 |
| EXOG | 0.02737792 | 1.22133474 | 1 |
| ATP5MD | 0.0117049 | 1.22045621 | 1 |
| DACT3 | 0.04210942 | 1.21938391 | 1 |
| DENND2C | 0.00419459 | 1.21904707 | 1 |
| TCF7 | 0.02015606 | 1.21897349 | 1 |
| FAM161A | 0.03787888 | 1.21790496 | 1 |
| DDOST | 0.00221361 | 1.21713373 | 1 |
| SETD9 | 0.03716833 | 1.21681743 | 1 |
| ADAMTSL3 | 0.03787619 | 1.21490902 | 1 |
| BAG2 | 0.00681733 | 1.21488921 | 1 |
| NEMP2 | 0.0099419 | 1.21473956 | 1 |
| CDKL5 | 0.0014197 | 1.21385525 | 1 |
| HDX | 0.02940135 | 1.21356519 | 1 |
| ANGEL2 | 0.00054809 | 1.21285228 | 1 |
| LRRC34 | 0.02870823 | 1.21092775 | 1 |
| NDUFB4 | 0.00441574 | 1.2108446 | 1 |
| LPCAT4 | 0.0006211 | 1.20727957 | 1 |
| RHBDD3 | 0.01475934 | 1.20655238 | 1 |
| CLIP3 | 8.82E-05 | 1.20609233 | 1 |
| ANAPC10 | 0.04348533 | 1.20590855 | 1 |
| GAS7 | 0.00022754 | 1.20558146 | 1 |
| TLCD5 | 0.01590306 | 1.20440423 | 1 |
| ZNF764 | 0.00895575 | 1.20420889 | 1 |
| RFT1 | 0.00015584 | 1.2038953 | 1 |
| ZNF599 | 0.00111786 | 1.20305569 | 1 |
| FIG4 | 0.01981705 | 1.20247793 | 1 |
| CD52 | 0.00440815 | 1.20075846 | 1 |
| LRFN1 | 0.01445366 | 1.19889685 | 1 |
| MMP16 | 0.02780054 | 1.19837505 | 1 |
| NDC1 | 0.01379913 | 1.19832719 | 1 |
| LAMC3 | 0.01260783 | 1.19658354 | 1 |

| | | | |
|---|---|---|---|
| MED14 | 0.00011635 | 1.19630833 | 1 |
| UBE2F | 0.00868366 | 1.19597647 | 1 |
| HAUS3 | 0.00973362 | 1.19556986 | 1 |
| ADAMTS7 | 0.00270814 | 1.19376985 | 1 |
| MILR1 | 0.00259305 | 1.19334139 | 1 |
| CEP19 | 0.02915733 | 1.19321336 | 1 |
| MPHOSPH9 | 0.00243744 | 1.19275302 | 1 |
| MMP15 | 0.0158063 | 1.19266 | 1 |
| FSTL3 | 0.00471475 | 1.19184639 | 1 |
| SCYL3 | 0.00267674 | 1.19129549 | 1 |
| GOLT1B | 0.00191352 | 1.1906147 | 1 |
| RSL1D1 | 0.00083926 | 1.1896499 | 1 |
| TYR | 0.0421829 | 1.18885714 | 1 |
| MAP6D1 | 0.04077409 | 1.18730194 | 1 |
| MID1 | 0.00186305 | 1.18713335 | 1 |
| ZNRD2 | 0.00015926 | 1.18561417 | 1 |
| IQSEC3 | 0.00190071 | 1.18544522 | 1 |
| KLHL7 | 0.00075614 | 1.18514686 | 1 |
| ARMC7 | 0.00031139 | 1.18458805 | 1 |
| RFX6 | 0.0421829 | 1.18373164 | 1 |
| CP | 0.03379567 | 1.18171944 | 1 |
| OCRL | 9.68E-05 | 1.17850562 | 1 |
| SBNO1 | 0.00012098 | 1.17849968 | 1 |
| PIR | 0.00130343 | 1.17776821 | 1 |
| PUS7L | 0.00063885 | 1.1772374 | 1 |
| MINPP1 | 0.00050345 | 1.17622723 | 1 |
| TMEM74B | 0.01166559 | 1.17344763 | 1 |
| APOM | 9.20E-05 | 1.17118573 | 1 |
| HIVEP3 | 0.01000974 | 1.17038526 | 1 |
| PUSL1 | 0.01148105 | 1.16883691 | 1 |
| GPT2 | 0.00295406 | 1.16866697 | 1 |
| IFITM10 | 0.00670504 | 1.16678987 | 1 |
| MLLT11 | 0.0417333 | 1.16606945 | 1 |
| MAGEF1 | 0.00020044 | 1.16339012 | 1 |
| ZNF514 | 0.00221842 | 1.16317937 | 1 |
| C11orf49 | 7.35E-05 | 1.16289566 | 1 |
| RCC1 | 0.0417333 | 1.16074745 | 1 |
| ZNF451 | 0.0014197 | 1.16054598 | 1 |
| ERICH3 | 0.0421829 | 1.16037709 | 1 |
| MYNN | 0.00012301 | 1.16029574 | 1 |
| RAB7B | 0.00933979 | 1.15844929 | 1 |
| GPBAR1 | 0.00701039 | 1.15824385 | 1 |
| DNAJC22 | 0.00625979 | 1.15781043 | 1 |

| | | | |
|---|---|---|---|
| G6PD | 0.00038965 | 1.15592553 | 1 |
| FASTKD3 | 0.00553339 | 1.15483797 | 1 |
| SPC25 | 0.01410723 | 1.15427133 | 1 |
| ZNF367 | 0.04077409 | 1.1513091 | 1 |
| DKK4 | 0.0421829 | 1.15084058 | 1 |
| CAPN12 | 0.01100345 | 1.15063035 | 1 |
| CASR | 0.01546747 | 1.15001052 | 1 |
| SUCLA2 | 0.00047304 | 1.14938995 | 1 |
| WBP11 | 0.01836751 | 1.14915676 | 1 |
| CERS4 | 0.00089761 | 1.14901091 | 1 |
| SEC24A | 0.00331156 | 1.14818557 | 1 |
| IDO1 | 0.00303354 | 1.14601491 | 1 |
| MEGF6 | 0.00035074 | 1.14592227 | 1 |
| GTF2H3 | 0.03787619 | 1.14300496 | 1 |
| MED30 | 0.00074074 | 1.14130357 | 1 |
| PFDN1 | 0.00036677 | 1.14107707 | 1 |
| PLAU | 0.00558298 | 1.14042866 | 1 |
| GPR1 | 0.0421829 | 1.13962806 | 1 |
| LIMK1 | 0.00027077 | 1.13945672 | 1 |
| XRCC5 | 0.00071052 | 1.13516591 | 1 |
| GNAI1 | 0.00269179 | 1.13477646 | 1 |
| ATL3 | 0.00014296 | 1.13401885 | 1 |
| TRPM4 | 9.37E-05 | 1.13285571 | 1 |
| CEP152 | 0.02204084 | 1.13028382 | 1 |
| MARVELD3 | 0.00156758 | 1.12925948 | 1 |
| GPR25 | 0.0417333 | 1.12824966 | 1 |
| KATNB1 | 0.03140476 | 1.12760558 | 1 |
| CPNE6 | 0.02647229 | 1.12450457 | 1 |
| MEF2C | 0.00032782 | 1.12436231 | 1 |
| FOXD4L5 | 0.00504778 | 1.12399005 | 1 |
| NCAPD2 | 0.00334017 | 1.1233981 | 1 |
| FADS2 | 0.00213005 | 1.12333582 | 1 |
| PPP3CA | 0.00010617 | 1.12221743 | 1 |
| GSTA4 | 0.00640422 | 1.12195437 | 1 |
| KCNN4 | 0.03023443 | 1.12116321 | 1 |
| PDE4A | 0.0009659 | 1.12063151 | 1 |
| RELL2 | 0.00862452 | 1.11854154 | 1 |
| WDFY4 | 0.01887176 | 1.11803028 | 1 |
| NUDT19 | 0.00019384 | 1.1179926 | 1 |
| ST7 | 0.00177784 | 1.11559076 | 1 |
| BRINP1 | 0.0421829 | 1.11456305 | 1 |
| RIPK2 | 0.00286973 | 1.1131956 | 1 |
| MCF2 | 0.04077409 | 1.112779 | 1 |

| | | | |
|---|---|---|---|
| CYB5RL | 0.01387099 | 1.11232541 | 1 |
| AMPH | 0.00415249 | 1.11180394 | 1 |
| THAP7 | 0.00290903 | 1.11175368 | 1 |
| CLIP2 | 0.00076133 | 1.11132868 | 1 |
| SASS6 | 0.00439156 | 1.11076709 | 1 |
| ABHD13 | 0.0028368 | 1.1094786 | 1 |
| PLIN2 | 0.00594317 | 1.10882824 | 1 |
| TOP1MT | 0.00019289 | 1.1076545 | 1 |
| P3H1 | 0.00345488 | 1.1066921 | 1 |
| HEBP2 | 0.00050958 | 1.10619126 | 1 |
| PROCR | 0.01653524 | 1.10400154 | 1 |
| ARHGEF25 | 0.00034845 | 1.10374061 | 1 |
| HOPX | 0.01546747 | 1.10356041 | 1 |
| KCNK13 | 0.00725808 | 1.10347594 | 1 |
| GTPBP8 | 0.0417333 | 1.1008149 | 1 |
| FGD4 | 0.00039486 | 1.1007736 | 1 |
| TMSB15B | 0.03983363 | 1.10041974 | 1 |
| FAM118B | 0.00221361 | 1.10009578 | 1 |
| ZNF10 | 0.02309857 | 1.09931299 | 1 |
| B4GALT2 | 0.00106457 | 1.09807152 | 1 |
| TMEM273 | 0.01760019 | 1.09785181 | 1 |
| SPATA33 | 0.00036728 | 1.09777554 | 1 |
| TFAP4 | 0.00877279 | 1.09725577 | 1 |
| FNDC1 | 0.04077409 | 1.09690501 | 1 |
| IL1RAPL2 | 0.0421829 | 1.09646588 | 1 |
| ANP32B | 0.00292489 | 1.09558789 | 1 |
| ZNF274 | 0.00052322 | 1.09440093 | 1 |
| CALCR | 0.03087076 | 1.09311273 | 1 |
| MCM9 | 0.00759474 | 1.09264229 | 1 |
| IER3 | 7.51E-05 | 1.09249893 | 1 |
| PRSS36 | 0.04077409 | 1.09206832 | 1 |
| PRPF18 | 0.00107848 | 1.08972132 | 1 |
| RNF6 | 0.00044628 | 1.08955606 | 1 |
| PDE7B | 0.00012301 | 1.08707857 | 1 |
| PPP2R5B | 0.00068766 | 1.08444551 | 1 |
| NFS1 | 0.0062752 | 1.0843358 | 1 |
| MTERF1 | 0.04671199 | 1.08431518 | 1 |
| MS4A14 | 0.00436945 | 1.08334376 | 1 |
| C8orf76 | 0.00122287 | 1.0821991 | 1 |
| MED23 | 0.00096486 | 1.0819242 | 1 |
| PLEKHA8 | 0.01037387 | 1.08019253 | 1 |
| ZSWIM4 | 0.00485427 | 1.080094 | 1 |
| STK26 | 0.02775365 | 1.079912 | 1 |

| | | | |
|---|---|---|---|
| IGF2BP1 | 0.0421829 | 1.0791072 | 1 |
| ZNF219 | 0.00177784 | 1.07793802 | 1 |
| FUT11 | 0.00127994 | 1.07598095 | 1 |
| RFX3 | 0.00091511 | 1.07549046 | 1 |
| FAM185A | 0.03710068 | 1.07346462 | 1 |
| TAF6L | 0.00403254 | 1.07340114 | 1 |
| RBPMS2 | 0.00053395 | 1.07314897 | 1 |
| DTD2 | 0.01440839 | 1.07070346 | 1 |
| NOS3 | 0.0016594 | 1.06899322 | 1 |
| CGAS | 0.00299481 | 1.06894085 | 1 |
| TSPAN5 | 0.00301297 | 1.06882261 | 1 |
| KLHL8 | 0.0004183 | 1.06830204 | 1 |
| OR51E2 | 0.02870823 | 1.06811796 | 1 |
| BMI1 | 0.00022544 | 1.06803054 | 1 |
| MYBPH | 0.03103471 | 1.06666833 | 1 |
| LILRB4 | 0.03891164 | 1.06659132 | 1 |
| MTMR6 | 0.00013111 | 1.06400344 | 1 |
| GMPPB | 0.00012304 | 1.06155325 | 1 |
| NOM1 | 0.00309769 | 1.06083613 | 1 |
| PRKAB2 | 0.00825211 | 1.05854275 | 1 |
| RNF139 | 0.00031685 | 1.05626273 | 1 |
| LRRC42 | 0.00166402 | 1.05591755 | 1 |
| TREM2 | 0.00012348 | 1.055517 | 1 |
| LAMA3 | 0.01784279 | 1.05546518 | 1 |
| NUDT8 | 0.02394403 | 1.05450277 | 1 |
| PHLDA2 | 0.02085684 | 1.05439268 | 1 |
| STMN1 | 0.00459688 | 1.05365417 | 1 |
| RCC2 | 0.00977344 | 1.05203117 | 1 |
| FERMT3 | 0.00029744 | 1.05186312 | 1 |
| AIMP1 | 8.36E-05 | 1.05121043 | 1 |
| SEC24D | 0.00130127 | 1.0499749 | 1 |
| PCNX2 | 0.00541167 | 1.04984904 | 1 |
| RUBCNL | 0.04617737 | 1.04843116 | 1 |
| ZNF184 | 0.0074001 | 1.04673307 | 1 |
| FGFR1OP2 | 0.00031669 | 1.0467313 | 1 |
| SCGB2A1 | 0.0421829 | 1.04377597 | 1 |
| CMAS | 0.00055673 | 1.04311361 | 1 |
| AMPD3 | 0.00828943 | 1.0430798 | 1 |
| BRWD3 | 0.00647909 | 1.04300089 | 1 |
| VIPAS39 | 0.00283787 | 1.04049651 | 1 |
| CINP | 0.00544304 | 1.04026571 | 1 |
| RAB3IL1 | 0.00022307 | 1.03791174 | 1 |
| PDZD3 | 0.00030032 | 1.03755393 | 1 |

| | | | |
|---|---|---|---|
| FUT7 | 0.02309857 | 1.03541346 | 1 |
| KCNMA1 | 0.00238054 | 1.03480307 | 1 |
| DNASE2 | 0.00151133 | 1.03393736 | 1 |
| ST3GAL4 | 0.00215889 | 1.03338152 | 1 |
| CNTLN | 0.00055921 | 1.03111999 | 1 |
| LHFPL2 | 0.00194139 | 1.02966599 | 1 |
| ABCC3 | 0.00020953 | 1.02640463 | 1 |
| FOXO1 | 0.00255036 | 1.02551854 | 1 |
| ATF7IP2 | 0.00766485 | 1.02490267 | 1 |
| PCNX3 | 0.00027354 | 1.02428076 | 1 |
| MYCN | 0.0260369 | 1.02387676 | 1 |
| PEBP4 | 0.04552837 | 1.02264069 | 1 |
| METRN | 0.00082174 | 1.02150452 | 1 |
| C15orf39 | 0.00185762 | 1.02049719 | 1 |
| ZDHHC12 | 0.00314469 | 1.01945326 | 1 |
| HACL1 | 0.00398548 | 1.01917396 | 1 |
| KRTAP13-2 | 0.03983363 | 1.01862384 | 1 |
| PDLIM4 | 0.02801587 | 1.01671286 | 1 |
| TMEM43 | 0.00052436 | 1.01665785 | 1 |
| NCAPG2 | 0.03264609 | 1.01520069 | 1 |
| TUBA1C | 0.00225618 | 1.01486969 | 1 |
| PCDHGB6 | 0.00984385 | 1.01367417 | 1 |
| C1GALT1 | 0.01591773 | 1.01251389 | 1 |
| DSPP | 0.0421829 | 1.01140597 | 1 |
| GLT8D1 | 0.00019829 | 1.00993741 | 1 |
| PNPLA1 | 0.03854222 | 1.00950097 | 1 |
| BTBD10 | 0.00023734 | 1.00949957 | 1 |
| ORC3 | 0.0083784 | 1.00935219 | 1 |
| ACTA2 | 0.00247702 | 1.00925242 | 1 |
| ZNF575 | 0.04077409 | 1.00913595 | 1 |
| SERPINI1 | 0.00452075 | 1.00842672 | 1 |
| HACD3 | 0.00059625 | 1.00811898 | 1 |
| SLC35C2 | 0.0015369 | 1.0078937 | 1 |
| SUV39H1 | 0.01656671 | 1.00721894 | 1 |
| RPIA | 0.00065156 | 1.00604149 | 1 |
| TRIM54 | 0.00260495 | 1.00566781 | 1 |
| PGM5 | 0.00082079 | 1.00533812 | 1 |
| PSMG3 | 0.00078358 | 1.00418714 | 1 |
| SNRPG | 0.00213005 | 1.00330379 | 1 |
| CYP2B6 | 0.00679167 | 1.00301316 | 1 |
| ITGB6 | 0.01697705 | 1.00291526 | 1 |
| MOSPD3 | 0.00933979 | 1.00186116 | 1 |
| TBC1D25 | 0.03578636 | 1.00181772 | 1 |

| | | | |
|---|---|---|---|
| LIME1 | 0.00171281 | 1.00110072 | 1 |
| RPUSD1 | 0.01777228 | 1.000899 | 1 |
| SLC2A11 | 0.00102738 | 1.00038244 | 1 |
| OR2F1 | 0.0421829 | 0.99986107 | 1 |
| MSR1 | 0.00038314 | 0.99974045 | 1 |
| MPPED2 | 0.00041144 | 0.99915492 | 1 |
| SMUG1 | 0.001705 | 0.99825956 | 1 |
| HOMER1 | 0.0096732 | 0.99812325 | 1 |
| GLTP | 0.01003801 | 0.99559157 | 1 |
| ABHD14A | 0.00561808 | 0.99449516 | 1 |
| RWDD4 | 0.00934393 | 0.99427128 | 1 |
| THEMIS2 | 0.00056113 | 0.99407513 | 1 |
| SIGLEC1 | 0.0018402 | 0.99368949 | 1 |
| COX20 | 0.00124691 | 0.99359627 | 1 |
| LTBP2 | 0.00934393 | 0.99356249 | 1 |
| FXR2 | 9.69E-05 | 0.99285445 | 1 |
| ABCB7 | 0.0003747 | 0.99101847 | 1 |
| GSTZ1 | 0.04499614 | 0.99092824 | 1 |
| EME2 | 0.00526871 | 0.9889884 | 1 |
| ANKRD37 | 0.02660379 | 0.98874096 | 1 |
| POLR3A | 7.06E-05 | 0.98806769 | 1 |
| SLC23A3 | 0.00970715 | 0.98587186 | 1 |
| SLC26A10 | 0.02870823 | 0.98505421 | 1 |
| MYOZ2 | 0.02103986 | 0.98478532 | 1 |
| TAF1C | 0.00066916 | 0.98433687 | 1 |
| TMEM177 | 0.00696831 | 0.98432484 | 1 |
| HIST1H4C | 0.0012542 | 0.98425855 | 1 |
| NAE1 | 0.00252833 | 0.98244985 | 1 |
| TEAD2 | 0.00036848 | 0.98202852 | 1 |
| NAA40 | 0.00489191 | 0.98176462 | 1 |
| TRIM37 | 0.00092744 | 0.98143601 | 1 |
| HPS3 | 0.00010344 | 0.98090056 | 1 |
| SOX17 | 0.00166753 | 0.98070495 | 1 |
| NCOA5 | 0.01012722 | 0.97990576 | 1 |
| THUMPD1 | 0.00011098 | 0.97981488 | 1 |
| TMEM33 | 0.00077538 | 0.97964123 | 1 |
| NADK2 | 0.00050428 | 0.97880215 | 1 |
| PPP5C | 0.00048964 | 0.97782141 | 1 |
| ADNP | 0.00071463 | 0.97654984 | 1 |
| C8orf88 | 0.00768031 | 0.97600136 | 1 |
| AP4B1 | 0.00171348 | 0.97549807 | 1 |
| PIH1D2 | 0.01260416 | 0.97548305 | 1 |
| P2RX7 | 0.0385382 | 0.97496964 | 1 |

| | | | |
|---|---|---|---|
| ARHGAP44 | 0.00533726 | 0.97357882 | 1 |
| TFAP2B | 0.00751456 | 0.97355959 | 1 |
| NUP155 | 0.00757299 | 0.97321037 | 1 |
| VWDE | 0.04077409 | 0.97255087 | 1 |
| MPV17L2 | 0.03185871 | 0.97212954 | 1 |
| FIGN | 0.00954646 | 0.97150448 | 1 |
| SLC24A3 | 0.04198129 | 0.97088856 | 1 |
| L3MBTL2 | 0.0035456 | 0.97085224 | 1 |
| AVIL | 0.00518194 | 0.97078428 | 1 |
| GOLGA8B | 0.00346247 | 0.96992616 | 1 |
| ABHD12 | 0.00706933 | 0.96867752 | 1 |
| LDLR | 0.00087952 | 0.96816211 | 1 |
| ERBB4 | 0.00063193 | 0.967721 | 1 |
| VCAN | 0.00187808 | 0.96770385 | 1 |
| NAP1L3 | 0.04372446 | 0.96713108 | 1 |
| RBM18 | 0.00212508 | 0.96686429 | 1 |
| TLR6 | 0.00933193 | 0.96682244 | 1 |
| GPRC5A | 0.00067086 | 0.96648354 | 1 |
| JMJD4 | 0.00275731 | 0.96614695 | 1 |
| ALDH16A1 | 0.00026796 | 0.96558623 | 1 |
| TBC1D7 | 0.00503025 | 0.96482424 | 1 |
| ZNF691 | 0.02896324 | 0.96449137 | 1 |
| DRAM1 | 0.00051608 | 0.96349694 | 1 |
| MPLKIP | 0.00023436 | 0.96323266 | 1 |
| SS18L2 | 0.00327665 | 0.96308835 | 1 |
| HPS4 | 0.00233945 | 0.96229762 | 1 |
| ZGLP1 | 0.03054598 | 0.96194117 | 1 |
| TMEM97 | 0.00830557 | 0.96191063 | 1 |
| ATP2A3 | 0.00050003 | 0.96050104 | 1 |
| TCTN3 | 0.0015369 | 0.95979035 | 1 |
| CFAP20 | 0.00010274 | 0.95960927 | 1 |
| BEST1 | 0.01013748 | 0.95934199 | 1 |
| ANTXR1 | 9.33E-05 | 0.95832818 | 1 |
| CEACAM19 | 0.02323393 | 0.95573816 | 1 |
| CFAP46 | 0.00898548 | 0.95561829 | 1 |
| TM6SF1 | 0.03313673 | 0.95539052 | 1 |
| FRAT2 | 0.00378012 | 0.95400077 | 1 |
| CRIP1 | 9.09E-05 | 0.95331303 | 1 |
| ZNF221 | 0.04113199 | 0.95288763 | 1 |
| PROS1 | 0.01196195 | 0.95286941 | 1 |
| SIGLEC11 | 0.02145674 | 0.95172145 | 1 |
| EFNA4 | 0.02939401 | 0.9513263 | 1 |
| C18orf54 | 0.01724429 | 0.95127398 | 1 |

| | | | |
|---|---|---|---|
| FCF1 | 0.00498011 | 0.95105272 | 1 |
| DYRK4 | 0.00572393 | 0.95092744 | 1 |
| TMEM251 | 0.00997643 | 0.94895571 | 1 |
| SPEG | 0.03255053 | 0.94888729 | 1 |
| TMC6 | 0.00108374 | 0.94853017 | 1 |
| AMZ2 | 0.01182955 | 0.9476327 | 1 |
| MTRF1 | 0.00141009 | 0.94700785 | 1 |
| SCNN1D | 0.02744554 | 0.9454798 | 1 |
| RAB5A | 0.0002412 | 0.9453251 | 1 |
| FLVCR1 | 0.04834406 | 0.94416912 | 1 |
| CYP7B1 | 0.0007974 | 0.94406537 | 1 |
| GPC6 | 0.01134523 | 0.943308 | 1 |
| PDE9A | 0.00684627 | 0.94321075 | 1 |
| AXIN2 | 0.01574848 | 0.9421138 | 1 |
| KCNK6 | 0.00181281 | 0.94048306 | 1 |
| HEATR6 | 0.01623685 | 0.9387397 | 1 |
| RENBP | 0.00141551 | 0.93865597 | 1 |
| RRP9 | 0.02491446 | 0.93832575 | 1 |
| SURF4 | 0.00030133 | 0.93830619 | 1 |
| TTC27 | 0.00052142 | 0.93776421 | 1 |
| RCN2 | 0.00052322 | 0.93694207 | 1 |
| FBXO25 | 0.00697475 | 0.93683654 | 1 |
| EDF1 | 0.00270826 | 0.93641417 | 1 |
| ZNF836 | 0.02921669 | 0.9355546 | 1 |
| TOR3A | 0.00222528 | 0.93356574 | 1 |
| CTNNBL1 | 0.00067068 | 0.93321873 | 1 |
| TRAF3IP3 | 0.01642088 | 0.93162589 | 1 |
| MAZ | 0.00059596 | 0.93106328 | 1 |
| RFLNB | 0.00039547 | 0.93079578 | 1 |
| TEP1 | 0.00468788 | 0.92970351 | 1 |
| SMLR1 | 0.0186475 | 0.92869733 | 1 |
| FIZ1 | 0.01852315 | 0.92854021 | 1 |
| IFT140 | 0.01182293 | 0.92757107 | 1 |
| SZT2 | 0.001497 | 0.92685522 | 1 |
| ZSWIM1 | 0.01037201 | 0.92661275 | 1 |
| CREBBP | 0.0002791 | 0.92656471 | 1 |
| SP140L | 0.00183829 | 0.9259844 | 1 |
| ROBO1 | 0.00113081 | 0.92523309 | 1 |
| TRAF3IP2 | 0.00726423 | 0.92508902 | 1 |
| LPCAT2 | 8.58E-05 | 0.92423695 | 1 |
| PARP8 | 0.00030396 | 0.92343502 | 1 |
| SOAT1 | 0.02622314 | 0.92273047 | 1 |
| GABPB1 | 0.01570579 | 0.9215919 | 1 |

| | | | |
|---|---|---|---|
| KLHL23 | 0.00776632 | 0.92150582 | 1 |
| D2HGDH | 0.00445426 | 0.92074141 | 1 |
| GSS | 0.00048964 | 0.92027917 | 1 |
| MAPK10 | 0.00036574 | 0.91972262 | 1 |
| CLMP | 0.04556667 | 0.91943288 | 1 |
| CEP97 | 0.0117121 | 0.91814342 | 1 |
| TIMP1 | 0.00026752 | 0.91810705 | 1 |
| TCF4 | 0.00035083 | 0.91749805 | 1 |
| RETREG3 | 0.00015775 | 0.91666779 | 1 |
| PIGP | 0.01343778 | 0.91639407 | 1 |
| INTS8 | 0.00103331 | 0.91605366 | 1 |
| ELOVL1 | 0.00012427 | 0.9160534 | 1 |
| SMIM20 | 0.00028188 | 0.91584278 | 1 |
| ABHD5 | 0.01319746 | 0.91490462 | 1 |
| NXPE3 | 0.00411574 | 0.91454177 | 1 |
| PAIP2B | 0.00231218 | 0.91453443 | 1 |
| FCGR3B | 0.03252181 | 0.91429862 | 1 |
| TMEM183A | 0.00057939 | 0.91427387 | 1 |
| SMPDL3A | 0.0044569 | 0.91406741 | 1 |
| TMEM132A | 0.01720434 | 0.91353837 | 1 |
| HSPBP1 | 0.00111522 | 0.91343386 | 1 |
| OSTF1 | 0.00017052 | 0.91093617 | 1 |
| MTPAP | 0.00412438 | 0.91078834 | 1 |
| HSD3B7 | 0.0085359 | 0.90999384 | 1 |
| ANO1 | 0.04520641 | 0.90807436 | 1 |
| SEC61A2 | 0.00701039 | 0.90719783 | 1 |
| XPO1 | 0.00022357 | 0.90671926 | 1 |
| MED10 | 0.00105164 | 0.90629817 | 1 |
| SF3A3 | 0.00268185 | 0.90594738 | 1 |
| LONRF2 | 0.02669996 | 0.90548917 | 1 |
| NALCN | 0.0417333 | 0.90488268 | 1 |
| TTYH2 | 0.01276602 | 0.90463925 | 1 |
| TMEM175 | 0.00178056 | 0.90329004 | 1 |
| ATP2C1 | 0.00011835 | 0.90316899 | 1 |
| FAIM | 0.00714929 | 0.90224578 | 1 |
| TAPBPL | 0.0006345 | 0.90191381 | 1 |
| CEP78 | 0.02438682 | 0.90178525 | 1 |
| FHIT | 0.01134523 | 0.90028563 | 1 |
| ZBTB9 | 0.01122524 | 0.90014867 | 1 |
| SURF1 | 0.00040965 | 0.89817186 | 1 |
| GSTA2 | 0.00701119 | 0.89665188 | 1 |
| AP1G2 | 0.00036389 | 0.89604103 | 1 |
| MYO5A | 0.00163209 | 0.89603546 | 1 |

| | | | |
|---|---|---|---|
| STAM | 0.00153913 | 0.8950116 | 1 |
| CYTH1 | 7.64E-05 | 0.8949378 | 1 |
| GALE | 0.00554016 | 0.8917589 | 1 |
| ABCA8 | 0.01918961 | 0.89157533 | 1 |
| RELL1 | 9.64E-05 | 0.89151803 | 1 |
| SPICE1 | 0.00594942 | 0.89086142 | 1 |
| TFB1M | 0.02389362 | 0.88986255 | 1 |
| PDS5A | 0.0007647 | 0.88953436 | 1 |
| VILL | 0.02704101 | 0.88872413 | 1 |
| CDCA4 | 0.02460238 | 0.88828857 | 1 |
| SCN4B | 0.02951577 | 0.88788175 | 1 |
| PSME2 | 0.01000974 | 0.88730011 | 1 |
| CD55 | 0.00054875 | 0.88696113 | 1 |
| B4GAT1 | 0.00180082 | 0.88618383 | 1 |
| MAK | 0.0417333 | 0.88608883 | 1 |
| VMAC | 0.00523045 | 0.88575441 | 1 |
| PARP6 | 0.00024307 | 0.88513009 | 1 |
| CSNK1A1 | 0.00072331 | 0.88427948 | 1 |
| CDK7 | 0.0409877 | 0.88418177 | 1 |
| ATP13A3 | 0.0003024 | 0.88338618 | 1 |
| SLC30A5 | 0.00044545 | 0.88261119 | 1 |
| PTP4A3 | 0.00181962 | 0.88253064 | 1 |
| STRADA | 0.02023032 | 0.88251924 | 1 |
| ZNF790 | 0.04348533 | 0.88158206 | 1 |
| TANGO6 | 0.00148499 | 0.88005383 | 1 |
| GTF2A1 | 0.0005035 | 0.87971447 | 1 |
| LRP1 | 0.00014375 | 0.87955669 | 1 |
| ZNF518B | 0.00425198 | 0.87863885 | 1 |
| MAPKAP1 | 0.00015498 | 0.87807526 | 1 |
| MIEF1 | 0.00272929 | 0.87787286 | 1 |
| ARL11 | 0.02744554 | 0.87703795 | 1 |
| PCED1A | 0.00012729 | 0.8766669 | 1 |
| IDH1 | 0.00010849 | 0.8765123 | 1 |
| PRR3 | 0.01977782 | 0.87636918 | 1 |
| EFCAB14 | 0.00010073 | 0.87630098 | 1 |
| MKKS | 0.00026355 | 0.87503867 | 1 |
| BAIAP2L1 | 0.02184252 | 0.8743354 | 1 |
| UBE2D4 | 0.0017579 | 0.87430068 | 1 |
| ZNF786 | 0.01898198 | 0.87360717 | 1 |
| CUL9 | 0.0066764 | 0.87350103 | 1 |
| BRSK1 | 0.02064445 | 0.87337921 | 1 |
| KIAA1324L | 0.00266583 | 0.87314657 | 1 |
| ECM1 | 0.00025394 | 0.87306674 | 1 |

| | | | |
|---|---|---|---|
| ETFBKMT | 0.01033408 | 0.8729911 | 1 |
| CASP8AP2 | 0.00109276 | 0.87251218 | 1 |
| FGF9 | 0.00567664 | 0.87180255 | 1 |
| GOLGA5 | 0.00050055 | 0.87166753 | 1 |
| FBLIM1 | 0.00045621 | 0.8716493 | 1 |
| ZNF277 | 0.00041455 | 0.87146585 | 1 |
| ZNF362 | 0.00016025 | 0.87019402 | 1 |
| UMPS | 0.00205061 | 0.86997678 | 1 |
| ARHGEF11 | 6.39E-05 | 0.86978962 | 1 |
| HK1 | 9.04E-05 | 0.86871729 | 1 |
| METTL25 | 0.00498011 | 0.86838283 | 1 |
| PSMD1 | 0.0001247 | 0.8679416 | 1 |
| IQCJ | 0.01399411 | 0.86738136 | 1 |
| CHFR | 0.00137434 | 0.867356 | 1 |
| PDE7A | 0.00060377 | 0.86703136 | 1 |
| RNF149 | 0.00104752 | 0.86680174 | 1 |
| C6orf136 | 0.03354378 | 0.86665096 | 1 |
| CCT2 | 0.00495049 | 0.86665079 | 1 |
| NAA30 | 0.00123137 | 0.86630369 | 1 |
| NNMT | 0.00013171 | 0.86604851 | 1 |
| SEMA4D | 0.00318395 | 0.86602909 | 1 |
| ERI1 | 0.00043376 | 0.86586315 | 1 |
| ASAP3 | 0.02869063 | 0.8656271 | 1 |
| ZNF8 | 0.00515743 | 0.86546261 | 1 |
| IL17RA | 0.00129631 | 0.86392756 | 1 |
| TMEM87A | 0.01584478 | 0.86323029 | 1 |
| C5orf22 | 0.01111257 | 0.86311414 | 1 |
| DNAJC18 | 0.00094239 | 0.86247893 | 1 |
| LSP1 | 0.00036585 | 0.86199673 | 1 |
| C20orf27 | 0.04315933 | 0.86197179 | 1 |
| NUCB2 | 0.00037859 | 0.86136027 | 1 |
| RAB29 | 0.00157314 | 0.86124936 | 1 |
| POLR3C | 0.00213005 | 0.86122018 | 1 |
| GPI | 0.00012599 | 0.86105998 | 1 |
| ACP1 | 0.01370281 | 0.86100559 | 1 |
| NUDT12 | 0.01008991 | 0.86080146 | 1 |
| METTL22 | 0.00044647 | 0.85906589 | 1 |
| NPRL3 | 0.00372838 | 0.85815161 | 1 |
| AGPS | 0.00086625 | 0.85664518 | 1 |
| PAICS | 0.0417333 | 0.85654071 | 1 |
| XPO6 | 0.00179515 | 0.85616417 | 1 |
| SLC17A5 | 0.00271691 | 0.85588982 | 1 |
| CCDC137 | 0.01065929 | 0.85536694 | 1 |

| | | | |
|---|---|---|---|
| PDGFA | 0.00061336 | 0.85531749 | 1 |
| ZNF524 | 0.01577953 | 0.85501436 | 1 |
| GP1BB | 0.01749137 | 0.85489504 | 1 |
| LATS1 | 0.00157704 | 0.85416551 | 1 |
| ZNF628 | 0.04236844 | 0.85321407 | 1 |
| MLLT3 | 0.00143003 | 0.85251584 | 1 |
| RAD51C | 0.00639764 | 0.85226126 | 1 |
| COL1A1 | 0.00578336 | 0.8520983 | 1 |
| USF1 | 0.00050692 | 0.85207741 | 1 |
| AP2S1 | 0.00868636 | 0.85125818 | 1 |
| WDR73 | 0.00207281 | 0.85111084 | 1 |
| XYLB | 0.02508682 | 0.8503186 | 1 |
| ZNF423 | 0.00058397 | 0.85005752 | 1 |
| CXCR4 | 0.00026669 | 0.85000662 | 1 |
| PROSER3 | 0.00547954 | 0.84971735 | 1 |
| HIF1A | 0.00016253 | 0.84960596 | 1 |
| FOXK1 | 0.00075269 | 0.84946384 | 1 |
| ATF1 | 0.00372521 | 0.84919468 | 1 |
| LIPA | 0.00052549 | 0.84911273 | 1 |
| EXOC3L1 | 0.01352451 | 0.84903659 | 1 |
| EXTL3 | 0.00275036 | 0.84893459 | 1 |
| FANCC | 0.01591542 | 0.8488683 | 1 |
| KIF1C | 0.00018298 | 0.84796485 | 1 |
| UTP3 | 0.00031966 | 0.84708592 | 1 |
| ELOVL7 | 0.00877725 | 0.84665865 | 1 |
| AP3M2 | 0.00839264 | 0.84624008 | 1 |
| ZFP90 | 0.00034141 | 0.84595144 | 1 |
| SRCAP | 0.0002041 | 0.84569043 | 1 |
| GAS2L1 | 0.00030373 | 0.84531573 | 1 |
| COL5A1 | 0.01066812 | 0.84420061 | 1 |
| ARSG | 0.04029729 | 0.84261879 | 1 |
| DPH3 | 0.0007568 | 0.84227901 | 1 |
| BTN2A1 | 0.00299423 | 0.84220073 | 1 |
| C1orf162 | 0.00023493 | 0.84155826 | 1 |
| NUBPL | 0.00221862 | 0.83991215 | 1 |
| STK40 | 0.00064879 | 0.83977632 | 1 |
| RBL2 | 0.00011363 | 0.83926673 | 1 |
| CRTC1 | 0.01204558 | 0.83913875 | 1 |
| XRCC4 | 0.02704808 | 0.83890038 | 1 |
| SH3KBP1 | 0.00089502 | 0.83887874 | 1 |
| DENND6B | 0.02985889 | 0.8386753 | 1 |
| ESS2 | 0.00263768 | 0.83814935 | 1 |
| UBQLN1 | 0.00304129 | 0.8375134 | 1 |

| | | | |
|---|---|---|---|
| APOBR | 0.00038527 | 0.83691601 | 1 |
| PDXP | 0.00564497 | 0.83579204 | 1 |
| PREP | 0.00121194 | 0.83559414 | 1 |
| TMEM123 | 0.0007264 | 0.83547819 | 1 |
| CMC2 | 0.00013656 | 0.83523183 | 1 |
| CARF | 0.00017008 | 0.83515106 | 1 |
| UBA6 | 0.00385879 | 0.83496253 | 1 |
| SALL3 | 0.04730337 | 0.83327721 | 1 |
| GRIP1 | 0.039477 | 0.83274146 | 1 |
| OXSM | 0.001712 | 0.83257037 | 1 |
| PLEKHG2 | 0.01327899 | 0.83222293 | 1 |
| TMEM80 | 0.01930519 | 0.83219352 | 1 |
| G6PC3 | 0.00577485 | 0.83216813 | 1 |
| DNAJC16 | 0.00680957 | 0.83131415 | 1 |
| TUBB4B | 0.00267434 | 0.83101804 | 1 |
| PHACTR1 | 0.00871178 | 0.83065824 | 1 |
| USH1C | 0.00010274 | 0.83045288 | 1 |
| LONRF3 | 0.02002046 | 0.83033716 | 1 |
| SNRPB2 | 0.01925588 | 0.82834467 | 1 |
| DUS4L | 0.03525059 | 0.82825612 | 1 |
| RTL6 | 0.0006541 | 0.82803228 | 1 |
| NUMBL | 0.00077896 | 0.8269437 | 1 |
| TBRG1 | 0.00678828 | 0.82675536 | 1 |
| ACSL1 | 0.00020289 | 0.82575405 | 1 |
| PYGM | 0.02512216 | 0.82496152 | 1 |
| CDS1 | 0.00952053 | 0.82398426 | 1 |
| BAG4 | 0.02413237 | 0.82366225 | 1 |
| P2RX1 | 0.0464536 | 0.82324049 | 1 |
| MED19 | 0.03361206 | 0.82254685 | 1 |
| ZMYND10 | 0.0417333 | 0.82210743 | 1 |
| SNX13 | 0.0005637 | 0.82114233 | 1 |
| SUCNR1 | 0.00198749 | 0.8210461 | 1 |
| POLA1 | 0.00270375 | 0.82076631 | 1 |
| LOXL1 | 0.0022223 | 0.81901023 | 1 |
| ADGRE5 | 0.00065646 | 0.81758261 | 1 |
| TNFRSF1B | 0.00032647 | 0.81734957 | 1 |
| YBX1 | 0.00018111 | 0.81617376 | 1 |
| LRRC59 | 0.00091134 | 0.81614275 | 1 |
| DIAPH2 | 0.00072735 | 0.81564384 | 1 |
| ZNF500 | 0.00081323 | 0.81527412 | 1 |
| PPIP5K2 | 0.00377475 | 0.81515847 | 1 |
| CDC14A | 0.0007747 | 0.8149666 | 1 |
| ZNF398 | 0.01030998 | 0.81457896 | 1 |

| | | | |
|---|---|---|---|
| C16orf54 | 0.03565524 | 0.81419053 | 1 |
| PJVK | 0.01475934 | 0.81129208 | 1 |
| MGME1 | 0.00623896 | 0.81118202 | 1 |
| SDK2 | 0.01445366 | 0.81085705 | 1 |
| SLC25A26 | 0.01197826 | 0.81082861 | 1 |
| PAPSS1 | 0.00044417 | 0.81076614 | 1 |
| RPN2 | 0.00017931 | 0.81073337 | 1 |
| NDUFA11 | 0.00042105 | 0.81028847 | 1 |
| MIS12 | 0.00241149 | 0.81019348 | 1 |
| NUP43 | 0.00098623 | 0.80999149 | 1 |
| SLC39A4 | 0.01524893 | 0.80988986 | 1 |
| BCL6B | 0.00398721 | 0.80966873 | 1 |
| UNC5C | 0.01391163 | 0.80861446 | 1 |
| STRIP1 | 0.00299707 | 0.80841379 | 1 |
| RNF135 | 0.00068195 | 0.80834329 | 1 |
| MGAT3 | 0.00386225 | 0.80801957 | 1 |
| NFYC | 0.00932589 | 0.80745823 | 1 |
| ZNF318 | 0.00185849 | 0.80681026 | 1 |
| TGS1 | 0.00340198 | 0.80628311 | 1 |
| IAH1 | 0.00163612 | 0.80611518 | 1 |
| LPGAT1 | 0.00063799 | 0.80604464 | 1 |
| SLC35G2 | 0.01098148 | 0.80602503 | 1 |
| TRPM6 | 0.02775365 | 0.80550466 | 1 |
| NIFK | 0.04396267 | 0.80547524 | 1 |
| ZSCAN21 | 0.00013579 | 0.8050669 | 1 |
| CHIC2 | 0.00288792 | 0.80456737 | 1 |
| ATAD2 | 0.0158063 | 0.8044049 | 1 |
| ARIH2 | 6.50E-05 | 0.80376087 | 1 |
| TBC1D15 | 0.00013588 | 0.8030274 | 1 |
| PLEKHH2 | 0.00337896 | 0.80285431 | 1 |
| TCOF1 | 0.0034579 | 0.80255566 | 1 |
| TYSND1 | 0.00106269 | 0.80248829 | 1 |
| STEAP3 | 0.00194297 | 0.80245159 | 1 |
| PUS1 | 9.90E-05 | 0.80174706 | 1 |
| PI16 | 0.00829332 | 0.80152567 | 1 |
| AP2M1 | 0.00043667 | 0.80097087 | 1 |
| LURAP1L | 0.0349498 | 0.80083982 | 1 |
| AC092835.1 | 0.00416253 | 0.80080278 | 1 |
| CHMP4A | 0.00028347 | 0.8002846 | 1 |
| SBF2 | 0.00019193 | 0.79989864 | 1 |
| RHOBTB2 | 0.00099048 | 0.7994573 | 1 |
| CNTNAP1 | 0.01872295 | 0.79929751 | 1 |
| TUSC3 | 0.00026247 | 0.79900836 | 1 |

| | | | |
|---|---|---|---|
| SLC22A11 | 0.00392493 | 0.7989931 | 1 |
| KRTCAP3 | 0.01496153 | 0.79820268 | 1 |
| ASB1 | 0.00558101 | 0.79765792 | 1 |
| ZCCHC24 | 0.00027735 | 0.79719089 | 1 |
| MAFF | 0.03608466 | 0.79713047 | 1 |
| CDC25B | 0.00046844 | 0.79710492 | 1 |
| ZSCAN29 | 0.00106778 | 0.79661324 | 1 |
| INTS14 | 0.00168178 | 0.79545249 | 1 |
| PSMB7 | 0.01087475 | 0.79437191 | 1 |
| RAI2 | 0.01453944 | 0.79362527 | 1 |
| EMC1 | 0.00039452 | 0.79322954 | 1 |
| EEF1E1 | 0.01361644 | 0.79260682 | 1 |
| TLN2 | 0.00371303 | 0.792578 | 1 |
| NUAK1 | 0.00196882 | 0.79172983 | 1 |
| SAE1 | 0.00736897 | 0.79165745 | 1 |
| FPGS | 0.00149521 | 0.79126961 | 1 |
| KRBA1 | 0.01701172 | 0.79121397 | 1 |
| C7orf26 | 0.00181222 | 0.79061167 | 1 |
| DGKG | 0.02848263 | 0.7898255 | 1 |
| APOD | 0.00516147 | 0.7897989 | 1 |
| COX17 | 0.00028419 | 0.78949043 | 1 |
| APC | 0.00077512 | 0.78927133 | 1 |
| GJA5 | 0.00066966 | 0.78891234 | 1 |
| ATG14 | 0.00182497 | 0.78886307 | 1 |
| YBEY | 0.01177323 | 0.78784506 | 1 |
| KCTD1 | 0.00510247 | 0.78783196 | 1 |
| SMC6 | 0.00351793 | 0.78743531 | 1 |
| C1QL1 | 0.0009663 | 0.78722954 | 1 |
| ECT2 | 0.00862452 | 0.78631792 | 1 |
| NAGK | 0.00073361 | 0.78528203 | 1 |
| LSM10 | 0.00040227 | 0.78513732 | 1 |
| MCUR1 | 0.00044707 | 0.78500193 | 1 |
| FAM102B | 0.00124525 | 0.7849508 | 1 |
| PKP4 | 0.01280737 | 0.78469687 | 1 |
| NHS | 0.00098362 | 0.78403493 | 1 |
| SPATA5 | 0.00986322 | 0.78389213 | 1 |
| METTL3 | 0.00026857 | 0.78273722 | 1 |
| MSRB1 | 0.00034998 | 0.78213311 | 1 |
| NIT2 | 0.00016846 | 0.78212469 | 1 |
| EXOSC10 | 0.00121126 | 0.78209106 | 1 |
| DLG4 | 0.00219063 | 0.7819931 | 1 |
| PRRC2B | 0.00042733 | 0.78180698 | 1 |
| DNAJC10 | 0.00037519 | 0.78113865 | 1 |

| | | | |
|---|---|---|---|
| STK19 | 0.00752814 | 0.7808774 | 1 |
| SAMD12 | 0.02348653 | 0.78059799 | 1 |
| MAPKAPK3 | 0.00016704 | 0.77900338 | 1 |
| NANS | 0.01229523 | 0.77894696 | 1 |
| INPP4A | 0.00059606 | 0.77892375 | 1 |
| SHROOM1 | 0.04871831 | 0.77887277 | 1 |
| DHX29 | 0.00102446 | 0.77865111 | 1 |
| ADAM9 | 0.00021912 | 0.77841088 | 1 |
| COX7B | 0.00098692 | 0.77835452 | 1 |
| CSPG4 | 0.00181738 | 0.7781 | 1 |
| MOSMO | 0.00383786 | 0.77734512 | 1 |
| CYP26B1 | 0.00645949 | 0.77733985 | 1 |
| SETX | 7.61E-05 | 0.77711648 | 1 |
| TNPO2 | 0.00017592 | 0.77686523 | 1 |
| PUM3 | 0.01018634 | 0.77626233 | 1 |
| SNRNP25 | 0.0017612 | 0.77566281 | 1 |
| KIF22 | 0.00839264 | 0.77495074 | 1 |
| FEZ1 | 0.00839264 | 0.77468215 | 1 |
| CTTNBP2NL | 0.00342248 | 0.77450656 | 1 |
| PHF1 | 0.00041458 | 0.77405125 | 1 |
| SLC6A6 | 0.00082462 | 0.77355716 | 1 |
| CPSF4 | 0.00107361 | 0.77310781 | 1 |
| PLEKHF1 | 0.03023443 | 0.77300708 | 1 |
| PIGV | 0.01170362 | 0.7727888 | 1 |
| CCN4 | 0.00340123 | 0.77167897 | 1 |
| ARL2BP | 0.00037884 | 0.7714953 | 1 |
| MAPK1IP1L | 0.00228016 | 0.77119815 | 1 |
| SIL1 | 0.00071348 | 0.77118023 | 1 |
| GALNT1 | 0.00225618 | 0.77101332 | 1 |
| CLEC1A | 0.01328519 | 0.77022054 | 1 |
| PRDX3 | 0.00099508 | 0.76977226 | 1 |
| FAM193A | 0.00048207 | 0.76974279 | 1 |
| ING2 | 0.00372523 | 0.7696469 | 1 |
| CAP2 | 0.01872295 | 0.76914058 | 1 |
| SNRK | 5.62E-05 | 0.76842983 | 1 |
| MAP6 | 0.00119268 | 0.76818958 | 1 |
| USP3 | 0.00170099 | 0.76749264 | 1 |
| E2F5 | 0.03255053 | 0.76725908 | 1 |
| SLCO5A1 | 0.02064445 | 0.76697078 | 1 |
| DNALI1 | 0.00659617 | 0.76674904 | 1 |
| VBP1 | 0.01434395 | 0.76635732 | 1 |
| VAT1L | 0.01007812 | 0.76597692 | 1 |
| RGS19 | 0.01989683 | 0.76572381 | 1 |

| | | | |
|---|---|---|---|
| NFIC | 6.94E-05 | 0.76494836 | 1 |
| FTCD | 0.04110217 | 0.76340475 | 1 |
| DNAJC21 | 0.0002082 | 0.76326187 | 1 |
| DENND5A | 0.00274967 | 0.76311429 | 1 |
| CD84 | 0.0060708 | 0.76285937 | 1 |
| CD180 | 0.04980502 | 0.7627704 | 1 |
| SLC25A51 | 0.014242 | 0.76251698 | 1 |
| PARL | 0.00159434 | 0.76237032 | 1 |
| CPSF6 | 0.009462 | 0.76210663 | 1 |
| PAM16 | 0.00111152 | 0.76142212 | 1 |
| ANAPC5 | 0.00084374 | 0.75965287 | 1 |
| GON4L | 0.00461453 | 0.75953681 | 1 |
| CEP104 | 0.00768729 | 0.75885639 | 1 |
| FAM49B | 0.02144993 | 0.75872153 | 1 |
| F13A1 | 0.00535584 | 0.75841288 | 1 |
| COASY | 0.00216848 | 0.75799217 | 1 |
| HHAT | 0.01172894 | 0.75767021 | 1 |
| MAPT | 0.00040941 | 0.7570151 | 1 |
| WRN | 0.00796299 | 0.75659324 | 1 |
| ADCY7 | 0.00942989 | 0.75644471 | 1 |
| ACAD10 | 0.01540948 | 0.75619186 | 1 |
| FAM110B | 0.04210141 | 0.75603467 | 1 |
| ENTR1 | 0.00206211 | 0.75565745 | 1 |
| KNSTRN | 0.00185977 | 0.75558792 | 1 |
| BPNT1 | 0.00052222 | 0.75551527 | 1 |
| RELN | 0.00953331 | 0.75427516 | 1 |
| MARVELD2 | 0.0099419 | 0.75372787 | 1 |
| HLF | 0.00671341 | 0.75359762 | 1 |
| RHBDF2 | 0.00231983 | 0.75298292 | 1 |
| STK11 | 0.00223142 | 0.75279416 | 1 |
| RARB | 0.00328289 | 0.75260686 | 1 |
| ZNF408 | 0.02319702 | 0.7523833 | 1 |
| TMA7 | 0.02783037 | 0.75220925 | 1 |
| LMBRD1 | 0.00011971 | 0.75216705 | 1 |
| SIGLEC9 | 0.00344721 | 0.75184777 | 1 |
| GTF2H1 | 0.00096117 | 0.75088278 | 1 |
| RNASEL | 0.00797566 | 0.75075572 | 1 |
| SIKE1 | 0.00024797 | 0.75054813 | 1 |
| ADAM20 | 0.01720434 | 0.75035288 | 1 |
| TXLNG | 0.03462597 | 0.74968085 | 1 |
| DMAC1 | 0.02726218 | 0.74960775 | 1 |
| TSPAN17 | 0.00033385 | 0.74959841 | 1 |
| EPRS | 0.00051841 | 0.74939601 | 1 |

| | | | |
|---|---|---|---|
| VEGFC | 0.03000926 | 0.74890719 | 1 |
| FAM160B2 | 0.00035219 | 0.74877941 | 1 |
| IQCA1 | 0.04372446 | 0.74810847 | 1 |
| CD58 | 0.00131187 | 0.74802288 | 1 |
| CCAR1 | 0.01033649 | 0.74702334 | 1 |
| COL15A1 | 0.02506977 | 0.74695377 | 1 |
| ZNF174 | 0.00811959 | 0.74692757 | 1 |
| AAAS | 0.00040602 | 0.74647779 | 1 |
| MOK | 0.02783858 | 0.7462924 | 1 |
| SLC36A1 | 0.00304509 | 0.74594935 | 1 |
| NGFR | 0.00316383 | 0.74564195 | 1 |
| DAGLB | 0.00553838 | 0.74486222 | 1 |
| MYPOP | 0.02184252 | 0.74477908 | 1 |
| BIRC6 | 0.00119145 | 0.74465445 | 1 |
| ZC3HC1 | 0.00850022 | 0.74428501 | 1 |
| TESK1 | 0.01261098 | 0.74406164 | 1 |
| NDUFA9 | 0.00499299 | 0.74374529 | 1 |
| MAP3K5 | 0.00056348 | 0.74237921 | 1 |
| FAM20B | 0.0007334 | 0.74199474 | 1 |
| TMEM44 | 0.00501322 | 0.7417705 | 1 |
| SLC8B1 | 0.00623695 | 0.74159284 | 1 |
| KHSRP | 0.00361592 | 0.74156854 | 1 |
| ATG5 | 0.00174388 | 0.74114618 | 1 |
| SNRNP48 | 0.00423365 | 0.74102473 | 1 |
| RSBN1 | 0.00104248 | 0.74025526 | 1 |
| ITGA4 | 0.01904079 | 0.73973908 | 1 |
| CERS2 | 0.00157833 | 0.73948811 | 1 |
| MMP19 | 0.01145859 | 0.73911825 | 1 |
| RAD50 | 0.01913567 | 0.73826861 | 1 |
| CDC26 | 0.04176457 | 0.73786936 | 1 |
| SLX4 | 0.02246742 | 0.73756845 | 1 |
| GSTO1 | 0.00166178 | 0.73712698 | 1 |
| WNT6 | 0.03551512 | 0.73568057 | 1 |
| HCFC2 | 0.00334316 | 0.73566816 | 1 |
| SREBF1 | 0.00085857 | 0.73560749 | 1 |
| ASPSCR1 | 0.00474062 | 0.73456903 | 1 |
| IL17RC | 0.00702934 | 0.73447271 | 1 |
| MAP3K11 | 0.00033076 | 0.7343203 | 1 |
| GTF3C1 | 0.0019644 | 0.73431239 | 1 |
| CD37 | 0.00812255 | 0.73428465 | 1 |
| MTMR14 | 0.00495725 | 0.73422158 | 1 |
| PSTPIP1 | 0.01376212 | 0.73369538 | 1 |
| COX4I1 | 0.00015006 | 0.73350232 | 1 |

| | | | |
|---|---|---|---|
| SLC26A6 | 0.00726918 | 0.73324203 | 1 |
| TEDC1 | 0.01784279 | 0.73321782 | 1 |
| PIP5K1C | 0.00296423 | 0.73258979 | 1 |
| TBCE | 0.01087302 | 0.73255869 | 1 |
| LSM12 | 0.02054419 | 0.7324632 | 1 |
| CPXM1 | 0.0032892 | 0.73232449 | 1 |
| FASTKD5 | 0.00512147 | 0.73128319 | 1 |
| EHBP1L1 | 0.00010827 | 0.73110927 | 1 |
| HBEGF | 0.04927725 | 0.730951 | 1 |
| FDFT1 | 0.00148379 | 0.7309392 | 1 |
| AP4E1 | 0.00438929 | 0.73087987 | 1 |
| MOSPD2 | 7.68E-05 | 0.7308157 | 1 |
| CLDN19 | 0.02960939 | 0.73070988 | 1 |
| IMPA1 | 0.00220376 | 0.73048602 | 1 |
| RWDD2A | 0.01983126 | 0.73025468 | 1 |
| BBS10 | 0.03177746 | 0.73006422 | 1 |
| IQCE | 0.00244166 | 0.72911146 | 1 |
| POLB | 0.03329415 | 0.72908853 | 1 |
| SLC49A3 | 0.00411015 | 0.72902414 | 1 |
| LCMT2 | 0.04043885 | 0.72875409 | 1 |
| CAMK1 | 0.00308212 | 0.72810451 | 1 |
| HDAC3 | 0.00442171 | 0.72806714 | 1 |
| RDX | 0.00226044 | 0.72765736 | 1 |
| MAP3K14 | 0.02327294 | 0.72758733 | 1 |
| CERT1 | 0.00073268 | 0.72709774 | 1 |
| KIAA1841 | 0.00377503 | 0.72690996 | 1 |
| MRVI1 | 0.00130389 | 0.72682556 | 1 |
| WDR35 | 0.00637953 | 0.72546701 | 1 |
| LYPD6B | 0.02899551 | 0.72509905 | 1 |
| EGFLAM | 0.01262454 | 0.72491579 | 1 |
| POSTN | 0.00261307 | 0.72374669 | 1 |
| DESI1 | 0.001023 | 0.72356193 | 1 |
| EPHA1 | 0.0381598 | 0.72350216 | 1 |
| LRRN2 | 0.01519877 | 0.7230632 | 1 |
| RNASE6 | 0.00386837 | 0.72236015 | 1 |
| EIF2D | 0.00354274 | 0.72187007 | 1 |
| RBBP9 | 0.00338267 | 0.7215216 | 1 |
| WASF1 | 0.01195239 | 0.72150919 | 1 |
| GALNT2 | 0.03251768 | 0.72131807 | 1 |
| SNRPN | 0.0011829 | 0.72125263 | 1 |
| RNPEP | 0.00320075 | 0.72123775 | 1 |
| PITHD1 | 0.00105164 | 0.72001705 | 1 |
| FBN1 | 0.00123322 | 0.71995962 | 1 |

| | | | |
|---|---|---|---|
| CREB3L2 | 0.00227757 | 0.7197038 | 1 |
| JAML | 0.0045938 | 0.71846059 | 1 |
| REPS2 | 0.0030287 | 0.71845501 | 1 |
| SEPTIN10 | 0.00089653 | 0.71740915 | 1 |
| ORMDL3 | 0.00074277 | 0.71725132 | 1 |
| SRPK2 | 0.00412438 | 0.71718532 | 1 |
| FOXC2 | 0.00236023 | 0.71715753 | 1 |
| WNT5A | 0.01164783 | 0.71704927 | 1 |
| CADM1 | 0.00409904 | 0.71704869 | 1 |
| RYK | 0.01721952 | 0.7168081 | 1 |
| RAD23B | 0.00867426 | 0.71536227 | 1 |
| CUTC | 0.00385242 | 0.71482623 | 1 |
| EIF4G2 | 0.00085496 | 0.714616 | 1 |
| MPP7 | 0.01837724 | 0.71373119 | 1 |
| SERPINB8 | 0.00238966 | 0.71367922 | 1 |
| HMCES | 0.00249912 | 0.71226707 | 1 |
| MRTFB | 0.00658364 | 0.71200563 | 1 |
| ZNF322 | 0.01620917 | 0.71180282 | 1 |
| TBC1D10C | 0.03490566 | 0.71170324 | 1 |
| CRKL | 0.00117994 | 0.71124859 | 1 |
| CABLES1 | 0.00362836 | 0.71117955 | 1 |
| SERINC3 | 0.00015464 | 0.7097418 | 1 |
| ZNF227 | 0.043715 | 0.7096458 | 1 |
| DVL2 | 0.0205797 | 0.70964413 | 1 |
| CD53 | 0.00112962 | 0.70956532 | 1 |
| SEC61B | 0.00014031 | 0.70948559 | 1 |
| RRP8 | 0.01897316 | 0.70935194 | 1 |
| AP3M1 | 0.00668235 | 0.70931339 | 1 |
| RAP2A | 0.00067569 | 0.70882153 | 1 |
| HCN3 | 0.04644879 | 0.70733152 | 1 |
| CDC42BPB | 0.00012181 | 0.70723083 | 1 |
| AMT | 0.00069954 | 0.70680863 | 1 |
| LMBR1L | 0.04249481 | 0.70654368 | 1 |
| PAIP2 | 0.00243547 | 0.70569677 | 1 |
| ARHGEF40 | 0.00134446 | 0.70554346 | 1 |
| FAM234A | 0.00201043 | 0.70545048 | 1 |
| ARF4 | 0.00443846 | 0.7051599 | 1 |
| IFT27 | 0.00703048 | 0.70480209 | 1 |
| IL13RA2 | 0.00245029 | 0.70411019 | 1 |
| RALY | 0.00866034 | 0.70394821 | 1 |
| VAMP7 | 0.00430114 | 0.70380174 | 1 |
| ETV6 | 0.00023197 | 0.70359224 | 1 |
| LMTK2 | 0.00523045 | 0.70307919 | 1 |

| | | | |
|---|---|---|---|
| ZNF841 | 0.02612874 | 0.70267976 | 1 |
| MEGF8 | 0.00724039 | 0.70264485 | 1 |
| MLH3 | 0.00206099 | 0.70229208 | 1 |
| PCDH17 | 0.0202365 | 0.70228089 | 1 |
| EHMT1 | 0.00334807 | 0.70220145 | 1 |
| NAGA | 0.00456411 | 0.7021265 | 1 |
| NIBAN2 | 0.0011056 | 0.702021 | 1 |
| SUFU | 0.0286182 | 0.70138678 | 1 |
| FAM167B | 0.01606987 | 0.7012426 | 1 |
| SCML1 | 0.01814369 | 0.70113105 | 1 |
| SEC16A | 0.00140494 | 0.701055 | 1 |
| SLC35D1 | 0.00656488 | 0.7004899 | 1 |
| TEC | 0.01836627 | 0.7004329 | 1 |
| PRPSAP2 | 0.02217394 | 0.70039686 | 1 |
| PRKRA | 0.00207602 | 0.700363 | 1 |
| FAM126B | 0.00424922 | 0.69999573 | 1 |
| PDE1A | 0.0001484 | 0.6996419 | 1 |
| TMEM30A | 0.00512537 | 0.69909652 | 1 |
| ALKBH2 | 0.017172 | 0.69848414 | 1 |
| MPP5 | 0.00031937 | 0.69816081 | 1 |
| ANAPC13 | 0.00479594 | 0.69795481 | 1 |
| FAM104B | 0.00617055 | 0.69789795 | 1 |
| PIH1D1 | 0.00168471 | 0.69691387 | 1 |
| RCAN2 | 0.00214766 | 0.69689727 | 1 |
| MTMR9 | 0.00132507 | 0.69573955 | 1 |
| ADAM8 | 0.04644879 | 0.69553696 | 1 |
| SULF1 | 0.00324753 | 0.6954379 | 1 |
| ACKR3 | 0.02741578 | 0.69431208 | 1 |
| TFPT | 0.00807004 | 0.69412513 | 1 |
| ITGA5 | 0.00211898 | 0.69379976 | 1 |
| TMEM266 | 0.01688647 | 0.69361269 | 1 |
| UNC93B1 | 0.00628421 | 0.69324149 | 1 |
| HNRNPC | 5.79E-05 | 0.69249293 | 1 |
| SULF2 | 0.00025532 | 0.69169859 | 1 |
| MDH2 | 0.00155121 | 0.69166522 | 1 |
| TAB2 | 0.00085365 | 0.69092794 | 1 |
| TENT5A | 0.00839572 | 0.69036326 | 1 |
| CD99 | 0.00073301 | 0.68997321 | 1 |
| EML1 | 0.00493734 | 0.68983707 | 1 |
| RAD51D | 0.00339656 | 0.68975088 | 1 |
| NAA80 | 0.03729744 | 0.68960914 | 1 |
| TGFB1I1 | 0.00074792 | 0.68957 | 1 |
| MFSD14A | 0.00392109 | 0.68955725 | 1 |

| | | | |
|---|---|---|---|
| DDX39A | 0.00462471 | 0.68934966 | 1 |
| USP40 | 0.00028002 | 0.68929722 | 1 |
| HCFC1R1 | 0.00027778 | 0.68923813 | 1 |
| NSF | 0.00104678 | 0.68831893 | 1 |
| TUBB2A | 0.02748648 | 0.68828289 | 1 |
| SGMS2 | 0.00460932 | 0.68826055 | 1 |
| MCRIP1 | 0.00095863 | 0.68784157 | 1 |
| MORF4L2 | 0.0018556 | 0.68732039 | 1 |
| KCTD10 | 0.00458371 | 0.68670429 | 1 |
| ARL8A | 0.00107385 | 0.6866565 | 1 |
| HNRNPLL | 0.01451486 | 0.68545308 | 1 |
| ADRA2C | 0.02820909 | 0.68495462 | 1 |
| TRPV1 | 0.03418586 | 0.68452089 | 1 |
| CDK11A | 0.02512216 | 0.68318693 | 1 |
| CSK | 0.00245677 | 0.68310121 | 1 |
| AXL | 0.00050806 | 0.68247423 | 1 |
| ZNF133 | 0.00289769 | 0.68240913 | 1 |
| SFT2D2 | 0.00077964 | 0.68240794 | 1 |
| AC073111.4 | 0.0094382 | 0.68231238 | 1 |
| GRB2 | 0.00435435 | 0.6820806 | 1 |
| DPY19L3 | 0.00617875 | 0.68115914 | 1 |
| PRKACB | 0.01012427 | 0.68114263 | 1 |
| FAAP100 | 0.00062833 | 0.68103305 | 1 |
| IPO5 | 0.00365971 | 0.68068982 | 1 |
| LRRC14 | 0.00308943 | 0.68065075 | 1 |
| KAT2A | 0.04291638 | 0.68046985 | 1 |
| NOP16 | 0.00031299 | 0.68028277 | 1 |
| ZDHHC1 | 0.00877725 | 0.67955465 | 1 |
| SLC15A4 | 0.00446666 | 0.67942597 | 1 |
| KDELR2 | 0.00228617 | 0.67926462 | 1 |
| BCL10 | 0.00168394 | 0.67901054 | 1 |
| PRELP | 0.00013664 | 0.67900543 | 1 |
| SH3GL1 | 0.00024271 | 0.6788214 | 1 |
| ISG20L2 | 0.00613888 | 0.67841911 | 1 |
| ZNF546 | 0.04477859 | 0.67796378 | 1 |
| FAM241A | 0.00297792 | 0.67787375 | 1 |
| GATAD2A | 0.00150608 | 0.67775971 | 1 |
| OXA1L | 0.00734408 | 0.67720002 | 1 |
| EEF1AKNMT | 0.00145802 | 0.67703619 | 1 |
| DHRSX | 0.00486712 | 0.67644394 | 1 |
| CCDC183 | 0.02608777 | 0.67600985 | 1 |
| PODXL2 | 0.03565524 | 0.67599097 | 1 |
| ZDHHC13 | 0.01120299 | 0.67564136 | 1 |

| | | | |
|---|---|---|---|
| RP9 | 0.02484733 | 0.67532014 | 1 |
| TAF9B | 0.00352798 | 0.67500211 | 1 |
| TPMT | 0.00953285 | 0.67466348 | 1 |
| ENTPD1 | 0.00115302 | 0.67450964 | 1 |
| PRPS2 | 0.0060386 | 0.67345058 | 1 |
| S100A9 | 0.00097238 | 0.67325653 | 1 |
| PDLIM7 | 0.00070886 | 0.6732422 | 1 |
| IDUA | 0.00434818 | 0.67255712 | 1 |
| SCN7A | 0.0182496 | 0.67211386 | 1 |
| PIK3R5 | 0.00954646 | 0.67208932 | 1 |
| TOMM40L | 0.01658181 | 0.67208465 | 1 |
| TPM4 | 0.00156562 | 0.67200028 | 1 |
| ZC3HAV1L | 0.03341534 | 0.67169187 | 1 |
| SIN3A | 0.00237041 | 0.67100407 | 1 |
| KYAT3 | 0.00287098 | 0.67072881 | 1 |
| SLC1A5 | 0.00104972 | 0.67034857 | 1 |
| ZC3H15 | 0.00086254 | 0.66970398 | 1 |
| SPATA6 | 0.00036015 | 0.66965174 | 1 |
| FRG2C | 0.04817113 | 0.6694059 | 1 |
| UGGT2 | 0.00938304 | 0.6691654 | 1 |
| PYGO2 | 0.00744686 | 0.66915442 | 1 |
| REL | 0.0097272 | 0.6689533 | 1 |
| ZFP14 | 0.00443183 | 0.6688589 | 1 |
| VPS13D | 0.00067882 | 0.66857289 | 1 |
| NPAS3 | 0.00452791 | 0.66820251 | 1 |
| BTK | 0.00343503 | 0.66752305 | 1 |
| C8orf58 | 0.0103013 | 0.6672518 | 1 |
| SPRY2 | 0.00222383 | 0.66720906 | 1 |
| SPINDOC | 0.02502981 | 0.6671658 | 1 |
| PCDH1 | 0.00253272 | 0.66684511 | 1 |
| RASGRF2 | 0.04616903 | 0.66661164 | 1 |
| MPG | 0.00102805 | 0.66618861 | 1 |
| NCBP3 | 0.00035693 | 0.66611383 | 1 |
| FAM20C | 0.00051885 | 0.66590353 | 1 |
| ABCC1 | 0.0004472 | 0.66574956 | 1 |
| TARBP1 | 0.0033859 | 0.66568983 | 1 |
| OPN3 | 0.00399312 | 0.66564153 | 1 |
| SLC48A1 | 0.01054816 | 0.66517602 | 1 |
| TUT7 | 9.35E-05 | 0.66477733 | 1 |
| TEN1 | 0.00076187 | 0.66431688 | 1 |
| DHDH | 0.01061001 | 0.66418993 | 1 |
| ATP2B1 | 0.00237444 | 0.664059 | 1 |
| C1GALT1C1 | 0.00099162 | 0.66386503 | 1 |

| | | | |
|---|---|---|---|
| PDGFB | 0.00116052 | 0.66366431 | 1 |
| GRAMD1A | 0.00442171 | 0.66311075 | 1 |
| SNCA | 0.00274946 | 0.66245288 | 1 |
| C6orf62 | 0.00377549 | 0.6620097 | 1 |
| PAG1 | 0.00144142 | 0.6616572 | 1 |
| SNX5 | 0.01967212 | 0.66110196 | 1 |
| HECTD1 | 0.00050241 | 0.66088352 | 1 |
| AP1M2 | 0.01138457 | 0.66061338 | 1 |
| COG7 | 0.01587918 | 0.66014925 | 1 |
| SPTLC3 | 0.01626017 | 0.65992541 | 1 |
| TM2D1 | 0.00274967 | 0.6597778 | 1 |
| B3GLCT | 0.00572393 | 0.6596116 | 1 |
| DVL1 | 0.00058101 | 0.65942024 | 1 |
| TMEM25 | 0.01545404 | 0.65915436 | 1 |
| OSER1 | 0.00042974 | 0.65878464 | 1 |
| ZBTB47 | 0.00122916 | 0.6580895 | 1 |
| PIGN | 0.02755014 | 0.6572432 | 1 |
| TRNP1 | 0.00958202 | 0.65642863 | 1 |
| GNL1 | 0.00231995 | 0.65611538 | 1 |
| KIF3A | 0.00841731 | 0.65592606 | 1 |
| GRAMD2B | 0.00204302 | 0.65544929 | 1 |
| DNAJC8 | 0.00211895 | 0.65540478 | 1 |
| CLIC2 | 0.00631061 | 0.65528546 | 1 |
| ARV1 | 0.01540487 | 0.65513076 | 1 |
| TGFBRAP1 | 0.00684359 | 0.65511156 | 1 |
| MPZL2 | 0.04817113 | 0.65472145 | 1 |
| RNF14 | 0.04791564 | 0.65454254 | 1 |
| HDDC2 | 0.00044685 | 0.65446437 | 1 |
| UNC119 | 0.00960028 | 0.65408766 | 1 |
| HDHD3 | 0.01446331 | 0.65370106 | 1 |
| RNF168 | 0.00416598 | 0.65360419 | 1 |
| ATP8B1 | 0.00147011 | 0.65352077 | 1 |
| ODC1 | 0.02700396 | 0.65323512 | 1 |
| SLC7A6OS | 0.00471926 | 0.65312973 | 1 |
| PRORP | 0.00419763 | 0.65296775 | 1 |
| IL17RD | 0.03327079 | 0.65271433 | 1 |
| CD48 | 0.03422765 | 0.65259192 | 1 |
| ATP8B2 | 0.00053925 | 0.65254962 | 1 |
| SCAF4 | 0.00254385 | 0.65249818 | 1 |
| F8 | 0.00225655 | 0.65213731 | 1 |
| DGUOK | 0.00055864 | 0.65210494 | 1 |
| GANC | 0.00462233 | 0.65204244 | 1 |
| CD151 | 0.00038106 | 0.65188872 | 1 |

| | | | |
|---|---|---|---|
| ARMH4 | 0.00109299 | 0.65174754 | 1 |
| AGBL5 | 0.02077897 | 0.65164668 | 1 |
| TMED10 | 0.00473169 | 0.65139479 | 1 |
| CCL14 | 0.00596662 | 0.65071795 | 1 |
| GLUL | 0.00014542 | 0.65048695 | 1 |
| TMEM184B | 0.00021434 | 0.64961651 | 1 |
| ZC2HC1A | 0.01511757 | 0.64943775 | 1 |
| TRIP11 | 0.00879058 | 0.64940131 | 1 |
| TAZ | 0.02955023 | 0.64915355 | 1 |
| POLR1E | 0.02246742 | 0.64893641 | 1 |
| NDUFAF2 | 0.0066764 | 0.64875702 | 1 |
| GSK3B | 0.00267513 | 0.64861962 | 1 |
| ROBO3 | 0.03973009 | 0.64840241 | 1 |
| PIK3CA | 0.00069627 | 0.64810707 | 1 |
| MTIF2 | 0.00172263 | 0.64781402 | 1 |
| SGCB | 0.00031388 | 0.64650947 | 1 |
| STK10 | 0.00426374 | 0.64600498 | 1 |
| PLD6 | 0.01460898 | 0.64584546 | 1 |
| AP2B1 | 0.01840408 | 0.64538677 | 1 |
| ARID1B | 0.00156484 | 0.6450839 | 1 |
| TRPC1 | 0.01021608 | 0.64504857 | 1 |
| ZNF763 | 0.0464536 | 0.64483128 | 1 |
| SNRNP40 | 0.01130692 | 0.64460271 | 1 |
| PCOLCE | 0.01810239 | 0.64448929 | 1 |
| TRABD | 0.00304031 | 0.64441359 | 1 |
| PPP1R11 | 0.00389658 | 0.64406738 | 1 |
| UBAP2L | 0.00397787 | 0.64389918 | 1 |
| MCM4 | 0.00895432 | 0.64314795 | 1 |
| ABCF2 | 0.00641412 | 0.64267814 | 1 |
| SLC39A14 | 0.01332796 | 0.64262026 | 1 |
| ALDH1A2 | 0.00191548 | 0.64199091 | 1 |
| WDR11 | 0.00346713 | 0.64193532 | 1 |
| SRSF3 | 0.00423523 | 0.64143131 | 1 |
| COL12A1 | 0.01440355 | 0.64125343 | 1 |
| AVPR1A | 0.04372446 | 0.64099454 | 1 |
| NGDN | 0.02070485 | 0.64089213 | 1 |
| ABCD1 | 0.00034768 | 0.64008119 | 1 |
| SHLD1 | 0.043715 | 0.64000368 | 1 |
| TARS | 0.00732179 | 0.63976191 | 1 |
| CD248 | 0.01858524 | 0.6397265 | 1 |
| DAPK3 | 0.00173434 | 0.63944741 | 1 |
| VCAM1 | 0.00016833 | 0.6391783 | 1 |
| CAVIN3 | 0.00360568 | 0.63881913 | 1 |

| | | | |
|---|---|---|---|
| GNL3 | 0.00725011 | 0.63863606 | 1 |
| ZGPAT | 0.00187327 | 0.63847096 | 1 |
| ORAI3 | 0.01704454 | 0.63794431 | 1 |
| EIF3D | 0.00101443 | 0.6370372 | 1 |
| TMEM150C | 0.00295535 | 0.63702844 | 1 |
| A1CF | 0.00814368 | 0.6367018 | 1 |
| GPATCH1 | 0.03117379 | 0.63657936 | 1 |
| YY1 | 0.00073878 | 0.63643765 | 1 |
| AKAP8L | 0.00289278 | 0.6363191 | 1 |
| PPIC | 0.00045711 | 0.63611026 | 1 |
| CYP24A1 | 0.00201849 | 0.63605495 | 1 |
| GSE1 | 0.00431597 | 0.63588068 | 1 |
| SCD | 0.0013557 | 0.63537652 | 1 |
| STOML2 | 0.01178951 | 0.63520025 | 1 |
| SHC1 | 0.00088461 | 0.63512128 | 1 |
| CFD | 0.04724006 | 0.6347721 | 1 |
| ACTG1 | 0.0001361 | 0.63460422 | 1 |
| TBX2 | 0.00012055 | 0.63456165 | 1 |
| WDR43 | 0.01074456 | 0.63435362 | 1 |
| KLHL2 | 0.00529306 | 0.63324004 | 1 |
| SEH1L | 0.00874023 | 0.63301244 | 1 |
| ATP1B3 | 0.00151133 | 0.63288076 | 1 |
| ZXDC | 0.00088747 | 0.63273309 | 1 |
| SUPT16H | 0.00188279 | 0.63245373 | 1 |
| MTA2 | 0.00110421 | 0.63239492 | 1 |
| ATG16L1 | 0.02755014 | 0.63207397 | 1 |
| ZNF569 | 0.01593136 | 0.63198059 | 1 |
| SCN2A | 0.0241828 | 0.63163783 | 1 |
| MYSM1 | 0.01987063 | 0.63155957 | 1 |
| FCHSD2 | 0.01808605 | 0.63148633 | 1 |
| NOVA2 | 0.00687382 | 0.63101528 | 1 |
| GPR180 | 0.01269167 | 0.63055063 | 1 |
| GMPPA | 0.03031373 | 0.6304754 | 1 |
| IDNK | 0.03236844 | 0.63042911 | 1 |
| SH3BGRL3 | 0.00054408 | 0.63021922 | 1 |
| SERBP1 | 0.00236845 | 0.63014336 | 1 |
| RTTN | 0.035405 | 0.62994867 | 1 |
| BBC3 | 0.01123446 | 0.62994702 | 1 |
| ARMC5 | 0.00937516 | 0.6298879 | 1 |
| SOX18 | 0.00041977 | 0.62931743 | 1 |
| EMILIN2 | 0.00203961 | 0.6287541 | 1 |
| PMS1 | 0.01896223 | 0.6280274 | 1 |
| RIN3 | 0.00097932 | 0.6279391 | 1 |

| | | | |
|---|---|---|---|
| ARPC1B | 0.00027701 | 0.62742319 | 1 |
| IGIP | 0.01008476 | 0.62720223 | 1 |
| SPRY4 | 0.0005516 | 0.62712903 | 1 |
| CDC23 | 0.00228224 | 0.62591192 | 1 |
| TTC7A | 0.00274994 | 0.62581189 | 1 |
| COX5B | 0.00417374 | 0.62557446 | 1 |
| SLIRP | 0.00066163 | 0.6253087 | 1 |
| TRPS1 | 0.00053472 | 0.62499651 | 1 |
| EXOC3L2 | 0.00350395 | 0.62489946 | 1 |
| LRRC1 | 0.03716369 | 0.62462456 | 1 |
| MFSD6 | 0.00092105 | 0.62394473 | 1 |
| AHI1 | 0.01370281 | 0.62374753 | 1 |
| TMEM184C | 0.00373548 | 0.62367281 | 1 |
| HGS | 0.00584797 | 0.62336402 | 1 |
| C6orf89 | 0.00553038 | 0.62301461 | 1 |
| PRKAR2B | 0.01922388 | 0.62282554 | 1 |
| PGP | 0.00955273 | 0.62269951 | 1 |
| DHX38 | 0.004856 | 0.62259252 | 1 |
| MED16 | 0.00043066 | 0.62202185 | 1 |
| SRSF8 | 0.00070883 | 0.62168946 | 1 |
| MYH11 | 0.00588717 | 0.62155555 | 1 |
| STRN | 0.00814593 | 0.6214317 | 1 |
| FUNDC2 | 0.00837624 | 0.62134906 | 1 |
| MPHOSPH10 | 0.002932 | 0.62134902 | 1 |
| LARS | 0.00129518 | 0.62134358 | 1 |
| MMP28 | 0.00610898 | 0.62127408 | 1 |
| RAI14 | 0.00060882 | 0.62106138 | 1 |
| OLFML1 | 0.00280156 | 0.62104836 | 1 |
| CD47 | 0.0012158 | 0.62053855 | 1 |
| ANO3 | 0.00789204 | 0.61967266 | 1 |
| APOC3 | 0.0473946 | 0.61962106 | 1 |
| GIGYF2 | 0.00045132 | 0.61954327 | 1 |
| RAB8A | 0.00320412 | 0.61951994 | 1 |
| MYO1E | 0.00013364 | 0.6188982 | 1 |
| EEA1 | 0.0003261 | 0.61865808 | 1 |
| TATDN2 | 0.00105078 | 0.61849718 | 1 |
| SYNPO | 0.00044791 | 0.61842747 | 1 |
| RUSC1 | 0.00233645 | 0.61805201 | 1 |
| UNC119B | 0.00558026 | 0.61785602 | 1 |
| EGLN3 | 0.02525518 | 0.6178313 | 1 |
| DOK2 | 0.01685833 | 0.61736412 | 1 |
| ANAPC1 | 0.04730337 | 0.61733457 | 1 |
| SLC29A1 | 0.00216528 | 0.61670144 | 1 |

| | | | |
|---|---|---|---|
| TOR1A | 0.00238911 | 0.61658627 | 1 |
| ADAMTS9 | 0.00025443 | 0.6162272 | 1 |
| NID1 | 0.001 | 0.61563535 | 1 |
| SAMD4A | 0.00564591 | 0.6156132 | 1 |
| CD209 | 0.01397979 | 0.61552511 | 1 |
| USP46 | 0.00617913 | 0.61525995 | 1 |
| VCL | 0.00017345 | 0.61500565 | 1 |
| HOXD11 | 0.02050085 | 0.61491999 | 1 |
| FBXW4 | 0.00075873 | 0.61440344 | 1 |
| NUP188 | 0.00511885 | 0.6140118 | 1 |
| ADAMTS1 | 0.00797653 | 0.61366633 | 1 |
| CLEC11A | 0.00984117 | 0.61287699 | 1 |
| NEK9 | 0.00690338 | 0.61268191 | 1 |
| FLI1 | 0.00568858 | 0.61235311 | 1 |
| PBX3 | 0.02702044 | 0.61178424 | 1 |
| PACRGL | 0.03123077 | 0.61176042 | 1 |
| ZNF268 | 0.01141548 | 0.6115518 | 1 |
| RGS5 | 0.00448413 | 0.61153566 | 1 |
| DCLRE1C | 0.03618228 | 0.61147619 | 1 |
| ZNF483 | 0.02903043 | 0.61108722 | 1 |
| ABLIM2 | 0.00353644 | 0.61028532 | 1 |
| AGGF1 | 0.00044946 | 0.60991215 | 1 |
| SLC22A18 | 0.00670113 | 0.60965786 | 1 |
| OXLD1 | 0.00202923 | 0.60952949 | 1 |
| WT1 | 0.00127278 | 0.60928076 | 1 |
| MAPK13 | 0.0125926 | 0.6085316 | 1 |
| CNOT6 | 0.02907924 | 0.60846012 | 1 |
| TRAM2 | 0.00497679 | 0.60837833 | 1 |
| PITPNM2 | 0.00468596 | 0.60835492 | 1 |
| ABRACL | 0.00041519 | 0.60803903 | 1 |
| SRP72 | 0.01391215 | 0.607937 | 1 |
| CLEC7A | 0.01256848 | 0.6076877 | 1 |
| GLS | 0.0028078 | 0.60761348 | 1 |
| LRRC4B | 0.0169713 | 0.60752309 | 1 |
| DNAJC2 | 0.0346101 | 0.60731522 | 1 |
| ATAT1 | 0.01312302 | 0.60725805 | 1 |
| MCF2L2 | 0.04151007 | 0.60710228 | 1 |
| ATP11A | 0.00739641 | 0.6070657 | 1 |
| SQOR | 0.00221986 | 0.60663928 | 1 |
| NEPRO | 0.00477393 | 0.60654532 | 1 |
| HEG1 | 0.00682618 | 0.6058163 | 1 |
| GNG10 | 0.0199821 | 0.60579832 | 1 |
| CLOCK | 0.00709796 | 0.60573539 | 1 |

| | | | |
|---|---|---|---|
| ARHGEF37 | 0.00493302 | 0.60548308 | 1 |
| GGPS1 | 0.00166299 | 0.60533961 | 1 |
| SPRED1 | 0.00675065 | 0.60532021 | 1 |
| LMAN2L | 0.01396154 | 0.60528936 | 1 |
| PITPNA | 0.00058675 | 0.60471921 | 1 |
| MICAL1 | 0.00205428 | 0.60450423 | 1 |
| GYPC | 0.00123462 | 0.60401624 | 1 |
| GAB1 | 0.00660089 | 0.60326608 | 1 |
| PCMT1 | 0.00232477 | 0.60312825 | 1 |
| GFM1 | 0.0058457 | 0.60305696 | 1 |
| YTHDF2 | 0.0067159 | 0.60302215 | 1 |
| RSPRY1 | 0.00298088 | 0.60293572 | 1 |
| NQO1 | 0.00108266 | 0.60283849 | 1 |
| NRBP1 | 0.00150605 | 0.60269535 | 1 |
| DYM | 0.0018427 | 0.60238981 | 1 |
| ARPC5L | 0.00217525 | 0.60188351 | 1 |
| EDNRA | 0.02795084 | 0.60160055 | 1 |
| PLCG1 | 0.00151394 | 0.6013163 | 1 |
| BCL7A | 0.00716902 | 0.60124389 | 1 |
| FABP4 | 0.00073743 | 0.601125 | 1 |
| UCHL3 | 0.04065212 | 0.60059446 | 1 |
| PLXNA2 | 0.00955328 | 0.60016286 | 1 |
| KIF5B | 0.00125766 | 0.59992528 | 1 |
| DAP | 0.00057819 | 0.59990707 | 1 |
| MBOAT2 | 0.0073674 | 0.59983175 | 1 |
| ZNHIT6 | 0.0245594 | 0.59915895 | 1 |
| MID1IP1 | 0.00348865 | 0.59851709 | 1 |
| TMEM189 | 0.01081183 | 0.59843685 | 1 |
| C12orf73 | 0.03511196 | 0.59800407 | 1 |
| AGO2 | 0.00145918 | 0.59798738 | 1 |
| ZNF703 | 0.00222383 | 0.59751569 | 1 |
| APLP1 | 0.01193459 | 0.59716119 | 1 |
| VEZF1 | 0.00847068 | 0.59685473 | 1 |
| OLFML2B | 0.00919208 | 0.59667637 | 1 |
| PPP2R2A | 0.00241005 | 0.59652025 | 1 |
| ADAMTS19 | 0.00440131 | 0.59589604 | 1 |
| CCNG1 | 0.00682777 | 0.59579401 | 1 |
| PWWP2A | 0.00062574 | 0.59573044 | 1 |
| DPY19L4 | 0.03041877 | 0.59563849 | 1 |
| PGD | 0.0010169 | 0.59549819 | 1 |
| FRK | 0.01198927 | 0.5952956 | 1 |
| REST | 0.00411723 | 0.59527493 | 1 |
| NOL9 | 0.00173743 | 0.59521808 | 1 |

| | | | |
|---|---|---|---|
| SLU7 | 0.0060269 | 0.59513129 | 1 |
| PAGR1 | 0.00093902 | 0.59504139 | 1 |
| IZUMO1 | 0.0473946 | 0.59498791 | 1 |
| ICMT | 0.00012116 | 0.59481757 | 1 |
| ATF4 | 0.00467525 | 0.5946813 | 1 |
| TASOR2 | 0.03037744 | 0.59467918 | 1 |
| C9orf72 | 0.01308137 | 0.59446273 | 1 |
| MIA2 | 0.00226443 | 0.59443293 | 1 |
| RUFY2 | 0.01562721 | 0.59425266 | 1 |
| ZNF664 | 0.00477575 | 0.59401291 | 1 |
| NDNF | 0.00357732 | 0.59356109 | 1 |
| TMED9 | 0.00251651 | 0.59348665 | 1 |
| KL | 0.00267038 | 0.59340865 | 1 |
| IKZF4 | 0.04095313 | 0.59335924 | 1 |
| PRMT7 | 0.00595883 | 0.59242344 | 1 |
| PRKCZ | 0.01510363 | 0.59230466 | 1 |
| PAK1 | 0.00929596 | 0.59230066 | 1 |
| TOP3B | 0.02020393 | 0.5919207 | 1 |
| ENSA | 0.00074721 | 0.59183641 | 1 |
| PRPF39 | 0.00886872 | 0.5916938 | 1 |
| SLC27A4 | 0.00373428 | 0.59149658 | 1 |
| CAVIN1 | 0.00048093 | 0.59122731 | 1 |
| GSTM3 | 0.0098909 | 0.59060289 | 1 |
| AARS2 | 0.0002882 | 0.59031785 | 1 |
| RPF1 | 0.01779886 | 0.59006344 | 1 |
| PDS5B | 0.00830311 | 0.58996612 | 1 |
| NCF2 | 0.00725141 | 0.5899359 | 1 |
| MRI1 | 0.00081629 | 0.58987718 | 1 |
| AASDH | 0.00141955 | 0.58964196 | 1 |
| ELL | 0.00166273 | 0.58923622 | 1 |
| GTF2B | 0.02454959 | 0.58836298 | 1 |
| SNN | 0.00420222 | 0.58827431 | 1 |
| KBTBD2 | 0.00087849 | 0.58818673 | 1 |
| CD36 | 0.02996456 | 0.58814059 | 1 |
| KCNAB2 | 0.0018672 | 0.58799682 | 1 |
| IL1R1 | 0.00221901 | 0.58730616 | 1 |
| S1PR3 | 0.00660101 | 0.58689705 | 1 |
| ARHGAP9 | 0.00491029 | 0.58686424 | 1 |
| WRAP73 | 0.03273626 | 0.58651948 | 1 |
| CDO1 | 0.02568792 | 0.58635493 | 1 |
| COPS8 | 0.00070826 | 0.58615663 | 1 |
| COPS5 | 0.00114667 | 0.58582265 | 1 |
| DSG2 | 0.03729748 | 0.58547402 | 1 |

| | | | |
|---|---|---|---|
| HSF4 | 0.00270897 | 0.58539693 | 1 |
| GSK3A | 0.00351862 | 0.58505816 | 1 |
| RNF11 | 0.00513714 | 0.58502904 | 1 |
| HIST1H2AC | 0.00244993 | 0.58322179 | 1 |
| KANK1 | 0.00274818 | 0.58317265 | 1 |
| CSF2RA | 0.00073085 | 0.58315716 | 1 |
| SNRPA | 0.01657923 | 0.58306703 | 1 |
| SUSD1 | 0.00959708 | 0.58283117 | 1 |
| ENOPH1 | 0.02505368 | 0.58279772 | 1 |
| STRAP | 0.00349453 | 0.58267169 | 1 |
| MAP1S | 0.01064612 | 0.58217685 | 1 |
| PPP2CB | 0.00069684 | 0.58145778 | 1 |
| OXNAD1 | 0.01525347 | 0.58142579 | 1 |
| ARHGEF6 | 0.00714363 | 0.5813515 | 1 |
| NBEAL2 | 0.0086351 | 0.58101204 | 1 |
| RHOJ | 0.03792332 | 0.58096751 | 1 |
| DPM1 | 0.00155247 | 0.58082296 | 1 |
| GSKIP | 0.01717465 | 0.58073467 | 1 |
| OXSR1 | 0.00743188 | 0.58068505 | 1 |
| TMX4 | 0.00057819 | 0.58052607 | 1 |
| C2orf74 | 0.00115047 | 0.58005429 | 1 |
| CCNH | 0.00692218 | 0.57988118 | 1 |
| FOXL1 | 0.01308686 | 0.57948184 | 1 |
| SEPHS1 | 0.00637589 | 0.57938687 | 1 |
| ME2 | 0.00443524 | 0.57842989 | 1 |
| LMO4 | 0.00980868 | 0.57830487 | 1 |
| AR | 0.00571577 | 0.57812427 | 1 |
| ESCO1 | 0.00088703 | 0.57802525 | 1 |
| MAP3K12 | 0.00885949 | 0.57790708 | 1 |
| LRRFIP1 | 0.00228314 | 0.5773091 | 1 |
| ODF2 | 0.00426043 | 0.57723063 | 1 |
| UEVLD | 0.02362299 | 0.57672019 | 1 |
| SERPINB9 | 0.00188611 | 0.57664627 | 1 |
| FOXRED2 | 0.04236844 | 0.57658541 | 1 |
| COA1 | 0.01785894 | 0.57626759 | 1 |
| SPP1 | 0.00034218 | 0.57614603 | 1 |
| LAPTM5 | 0.0005214 | 0.57596987 | 1 |
| PREPL | 0.00214746 | 0.57592439 | 1 |
| IMPACT | 0.00561916 | 0.5755656 | 1 |
| PHF21A | 0.01733429 | 0.57536003 | 1 |
| CDIP1 | 0.01216405 | 0.57506029 | 1 |
| RIOX2 | 0.01195348 | 0.57491299 | 1 |
| ST6GALNAC | 0.00335652 | 0.57476936 | 1 |

| | | | |
|---|---|---|---|
| MAFB | 0.00071945 | 0.57471776 | 1 |
| UBA2 | 0.00441139 | 0.57440826 | 1 |
| GCC2 | 0.00457371 | 0.57419717 | 1 |
| ALG13 | 0.00083095 | 0.57385362 | 1 |
| SMAP1 | 0.01580853 | 0.5737306 | 1 |
| GMDS | 0.03080481 | 0.57328545 | 1 |
| GHR | 0.01276296 | 0.57289463 | 1 |
| GAMT | 0.00809293 | 0.57274841 | 1 |
| KDM1B | 0.01794464 | 0.57261075 | 1 |
| FBXO44 | 0.03336914 | 0.57234235 | 1 |
| ZSCAN20 | 0.02573871 | 0.57189345 | 1 |
| KMT2B | 0.00578336 | 0.57186548 | 1 |
| TXNRD3 | 0.02849411 | 0.57177521 | 1 |
| BECN1 | 0.01757602 | 0.57159154 | 1 |
| PI4K2A | 0.00613372 | 0.57156834 | 1 |
| RAP2C | 0.00214998 | 0.57151469 | 1 |
| SLC40A1 | 0.00568993 | 0.57140908 | 1 |
| LACC1 | 0.03773899 | 0.57134857 | 1 |
| COLGALT1 | 0.00282714 | 0.57120914 | 1 |
| KHDRBS1 | 0.00185047 | 0.57110202 | 1 |
| LMO7 | 0.00251729 | 0.57070332 | 1 |
| SLC25A40 | 0.02091139 | 0.57067684 | 1 |
| MNAT1 | 0.00868135 | 0.57057196 | 1 |
| ACAD11 | 0.00437951 | 0.57053822 | 1 |
| ATAD3A | 0.02687174 | 0.57041858 | 1 |
| GNAI2 | 0.00200548 | 0.57034783 | 1 |
| SEC24C | 0.00146141 | 0.57025274 | 1 |
| EGR1 | 0.01481836 | 0.56996289 | 1 |
| ALDH18A1 | 0.00620861 | 0.56935174 | 1 |
| GLIPR1 | 0.00477616 | 0.56911988 | 1 |
| CTDNEP1 | 0.02343773 | 0.56888099 | 1 |
| CLK1 | 0.00339603 | 0.5688129 | 1 |
| NET1 | 0.00405285 | 0.56867548 | 1 |
| COMMD6 | 0.00266137 | 0.56850037 | 1 |
| TMEM173 | 0.0021061 | 0.56822333 | 1 |
| NOTCH3 | 0.00130582 | 0.56801925 | 1 |
| NPEPPS | 0.00058241 | 0.56792061 | 1 |
| STK38L | 0.00334851 | 0.56767962 | 1 |
| ATPAF1 | 0.00481393 | 0.56747432 | 1 |
| CTSV | 0.00226681 | 0.56665865 | 1 |
| SENP5 | 0.00662519 | 0.5666496 | 1 |
| SFT2D3 | 0.0014859 | 0.56640258 | 1 |
| MARCKS | 0.00079208 | 0.56638338 | 1 |

| | | | |
|---|---|---|---|
| PPRC1 | 0.00134751 | 0.56630023 | 1 |
| BRD2 | 0.00089026 | 0.56620795 | 1 |
| ARL6IP1 | 0.01767194 | 0.56618737 | 1 |
| DSEL | 0.02605186 | 0.5657674 | 1 |
| POLD3 | 0.0104468 | 0.56573155 | 1 |
| HNRNPK | 0.00313813 | 0.56547048 | 1 |
| ZNF823 | 0.00026041 | 0.56512032 | 1 |
| ZC3H7A | 0.02907009 | 0.56510944 | 1 |
| FAM151A | 0.01441506 | 0.56475719 | 1 |
| EPN2 | 0.00986953 | 0.56466764 | 1 |
| MICALL1 | 0.01018123 | 0.5644565 | 1 |
| BPGM | 0.00252385 | 0.56445365 | 1 |
| ZNF655 | 0.00176343 | 0.56407477 | 1 |
| ACE2 | 0.0337303 | 0.56399831 | 1 |
| DNAJC5 | 0.00461333 | 0.56388186 | 1 |
| TMEM179B | 0.01205165 | 0.56291332 | 1 |
| WIPF1 | 0.0010144 | 0.5622922 | 1 |
| FBXO21 | 0.00952457 | 0.56207739 | 1 |
| CD320 | 0.03454774 | 0.56202508 | 1 |
| KLHDC9 | 0.03251768 | 0.56186269 | 1 |
| TEX261 | 0.00867491 | 0.56178121 | 1 |
| TRAPPC6B | 0.01016459 | 0.56156827 | 1 |
| C12orf4 | 0.00457068 | 0.56111897 | 1 |
| ARFIP1 | 0.00359428 | 0.5609933 | 1 |
| POGLUT3 | 0.01677623 | 0.56057412 | 1 |
| DNAH1 | 0.00671642 | 0.56037285 | 1 |
| MKNK1 | 0.00161225 | 0.55991668 | 1 |
| INTS9 | 0.01210536 | 0.55943916 | 1 |
| HEXD | 0.0051062 | 0.55922181 | 1 |
| NECTIN2 | 0.00139698 | 0.55873156 | 1 |
| TXNDC9 | 0.02358357 | 0.55854208 | 1 |
| FAM199X | 0.00101488 | 0.55845025 | 1 |
| TIE1 | 0.00805216 | 0.55842441 | 1 |
| TMEM256 | 0.00085671 | 0.55839671 | 1 |
| C1orf174 | 0.01364245 | 0.5583497 | 1 |
| WWP1 | 0.01984134 | 0.55823594 | 1 |
| LLGL1 | 0.01299556 | 0.5582064 | 1 |
| HDGFL3 | 0.01423845 | 0.55793885 | 1 |
| CHST2 | 0.01242402 | 0.55774377 | 1 |
| POGLUT2 | 0.00662661 | 0.55679321 | 1 |
| CSNK2A1 | 0.04097486 | 0.55676774 | 1 |
| ZNF146 | 0.01355423 | 0.55673163 | 1 |
| TAF6 | 0.00095927 | 0.5566935 | 1 |

| | | | |
|---|---|---|---|
| TTBK2 | 0.01570505 | 0.55600415 | 1 |
| CAV1 | 0.03556324 | 0.55583396 | 1 |
| PCBD1 | 6.83E-05 | 0.55573934 | 1 |
| ARL6 | 0.00067275 | 0.55562543 | 1 |
| EAF2 | 0.00784297 | 0.55560127 | 1 |
| CCDC134 | 0.01632515 | 0.55491062 | 1 |
| SMC1A | 0.00250266 | 0.55470823 | 1 |
| NOP2 | 0.02962345 | 0.55447953 | 1 |
| UBE2G2 | 0.00016574 | 0.55428432 | 1 |
| ACOT8 | 0.01001718 | 0.55421233 | 1 |
| CYP17A1 | 0.00985139 | 0.55414568 | 1 |
| FUCA2 | 0.02060311 | 0.55395818 | 1 |
| ARL4C | 0.03237772 | 0.55355117 | 1 |
| PKDCC | 0.00208912 | 0.55313911 | 1 |
| TRAPPC11 | 0.00236386 | 0.55283907 | 1 |
| MMGT1 | 0.04371457 | 0.55229953 | 1 |
| VIM | 9.21E-05 | 0.55226602 | 1 |
| CPNE2 | 0.02476549 | 0.5521133 | 1 |
| WWC3 | 0.00311038 | 0.55158148 | 1 |
| PRICKLE2 | 0.01346942 | 0.55147836 | 1 |
| HOOK2 | 0.01549854 | 0.55116916 | 1 |
| VHL | 0.00755262 | 0.55115612 | 1 |
| CAV2 | 0.00339999 | 0.55105507 | 1 |
| UBE2V1 | 0.00285996 | 0.55064279 | 1 |
| HADHA | 0.00680471 | 0.55030332 | 1 |
| CASP10 | 0.04895491 | 0.5497459 | 1 |
| RHOBTB3 | 0.00698335 | 0.54903477 | 1 |
| PTAFR | 0.02611081 | 0.54846829 | 1 |
| UBE3C | 0.01293249 | 0.5484544 | 1 |
| PTS | 0.00301596 | 0.54835513 | 1 |
| DAPK2 | 0.0003407 | 0.54812122 | 1 |
| ROBO2 | 0.02431683 | 0.54776938 | 1 |
| JMJD8 | 0.00298089 | 0.547384 | 1 |
| TMEM88 | 0.01448212 | 0.54724996 | 1 |
| REC8 | 0.01746506 | 0.54706525 | 1 |
| WAPL | 0.0103374 | 0.5468524 | 1 |
| PHKG2 | 0.01766379 | 0.54645246 | 1 |
| EMP1 | 0.02271188 | 0.54627576 | 1 |
| HEXIM1 | 0.00212472 | 0.5458277 | 1 |
| NKD2 | 0.02195491 | 0.54564269 | 1 |
| PARVA | 0.00098779 | 0.54561652 | 1 |
| SETDB1 | 0.01365932 | 0.54522678 | 1 |
| LGALS3 | 0.00051924 | 0.5444039 | 1 |

| | | | |
|---|---|---|---|
| PFKP | 0.0161665 | 0.5443786 | 1 |
| MAFG | 0.01564616 | 0.54419669 | 1 |
| MARF1 | 0.01511654 | 0.54373696 | 1 |
| C5AR1 | 0.01450982 | 0.54369411 | 1 |
| PCDHA10 | 0.04158984 | 0.54354232 | 1 |
| SLC5A3 | 0.00489243 | 0.54348925 | 1 |
| UBOX5 | 0.01422077 | 0.54252578 | 1 |
| PPP1R14B | 0.01825055 | 0.5424836 | 1 |
| RASGEF1B | 0.00030555 | 0.54235819 | 1 |
| PLEK | 0.00392163 | 0.54208909 | 1 |
| TMEM120B | 0.02641498 | 0.54143779 | 1 |
| POLH | 0.01196619 | 0.54141557 | 1 |
| SYAP1 | 0.00109287 | 0.54066816 | 1 |
| EIF5B | 0.01560729 | 0.54059475 | 1 |
| RCBTB1 | 0.00385278 | 0.54054731 | 1 |
| RSAD1 | 0.01297253 | 0.54043606 | 1 |
| SEPTIN4 | 0.00013632 | 0.54021932 | 1 |
| NCBP1 | 0.01182738 | 0.53967616 | 1 |
| SH3BGRL | 0.00059941 | 0.53908952 | 1 |
| SFXN4 | 0.02372065 | 0.53899935 | 1 |
| MXRA8 | 0.00072595 | 0.53884214 | 1 |
| PFDN5 | 0.00436224 | 0.53872595 | 1 |
| RNF19B | 0.01962231 | 0.53848138 | 1 |
| ARL13B | 0.00267317 | 0.53840798 | 1 |
| ACE | 0.01970605 | 0.53817753 | 1 |
| MED4 | 0.00379749 | 0.5381755 | 1 |
| CWC27 | 0.02765047 | 0.53806174 | 1 |
| BAZ2A | 0.00017378 | 0.53782305 | 1 |
| C1QA | 0.00590066 | 0.53780244 | 1 |
| ABCA7 | 0.01136475 | 0.53763902 | 1 |
| LUM | 0.01012634 | 0.53729604 | 1 |
| LCP1 | 0.00129827 | 0.5369195 | 1 |
| MTHFD1 | 0.01006053 | 0.53686793 | 1 |
| BCKDHA | 0.00726427 | 0.53664886 | 1 |
| CERCAM | 0.03227788 | 0.53654992 | 1 |
| TMEM50A | 0.00264094 | 0.53651771 | 1 |
| DLGAP4 | 0.00072012 | 0.53613918 | 1 |
| HM13 | 0.00132394 | 0.53608891 | 1 |
| HOXC8 | 0.03299647 | 0.535907 | 1 |
| PPP1R9B | 0.00015436 | 0.5358959 | 1 |
| RRS1 | 0.03794047 | 0.53589197 | 1 |
| RAB4B | 0.02193529 | 0.53575915 | 1 |
| HOXC5 | 0.03813317 | 0.53567931 | 1 |

| | | | |
|---|---|---|---|
| CACNB3 | 0.04695616 | 0.53552412 | 1 |
| NPHS2 | 0.00241471 | 0.53485593 | 1 |
| CEBPZOS | 0.00738875 | 0.53480697 | 1 |
| CCDC71 | 0.03282222 | 0.53446074 | 1 |
| CD46 | 0.00163653 | 0.53441189 | 1 |
| ITGA3 | 0.00032191 | 0.53427051 | 1 |
| PLXNA1 | 0.02042995 | 0.53382878 | 1 |
| ANKRD52 | 0.04048979 | 0.53333416 | 1 |
| DNASE1L1 | 0.01290041 | 0.53325817 | 1 |
| THUMPD3 | 0.04949203 | 0.53299525 | 1 |
| COG1 | 0.01387615 | 0.53290608 | 1 |
| NRG3 | 0.00975819 | 0.5327143 | 1 |
| CD164 | 0.00363205 | 0.53252089 | 1 |
| METAP2 | 0.01049486 | 0.53197408 | 1 |
| RNGTT | 0.00724039 | 0.53193354 | 1 |
| GAS1 | 0.00035217 | 0.53083634 | 1 |
| NR2C1 | 0.02899073 | 0.53030555 | 1 |
| SACM1L | 0.00588799 | 0.5301397 | 1 |
| EXOSC9 | 0.00639433 | 0.52940688 | 1 |
| ACSL4 | 0.01421526 | 0.52898703 | 1 |
| PYROXD2 | 0.0470567 | 0.52892101 | 1 |
| ZNF660 | 0.0473946 | 0.52885059 | 1 |
| LOXL2 | 0.02051517 | 0.52839315 | 1 |
| ZC3HAV1 | 0.01422261 | 0.528063 | 1 |
| STN1 | 0.00121381 | 0.527968 | 1 |
| FAM168A | 0.01070205 | 0.5279568 | 1 |
| DOCK8 | 0.03041669 | 0.52766281 | 1 |
| FAM81A | 0.00405527 | 0.52763348 | 1 |
| ALYREF | 0.03203905 | 0.52759392 | 1 |
| NUDT21 | 0.0018555 | 0.52737153 | 1 |
| STX11 | 0.00166273 | 0.52700283 | 1 |
| MPEG1 | 0.01922623 | 0.52695095 | 1 |
| MED13 | 0.01563514 | 0.52675567 | 1 |
| ZBTB2 | 0.03581181 | 0.52665705 | 1 |
| ABCA2 | 0.00166204 | 0.52648505 | 1 |
| GSPT1 | 0.02342499 | 0.52627022 | 1 |
| KDM4B | 0.00290456 | 0.52589383 | 1 |
| SPG21 | 0.00470956 | 0.52581351 | 1 |
| PHETA2 | 0.00719694 | 0.52572495 | 1 |
| PVR | 0.00423012 | 0.52551776 | 1 |
| C19orf53 | 0.01917422 | 0.52509797 | 1 |
| ZNF169 | 0.00907706 | 0.52438061 | 1 |
| PIGT | 0.01720473 | 0.52427222 | 1 |

| | | | |
|---|---|---|---|
| TMED1 | 0.00711193 | 0.52419466 | 1 |
| SLC41A3 | 0.02675655 | 0.5240992 | 1 |
| LAMC1 | 0.00347832 | 0.52395931 | 1 |
| ALPK2 | 0.00344834 | 0.52378068 | 1 |
| SMDT1 | 0.01605175 | 0.5236632 | 1 |
| CLCN4 | 0.03691756 | 0.52359772 | 1 |
| PAPOLA | 0.00982348 | 0.52328595 | 1 |
| TNKS2 | 0.00916451 | 0.52234696 | 1 |
| LBH | 0.00601503 | 0.5222427 | 1 |
| UCP1 | 0.02939401 | 0.52208224 | 1 |
| CSF2RB | 0.04211656 | 0.52191946 | 1 |
| CYP1B1 | 0.01438633 | 0.52174857 | 1 |
| ALS2CL | 0.00347026 | 0.52170795 | 1 |
| HDHD2 | 0.01257348 | 0.52120266 | 1 |
| PTPN9 | 0.01041434 | 0.52098949 | 1 |
| ZFP36L1 | 0.00116919 | 0.5209763 | 1 |
| TNFRSF25 | 0.01620917 | 0.52087127 | 1 |
| C19orf47 | 0.0132247 | 0.5205167 | 1 |
| ANKLE2 | 0.00040753 | 0.51969232 | 1 |
| NDST1 | 0.00334843 | 0.51966299 | 1 |
| ECE1 | 0.00088282 | 0.51905445 | 1 |
| MAPK6 | 0.01033809 | 0.51811968 | 1 |
| KCNE4 | 0.01611126 | 0.51788651 | 1 |
| LARP1 | 0.01302821 | 0.517785 | 1 |
| TUSC2 | 0.00653838 | 0.51775801 | 1 |
| ZNF44 | 0.00858203 | 0.51691864 | 1 |
| TOPBP1 | 0.02696485 | 0.5168425 | 1 |
| CLIC1 | 0.00100632 | 0.51593213 | 1 |
| NPC2 | 7.35E-05 | 0.51591568 | 1 |
| COPZ2 | 0.03960283 | 0.5154985 | 1 |
| IPO9 | 0.00125213 | 0.51542939 | 1 |
| TXNDC11 | 0.00271265 | 0.5154079 | 1 |
| GFM2 | 0.03756657 | 0.51531249 | 1 |
| LYPLAL1 | 0.00512345 | 0.51502196 | 1 |
| ACAP3 | 0.02870457 | 0.51496984 | 1 |
| CAPNS1 | 0.00569534 | 0.5146652 | 1 |
| SS18L1 | 0.01486181 | 0.51423451 | 1 |
| MAP2K3 | 0.00298764 | 0.51418317 | 1 |
| ALDH7A1 | 0.00417647 | 0.51364343 | 1 |
| RNF7 | 0.00053479 | 0.51310689 | 1 |
| ACTB | 0.0010903 | 0.51234404 | 1 |
| SSR2 | 0.00022759 | 0.51198821 | 1 |
| SH3RF3 | 0.00117317 | 0.51171725 | 1 |

| | | | |
|---|---|---|---|
| ARMT1 | 0.02485843 | 0.51169606 | 1 |
| ESD | 0.01763473 | 0.51128884 | 1 |
| RAB39A | 0.04090519 | 0.51123991 | 1 |
| TTC12 | 0.01769224 | 0.51114717 | 1 |
| COP1 | 0.01988047 | 0.51089946 | 1 |
| PLA2G4B | 0.0265354 | 0.51046697 | 1 |
| PEMT | 0.00604241 | 0.51020169 | 1 |
| SUPT7L | 0.0089242 | 0.50976731 | 1 |
| GNG2 | 0.00478004 | 0.50968436 | 1 |
| TSG101 | 0.02735091 | 0.50952296 | 1 |
| VSIG4 | 0.0024754 | 0.50944098 | 1 |
| YWHAH | 0.00188272 | 0.50939353 | 1 |
| AKIP1 | 0.02548167 | 0.5093517 | 1 |
| MAGED1 | 0.00084665 | 0.50933628 | 1 |
| CACUL1 | 0.00226443 | 0.50930741 | 1 |
| TWSG1 | 0.00799408 | 0.50922495 | 1 |
| DCTN2 | 0.00046425 | 0.50910803 | 1 |
| LDLRAP1 | 0.00469266 | 0.50865335 | 1 |
| SH3BP5L | 0.01149097 | 0.50861379 | 1 |
| ANXA2 | 0.00076383 | 0.50857815 | 1 |
| SLC22A13 | 0.00670942 | 0.50853924 | 1 |
| RPRD2 | 0.03551005 | 0.50797545 | 1 |
| SPA17 | 0.01485704 | 0.50772132 | 1 |
| PTPN18 | 0.02431683 | 0.50770206 | 1 |
| CHCHD7 | 0.00992247 | 0.50762789 | 1 |
| YIF1B | 0.04702973 | 0.50734973 | 1 |
| AGTRAP | 0.00501537 | 0.50694268 | 1 |
| MLLT6 | 0.00137507 | 0.50677535 | 1 |
| NXT2 | 0.01815002 | 0.50653408 | 1 |
| USP10 | 0.01816515 | 0.5064961 | 1 |
| LRRC19 | 0.04570994 | 0.50613647 | 1 |
| ATR | 0.00576275 | 0.5055449 | 1 |
| PSMA3 | 0.03099388 | 0.50542895 | 1 |
| PAF1 | 0.00438055 | 0.50511643 | 1 |
| RANBP10 | 0.00346015 | 0.50436984 | 1 |
| MAP4K4 | 0.01694242 | 0.50416712 | 1 |
| PBDC1 | 0.03304434 | 0.50404427 | 1 |
| ANKIB1 | 0.00692227 | 0.50399176 | 1 |
| ARHGAP18 | 0.00107107 | 0.50373919 | 1 |
| PSPC1 | 0.01541751 | 0.50369481 | 1 |
| KARS | 0.01332862 | 0.50341599 | 1 |
| MEX3D | 0.01252281 | 0.50320113 | 1 |
| SLC37A4 | 0.00868395 | 0.50318147 | 1 |

| | | | |
|---|---|---|---|
| VRK3 | 0.00470276 | 0.50307377 | 1 |
| FEM1B | 9.79E-05 | 0.50265783 | 1 |
| GLB1L | 0.02568792 | 0.502585 | 1 |
| SP3 | 0.01001428 | 0.50232623 | 1 |
| TMEM147 | 0.00626933 | 0.50219155 | 1 |
| CSGALNACT | 0.01813839 | 0.50214042 | 1 |
| MRC2 | 0.04911956 | 0.5017794 | 1 |
| TANGO2 | 0.00300294 | 0.50145248 | 1 |
| RBM6 | 0.00512711 | 0.50144247 | 1 |
| SGF29 | 0.03360786 | 0.50142534 | 1 |
| MGP | 0.00465543 | 0.50123092 | 1 |
| SLC22A12 | 0.00226311 | 0.50122579 | 1 |
| CHRD | 0.00016379 | 0.50097338 | 1 |
| ATF2 | 0.00422245 | 0.50091693 | 1 |
| MDM2 | 0.01281912 | 0.50037913 | 1 |
| LAMTOR3 | 0.00315021 | 0.50026748 | 1 |
| ACSF3 | 0.00238911 | 0.50001881 | 1 |
| ENY2 | 0.00607977 | 0.50000942 | 1 |
| PLK1 | 0.04775027 | -0.5036916 | 1 |
| SAT1 | 0.00399679 | -0.5054076 | 1 |
| H6PD | 0.02277416 | -0.5064011 | 1 |
| LILRA2 | 0.01436996 | -0.5066953 | 1 |
| MACO1 | 0.0012652 | -0.5076007 | 1 |
| DENR | 0.02795084 | -0.5144619 | 1 |
| SIRT3 | 0.03200691 | -0.5188922 | 1 |
| TPGS1 | 0.04293446 | -0.5460121 | 1 |
| PLD1 | 0.02450323 | -0.5460454 | 1 |
| ZNF16 | 0.04303198 | -0.5554995 | 1 |
| RET | 0.03493522 | -0.5602956 | 1 |
| CYB5A | 0.03943396 | -0.5604678 | 1 |
| ELK1 | 0.01984693 | -0.5622062 | 1 |
| PLGRKT | 0.02227622 | -0.5710816 | 1 |
| MFHAS1 | 0.04701898 | -0.5741154 | 1 |
| GALNT7 | 0.01887176 | -0.5758619 | 1 |
| DCTN6 | 0.04252438 | -0.5830345 | 1 |
| NRF1 | 0.02107032 | -0.5911732 | 1 |
| WDR3 | 0.01715601 | -0.5975079 | 1 |
| ZW10 | 0.04861922 | -0.6035198 | 1 |
| COQ8A | 0.03010468 | -0.6079928 | 1 |
| ISG20 | 0.01423845 | -0.6091734 | 1 |
| ERG28 | 0.03922949 | -0.6232537 | 1 |
| PLAT | 0.01560339 | -0.628667 | 1 |
| PALM3 | 0.03369465 | -0.6340092 | 1 |

| | | | |
|---|---|---|---|
| DHRS7B | 0.02888311 | -0.6368254 | 1 |
| CKAP5 | 0.00058848 | -0.639226 | 1 |
| TSTA3 | 0.01248423 | -0.6434608 | 1 |
| RNASE4 | 0.00228137 | -0.6440715 | 1 |
| RNLS | 0.0470567 | -0.6461619 | 1 |
| NDUFA6 | 0.00672133 | -0.6464635 | 1 |
| PCGF3 | 0.02237363 | -0.6496746 | 1 |
| DGCR8 | 0.04469253 | -0.6499603 | 1 |
| CHRM3 | 0.0496991 | -0.6554738 | 1 |
| TLE3 | 0.02233734 | -0.6588716 | 1 |
| SPIN4 | 0.04375981 | -0.6616267 | 1 |
| RARG | 0.0310878 | -0.6646545 | 1 |
| SYNE1 | 0.00084151 | -0.6748885 | 1 |
| ZBTB16 | 0.01937217 | -0.6815733 | 1 |
| HEYL | 0.01423611 | -0.6922051 | 1 |
| ZNF438 | 0.0048582 | -0.693829 | 1 |
| FBXO38 | 0.03855807 | -0.6971145 | 1 |
| NDRG3 | 0.04908377 | -0.6982605 | 1 |
| UBIAD1 | 0.04097486 | -0.7011282 | 1 |
| SLC12A2 | 0.04211621 | -0.7144017 | 1 |
| P2RY13 | 0.04611249 | -0.733473 | 1 |
| KDELR1 | 0.04362436 | -0.7406615 | 1 |
| HABP2 | 0.02756626 | -0.767942 | 1 |
| SDHAF4 | 0.04185864 | -0.771742 | 1 |
| ARMCX6 | 0.00850022 | -0.7762681 | 1 |
| HSPA4L | 0.00855629 | -0.7825517 | 1 |
| MEAK7 | 0.01938057 | -0.7995996 | 1 |
| EPOP | 0.04082339 | -0.825885 | 1 |
| ZBTB26 | 0.00720505 | -0.8296398 | 1 |
| GPANK1 | 0.03289954 | -0.8340402 | 1 |
| CNBP | 0.002807 | -0.8350022 | 1 |
| PXMP4 | 0.02266694 | -0.8449737 | 1 |
| HSD11B2 | 0.01287745 | -0.8560668 | 1 |
| SVBP | 0.01252775 | -0.8609945 | 1 |
| OTUD7B | 0.03437586 | -0.8766677 | 1 |
| CDK2AP2 | 0.0232337 | -0.8863926 | 1 |
| RIIAD1 | 0.02939401 | -0.8933031 | 1 |
| RABEPK | 0.04372738 | -0.8949431 | 1 |
| NFYB | 0.03418015 | -0.9146634 | 1 |
| SOX7 | 0.02116617 | -0.9215679 | 1 |
| VPS18 | 0.04239403 | -0.9458148 | 1 |
| SOWAHD | 0.0417333 | -0.963136 | 1 |
| OR2T10 | 0.02352405 | -0.9802574 | 1 |

| | | | |
|---|---|---|---|
| ZFP30 | 0.04283826 | -0.9894203 | 1 |
| KLF9 | 0.0032348 | -1.0107527 | 1 |
| BCDIN3D | 0.03640275 | -1.0588007 | 1 |
| RNF2 | 0.02073347 | -1.094659 | 1 |
| GPR34 | 0.04232015 | -1.1604876 | 1 |
| EPCAM | 0.00966354 | -1.1715262 | 1 |
| SLC7A6 | 0.02133996 | -1.2803542 | 1 |
| TM2D2 | 0.02949809 | -1.3184527 | 1 |
| PPP2R2B | 0.04721253 | -1.3671961 | 1 |
| KRT19 | 0.01087302 | -1.7118316 | 1 |
| SATB2 | 0.03565524 | -1.8759845 | 1 |
| ANKRD66 | 0.04609245 | -1.8804868 | 1 |
| RFX8 | 0.01927611 | -1.9580787 | 1 |
| CRTAM | 0.04886488 | -1.9823445 | 1 |
| TBX19 | 0.02565142 | -1.9830131 | 1 |
| NPVF | 0.03754192 | -1.9936118 | 1 |
| LGALS16 | 0.03889679 | -2.0518444 | 1 |
| SLC16A13 | 0.03378795 | -2.0742751 | 1 |
| RFX4 | 0.04624818 | -2.0785022 | 1 |
| HTR3D | 0.04029296 | -2.084097 | 1 |
| ADRB1 | 0.01836627 | -2.1186928 | 1 |
| IGHG3 | 0.01803627 | -2.1545512 | 1 |
| SIX1 | 0.03862973 | -2.1606396 | 1 |
| ASMT | 0.04746699 | -2.1851079 | 1 |
| LYRM2 | 0.01418627 | -2.2973686 | 1 |
| PTPRH | 0.03754192 | -2.3036956 | 1 |
| KCNJ6 | 0.0152724 | -2.3084812 | 1 |
| KRT82 | 0.04624818 | -2.3203039 | 1 |
| COL9A1 | 0.03754192 | -2.3649342 | 1 |
| CTRC | 0.03889679 | -2.4010409 | 1 |
| ATP5PO | 0.04507161 | -2.4278859 | 1 |
| MAGEC2 | 0.03889679 | -2.4485451 | 1 |
| ASCL3 | 0.03754192 | -2.4869118 | 1 |
| LOR | 0.00734614 | -2.5992736 | 1 |
| ERICH5 | 0.03993018 | -2.6473599 | 1 |
| IGHA2 | 0.02087372 | -2.6509321 | 1 |
| GNG4 | 0.02087372 | -2.6528816 | 1 |
| SPATA32 | 0.0394759 | -2.6580261 | 1 |
| TP53AIP1 | 0.03622743 | -2.6582677 | 1 |
| SLC18A3 | 0.0438895 | -2.6630604 | 1 |
| KCNJ13 | 0.04624818 | -2.6815439 | 1 |
| C22orf23 | 0.02087372 | -2.6829125 | 1 |
| HSD3B1 | 0.02087372 | -2.7225525 | 1 |

| | | | |
|---|---|---|---|
| FSD1 | 0.03889679 | -2.7952753 | 1 |
| TG | 0.03862973 | -2.8298534 | 1 |
| MORC1 | 0.04345369 | -2.859419 | 1 |
| CCDC87 | 0.0409437 | -2.9343235 | 1 |
| CACNG6 | 0.0421829 | -2.9926842 | 1 |
| GPRC5D | 0.03754192 | -3.0465303 | 1 |
| TAFA3 | 0.00492061 | -3.1405329 | 1 |
| CCK | 0.0421829 | -3.2353984 | 1 |
| KIF24 | 0.01316886 | -3.4057127 | 1 |
| GFI1 | 0.0421829 | -3.4630522 | 1 |
| SH2D1B | 0.04345369 | -3.6060867 | 1 |
| GPR156 | 0.00734614 | -3.7006034 | 1 |
| SPZ1 | 0.04345369 | -3.719725 | 1 |
| LOXHD1 | 0.03622743 | -3.7392221 | 1 |
| OR6M1 | 0.00092557 | -3.787215 | 1 |
| CACNG3 | 0.01316886 | -3.8531616 | 1 |
| TMEM221 | 0.01366921 | -3.9464552 | 1 |
| GRM8 | 0.03622743 | -3.9594685 | 1 |
| PRDM9 | 0.03622743 | -3.9890571 | 1 |
| TSSK1B | 0.01316886 | -4.0970273 | 1 |
| ZIC5 | 0.0409437 | -4.1511897 | 1 |
| C5orf67 | 0.02087372 | -5.1156887 | 1 |

### Supplemental Table S3

**Differential Gene Expression Analysis Class III LN glomeruli vs Class V LN glomeruli**

Differential gene expression analysis performed using FindMarkers comparing gene expression in 55uM spots from all Class III LN glomeruli to Class V LN glomeruli. For exploratory analysis log2FC>|0.5|, p <0.05 was included in the list of differentially expressed genes.

| | p_val | avg_log2FC | p_val_adj |
|---|---|---|---|
| CD163 | 9.48E-23 | 4.11779456 | 1.71E-18 |
| C1QC | 5.92E-22 | 2.67399237 | 1.07E-17 |
| C1QB | 3.28E-19 | 2.73164593 | 5.94E-15 |
| FKBP5 | 1.56E-18 | 2.99564086 | 2.81E-14 |
| C1QA | 8.97E-18 | 2.54513305 | 1.62E-13 |
| BCAM | 1.62E-15 | -0.9427351 | 2.93E-11 |
| SPI1 | 7.04E-15 | 1.88854602 | 1.27E-10 |
| FCER1G | 1.15E-14 | 2.30017974 | 2.07E-10 |
| FCGR3A | 2.82E-14 | 3.2117117 | 5.10E-10 |
| LIPA | 5.93E-14 | 2.07202513 | 1.07E-09 |
| SRGN | 1.22E-13 | 1.50456326 | 2.20E-09 |
| TIMP3 | 1.48E-13 | 1.15566846 | 2.67E-09 |
| IFI30 | 1.48E-13 | 2.00279901 | 2.67E-09 |
| CD74 | 1.64E-13 | 0.70285362 | 2.96E-09 |
| CSF1R | 1.73E-13 | 1.63419571 | 3.13E-09 |
| FABP4 | 3.17E-13 | 1.76344915 | 5.73E-09 |
| IGFBP5 | 4.11E-13 | -0.8521109 | 7.43E-09 |
| ERRFI1 | 4.94E-13 | 2.17443901 | 8.94E-09 |
| MS4A7 | 8.25E-13 | 2.09705678 | 1.49E-08 |
| ITGB2 | 9.81E-13 | 2.02782322 | 1.77E-08 |
| CD53 | 1.67E-12 | 4.30156687 | 3.01E-08 |
| FTH1 | 1.81E-12 | 0.78450539 | 3.28E-08 |
| CTSS | 3.39E-12 | 1.66299774 | 6.12E-08 |
| PLCG2 | 4.03E-12 | 1.88592867 | 7.29E-08 |
| ZBTB16 | 4.07E-12 | 2.18607286 | 7.36E-08 |
| CD14 | 4.29E-12 | 1.61424491 | 7.75E-08 |
| HK3 | 5.74E-12 | 3.16023009 | 1.04E-07 |
| SERPING1 | 1.33E-11 | 1.3105491 | 2.41E-07 |
| SYNE1 | 1.41E-11 | 1.07057015 | 2.55E-07 |
| LCP1 | 1.57E-11 | 1.60276198 | 2.83E-07 |
| ITGAX | 1.79E-11 | 1.94558255 | 3.25E-07 |
| CTSB | 1.95E-11 | 0.83925035 | 3.53E-07 |
| SPOCK1 | 2.39E-11 | -1.1513422 | 4.32E-07 |
| MS4A6A | 2.61E-11 | 1.72199961 | 4.73E-07 |
| MAOB | 4.08E-11 | 2.28658032 | 7.39E-07 |
| TYROBP | 4.52E-11 | 1.78552403 | 8.17E-07 |
| MSR1 | 6.09E-11 | 1.5032529 | 1.10E-06 |
| IRAK3 | 6.41E-11 | 1.80042346 | 1.16E-06 |
| SPARC | 7.27E-11 | -0.7438394 | 1.32E-06 |
| ARRB2 | 8.64E-11 | 1.57173853 | 1.56E-06 |
| LAPTM5 | 1.01E-10 | 1.64845123 | 1.82E-06 |

| | | | |
|---|---|---|---|
| PTGDS | 1.24E-10 | -1.0659532 | 2.24E-06 |
| EPB41L3 | 1.29E-10 | 1.21183825 | 2.33E-06 |
| LILRB2 | 1.31E-10 | 2.28062052 | 2.38E-06 |
| C3AR1 | 1.34E-10 | 3.42170686 | 2.42E-06 |
| CORO1A | 1.37E-10 | 1.33768758 | 2.47E-06 |
| DEPP1 | 2.55E-10 | 1.81951574 | 4.60E-06 |
| VSIG4 | 2.74E-10 | 2.50456988 | 4.95E-06 |
| CEBPD | 2.87E-10 | 1.48881538 | 5.18E-06 |
| TLR4 | 3.47E-10 | 1.85023805 | 6.28E-06 |
| PLEK | 3.51E-10 | 3.68442026 | 6.35E-06 |
| PLAUR | 3.80E-10 | 2.69299267 | 6.87E-06 |
| MKLN1 | 4.09E-10 | 1.60283023 | 7.40E-06 |
| UACA | 4.13E-10 | -1.0737662 | 7.47E-06 |
| CRYAB | 4.19E-10 | 1.63067661 | 7.58E-06 |
| MT2A | 5.23E-10 | 1.2323397 | 9.46E-06 |
| EMILIN2 | 5.59E-10 | 1.59834995 | 1.01E-05 |
| CST3 | 5.81E-10 | 0.80630027 | 1.05E-05 |
| BAHCC1 | 5.84E-10 | 2.24252866 | 1.06E-05 |
| HCLS1 | 5.92E-10 | 1.34825591 | 1.07E-05 |
| HMOX1 | 6.28E-10 | 1.4262227 | 1.14E-05 |
| C5AR1 | 6.29E-10 | 2.45790075 | 1.14E-05 |
| GRHPR | 6.39E-10 | 1.86701024 | 1.15E-05 |
| TYMP | 6.95E-10 | 1.68833584 | 1.26E-05 |
| CEBPB | 7.11E-10 | 1.27309873 | 1.29E-05 |
| A2M | 7.86E-10 | 0.76749749 | 1.42E-05 |
| ADCY3 | 9.11E-10 | 1.4390643 | 1.65E-05 |
| SHMT1 | 1.04E-09 | 1.4322501 | 1.87E-05 |
| CORO1C | 1.16E-09 | 1.54505424 | 2.10E-05 |
| PIK3AP1 | 1.20E-09 | 1.86838523 | 2.18E-05 |
| CD300A | 1.31E-09 | 3.85589893 | 2.36E-05 |
| NNMT | 1.33E-09 | 2.66126957 | 2.40E-05 |
| TAGLN | 1.33E-09 | 1.38620407 | 2.40E-05 |
| ANKS1A | 1.43E-09 | 1.28612982 | 2.58E-05 |
| INMT | 1.47E-09 | 1.40923587 | 2.65E-05 |
| PLAT | 1.53E-09 | 1.08211009 | 2.76E-05 |
| RUNX1 | 1.80E-09 | 2.52979349 | 3.26E-05 |
| LYN | 2.16E-09 | 1.35329399 | 3.91E-05 |
| NPNT | 2.20E-09 | -0.9398035 | 3.97E-05 |
| ADAMTS2 | 2.27E-09 | 1.85453378 | 4.11E-05 |
| FGD2 | 2.94E-09 | 2.50324219 | 5.32E-05 |
| DPEP1 | 2.98E-09 | 1.47645523 | 5.40E-05 |
| HIF3A | 3.01E-09 | 1.53378426 | 5.45E-05 |
| S100A1 | 3.31E-09 | 2.21864694 | 5.98E-05 |
| RNF152 | 4.46E-09 | 1.51712572 | 8.06E-05 |
| SLC11A1 | 4.55E-09 | 2.66997748 | 8.22E-05 |
| CD68 | 4.71E-09 | 1.29550645 | 8.51E-05 |
| MMADHC | 4.82E-09 | 2.62629581 | 8.72E-05 |
| NDNF | 4.90E-09 | -0.9737156 | 8.86E-05 |
| CALML3 | 5.93E-09 | 2.41420353 | 0.00010732 |
| COX11 | 6.40E-09 | 0.81633561 | 0.00011582 |
| CYTH4 | 6.57E-09 | 1.65282549 | 0.00011876 |
| GPAT3 | 6.68E-09 | 0.97695262 | 0.00012087 |
| LRRC25 | 6.71E-09 | 3.30975096 | 0.00012129 |

| | | | |
|---|---|---|---|
| SPOCK2 | 6.80E-09 | -0.8199089 | 0.00012294 |
| CD36 | 7.04E-09 | 2.2324617 | 0.00012739 |
| TPPP3 | 7.07E-09 | -1.2178581 | 0.0001279 |
| CCDC88A | 7.57E-09 | 1.22050392 | 0.00013696 |
| CWC15 | 8.56E-09 | 1.02727365 | 0.00015488 |
| CPVL | 9.87E-09 | 1.5255612 | 0.00017847 |
| SWAP70 | 9.89E-09 | 1.07549761 | 0.00017879 |
| ITGA3 | 1.15E-08 | -0.9324932 | 0.00020873 |
| RAD23A | 1.21E-08 | 1.08658279 | 0.00021897 |
| PTPRO | 1.37E-08 | -1.0583545 | 0.00024797 |
| RDH14 | 1.40E-08 | 1.21668843 | 0.00025301 |
| CXCL16 | 1.49E-08 | 0.96888178 | 0.00026904 |
| PLA2R1 | 1.49E-08 | -1.0417735 | 0.00026907 |
| TLE2 | 1.57E-08 | 1.11985029 | 0.00028449 |
| MTIF2 | 1.80E-08 | 1.88374956 | 0.00032555 |
| SLC15A3 | 1.99E-08 | 1.22872598 | 0.00036031 |
| PCOLCE2 | 2.08E-08 | -1.1951319 | 0.00037566 |
| CYP24A1 | 2.11E-08 | 3.24738714 | 0.00038143 |
| SEC61B | 2.37E-08 | 2.09489004 | 0.00042868 |
| CYP20A1 | 2.65E-08 | 1.00408437 | 0.00047871 |
| HOXB4 | 2.69E-08 | 1.19594367 | 0.00048571 |
| C1orf162 | 2.77E-08 | 2.44495216 | 0.00050046 |
| SDC1 | 3.18E-08 | 1.87449559 | 0.00057532 |
| ANAPC7 | 3.18E-08 | 1.52438527 | 0.00057532 |
| FAM118A | 3.20E-08 | 2.16462742 | 0.00057948 |
| PLPBP | 3.29E-08 | 0.81300037 | 0.00059429 |
| LAIR1 | 3.35E-08 | 2.41869422 | 0.00060535 |
| DCN | 3.39E-08 | -0.7403096 | 0.00061304 |
| CDKN1A | 3.55E-08 | 1.94790387 | 0.00064135 |
| ALB | 3.56E-08 | 2.13197369 | 0.000643 |
| IKZF1 | 3.84E-08 | 1.52175604 | 0.00069454 |
| SH3BGRL3 | 3.84E-08 | 0.9251362 | 0.00069525 |
| SMAP2 | 3.94E-08 | 1.23185291 | 0.00071215 |
| ECHDC1 | 3.95E-08 | 1.00280297 | 0.00071397 |
| DLST | 4.09E-08 | 1.54958451 | 0.0007398 |
| ARHGAP10 | 4.57E-08 | 1.77811458 | 0.00082657 |
| ZFP36 | 4.57E-08 | 1.01120551 | 0.00082697 |
| KCNAB2 | 4.81E-08 | 1.23220397 | 0.0008692 |
| ARPC4 | 4.85E-08 | 1.10481937 | 0.0008764 |
| ACY3 | 5.13E-08 | 2.00041424 | 0.00092768 |
| C7 | 5.46E-08 | 0.99779896 | 0.00098692 |
| RAC2 | 5.55E-08 | 2.69319442 | 0.00100361 |
| SAT1 | 5.64E-08 | 0.88250552 | 0.00101961 |
| KLHL21 | 6.04E-08 | 1.08316725 | 0.00109194 |
| NRGN | 6.08E-08 | -1.2887519 | 0.00109951 |
| PDK4 | 6.15E-08 | 1.30468039 | 0.00111195 |
| NAMPT | 6.16E-08 | 0.97543515 | 0.00111399 |
| FABP5 | 6.40E-08 | 2.21195494 | 0.00115817 |
| FAM107A | 6.55E-08 | 1.67534687 | 0.00118497 |
| JUN | 6.80E-08 | 1.0760138 | 0.00123037 |
| HTRA1 | 7.17E-08 | -0.9052449 | 0.00129651 |
| TRPC4AP | 7.40E-08 | 0.56064114 | 0.00133824 |
| SLC12A6 | 8.14E-08 | 1.37926875 | 0.00147151 |

| | | | |
|---|---|---|---|
| APOD | 8.19E-08 | 1.97934635 | 0.00148129 |
| NTN4 | 8.31E-08 | 0.75079255 | 0.0015025 |
| TNFAIP3 | 8.64E-08 | 0.56425438 | 0.00156272 |
| CD302 | 8.82E-08 | 2.11234026 | 0.00159441 |
| IL1R1 | 8.95E-08 | 0.94679601 | 0.00161927 |
| DOK2 | 9.23E-08 | 1.70574563 | 0.00166877 |
| ATXN1L | 9.36E-08 | 0.88778757 | 0.00169312 |
| SAMHD1 | 9.71E-08 | 1.16517805 | 0.00175588 |
| TRPV2 | 9.94E-08 | 1.20528365 | 0.00179808 |
| THEMIS2 | 1.00E-07 | 2.11249819 | 0.00181132 |
| IL1RL1 | 1.17E-07 | 0.9422104 | 0.00211691 |
| 8-Mar | 1.18E-07 | 1.62528867 | 0.00212901 |
| IL10RA | 1.20E-07 | 1.76998463 | 0.00216153 |
| RNASE6 | 1.20E-07 | 4.03219543 | 0.00216923 |
| C1QL1 | 1.27E-07 | 1.93807407 | 0.00228811 |
| C9orf16 | 1.40E-07 | 1.52597777 | 0.00253564 |
| PCK1 | 1.44E-07 | 1.21948857 | 0.00259688 |
| SIGLEC1 | 1.46E-07 | 2.46699315 | 0.00263465 |
| MYO9B | 1.46E-07 | 1.18841748 | 0.00264041 |
| CMBL | 1.55E-07 | 1.28596302 | 0.0028061 |
| NPHS2 | 1.59E-07 | -0.722458 | 0.002867 |
| LGALS9 | 1.60E-07 | 1.63458189 | 0.0028861 |
| PPIE | 1.63E-07 | 1.73835367 | 0.00294692 |
| GALNT10 | 1.66E-07 | 1.21260323 | 0.00299661 |
| MSTO1 | 1.75E-07 | 1.16493991 | 0.00315622 |
| STK4 | 1.77E-07 | 1.45172705 | 0.00320245 |
| BCL6 | 1.81E-07 | 1.61745365 | 0.00326899 |
| ATP1A2 | 1.83E-07 | 1.64389298 | 0.00331525 |
| PIK3R1 | 1.87E-07 | 0.99519861 | 0.00337842 |
| IFNGR1 | 1.88E-07 | 0.91888974 | 0.00339771 |
| GGT5 | 1.93E-07 | 1.43999253 | 0.00348822 |
| ARHGEF28 | 1.94E-07 | 1.10345933 | 0.0035004 |
| CYP26B1 | 2.01E-07 | 1.32378075 | 0.00363703 |
| KYNU | 2.11E-07 | 1.89274271 | 0.00381042 |
| METRNL | 2.18E-07 | 0.87611723 | 0.00394329 |
| PIGG | 2.24E-07 | 0.96509845 | 0.00405577 |
| NUP50 | 2.24E-07 | 0.688108 | 0.00405577 |
| LCP2 | 2.29E-07 | 1.04941994 | 0.00414689 |
| PPP1R15B | 2.37E-07 | 1.09355539 | 0.00428493 |
| PLEKHM2 | 2.60E-07 | 0.99676297 | 0.0047055 |
| CRIM1 | 2.61E-07 | -0.8274464 | 0.00471353 |
| ARAP1 | 2.71E-07 | 0.94927402 | 0.00489904 |
| NEK6 | 2.72E-07 | 1.0707234 | 0.00491155 |
| KAT6A | 2.83E-07 | 1.05190311 | 0.00511345 |
| ATG101 | 2.87E-07 | 1.2804816 | 0.00518804 |
| LST1 | 2.91E-07 | 1.98527722 | 0.00526287 |
| NCKAP1L | 2.92E-07 | 1.38160772 | 0.00527598 |
| SKAP2 | 2.92E-07 | 1.56064191 | 0.00528716 |
| RBM47 | 3.07E-07 | 1.00908356 | 0.00555735 |
| FLNA | 3.09E-07 | 1.01730742 | 0.00559291 |
| DNASE1L3 | 3.13E-07 | 1.72669981 | 0.00565327 |
| PLEKHO2 | 3.16E-07 | 0.89613143 | 0.00571319 |
| S100A9 | 3.19E-07 | 1.20771912 | 0.00577078 |

| | | | |
|---|---|---|---|
| USP2 | 3.35E-07 | 2.03927018 | 0.00606609 |
| PIK3CD | 3.38E-07 | 1.78763429 | 0.00611566 |
| FPR3 | 3.41E-07 | 1.94527555 | 0.00616793 |
| LSP1 | 3.64E-07 | 1.03090067 | 0.00657811 |
| IFNAR2 | 3.71E-07 | 0.63822757 | 0.00670793 |
| TNFRSF1B | 3.72E-07 | 1.39473704 | 0.00673067 |
| AOX1 | 3.75E-07 | 1.5378277 | 0.00677924 |
| DHCR24 | 3.87E-07 | 1.52330864 | 0.00698997 |
| ZNF385A | 3.91E-07 | 1.5223953 | 0.00707862 |
| THNSL2 | 4.09E-07 | 1.3359099 | 0.00740302 |
| NPL | 4.10E-07 | 1.12626747 | 0.00742071 |
| SLCO2A1 | 4.13E-07 | 0.82069502 | 0.00747154 |
| B2M | 4.16E-07 | 0.53999526 | 0.00752972 |
| MPP1 | 4.25E-07 | 1.04109923 | 0.00768638 |
| CYTH1 | 4.27E-07 | 0.935156 | 0.00771564 |
| GALT | 4.33E-07 | 1.20893904 | 0.00782394 |
| PPIL3 | 4.37E-07 | 2.07737651 | 0.00790087 |
| PCGF5 | 4.46E-07 | 0.9392916 | 0.00805835 |
| ACTN4 | 4.60E-07 | -0.711261 | 0.0083171 |
| ADGRF5 | 4.64E-07 | 0.72657982 | 0.00838384 |
| NAXE | 4.75E-07 | 0.62540764 | 0.00858561 |
| PIPOX | 4.84E-07 | 1.41348134 | 0.0087472 |
| FAM241A | 5.20E-07 | 1.28338351 | 0.00940129 |
| KLF6 | 5.20E-07 | 0.91161116 | 0.00940747 |
| HAVCR2 | 5.24E-07 | 1.47244264 | 0.00948265 |
| VRK2 | 5.37E-07 | 1.60957936 | 0.00971692 |
| FERMT3 | 5.49E-07 | 0.81948597 | 0.00992502 |
| RPP30 | 5.52E-07 | 1.39463496 | 0.00999073 |
| PTGIS | 5.54E-07 | 1.44021582 | 0.01002024 |
| AFG3L2 | 5.73E-07 | 0.51662389 | 0.01036501 |
| TRAPPC13 | 5.74E-07 | 1.26074273 | 0.01037819 |
| MATN3 | 5.74E-07 | 1.62213199 | 0.01038886 |
| PIGF | 5.93E-07 | 1.25031286 | 0.0107213 |
| RPF2 | 6.02E-07 | 2.33957141 | 0.01089604 |
| FOS | 6.07E-07 | 1.07584883 | 0.01096931 |
| LGALS3 | 6.38E-07 | 0.8787024 | 0.01153028 |
| RAB35 | 6.54E-07 | 0.90293424 | 0.01183009 |
| PSMA4 | 6.73E-07 | 0.55152097 | 0.01216489 |
| POR | 6.96E-07 | 0.92354378 | 0.0125961 |
| KIAA1109 | 7.26E-07 | 0.88075273 | 0.01313338 |
| KCNMB4 | 7.31E-07 | 1.90884058 | 0.01322397 |
| ARMH4 | 7.34E-07 | -1.1024982 | 0.01328026 |
| USO1 | 7.43E-07 | 1.20219014 | 0.01344523 |
| TCN2 | 7.50E-07 | 0.96828931 | 0.01356313 |
| RBM22 | 7.78E-07 | 1.32234105 | 0.01406592 |
| ELMO1 | 7.88E-07 | 1.65661228 | 0.01425416 |
| SLCO2B1 | 8.01E-07 | 0.84477006 | 0.01447961 |
| PDZD2 | 8.06E-07 | 1.6342248 | 0.01457746 |
| GPX3 | 8.25E-07 | 1.13339014 | 0.01491923 |
| CDYL2 | 8.26E-07 | 1.79086177 | 0.0149407 |
| PACSIN3 | 8.42E-07 | 1.63228091 | 0.01522637 |
| DAG1 | 8.42E-07 | -1.0139727 | 0.01523404 |
| AKNA | 8.45E-07 | 1.3349798 | 0.0152898 |

| | | | |
|---|---|---|---|
| SMARCD1 | 8.54E-07 | 1.03461117 | 0.01543878 |
| ADAM12 | 9.10E-07 | 1.41282014 | 0.01645546 |
| ASCC1 | 9.15E-07 | 1.5216316 | 0.01655568 |
| RASSF4 | 9.18E-07 | 1.4126026 | 0.01660909 |
| S100A8 | 9.40E-07 | 1.21791699 | 0.01699895 |
| APBB1IP | 9.45E-07 | 1.51907927 | 0.01708756 |
| UCP2 | 9.68E-07 | 0.65227743 | 0.01750292 |
| COPG2 | 9.69E-07 | 1.29534611 | 0.01751943 |
| CAB39 | 1.00E-06 | 0.66212428 | 0.01808267 |
| CSRNP1 | 1.01E-06 | 1.17897185 | 0.0183368 |
| NCF2 | 1.05E-06 | 0.92574971 | 0.01892529 |
| DHCR7 | 1.06E-06 | 2.69857814 | 0.01918306 |
| TBC1D24 | 1.06E-06 | 1.52884556 | 0.01925673 |
| PILRA | 1.08E-06 | 0.87087077 | 0.01947176 |
| CYBA | 1.08E-06 | 0.75533189 | 0.01948412 |
| PTK2B | 1.10E-06 | 1.15626845 | 0.01982144 |
| AGT | 1.10E-06 | 1.86903554 | 0.01990204 |
| ELL2 | 1.11E-06 | 0.90950437 | 0.02008771 |
| PRDM4 | 1.17E-06 | 1.23783845 | 0.0211853 |
| TAF4 | 1.18E-06 | 1.03032103 | 0.02132684 |
| ECHDC3 | 1.21E-06 | 1.40491269 | 0.02179928 |
| BLVRB | 1.30E-06 | 0.88848951 | 0.02351888 |
| MOCS1 | 1.32E-06 | 1.56350107 | 0.02382259 |
| LHFPL2 | 1.34E-06 | 0.58538161 | 0.02418948 |
| AXL | 1.37E-06 | 0.56893302 | 0.0248335 |
| SNAI2 | 1.38E-06 | 0.8657444 | 0.0249559 |
| CYP4A11 | 1.39E-06 | 1.77991905 | 0.02505536 |
| CHST2 | 1.39E-06 | 1.72314882 | 0.02507214 |
| CHDH | 1.40E-06 | 2.23132876 | 0.02526834 |
| NMRAL1 | 1.40E-06 | 1.28341275 | 0.02534371 |
| ZSWIM6 | 1.41E-06 | 1.14110312 | 0.0254545 |
| ATE1 | 1.43E-06 | 0.66000413 | 0.02593269 |
| TMEM44 | 1.44E-06 | 2.48283498 | 0.02605765 |
| TTC33 | 1.53E-06 | 1.27235619 | 0.02758732 |
| VSIR | 1.54E-06 | 1.0900482 | 0.02781843 |
| AIMP2 | 1.56E-06 | 1.02728246 | 0.02822078 |
| PCSK6 | 1.57E-06 | 1.44431479 | 0.02838103 |
| TCF20 | 1.57E-06 | 1.23083211 | 0.02846267 |
| NAPRT | 1.59E-06 | 0.85635939 | 0.02868577 |
| FMNL1 | 1.60E-06 | 1.47362436 | 0.02901512 |
| SNX5 | 1.62E-06 | 1.23483029 | 0.02933811 |
| CLCNKA | 1.69E-06 | 1.95961563 | 0.03053294 |
| SPP1 | 1.69E-06 | 0.99310962 | 0.03054532 |
| SLC8A1 | 1.70E-06 | 0.57112103 | 0.03070003 |
| ABCF2 | 1.71E-06 | 1.32885398 | 0.0310017 |
| DBI | 1.72E-06 | 0.85731062 | 0.03101865 |
| MOB3A | 1.72E-06 | 1.14307877 | 0.0311358 |
| CD4 | 1.75E-06 | 0.85367373 | 0.03158045 |
| CCN2 | 1.75E-06 | -0.7883265 | 0.03171241 |
| NR1H3 | 1.76E-06 | 0.89444226 | 0.03174289 |
| IL1RN | 1.83E-06 | 4.37945667 | 0.03312631 |
| HTT | 1.84E-06 | 0.52969068 | 0.03324207 |
| MPEG1 | 1.85E-06 | 1.09106631 | 0.03342437 |

| | | | |
|---|---|---|---|
| ZBTB11 | 1.85E-06 | 1.11904059 | 0.0335135 |
| SLC35D2 | 1.86E-06 | 0.50249837 | 0.03361218 |
| DHDH | 1.86E-06 | 1.36440694 | 0.03368072 |
| VWA5A | 1.89E-06 | 1.40886333 | 0.03417532 |
| CSF3R | 1.91E-06 | 1.15156415 | 0.03451443 |
| LBR | 1.91E-06 | 0.54584528 | 0.03453299 |
| PDK2 | 1.91E-06 | 0.88355188 | 0.0345488 |
| CRIPT | 1.91E-06 | 0.65529166 | 0.03460499 |
| ACIN1 | 1.92E-06 | 0.58078914 | 0.03477356 |
| CCDC84 | 1.94E-06 | 1.24425415 | 0.03504789 |
| IFT122 | 1.94E-06 | 0.91789117 | 0.03507254 |
| SLC27A2 | 1.94E-06 | 0.87344983 | 0.03510172 |
| CD40 | 1.95E-06 | 1.04094944 | 0.03525205 |
| SNX33 | 1.96E-06 | 1.12207324 | 0.03536616 |
| CD93 | 1.96E-06 | 0.80516384 | 0.0353918 |
| RGCC | 1.97E-06 | 1.56784248 | 0.03569768 |
| RHOG | 1.98E-06 | 0.8010816 | 0.03577965 |
| HPD | 2.00E-06 | 1.16148226 | 0.03623482 |
| CHST11 | 2.03E-06 | 1.3251469 | 0.03664269 |
| RAP2B | 2.04E-06 | 0.65020686 | 0.03694168 |
| HEATR3 | 2.05E-06 | 1.37481539 | 0.03698981 |
| CDC34 | 2.05E-06 | 0.95226685 | 0.03713185 |
| NBN | 2.06E-06 | 1.07512138 | 0.03725293 |
| KBTBD2 | 2.07E-06 | 0.69710802 | 0.03743615 |
| AOAH | 2.08E-06 | 1.82127557 | 0.03769866 |
| SERPINE1 | 2.15E-06 | 2.53611318 | 0.0388895 |
| FCGBP | 2.20E-06 | 1.76193362 | 0.03978474 |
| ZNF664 | 2.23E-06 | 0.87787267 | 0.04027918 |
| INPP5D | 2.23E-06 | 1.99272156 | 0.04033568 |
| OSMR | 2.23E-06 | 1.02500823 | 0.04035099 |
| WFDC2 | 2.24E-06 | 0.80735837 | 0.04044473 |
| ABI3 | 2.27E-06 | 0.8371064 | 0.04113489 |
| CCND3 | 2.32E-06 | 0.92575116 | 0.04188584 |
| FGR | 2.32E-06 | 1.36916977 | 0.04195959 |
| WAS | 2.33E-06 | 1.34479711 | 0.04220009 |
| LPL | 2.36E-06 | 1.01395177 | 0.04272486 |
| CPM | 2.40E-06 | 0.79029749 | 0.04344777 |
| SC5D | 2.42E-06 | 1.26831701 | 0.04374545 |
| MT1E | 2.43E-06 | 1.22487483 | 0.04390487 |
| EHHADH | 2.44E-06 | 1.2142615 | 0.04413073 |
| RARS | 2.50E-06 | 0.76889132 | 0.04522437 |
| FCN3 | 2.50E-06 | 0.76181488 | 0.045289 |
| ITGB8 | 2.53E-06 | -0.8522425 | 0.0457186 |
| CYBB | 2.53E-06 | 1.24186201 | 0.04576293 |
| PKN3 | 2.54E-06 | 2.222321 | 0.04602446 |
| COL4A3 | 2.58E-06 | -0.8840376 | 0.04668215 |
| POLR2K | 2.60E-06 | 0.86284229 | 0.04709483 |
| SBSPON | 2.63E-06 | 0.51723491 | 0.04749437 |
| PLCB2 | 2.65E-06 | 1.64059254 | 0.04797673 |
| PATL1 | 2.66E-06 | 1.53727096 | 0.04805978 |
| WDR3 | 2.68E-06 | 1.32828192 | 0.04854768 |
| ITPKC | 2.69E-06 | 1.43121746 | 0.04863579 |
| KCNJ15 | 2.69E-06 | 0.74739344 | 0.04871269 |

| | | | |
|---|---|---|---|
| PER1 | 2.70E-06 | 0.9324495 | 0.04883588 |
| PSMB6 | 2.76E-06 | 0.82585358 | 0.04993184 |
| SENP3 | 2.80E-06 | 1.58550321 | 0.05068653 |
| EFHD2 | 2.83E-06 | 0.91235415 | 0.05109611 |
| PRKCD | 2.83E-06 | 0.71282094 | 0.05119987 |
| HSPB2 | 2.93E-06 | 1.99341692 | 0.05292576 |
| AHDC1 | 2.99E-06 | 0.71554566 | 0.05412654 |
| MXD1 | 3.02E-06 | 3.0710252 | 0.05456216 |
| CYFIP2 | 3.02E-06 | 1.16739924 | 0.05466678 |
| ALDH3B1 | 3.04E-06 | 1.41186407 | 0.05504098 |
| LRRFIP2 | 3.12E-06 | 0.91190095 | 0.05651518 |
| DOCK10 | 3.14E-06 | 0.70105501 | 0.0567226 |
| XPNPEP2 | 3.14E-06 | 1.22499173 | 0.05685684 |
| AMOTL2 | 3.15E-06 | 0.85805017 | 0.05698567 |
| EGLN2 | 3.19E-06 | 0.91635652 | 0.0577577 |
| DOCK2 | 3.20E-06 | 0.6913194 | 0.05779567 |
| ACTR2 | 3.24E-06 | 1.02025897 | 0.05853714 |
| PLEKHO1 | 3.24E-06 | 0.69507946 | 0.05868223 |
| TXNDC9 | 3.25E-06 | 2.73620441 | 0.05871711 |
| SLC30A6 | 3.26E-06 | 1.69966332 | 0.05888256 |
| LNPK | 3.30E-06 | 0.82388487 | 0.05963745 |
| RRAGD | 3.38E-06 | 0.98697437 | 0.06108613 |
| ENOPH1 | 3.38E-06 | 0.68574425 | 0.06114717 |
| APOBR | 3.39E-06 | 0.71608669 | 0.06139163 |
| IFNGR2 | 3.55E-06 | 0.88480316 | 0.0642332 |
| VMP1 | 3.57E-06 | 0.86564797 | 0.06454771 |
| MYO1F | 3.58E-06 | 1.76790628 | 0.06467961 |
| EPB41 | 3.59E-06 | 1.14150661 | 0.06488598 |
| IRF8 | 3.67E-06 | 1.01631853 | 0.06637347 |
| KIAA0100 | 3.80E-06 | 0.75907658 | 0.06873269 |
| LZTS1 | 3.87E-06 | 3.73781302 | 0.06992624 |
| SLC4A9 | 3.89E-06 | 1.3651585 | 0.07034793 |
| RCN3 | 3.92E-06 | 0.59017572 | 0.07083224 |
| SOSTDC1 | 3.92E-06 | 1.50590759 | 0.07091881 |
| MTHFS | 3.92E-06 | 1.53516092 | 0.07095304 |
| RNASET2 | 3.95E-06 | 1.40925952 | 0.07145611 |
| ATG4D | 3.96E-06 | 0.55172737 | 0.07153477 |
| SF3A3 | 3.96E-06 | 1.675149 | 0.07160189 |
| MEGF8 | 3.96E-06 | 1.05221916 | 0.07170198 |
| LYVE1 | 3.99E-06 | 2.88695802 | 0.07212798 |
| GGPS1 | 3.99E-06 | 2.80860307 | 0.07212798 |
| PCOLCE | 4.02E-06 | 0.95468531 | 0.07264197 |
| ARHGEF1 | 4.04E-06 | 0.66027068 | 0.07300789 |
| NFKB1 | 4.05E-06 | 1.17012566 | 0.0732539 |
| CHCHD1 | 4.06E-06 | 1.09396028 | 0.07350321 |
| EIF2B2 | 4.08E-06 | 0.73851281 | 0.0737656 |
| H6PD | 4.10E-06 | 1.33360997 | 0.07410261 |
| ACTB | 4.16E-06 | 0.67867273 | 0.07523506 |
| DNAJA3 | 4.18E-06 | 1.05106598 | 0.07557401 |
| NENF | 4.18E-06 | 0.95301767 | 0.07562334 |
| PSMA2 | 4.18E-06 | 0.59736039 | 0.07562959 |
| CIC | 4.21E-06 | 0.82246882 | 0.07614505 |
| KCNMB1 | 4.25E-06 | 2.59858307 | 0.07682736 |

| | | | |
|---|---|---|---|
| ASRGL1 | 4.26E-06 | 1.25091547 | 0.07698761 |
| VAV1 | 4.28E-06 | 2.3630179 | 0.07736568 |
| ZNF526 | 4.28E-06 | 2.08171192 | 0.07749007 |
| PELI1 | 4.29E-06 | 1.34944435 | 0.07750312 |
| IGFBP6 | 4.34E-06 | 1.47400614 | 0.07842324 |
| CES2 | 4.36E-06 | 1.06388034 | 0.07879864 |
| PODXL | 4.44E-06 | -0.6036912 | 0.0803792 |
| WDR70 | 4.51E-06 | 0.57654853 | 0.08160685 |
| SDHAF3 | 4.51E-06 | 1.42812416 | 0.08162795 |
| ATP6V0A2 | 4.52E-06 | 0.55278824 | 0.08175232 |
| ITM2C | 4.52E-06 | -0.8625128 | 0.08179675 |
| GCDH | 4.59E-06 | 2.79443496 | 0.08296673 |
| MRTFA | 4.59E-06 | 1.11590895 | 0.08302944 |
| TKFC | 4.60E-06 | 1.09225609 | 0.08326214 |
| PPIF | 4.78E-06 | 1.28754863 | 0.08641374 |
| STRN4 | 4.80E-06 | 0.7264446 | 0.08687001 |
| TIGD5 | 4.82E-06 | 0.82504986 | 0.08715148 |
| P2RX4 | 4.85E-06 | 1.00381342 | 0.08776912 |
| PRKAA1 | 4.87E-06 | 0.64824894 | 0.08803166 |
| YARS2 | 4.87E-06 | 1.64991052 | 0.08808766 |
| ITPR2 | 4.87E-06 | 0.80963806 | 0.08815875 |
| EIF2B4 | 4.88E-06 | 2.77392305 | 0.08833092 |
| SPAG5 | 4.91E-06 | 0.72894973 | 0.08875466 |
| NDUFAB1 | 4.94E-06 | 0.52216464 | 0.08930171 |
| FOXO3 | 4.98E-06 | 1.09618555 | 0.09002384 |
| FGF1 | 4.98E-06 | -0.6929856 | 0.09010902 |
| EXTL3 | 4.99E-06 | 0.74518814 | 0.09022451 |
| CCDC88C | 5.06E-06 | 1.28270585 | 0.09147291 |
| DCUN1D1 | 5.07E-06 | 1.46578368 | 0.09171489 |
| JDP2 | 5.10E-06 | 0.68511776 | 0.09224156 |
| CLDN7 | 5.26E-06 | 0.57572433 | 0.09516896 |
| GIT1 | 5.30E-06 | 0.77676629 | 0.09579959 |
| GTF3C2 | 5.44E-06 | 1.52739612 | 0.09845511 |
| FBXW11 | 5.48E-06 | 0.88486434 | 0.09909396 |
| AZIN1 | 5.58E-06 | 0.97473963 | 0.10084591 |
| GMIP | 5.59E-06 | 3.89514712 | 0.10108562 |
| PLA2G7 | 5.59E-06 | 3.74136688 | 0.10108562 |
| PRAM1 | 5.59E-06 | 3.67811604 | 0.10108562 |
| PEX1 | 5.61E-06 | 0.8854848 | 0.10152344 |
| SIDT2 | 5.70E-06 | 1.68007102 | 0.10307362 |
| ADCY7 | 5.77E-06 | 1.19628542 | 0.10434882 |
| PLEKHM3 | 5.77E-06 | 1.00582293 | 0.10434882 |
| EMC3 | 5.83E-06 | 0.8172721 | 0.1053708 |
| MPST | 5.86E-06 | 1.16338694 | 0.10600903 |
| AP5Z1 | 5.91E-06 | 0.95652144 | 0.10680943 |
| DNTTIP1 | 5.93E-06 | 1.17850521 | 0.10723956 |
| NAGK | 5.93E-06 | 0.92036095 | 0.10725083 |
| PRLR | 5.99E-06 | 1.47819415 | 0.10840149 |
| PRAP1 | 6.03E-06 | 1.21495778 | 0.10912804 |
| POLR3A | 6.05E-06 | 0.65971316 | 0.10933273 |
| CD276 | 6.06E-06 | 1.02755184 | 0.10963232 |
| FOLR2 | 6.10E-06 | 1.216264 | 0.11033353 |
| COL5A2 | 6.11E-06 | 1.1107331 | 0.11043137 |

| | | | |
|---|---|---|---|
| NT5C | 6.11E-06 | 0.63125248 | 0.11043137 |
| CCM2 | 6.11E-06 | 0.68995044 | 0.11054226 |
| HCST | 6.16E-06 | 1.64097803 | 0.11140974 |
| PLXNA2 | 6.18E-06 | 1.79274878 | 0.1117383 |
| BRWD1 | 6.33E-06 | 0.78568642 | 0.11439046 |
| TIMM17A | 6.33E-06 | 1.07989062 | 0.1145564 |
| SLC41A1 | 6.42E-06 | 0.6264717 | 0.11614674 |
| ZZEF1 | 6.49E-06 | 0.99713609 | 0.11729198 |
| MAVS | 6.51E-06 | 0.88248185 | 0.1178008 |
| CHRDL1 | 6.63E-06 | 0.87636589 | 0.11982386 |
| NES | 6.63E-06 | -0.5613505 | 0.11995021 |
| RDH11 | 6.64E-06 | 0.77322672 | 0.12000134 |
| NCS1 | 6.67E-06 | 1.10655106 | 0.12056144 |
| CSF1 | 6.72E-06 | 0.78593935 | 0.12158863 |
| ADH1B | 6.77E-06 | 1.28556732 | 0.12237053 |
| KCNC3 | 6.82E-06 | 1.05922866 | 0.12331658 |
| FAM49A | 6.83E-06 | 0.98690288 | 0.1234333 |
| KLK6 | 6.88E-06 | -1.6255628 | 0.12434581 |
| FNTA | 6.96E-06 | 1.00629001 | 0.12595903 |
| CD52 | 6.97E-06 | 2.07198272 | 0.1260963 |
| VDAC2 | 6.97E-06 | 0.62513989 | 0.12612579 |
| HNMT | 7.00E-06 | 1.09528159 | 0.1266095 |
| CHST15 | 7.03E-06 | 1.80737154 | 0.12711085 |
| NRF1 | 7.09E-06 | 1.62078975 | 0.12830914 |
| TLR2 | 7.16E-06 | 0.80021903 | 0.12947424 |
| TLE1 | 7.31E-06 | 0.83400464 | 0.13224266 |
| USP11 | 7.35E-06 | 0.54532546 | 0.13283695 |
| ELL | 7.41E-06 | 1.02593728 | 0.13407211 |
| MED6 | 7.42E-06 | 0.83886348 | 0.13418076 |
| STAB1 | 7.48E-06 | 1.10098886 | 0.13521272 |
| IKBKB | 7.60E-06 | 0.98607873 | 0.13736342 |
| PEX16 | 7.71E-06 | 0.70251935 | 0.13939509 |
| PUM3 | 7.75E-06 | -0.6971053 | 0.14022595 |
| ULK2 | 7.80E-06 | 0.98079943 | 0.1410531 |
| PPP3CA | 7.83E-06 | 0.89889018 | 0.14168721 |
| KLHDC1 | 7.86E-06 | 1.57707047 | 0.14220315 |
| TM4SF18 | 7.87E-06 | 0.62209517 | 0.14225844 |
| MICAL2 | 7.87E-06 | 0.91874814 | 0.14228344 |
| TAB3 | 7.92E-06 | 0.92994147 | 0.14324193 |
| PLXNC1 | 7.96E-06 | 1.64576046 | 0.14401588 |
| MED9 | 7.97E-06 | 1.37963157 | 0.14404921 |
| PKLR | 8.03E-06 | 1.07997151 | 0.14528636 |
| SETD4 | 8.09E-06 | 0.73937382 | 0.14629693 |
| ASS1 | 8.09E-06 | 1.01874666 | 0.14636044 |
| FABP1 | 8.10E-06 | 1.35302724 | 0.14651679 |
| CHCHD3 | 8.12E-06 | 1.77067877 | 0.14682066 |
| APOM | 8.15E-06 | 1.44289951 | 0.14740301 |
| SENP1 | 8.27E-06 | 0.71537426 | 0.14963952 |
| TRIM3 | 8.33E-06 | 1.07016574 | 0.15056254 |
| ZBTB7B | 8.38E-06 | 0.82305733 | 0.15146836 |
| SLC13A3 | 8.44E-06 | 1.30630934 | 0.15268146 |
| POGK | 8.55E-06 | 0.75876598 | 0.15465842 |
| PARD6B | 8.63E-06 | 0.83894133 | 0.15607272 |

| | | | |
|---|---|---|---|
| TRANK1 | 8.63E-06 | 0.93372248 | 0.15608015 |
| FBH1 | 8.67E-06 | 0.69430543 | 0.15687256 |
| MGST1 | 8.75E-06 | 1.34700283 | 0.1582857 |
| MXD4 | 8.79E-06 | 0.74711954 | 0.15896313 |
| CFH | 8.79E-06 | 0.9745078 | 0.15897179 |
| ATF7 | 8.82E-06 | 0.80058506 | 0.15942272 |
| RGL1 | 8.90E-06 | 0.85605696 | 0.16094817 |
| PIN1 | 8.99E-06 | 0.94079 | 0.1624975 |
| ITGAL | 9.03E-06 | 1.65603491 | 0.1633294 |
| KIAA1549 | 9.17E-06 | 1.1197463 | 0.16588534 |
| LSM2 | 9.21E-06 | 0.74268661 | 0.16663748 |
| TMED8 | 9.37E-06 | 1.45322873 | 0.16945685 |
| TBX3 | 9.39E-06 | 0.72228643 | 0.16974384 |
| EAF2 | 9.45E-06 | 1.28337541 | 0.17081751 |
| BID | 9.45E-06 | 0.89006801 | 0.17082772 |
| SH3BP1 | 9.45E-06 | 1.77060617 | 0.17089921 |
| EIF5 | 9.51E-06 | 0.75159544 | 0.17202681 |
| PES1 | 9.52E-06 | 0.93130336 | 0.17212848 |
| FAM49B | 9.52E-06 | 1.257541 | 0.1721409 |
| PGLS | 9.52E-06 | 0.86084993 | 0.17216261 |
| F8 | 9.53E-06 | 0.88242843 | 0.1723061 |
| SHISA5 | 9.53E-06 | 0.66508263 | 0.17237722 |
| CIZ1 | 9.56E-06 | 0.73820558 | 0.17284942 |
| SNX10 | 9.71E-06 | 1.31399341 | 0.17563535 |
| ACAA1 | 9.88E-06 | 0.8228449 | 0.17872579 |
| EED | 1.00E-05 | 2.09394656 | 0.18103999 |
| SLC20A1 | 1.01E-05 | 1.03068877 | 0.18196713 |
| FABP3 | 1.01E-05 | 1.08998198 | 0.18254574 |
| ARRB1 | 1.04E-05 | 0.61327969 | 0.18864459 |
| GHR | 1.05E-05 | 0.83753275 | 0.19044239 |
| HSPBP1 | 1.07E-05 | 0.80713624 | 0.19415021 |
| PACRG | 1.08E-05 | 2.71291546 | 0.19445341 |
| TRHDE | 1.08E-05 | 0.9434125 | 0.19571676 |
| ACOX1 | 1.09E-05 | 0.74839713 | 0.19661004 |
| NFIB | 1.10E-05 | 0.77551054 | 0.19808863 |
| P4HA2 | 1.10E-05 | 1.29001544 | 0.19834154 |
| RAB7B | 1.10E-05 | 0.56374578 | 0.1992511 |
| NABP1 | 1.11E-05 | 0.94734557 | 0.19990434 |
| UBC | 1.11E-05 | 0.92517464 | 0.20004656 |
| SBF2 | 1.13E-05 | 0.84432373 | 0.20517332 |
| ADI1 | 1.14E-05 | 0.73291068 | 0.2053593 |
| ACBD6 | 1.14E-05 | 0.84222939 | 0.20600191 |
| SUSD2 | 1.14E-05 | 0.7428766 | 0.2065516 |
| DESI1 | 1.15E-05 | 0.64039055 | 0.20857164 |
| MME | 1.16E-05 | -0.6212305 | 0.2092086 |
| BAD | 1.17E-05 | 0.86366785 | 0.2110154 |
| ZDHHC5 | 1.17E-05 | 0.78468982 | 0.21208889 |
| SBF1 | 1.20E-05 | 1.07273971 | 0.21645557 |
| TBK1 | 1.20E-05 | 0.62820702 | 0.21763077 |
| AQP1 | 1.22E-05 | 0.62921253 | 0.22006363 |
| KRIT1 | 1.22E-05 | 0.95271497 | 0.22059411 |
| PTPRE | 1.22E-05 | 0.94319809 | 0.22113081 |
| GPATCH3 | 1.23E-05 | 0.70294404 | 0.22210591 |

| | | | |
|---|---|---|---|
| MLXIPL | 1.23E-05 | 0.99389644 | 0.22220151 |
| RPP25L | 1.23E-05 | 1.27005267 | 0.22283062 |
| SEMA4D | 1.23E-05 | 0.79656832 | 0.2231302 |
| STYX | 1.24E-05 | 1.24122062 | 0.22382933 |
| SLC43A3 | 1.27E-05 | 2.15299815 | 0.22892814 |
| CAPN15 | 1.27E-05 | 1.23266896 | 0.22901943 |
| TMEM165 | 1.27E-05 | 0.79297099 | 0.22920368 |
| TUBG2 | 1.29E-05 | 0.91071851 | 0.23259676 |
| DDC | 1.29E-05 | 3.21736755 | 0.23417999 |
| CCDC115 | 1.30E-05 | 0.78026092 | 0.23470403 |
| F13A1 | 1.31E-05 | 1.27491304 | 0.23622412 |
| TNFAIP8 | 1.32E-05 | 1.95082163 | 0.23831229 |
| SLC25A10 | 1.32E-05 | 1.00862757 | 0.23925357 |
| ACOT13 | 1.34E-05 | 0.832595 | 0.24242204 |
| GPM6B | 1.34E-05 | 3.14984748 | 0.24272856 |
| VARS | 1.34E-05 | 0.85388172 | 0.24279494 |
| BICD2 | 1.35E-05 | 0.88632297 | 0.24363327 |
| RASL12 | 1.35E-05 | 1.77684468 | 0.24387153 |
| ARHGAP29 | 1.35E-05 | -0.8780641 | 0.24410847 |
| CEP170B | 1.35E-05 | 1.10143166 | 0.24455737 |
| CCDC186 | 1.36E-05 | 0.74850673 | 0.2451203 |
| FMN1 | 1.36E-05 | 1.084916 | 0.24570961 |
| MTCH2 | 1.37E-05 | 1.25612501 | 0.24690232 |
| SIGLEC14 | 1.37E-05 | 3.11768795 | 0.24711369 |
| IFITM3 | 1.37E-05 | 0.58628101 | 0.24806673 |
| MRFAP1 | 1.39E-05 | 1.1668264 | 0.25050222 |
| DGAT1 | 1.40E-05 | 0.83775674 | 0.25259424 |
| PRKAR1B | 1.40E-05 | 1.03710172 | 0.25302601 |
| NADK2 | 1.40E-05 | 0.98289071 | 0.25302601 |
| GPR108 | 1.44E-05 | 1.09561477 | 0.26125634 |
| ACE | 1.45E-05 | 1.13560658 | 0.26199452 |
| SGF29 | 1.45E-05 | 1.11927761 | 0.26286281 |
| ACO1 | 1.47E-05 | 0.79385506 | 0.26633574 |
| BTBD6 | 1.49E-05 | 0.60021115 | 0.26957199 |
| ABCA1 | 1.50E-05 | 0.66354466 | 0.27136121 |
| FAM217B | 1.51E-05 | 1.51281112 | 0.27229812 |
| NCF4 | 1.51E-05 | 2.34762601 | 0.27380526 |
| ADAP2 | 1.52E-05 | 0.75946416 | 0.27443631 |
| RASL11A | 1.53E-05 | 0.7993808 | 0.27682569 |
| MT1X | 1.53E-05 | 0.57096434 | 0.27697936 |
| DNAJB2 | 1.53E-05 | 0.76385133 | 0.27725118 |
| ANGEL2 | 1.54E-05 | 0.69960013 | 0.27761451 |
| ATP11B | 1.54E-05 | 0.91090211 | 0.27762347 |
| MS4A4A | 1.55E-05 | 0.90163436 | 0.27952344 |
| CCDC88B | 1.55E-05 | 2.26447841 | 0.27999393 |
| SNRNP27 | 1.55E-05 | 1.45656408 | 0.28086774 |
| DPY19L3 | 1.56E-05 | 0.58198203 | 0.2828355 |
| C8orf88 | 1.58E-05 | 2.09469406 | 0.28501861 |
| UBXN11 | 1.62E-05 | 0.78907424 | 0.29219953 |
| SLC39A8 | 1.63E-05 | 1.25990784 | 0.2955452 |
| NDUFB6 | 1.64E-05 | 1.17746274 | 0.29603685 |
| ARHGAP30 | 1.64E-05 | 1.3093997 | 0.2970799 |
| ADCY5 | 1.68E-05 | 1.8611816 | 0.30309237 |

| | | | |
|---|---|---|---|
| USP37 | 1.68E-05 | 1.35080051 | 0.30373374 |
| ANAPC15 | 1.69E-05 | 1.0966586 | 0.30510137 |
| PLIN3 | 1.69E-05 | 0.78182685 | 0.30558296 |
| PTPN18 | 1.70E-05 | 1.49695448 | 0.30825501 |
| ELK4 | 1.72E-05 | 0.83331585 | 0.31185027 |
| DUSP15 | 1.76E-05 | 1.16417021 | 0.31747343 |
| LMO7 | 1.78E-05 | 0.79507227 | 0.32260833 |
| IL6R | 1.79E-05 | 1.1902963 | 0.32368215 |
| MDH2 | 1.80E-05 | 0.62551979 | 0.32496556 |
| AAMDC | 1.82E-05 | 0.80930239 | 0.32844514 |
| TFCP2L1 | 1.82E-05 | 0.9154603 | 0.32848431 |
| FBLIM1 | 1.82E-05 | 1.0308767 | 0.32862459 |
| REL | 1.83E-05 | 1.1007352 | 0.33180407 |
| PPP6R1 | 1.84E-05 | 0.81841448 | 0.33289423 |
| SEMA5A | 1.84E-05 | -0.860843 | 0.33359079 |
| RASAL2 | 1.86E-05 | 1.0643588 | 0.33594751 |
| SLC37A4 | 1.86E-05 | 0.90471394 | 0.33691873 |
| LIG3 | 1.87E-05 | 1.40894412 | 0.33784792 |
| MT1G | 1.87E-05 | 0.74080745 | 0.33880828 |
| N6AMT1 | 1.89E-05 | 0.52069031 | 0.34189653 |
| PHYH | 1.89E-05 | 1.03536158 | 0.34240672 |
| C19orf25 | 1.91E-05 | 0.87196553 | 0.34514879 |
| VPS8 | 1.92E-05 | 1.048045 | 0.34749293 |
| TTYH3 | 1.93E-05 | 0.82746689 | 0.34865355 |
| POU3F3 | 1.93E-05 | 0.80954126 | 0.3498467 |
| TTC27 | 1.99E-05 | 0.79383112 | 0.36053891 |
| EVA1C | 2.00E-05 | 1.67082243 | 0.36243835 |
| CPAMD8 | 2.00E-05 | 1.18451053 | 0.36243835 |
| SOCS4 | 2.01E-05 | 0.71669145 | 0.36291779 |
| VASN | 2.01E-05 | -0.9775434 | 0.36383391 |
| SLC25A13 | 2.03E-05 | 0.7747601 | 0.36767257 |
| PSMA3 | 2.05E-05 | 0.73260564 | 0.37071823 |
| PCCA | 2.05E-05 | 0.67723235 | 0.3710408 |
| HOGA1 | 2.05E-05 | 1.00152176 | 0.37114666 |
| SELENOO | 2.06E-05 | 0.70025107 | 0.37258417 |
| UBA1 | 2.07E-05 | 0.58561182 | 0.37446737 |
| NAPSA | 2.07E-05 | 1.36673462 | 0.37450341 |
| MITF | 2.08E-05 | 0.67748789 | 0.37604987 |
| KMT2B | 2.09E-05 | 0.5313729 | 0.37772904 |
| CA11 | 2.09E-05 | 1.01775177 | 0.37869937 |
| ATP11C | 2.11E-05 | 0.85751649 | 0.38195312 |
| GPR34 | 2.11E-05 | 2.60068989 | 0.38227091 |
| DIAPH2 | 2.12E-05 | 1.15315101 | 0.3829391 |
| CBR1 | 2.12E-05 | 0.74455176 | 0.38427025 |
| PTPRJ | 2.13E-05 | 0.86246233 | 0.38462339 |
| ING3 | 2.15E-05 | 1.43223637 | 0.38832744 |
| EDC4 | 2.15E-05 | 0.71051409 | 0.38960526 |
| ZNF740 | 2.17E-05 | 0.9101906 | 0.39310842 |
| PCYT2 | 2.18E-05 | 1.30192651 | 0.39483644 |
| CAMSAP1 | 2.20E-05 | 0.87508351 | 0.39785357 |
| CRIP1 | 2.21E-05 | 1.03093782 | 0.40000198 |
| SORD | 2.22E-05 | 1.54133717 | 0.40098966 |
| PLG | 2.23E-05 | 1.00896059 | 0.40343222 |

| | | | |
|---|---|---|---|
| MYO10 | 2.23E-05 | 1.78885481 | 0.40405924 |
| ARPC3 | 2.24E-05 | 0.94410299 | 0.40505044 |
| PIM1 | 2.26E-05 | 0.84514138 | 0.40899829 |
| DDIT4 | 2.27E-05 | 0.97955104 | 0.40979745 |
| IFI16 | 2.27E-05 | 0.60313235 | 0.41124323 |
| MERTK | 2.27E-05 | 0.82141152 | 0.41131431 |
| SMC5 | 2.30E-05 | 0.91622831 | 0.4161634 |
| MUC20 | 2.31E-05 | 0.84891246 | 0.41728728 |
| LRATD2 | 2.31E-05 | 1.02080664 | 0.41833241 |
| TPM1 | 2.32E-05 | 0.62838115 | 0.41947549 |
| SOST | 2.34E-05 | -1.1463376 | 0.42251172 |
| RND3 | 2.39E-05 | 1.89068749 | 0.43171444 |
| DENND3 | 2.42E-05 | 0.52451436 | 0.43790798 |
| FUZ | 2.42E-05 | 0.58433703 | 0.43797177 |
| UBAC1 | 2.43E-05 | 0.63205335 | 0.43986193 |
| POLR2B | 2.44E-05 | 0.74122463 | 0.4404779 |
| NUDT22 | 2.44E-05 | 0.6947669 | 0.44205042 |
| ARG2 | 2.46E-05 | 1.14541113 | 0.44453188 |
| ARHGAP9 | 2.46E-05 | 2.03321536 | 0.44467297 |
| DNPH1 | 2.46E-05 | 0.95981609 | 0.44540676 |
| CCNC | 2.47E-05 | 1.16356662 | 0.44605266 |
| STEAP3 | 2.48E-05 | 0.89695094 | 0.44782492 |
| ATL3 | 2.49E-05 | 0.53466646 | 0.45035108 |
| METTL7B | 2.50E-05 | 0.74886131 | 0.45229689 |
| AJUBA | 2.51E-05 | 0.791521 | 0.45413358 |
| MEN1 | 2.51E-05 | 1.98179483 | 0.45468122 |
| VPS72 | 2.54E-05 | 0.67456589 | 0.45860356 |
| TTC9C | 2.54E-05 | 1.09234152 | 0.45895875 |
| STAM2 | 2.56E-05 | 0.62383425 | 0.46208548 |
| FAM133B | 2.61E-05 | 1.25483327 | 0.47188635 |
| MMP19 | 2.65E-05 | 0.96646923 | 0.47910734 |
| SNX11 | 2.65E-05 | 0.93533274 | 0.47921816 |
| CCDC71L | 2.67E-05 | 1.57645625 | 0.48337991 |
| OSGEP | 2.68E-05 | 0.90023866 | 0.48553292 |
| VDR | 2.69E-05 | 0.83088837 | 0.48609566 |
| CNOT7 | 2.69E-05 | 1.61356268 | 0.48665821 |
| GGACT | 2.69E-05 | 1.66829517 | 0.48666501 |
| TFEC | 2.69E-05 | 0.86886275 | 0.4873478 |
| GNE | 2.70E-05 | 1.31550504 | 0.48916817 |
| CBFB | 2.72E-05 | 0.69934218 | 0.49218506 |
| LTA4H | 2.72E-05 | 0.65655891 | 0.49279681 |
| PIAS1 | 2.74E-05 | 1.13904824 | 0.49632356 |
| PPIB | 2.75E-05 | 0.52605383 | 0.49678516 |
| NAXD | 2.75E-05 | 0.67905006 | 0.4973069 |
| JMJD1C | 2.78E-05 | 0.7177566 | 0.50211939 |
| DAGLB | 2.80E-05 | 2.36453783 | 0.50595222 |
| PLEKHH2 | 2.81E-05 | 0.68365264 | 0.50755783 |
| LRRC27 | 2.81E-05 | 0.76407606 | 0.50817212 |
| DHX35 | 2.81E-05 | 0.99778365 | 0.50900095 |
| SPRYD7 | 2.82E-05 | 1.02856126 | 0.50964304 |
| ARHGAP6 | 2.83E-05 | 2.11436631 | 0.51101292 |
| C18orf25 | 2.83E-05 | 0.65276829 | 0.51190915 |
| ZBTB5 | 2.84E-05 | 0.72204604 | 0.51336348 |

| | | | |
|---|---|---|---|
| OCLN | 2.85E-05 | 0.52475772 | 0.51616825 |
| PTGER2 | 2.86E-05 | 0.97965158 | 0.51641972 |
| ANAPC4 | 2.87E-05 | 0.61269733 | 0.51815332 |
| SNAP47 | 2.90E-05 | 0.98707821 | 0.52427129 |
| SF3A1 | 2.91E-05 | 0.62872153 | 0.52660103 |
| TREM2 | 2.94E-05 | 1.19671262 | 0.53174263 |
| GSTO1 | 2.94E-05 | 0.62481859 | 0.53221151 |
| AK6 | 2.95E-05 | 1.19413273 | 0.53365754 |
| ARPIN | 2.96E-05 | 0.84263343 | 0.53513167 |
| PDE1C | 2.98E-05 | 2.70023678 | 0.53834526 |
| DOCK9 | 2.98E-05 | 0.52032583 | 0.53880178 |
| RPF1 | 2.98E-05 | 0.6608703 | 0.53928796 |
| ZNF185 | 3.02E-05 | 0.54560069 | 0.5458838 |
| E4F1 | 3.02E-05 | 1.08773166 | 0.5469514 |
| EFL1 | 3.03E-05 | 1.33173152 | 0.54838543 |
| DCBLD2 | 3.04E-05 | 0.55478693 | 0.54900363 |
| RILP | 3.04E-05 | 1.03128076 | 0.55019272 |
| ASNSD1 | 3.05E-05 | 0.7897165 | 0.5522395 |
| SLA | 3.06E-05 | 0.74117836 | 0.55351791 |
| WASHC4 | 3.07E-05 | 0.90766843 | 0.55606771 |
| SAP30 | 3.11E-05 | 1.29083979 | 0.56324713 |
| SULF1 | 3.13E-05 | -1.6986665 | 0.56654721 |
| EDC3 | 3.14E-05 | 1.4890945 | 0.5670772 |
| FAM78A | 3.15E-05 | 2.81096461 | 0.56885655 |
| LRP1 | 3.15E-05 | 0.9250065 | 0.56913947 |
| CCDC191 | 3.19E-05 | 0.91716918 | 0.57608844 |
| DOC2B | 3.22E-05 | 1.65047137 | 0.58205684 |
| MT1F | 3.23E-05 | 0.83849352 | 0.58375917 |
| SMAD1 | 3.25E-05 | 0.9743077 | 0.5870138 |
| WDR86 | 3.26E-05 | 1.01008818 | 0.58989495 |
| MITD1 | 3.28E-05 | 0.81726194 | 0.59330462 |
| SLC25A27 | 3.28E-05 | 1.26300533 | 0.5939425 |
| CCDC8 | 3.31E-05 | 1.36910325 | 0.59932829 |
| CCDC127 | 3.34E-05 | 1.73812761 | 0.60377939 |
| PARVG | 3.35E-05 | 0.72700255 | 0.60510856 |
| KIAA0930 | 3.35E-05 | 0.75913525 | 0.60592882 |
| MASTL | 3.36E-05 | 0.76345945 | 0.60752284 |
| LRRC6 | 3.36E-05 | 0.74801713 | 0.60752284 |
| MROH7 | 3.36E-05 | 1.71144117 | 0.60806778 |
| GPR65 | 3.36E-05 | 3.40192728 | 0.60820394 |
| SLC39A6 | 3.37E-05 | 0.53134326 | 0.60935515 |
| GTF2F1 | 3.39E-05 | 0.53396699 | 0.61372262 |
| VTA1 | 3.39E-05 | 0.77598705 | 0.61385206 |
| LEO1 | 3.40E-05 | 0.57699503 | 0.61564067 |
| ZDHHC7 | 3.41E-05 | 0.65878392 | 0.61710307 |
| ARPC1B | 3.42E-05 | 0.77252854 | 0.61837157 |
| PNP | 3.43E-05 | 0.82601559 | 0.62085178 |
| PPM1G | 3.44E-05 | 0.78748153 | 0.62235875 |
| KLF9 | 3.45E-05 | 1.06126345 | 0.62418162 |
| FHL5 | 3.47E-05 | 1.48329096 | 0.62772907 |
| CLIP2 | 3.48E-05 | 0.59284705 | 0.62886324 |
| SMIM32 | 3.49E-05 | 0.6444501 | 0.6311773 |
| ZBTB25 | 3.54E-05 | 0.73601459 | 0.64093581 |

| | | | |
|---|---|---|---|
| SEMA3C | 3.55E-05 | 0.97598108 | 0.64153567 |
| ALDH4A1 | 3.55E-05 | 0.86907658 | 0.64173675 |
| OGFOD1 | 3.55E-05 | 0.79869255 | 0.64175886 |
| GLMP | 3.58E-05 | 0.92328739 | 0.64790305 |
| GAR1 | 3.59E-05 | 1.45464547 | 0.64931492 |
| RIMKLB | 3.60E-05 | 0.84869352 | 0.6508656 |
| IRGQ | 3.62E-05 | 0.52120592 | 0.65538767 |
| DMRTA1 | 3.65E-05 | 0.92580288 | 0.65969942 |
| CPSF4 | 3.66E-05 | 0.59218507 | 0.66137657 |
| MED18 | 3.68E-05 | 1.15980885 | 0.66510264 |
| XPO4 | 3.68E-05 | 0.65215367 | 0.66611238 |
| TUT7 | 3.68E-05 | 0.58855961 | 0.666359 |
| PPM1M | 3.69E-05 | 1.20410338 | 0.66707633 |
| TNFAIP8L1 | 3.69E-05 | 0.65376389 | 0.6680392 |
| PPA2 | 3.70E-05 | 0.91143996 | 0.669087 |
| BRD3 | 3.75E-05 | 0.51958198 | 0.67783219 |
| NUP85 | 3.77E-05 | 0.64217251 | 0.68150373 |
| VASP | 3.79E-05 | 0.8887834 | 0.68480532 |
| SLC7A9 | 3.80E-05 | 1.21323362 | 0.68756927 |
| HCK | 3.84E-05 | 1.44573601 | 0.69432244 |
| DCAF5 | 3.85E-05 | 0.96388509 | 0.69629452 |
| NFKBIZ | 3.88E-05 | 0.65464434 | 0.70123563 |
| CMKLR1 | 3.88E-05 | 1.13160105 | 0.70200839 |
| ZNF649 | 3.89E-05 | 1.5912157 | 0.70283565 |
| CD86 | 3.90E-05 | 0.89861295 | 0.70588709 |
| TYW3 | 3.95E-05 | 1.04946066 | 0.71463308 |
| DHDDS | 3.96E-05 | 0.77097496 | 0.71574048 |
| KTI12 | 3.97E-05 | 1.15287095 | 0.71763442 |
| MOB4 | 4.01E-05 | 0.5451234 | 0.72539801 |
| LRFN3 | 4.06E-05 | 0.72312948 | 0.73452354 |
| ARRDC2 | 4.07E-05 | 0.68284731 | 0.73608715 |
| LPIN1 | 4.07E-05 | 0.92876791 | 0.73655351 |
| BRCC3 | 4.08E-05 | 0.84950216 | 0.73742163 |
| DOCK1 | 4.09E-05 | 0.66232724 | 0.73922013 |
| SH3TC1 | 4.12E-05 | 0.52862278 | 0.74431001 |
| MPHOSPH10 | 4.16E-05 | 0.71746934 | 0.75260754 |
| COMMD7 | 4.16E-05 | 0.90576457 | 0.7527623 |
| SLC30A2 | 4.18E-05 | 0.84316549 | 0.75591886 |
| PAFAH2 | 4.24E-05 | 0.69863335 | 0.7674679 |
| BIN1 | 4.26E-05 | 0.56786994 | 0.77061537 |
| SNX6 | 4.27E-05 | 0.69761091 | 0.77159711 |
| EPS8L1 | 4.29E-05 | 1.40707765 | 0.77625731 |
| CKAP5 | 4.32E-05 | 0.58291097 | 0.78118931 |
| MSH6 | 4.35E-05 | 1.03689236 | 0.78741536 |
| ZBTB39 | 4.36E-05 | 1.82259832 | 0.7881859 |
| SLC9A8 | 4.36E-05 | 1.56165531 | 0.7881859 |
| MEIS2 | 4.36E-05 | 0.6575343 | 0.78913411 |
| CDKN2B | 4.37E-05 | 1.6098681 | 0.79024497 |
| OXNAD1 | 4.39E-05 | 0.80786134 | 0.79444294 |
| ETNK2 | 4.40E-05 | 0.83270047 | 0.79549989 |
| POU5F2 | 4.44E-05 | 1.99557563 | 0.80215731 |
| ELOF1 | 4.44E-05 | 0.67095562 | 0.80247842 |
| DHX32 | 4.49E-05 | 1.52246096 | 0.81256668 |

| | | | |
|---|---|---|---|
| CDH1 | 4.51E-05 | 0.93271688 | 0.81537816 |
| PSMC4 | 4.54E-05 | 0.73570721 | 0.8215503 |
| AKAP12 | 4.56E-05 | 0.79698445 | 0.82460847 |
| HM13 | 4.57E-05 | 0.57665198 | 0.82622581 |
| PTPN3 | 4.59E-05 | 0.79043252 | 0.83100166 |
| SELENBP1 | 4.60E-05 | 1.01818331 | 0.83201955 |
| BCL2L1 | 4.61E-05 | 0.6641554 | 0.83385605 |
| CD58 | 4.62E-05 | 0.60104368 | 0.83506759 |
| KIAA1328 | 4.65E-05 | 0.76532679 | 0.84173897 |
| RAB40C | 4.66E-05 | 0.64068233 | 0.84248779 |
| PGS1 | 4.69E-05 | 0.74086577 | 0.84831121 |
| NKRF | 4.69E-05 | 0.62149772 | 0.84887056 |
| ECM1 | 4.71E-05 | -1.7305812 | 0.85203338 |
| CLDND1 | 4.71E-05 | 0.94557202 | 0.85267245 |
| AKAP10 | 4.72E-05 | 1.00226397 | 0.85286377 |
| ADAMTS15 | 4.74E-05 | 0.95618718 | 0.85700262 |
| FKTN | 4.80E-05 | 1.05351465 | 0.86769639 |
| MAOA | 4.81E-05 | 0.8460281 | 0.86939665 |
| TCIRG1 | 4.83E-05 | 0.64999068 | 0.8735932 |
| CDC40 | 4.85E-05 | 1.71221491 | 0.87668603 |
| RTCB | 4.85E-05 | 0.5133577 | 0.87675641 |
| CALHM2 | 4.91E-05 | 0.66612326 | 0.88711503 |
| GCHFR | 4.93E-05 | 0.63511533 | 0.89088092 |
| DCXR | 4.97E-05 | 1.03593075 | 0.89813862 |
| COTL1 | 4.97E-05 | 0.64636004 | 0.89813862 |
| YAE1 | 4.97E-05 | 0.75205042 | 0.8985632 |
| BET1L | 4.98E-05 | 0.59805257 | 0.90022184 |
| POLD3 | 5.01E-05 | 1.02455083 | 0.90688994 |
| BMP6 | 5.03E-05 | 1.92228106 | 0.90938581 |
| CEP89 | 5.06E-05 | 1.3256999 | 0.91423608 |
| ZYX | 5.08E-05 | 0.90089667 | 0.9192473 |
| NLN | 5.13E-05 | 0.65675276 | 0.92797539 |
| IFIH1 | 5.17E-05 | 0.85118663 | 0.93461432 |
| ILRUN | 5.19E-05 | 0.59907923 | 0.93805906 |
| PPP5C | 5.20E-05 | 0.8092998 | 0.93963669 |
| ASAP1 | 5.23E-05 | 0.66360775 | 0.94595663 |
| KMO | 5.25E-05 | 1.20505144 | 0.95008408 |
| PIK3CA | 5.25E-05 | 0.86590977 | 0.95021828 |
| UBE2J2 | 5.26E-05 | 0.76021121 | 0.9521362 |
| TRIR | 5.28E-05 | 0.69367522 | 0.95415705 |
| MTRF1 | 5.31E-05 | 0.82594041 | 0.95996413 |
| IKBKG | 5.31E-05 | 0.58700673 | 0.95999158 |
| PDCL | 5.33E-05 | 0.75707093 | 0.96353201 |
| CD82 | 5.36E-05 | 1.04342066 | 0.96849692 |
| KCNJ16 | 5.42E-05 | 0.7489687 | 0.98025216 |
| TMEM37 | 5.42E-05 | 0.8370527 | 0.98043978 |
| ZNF592 | 5.45E-05 | 0.8736934 | 0.98523269 |
| SNX9 | 5.47E-05 | 0.67200903 | 0.98895476 |
| ZNF207 | 5.47E-05 | 1.39516154 | 0.98993471 |
| ERAL1 | 5.48E-05 | 0.82123962 | 0.99087386 |
| STRAP | 5.49E-05 | 0.53289466 | 0.99218144 |
| CRAMP1 | 5.50E-05 | 0.64752743 | 0.9938792 |
| DSE | 5.52E-05 | 0.6372088 | 0.99775716 |

| | | | |
|---|---|---|---|
| NHEJ1 | 5.52E-05 | 0.84002735 | 0.99848463 |
| BMERB1 | 5.53E-05 | 0.59915328 | 0.99988896 |
| OR10D3 | 0.00354541 | 3.86448341 | 1 |
| PRG2 | 0.01248949 | 3.73165463 | 1 |
| GALNT15 | 0.00036661 | 3.69017663 | 1 |
| CLEC12A | 0.00051031 | 3.53881202 | 1 |
| CHAF1A | 6.75E-05 | 3.47050196 | 1 |
| ZNF695 | 0.02334027 | 3.42047209 | 1 |
| PRAMEF11 | 0.02334027 | 3.37380943 | 1 |
| ZDHHC15 | 0.00098186 | 3.28186221 | 1 |
| SERPINB12 | 0.03191766 | 3.27129448 | 1 |
| TCEAL6 | 0.00666945 | 3.23427087 | 1 |
| RUNX3 | 0.00070866 | 3.21779094 | 1 |
| ULBP3 | 0.01248949 | 3.21777071 | 1 |
| ZNF676 | 0.00051031 | 3.21521752 | 1 |
| SST | 0.00135096 | 3.21129857 | 1 |
| CEP85 | 0.00013388 | 3.17270299 | 1 |
| LILRA6 | 0.00486606 | 3.1555587 | 1 |
| MS4A10 | 0.02334027 | 3.13950153 | 1 |
| OLR1 | 0.00018779 | 3.13662643 | 1 |
| MPL | 0.04369348 | 3.11309881 | 1 |
| FCRL5 | 0.00486606 | 3.09693042 | 1 |
| ZIC4 | 0.04369348 | 3.09058331 | 1 |
| CAMK2N2 | 0.02334027 | 3.08922803 | 1 |
| DENND2C | 0.00036661 | 3.07887532 | 1 |
| VENTX | 0.00257911 | 3.07133885 | 1 |
| KCTD16 | 0.02334027 | 3.06377802 | 1 |
| NDST2 | 7.70E-05 | 3.04185581 | 1 |
| KNTC1 | 0.00098186 | 3.03143833 | 1 |
| SLC29A3 | 0.00051031 | 3.02787001 | 1 |
| TLX2 | 0.01248949 | 3.02230655 | 1 |
| TRIM15 | 0.00135745 | 3.00518781 | 1 |
| VRTN | 0.02334027 | 2.99169851 | 1 |
| TGM1 | 0.00666945 | 2.98230511 | 1 |
| ADPRHL1 | 0.00257911 | 2.97854642 | 1 |
| SPIC | 0.00098186 | 2.97766029 | 1 |
| NHLH1 | 0.04369348 | 2.9703991 | 1 |
| KLK5 | 0.00051031 | 2.96810977 | 1 |
| PNMA6A | 0.00486606 | 2.96089551 | 1 |
| AIM2 | 0.00354541 | 2.95944651 | 1 |
| ACTA1 | 0.03191766 | 2.95895062 | 1 |
| VEGFD | 0.00051031 | 2.9440419 | 1 |
| SCN1A | 0.01707465 | 2.94186193 | 1 |
| CXCL2 | 0.00070866 | 2.93991142 | 1 |
| GZMK | 0.01248949 | 2.93318642 | 1 |
| OR11H6 | 0.02334027 | 2.92990112 | 1 |
| ZNF396 | 0.01248949 | 2.92293721 | 1 |
| HENMT1 | 0.00051031 | 2.92244523 | 1 |
| ATOH7 | 0.00486606 | 2.90965202 | 1 |
| ZNF607 | 0.01707465 | 2.90632094 | 1 |
| LAIR2 | 0.03191766 | 2.9019749 | 1 |
| NR0B2 | 0.01248949 | 2.89928727 | 1 |
| MYOCOS | 0.01707465 | 2.89803798 | 1 |

| | | | |
|---|---|---|---|
| CCL3 | 6.29E-05 | 2.89539114 | 1 |
| L1TD1 | 0.00486606 | 2.89100919 | 1 |
| C19orf33 | 0.00135745 | 2.88832878 | 1 |
| ZNF850 | 0.01707465 | 2.87988276 | 1 |
| ADGRF2 | 0.00354541 | 2.85307327 | 1 |
| ANXA8 | 0.03191766 | 2.85240499 | 1 |
| TNFSF9 | 0.02334027 | 2.84955828 | 1 |
| SLC35E4 | 0.00135745 | 2.81965908 | 1 |
| CREB3L4 | 0.00098186 | 2.81563417 | 1 |
| CXCL9 | 0.00113901 | 2.81474693 | 1 |
| CDH12 | 0.01248949 | 2.81263295 | 1 |
| TSHB | 0.02334027 | 2.8118127 | 1 |
| MT3 | 0.00187288 | 2.79054165 | 1 |
| PCDHA12 | 0.01391824 | 2.78658031 | 1 |
| OVOL2 | 0.00666945 | 2.78380919 | 1 |
| NKX2-6 | 0.00257911 | 2.77917134 | 1 |
| MRGPRX2 | 0.03191766 | 2.77779758 | 1 |
| ZNF792 | 0.00218359 | 2.77132362 | 1 |
| GPRIN1 | 0.04369348 | 2.7645698 | 1 |
| AZU1 | 0.04369348 | 2.75526907 | 1 |
| OR2H2 | 0.04369348 | 2.75299177 | 1 |
| IL2RB | 0.00354541 | 2.75106788 | 1 |
| CXCR5 | 0.02334027 | 2.73984247 | 1 |
| POMC | 0.01248949 | 2.73409218 | 1 |
| DEFA3 | 0.02334027 | 2.73218226 | 1 |
| BECN2 | 0.03191766 | 2.71584365 | 1 |
| PCDHB5 | 0.00051031 | 2.69533465 | 1 |
| USP17L2 | 0.00032137 | 2.6932786 | 1 |
| SCYGR4 | 0.03191766 | 2.69284162 | 1 |
| VN1R4 | 0.04369348 | 2.69087938 | 1 |
| TTC26 | 0.00019128 | 2.69028188 | 1 |
| TACC3 | 7.84E-05 | 2.69018313 | 1 |
| INS | 0.04369348 | 2.68921858 | 1 |
| CHRM2 | 0.04369348 | 2.68465871 | 1 |
| CEACAM5 | 0.02334027 | 2.65669669 | 1 |
| C16orf95 | 0.00098186 | 2.65546956 | 1 |
| SMOC1 | 0.00036661 | 2.65174096 | 1 |
| LCE2A | 0.01707465 | 2.63724272 | 1 |
| ITLN2 | 0.00913074 | 2.6241273 | 1 |
| ICAM5 | 0.01248949 | 2.61957091 | 1 |
| SDK1 | 0.00041039 | 2.59712382 | 1 |
| PAOX | 0.00085411 | 2.59538971 | 1 |
| CACNA1G | 0.02334027 | 2.58809248 | 1 |
| KERA | 0.01248949 | 2.57342494 | 1 |
| SPTA1 | 0.01707465 | 2.56561835 | 1 |
| IGHE | 0.00666945 | 2.56214374 | 1 |
| CD5L | 0.00027839 | 2.55610206 | 1 |
| CNTFR | 0.01707465 | 2.55415291 | 1 |
| SMYD1 | 0.00666945 | 2.55294044 | 1 |
| DHRS7C | 0.03191766 | 2.55028436 | 1 |
| SPOCD1 | 0.00486606 | 2.54181064 | 1 |
| SGPP2 | 0.00052201 | 2.54034385 | 1 |
| OR2A12 | 0.02334027 | 2.53447072 | 1 |

| | | | |
|---|---|---|---|
| FBXO24 | 0.00486606 | 2.53432571 | 1 |
| HS3ST1 | 0.00257911 | 2.53407288 | 1 |
| MFSD4A | 0.01248949 | 2.53377782 | 1 |
| ARHGAP15 | 8.11E-05 | 2.53201486 | 1 |
| NR4A2 | 0.00354541 | 2.53031203 | 1 |
| CD300E | 0.00057835 | 2.53020576 | 1 |
| OR4C13 | 0.04369348 | 2.5281152 | 1 |
| FAM71F2 | 0.00354541 | 2.52792333 | 1 |
| THAP10 | 0.01707465 | 2.52198611 | 1 |
| USP43 | 0.00257911 | 2.51569231 | 1 |
| KRT85 | 0.00486606 | 2.5129079 | 1 |
| GZMA | 0.00486606 | 2.50645295 | 1 |
| SAMD15 | 0.03191766 | 2.50426096 | 1 |
| AIFM3 | 0.01798766 | 2.50394281 | 1 |
| SLURP2 | 0.00913074 | 2.49756269 | 1 |
| TMEM225B | 0.00666945 | 2.49490129 | 1 |
| WDR31 | 0.0054824 | 2.47846584 | 1 |
| ZSWIM3 | 0.00354541 | 2.47817664 | 1 |
| WNT6 | 0.00070866 | 2.4775299 | 1 |
| CRCT1 | 0.01248949 | 2.47572101 | 1 |
| UPK1B | 0.00486606 | 2.47453954 | 1 |
| UTP14A | 0.00257911 | 2.47108525 | 1 |
| PHOX2B | 0.01707465 | 2.43492009 | 1 |
| FCRL6 | 0.04369348 | 2.43478523 | 1 |
| CCDC112 | 0.00187288 | 2.43127244 | 1 |
| C11orf65 | 0.00187288 | 2.42542586 | 1 |
| LY75 | 0.00043161 | 2.42527956 | 1 |
| PRRG3 | 0.03191766 | 2.42341701 | 1 |
| ACTG2 | 0.01707465 | 2.4195494 | 1 |
| DDX25 | 0.00666945 | 2.4155624 | 1 |
| RFX2 | 0.01075913 | 2.41440823 | 1 |
| SLC4A3 | 0.02334027 | 2.41303265 | 1 |
| TNFAIP6 | 0.01248949 | 2.4128736 | 1 |
| NHLRC4 | 0.00082322 | 2.41151755 | 1 |
| UGT1A4 | 0.00486606 | 2.41148214 | 1 |
| PLAC8 | 0.01707465 | 2.40531084 | 1 |
| C3orf62 | 0.00913074 | 2.40251438 | 1 |
| CXCL8 | 0.00106926 | 2.40165637 | 1 |
| MRLN | 0.00486606 | 2.39998257 | 1 |
| HCAR3 | 0.01248949 | 2.39776293 | 1 |
| SLC17A8 | 0.02334027 | 2.39612866 | 1 |
| KCNK15 | 0.00666945 | 2.39565572 | 1 |
| NMB | 0.00354541 | 2.39528802 | 1 |
| STAMBPL1 | 0.0033313 | 2.38269263 | 1 |
| MEDAG | 0.04369348 | 2.38234096 | 1 |
| POP1 | 0.00012003 | 2.38177357 | 1 |
| SLA2 | 0.02334027 | 2.37541362 | 1 |
| RD3L | 0.02334027 | 2.37180105 | 1 |
| DNAJB8 | 0.03191766 | 2.36482338 | 1 |
| SFTPD | 0.00257911 | 2.3621446 | 1 |
| TENT5D | 0.04369348 | 2.35828455 | 1 |
| ADAMTSL4 | 0.00084011 | 2.35732873 | 1 |
| C17orf113 | 0.00060802 | 2.35459879 | 1 |

| | | | |
|---|---|---|---|
| C4orf48 | 0.00060962 | 2.35081917 | 1 |
| C6orf141 | 0.01248949 | 2.34899252 | 1 |
| OTULINL | 0.00025517 | 2.3477973 | 1 |
| ZWILCH | 0.02663715 | 2.34555713 | 1 |
| CTXND1 | 0.04369348 | 2.3455286 | 1 |
| KLRB1 | 0.04845233 | 2.34374998 | 1 |
| C14orf180 | 0.01883557 | 2.34289833 | 1 |
| CD163L1 | 0.04369348 | 2.34066212 | 1 |
| TMEM95 | 0.02334027 | 2.33995439 | 1 |
| SGCA | 0.0016181 | 2.33277258 | 1 |
| ZNF773 | 0.00486606 | 2.32756334 | 1 |
| FGFBP3 | 0.01370469 | 2.3243442 | 1 |
| AFF3 | 0.00017391 | 2.32363478 | 1 |
| CEL | 0.00059798 | 2.32285775 | 1 |
| ODF4 | 0.02334027 | 2.32169737 | 1 |
| IGHG3 | 0.03978147 | 2.32119904 | 1 |
| LCE1C | 0.02334027 | 2.32087528 | 1 |
| EPHA3 | 0.00060802 | 2.32029003 | 1 |
| SEMA3E | 0.00913074 | 2.3200143 | 1 |
| STX1A | 0.01248949 | 2.31947121 | 1 |
| NAA11 | 0.01248949 | 2.31608089 | 1 |
| GPR137C | 0.01248949 | 2.31490601 | 1 |
| XDH | 0.00913074 | 2.30911166 | 1 |
| CSNKA2IP | 0.04369348 | 2.2932367 | 1 |
| QRICH2 | 0.00257911 | 2.29180909 | 1 |
| ZNF630 | 0.00119669 | 2.29060132 | 1 |
| ZBTB32 | 0.04369348 | 2.2886225 | 1 |
| SSTR2 | 0.00913074 | 2.28824215 | 1 |
| USP17L1 | 0.01248949 | 2.28469853 | 1 |
| OR52W1 | 0.04369348 | 2.28258933 | 1 |
| CCDC173 | 0.04369348 | 2.28002649 | 1 |
| PRSS38 | 0.04369348 | 2.27814042 | 1 |
| FBXL7 | 5.74E-05 | 2.27548305 | 1 |
| SPRR1B | 0.01707465 | 2.26701142 | 1 |
| CXorf21 | 0.00032137 | 2.26586152 | 1 |
| CNTN5 | 0.00486606 | 2.26483032 | 1 |
| OR52J3 | 0.04369348 | 2.2589703 | 1 |
| ENPP3 | 0.00043161 | 2.25718062 | 1 |
| KRTAP11-1 | 0.01248949 | 2.25683287 | 1 |
| TRIM7 | 0.00045706 | 2.25662765 | 1 |
| SLC16A8 | 0.04369348 | 2.25625023 | 1 |
| PAPPA2 | 0.01707465 | 2.25549425 | 1 |
| HOXD12 | 0.01248949 | 2.25106198 | 1 |
| EYA1 | 0.00913074 | 2.25066528 | 1 |
| MS4A14 | 0.00042443 | 2.24904178 | 1 |
| TPH1 | 0.00666945 | 2.24653495 | 1 |
| CD70 | 0.01707465 | 2.24535141 | 1 |
| KNL1 | 0.00913074 | 2.2452491 | 1 |
| TIMP4 | 0.00257911 | 2.24426106 | 1 |
| CFAP161 | 0.03191766 | 2.24292763 | 1 |
| MRM1 | 0.00211391 | 2.24016824 | 1 |
| NUTM2A | 0.02334027 | 2.23999935 | 1 |
| GPR63 | 0.02334027 | 2.23864094 | 1 |

| | | | |
|---|---|---|---|
| COQ2 | 0.00015926 | 2.23850534 | 1 |
| GEN1 | 0.00401617 | 2.23739376 | 1 |
| SIGLEC15 | 0.01707465 | 2.23619056 | 1 |
| GSX1 | 0.01248949 | 2.23201235 | 1 |
| OR13C9 | 0.00486606 | 2.22376525 | 1 |
| FAM218A | 0.00666945 | 2.22243776 | 1 |
| AVP | 0.01248949 | 2.21922811 | 1 |
| IRX6 | 0.04369348 | 2.20743205 | 1 |
| CEP295NL | 0.00044633 | 2.20710504 | 1 |
| ZSWIM7 | 0.00031678 | 2.20637155 | 1 |
| OR13C2 | 0.04369348 | 2.20635523 | 1 |
| LRAT | 0.00666945 | 2.20330645 | 1 |
| TRDC | 0.00354541 | 2.20246237 | 1 |
| EIF3E | 0.01707465 | 2.20221397 | 1 |
| TRPV5 | 0.01248949 | 2.20151025 | 1 |
| OR4K13 | 0.00486606 | 2.19588115 | 1 |
| ANKK1 | 0.04369348 | 2.19172182 | 1 |
| PSORS1C1 | 0.00913074 | 2.18893777 | 1 |
| ZNF765 | 0.00119649 | 2.1798373 | 1 |
| PLPP2 | 0.00583111 | 2.17905783 | 1 |
| KRT82 | 0.03191766 | 2.1782984 | 1 |
| TGFBR3L | 0.02334027 | 2.17790451 | 1 |
| HIST1H4J | 0.01707465 | 2.17670755 | 1 |
| PGA5 | 0.01248949 | 2.17617234 | 1 |
| SCRT1 | 0.03191766 | 2.17249226 | 1 |
| RASGRP4 | 0.00085411 | 2.17157427 | 1 |
| HOMER2 | 0.00486606 | 2.17112463 | 1 |
| TREML1 | 0.01707465 | 2.17028098 | 1 |
| TMEM171 | 0.00121624 | 2.16920642 | 1 |
| AGXT | 0.04369348 | 2.16778967 | 1 |
| ARHGEF4 | 0.01707465 | 2.16337578 | 1 |
| CCDC170 | 0.01826651 | 2.16314844 | 1 |
| CLPS | 0.01248949 | 2.16293018 | 1 |
| KDF1 | 0.00913074 | 2.16173676 | 1 |
| FAM177B | 0.01707465 | 2.16118885 | 1 |
| ADAD1 | 0.02334027 | 2.15992323 | 1 |
| SMKR1 | 0.01248949 | 2.15634075 | 1 |
| HTR1F | 0.02334027 | 2.1560471 | 1 |
| CD80 | 0.00060006 | 2.15485686 | 1 |
| AOC3 | 0.0016181 | 2.15449282 | 1 |
| ARHGEF39 | 0.04369348 | 2.15385785 | 1 |
| GABRR2 | 0.00313567 | 2.1523306 | 1 |
| FBXO5 | 0.02623829 | 2.15043623 | 1 |
| SP140 | 0.00159194 | 2.14941507 | 1 |
| C4orf33 | 0.00102748 | 2.14028823 | 1 |
| MAPK8IP2 | 0.01707465 | 2.13933498 | 1 |
| ANKRD34A | 0.00805279 | 2.13670536 | 1 |
| CENPW | 0.00115788 | 2.13644744 | 1 |
| GP9 | 0.04369348 | 2.13521849 | 1 |
| ACKR2 | 0.00666945 | 2.13327083 | 1 |
| RTL4 | 0.01248949 | 2.1295881 | 1 |
| NLRP11 | 0.01248949 | 2.12520006 | 1 |
| CCDC77 | 0.00225537 | 2.12379544 | 1 |

| | | | |
|---|---|---|---|
| LEFTY1 | 0.00376678 | 2.11869951 | 1 |
| NIM1K | 0.02334027 | 2.11783184 | 1 |
| GPLD1 | 0.01248949 | 2.11478553 | 1 |
| DES | 0.00573986 | 2.11381521 | 1 |
| NMRK2 | 0.00913074 | 2.11312226 | 1 |
| FRMPD2 | 0.00913074 | 2.11180773 | 1 |
| TIMD4 | 0.01971824 | 2.11170089 | 1 |
| OXCT2 | 0.01826651 | 2.10972613 | 1 |
| ABCC11 | 0.01707465 | 2.10342074 | 1 |
| CLCA4 | 0.03191766 | 2.10264174 | 1 |
| SSTR5 | 0.02334027 | 2.10044089 | 1 |
| MOV10L1 | 0.00913074 | 2.10029961 | 1 |
| EPHA5 | 0.04245894 | 2.09854672 | 1 |
| KIRREL3 | 0.02334027 | 2.0979224 | 1 |
| REM1 | 0.00913074 | 2.09683832 | 1 |
| STAP1 | 0.00995342 | 2.09676995 | 1 |
| CFAP73 | 0.00486606 | 2.09461905 | 1 |
| PCDHA3 | 0.03594207 | 2.09404237 | 1 |
| WNT2 | 0.01248949 | 2.09346674 | 1 |
| DNAH8 | 0.01707465 | 2.09304622 | 1 |
| IRX2 | 0.00117704 | 2.09244787 | 1 |
| GAL3ST3 | 0.00086833 | 2.08799635 | 1 |
| PAQR6 | 0.02334027 | 2.08732529 | 1 |
| PLAC8L1 | 0.01707465 | 2.08599956 | 1 |
| CEBPE | 0.00666945 | 2.08247141 | 1 |
| CFAP77 | 0.01248949 | 2.07890916 | 1 |
| ADGRE2 | 0.000443 | 2.07587442 | 1 |
| FCGR1A | 0.00913074 | 2.07543842 | 1 |
| ZNF660 | 0.00354541 | 2.07346096 | 1 |
| ADAMTS3 | 0.01248949 | 2.07135511 | 1 |
| CCNE1 | 0.01370469 | 2.07029997 | 1 |
| FER1L5 | 0.04369348 | 2.06669881 | 1 |
| CDC45 | 0.00666945 | 2.06494965 | 1 |
| FPR1 | 0.00298834 | 2.06469494 | 1 |
| AKAP3 | 0.01075913 | 2.06452486 | 1 |
| TRIM46 | 0.00666945 | 2.06222885 | 1 |
| NMNAT2 | 0.01248949 | 2.05944951 | 1 |
| SLC22A25 | 0.04369348 | 2.05430742 | 1 |
| ACP7 | 0.01707465 | 2.05172323 | 1 |
| TCP10L | 0.02545603 | 2.05113917 | 1 |
| OR10A7 | 0.02334027 | 2.04988279 | 1 |
| PRR35 | 0.00667218 | 2.04910745 | 1 |
| NRL | 0.00032682 | 2.04637934 | 1 |
| GPRC6A | 0.03191766 | 2.04421608 | 1 |
| AC132217.2 | 0.01707465 | 2.04179451 | 1 |
| OR4K1 | 0.01248949 | 2.03709428 | 1 |
| NPTX1 | 0.01457662 | 2.0354812 | 1 |
| HRNR | 0.01248949 | 2.03486685 | 1 |
| GSG1L | 0.02334027 | 2.0337937 | 1 |
| UQCRHL | 0.01707465 | 2.02629915 | 1 |
| FAM117B | 0.00134738 | 2.02428372 | 1 |
| CCT8L2 | 0.00229206 | 2.02358265 | 1 |
| RNASE11 | 0.0342393 | 2.02260479 | 1 |

| | | | |
|---|---|---|---|
| C2orf83 | 0.02334027 | 2.01755098 | 1 |
| RCC1 | 0.01248949 | 2.0173825 | 1 |
| TMSB15B.1 | 0.03191766 | 2.01614559 | 1 |
| CES4A | 0.00033237 | 2.01536627 | 1 |
| GPR39 | 0.01011012 | 2.01351184 | 1 |
| SUV39H2 | 0.00913074 | 2.01015749 | 1 |
| POU2F2 | 0.00024279 | 2.00962531 | 1 |
| SSU72P8 | 0.03191766 | 2.00350106 | 1 |
| NACAD | 0.02334027 | 1.99685746 | 1 |
| OR5J2 | 0.03191766 | 1.99643388 | 1 |
| SCN2B | 0.02663715 | 1.99582919 | 1 |
| SH3PXD2B | 0.00159194 | 1.99546279 | 1 |
| NEURL2 | 0.000338 | 1.99431403 | 1 |
| PGBD5 | 0.01248949 | 1.99316822 | 1 |
| ZFP42 | 0.0078049 | 1.99205404 | 1 |
| NT5C3A | 0.00024279 | 1.99178232 | 1 |
| CER1 | 0.02334027 | 1.9915458 | 1 |
| SMIM21 | 0.04369348 | 1.98844629 | 1 |
| ALKBH8 | 0.01026902 | 1.98670779 | 1 |
| PODNL1 | 0.00486606 | 1.98612541 | 1 |
| IL1RAPL1 | 0.04369348 | 1.98451161 | 1 |
| FA2H | 0.03191766 | 1.98406944 | 1 |
| CD1D | 0.00421295 | 1.98249775 | 1 |
| SPINK5 | 0.01248949 | 1.9816102 | 1 |
| PGLYRP3 | 0.03191766 | 1.98056514 | 1 |
| CHRNA3 | 0.02334027 | 1.97759186 | 1 |
| MIXL1 | 0.04369348 | 1.97757988 | 1 |
| CRLF1 | 0.00732954 | 1.97692984 | 1 |
| DBX2 | 0.03191766 | 1.97689581 | 1 |
| 1-Mar | 9.32E-05 | 1.97403717 | 1 |
| RLIM | 0.04634872 | 1.97025767 | 1 |
| ATP8B4 | 0.00026336 | 1.96945339 | 1 |
| CYP7A1 | 0.00913074 | 1.96888366 | 1 |
| KLRG1 | 0.01391824 | 1.9646384 | 1 |
| NR1I3 | 0.00221921 | 1.96308094 | 1 |
| DCPS | 0.00023387 | 1.9613871 | 1 |
| LCE2C | 0.01248949 | 1.96083927 | 1 |
| ZP3 | 0.01707465 | 1.95797018 | 1 |
| FITM1 | 0.01707465 | 1.9570031 | 1 |
| ABO | 0.00121624 | 1.95681422 | 1 |
| MIA | 0.01248949 | 1.95407489 | 1 |
| TPD52L3 | 0.02334027 | 1.95268675 | 1 |
| PRF1 | 0.00086833 | 1.9525474 | 1 |
| KCND2 | 0.01248949 | 1.95242091 | 1 |
| EFNA5 | 0.00032824 | 1.95173724 | 1 |
| CROCC2 | 0.02334027 | 1.94994269 | 1 |
| COL11A1 | 0.01248949 | 1.94988346 | 1 |
| OSM | 0.00486606 | 1.94918759 | 1 |
| SLC30A3 | 0.03191766 | 1.94886658 | 1 |
| OR4C11 | 0.02334027 | 1.94825625 | 1 |
| ILDR1 | 0.007914 | 1.94806681 | 1 |
| HS3ST5 | 0.04369348 | 1.94781991 | 1 |
| GRIK1 | 0.01707465 | 1.94699485 | 1 |

| | | | |
|---|---|---|---|
| SLC34A2 | 0.01707465 | 1.94638804 | 1 |
| CRMP1 | 0.02334027 | 1.9417565 | 1 |
| TCERG1L | 0.03191766 | 1.9412764 | 1 |
| CREG2 | 0.03191766 | 1.94066253 | 1 |
| IRAK2 | 0.00023872 | 1.93998033 | 1 |
| ACTC1 | 0.00913074 | 1.93919351 | 1 |
| PRSS1 | 0.03191766 | 1.93238731 | 1 |
| SIRT4 | 0.00372471 | 1.93059453 | 1 |
| CDH17 | 0.01707465 | 1.92802649 | 1 |
| PPL | 0.00167161 | 1.92658505 | 1 |
| ZNF79 | 0.00666945 | 1.92377478 | 1 |
| KCNK2 | 0.00913074 | 1.92059596 | 1 |
| MGAT4D | 0.02334027 | 1.91743528 | 1 |
| SYCP1 | 0.04369348 | 1.9168073 | 1 |
| OR14C36 | 0.04369348 | 1.91536321 | 1 |
| LRRC75A | 0.03191766 | 1.91465339 | 1 |
| SVOPL | 0.02334027 | 1.91445561 | 1 |
| SCG2 | 0.03191766 | 1.91424877 | 1 |
| IRX5 | 0.00089742 | 1.91212258 | 1 |
| TAF7L | 0.02704124 | 1.91074617 | 1 |
| UBQLN4 | 0.00017258 | 1.90976378 | 1 |
| EREG | 0.01248949 | 1.90430978 | 1 |
| ACBD7 | 0.02334027 | 1.90393268 | 1 |
| TMPRSS4 | 0.02334027 | 1.89788095 | 1 |
| ALOX15B | 0.03191766 | 1.89689882 | 1 |
| SH3TC2 | 0.03191766 | 1.89287715 | 1 |
| WDR97 | 0.02334027 | 1.89214529 | 1 |
| UMODL1 | 0.02334027 | 1.88731932 | 1 |
| LIN7B | 0.0004698 | 1.88551593 | 1 |
| VWA2 | 0.01092706 | 1.88516293 | 1 |
| CCNB3 | 0.03191766 | 1.88443587 | 1 |
| LRRC23 | 0.01003053 | 1.88402391 | 1 |
| GATA6 | 0.00913074 | 1.88303859 | 1 |
| CDC6 | 0.01248949 | 1.88178235 | 1 |
| ZXDB | 0.00401617 | 1.88126034 | 1 |
| IL1R2 | 0.00063911 | 1.86944792 | 1 |
| PIGBOS1 | 0.01043014 | 1.86934878 | 1 |
| KCNC2 | 0.03191766 | 1.86859627 | 1 |
| LCE1A | 0.02334027 | 1.86857606 | 1 |
| IL11 | 0.03191766 | 1.86761902 | 1 |
| REELD1 | 0.01248949 | 1.86544399 | 1 |
| ARRDC5 | 0.00486606 | 1.86258688 | 1 |
| C9orf135 | 0.02334027 | 1.85978292 | 1 |
| TMEM177 | 0.00066064 | 1.85971015 | 1 |
| KLRD1 | 0.00883499 | 1.85915497 | 1 |
| CCDC57 | 8.73E-05 | 1.85877052 | 1 |
| GREB1 | 0.01912588 | 1.85566221 | 1 |
| SYT16 | 0.02334027 | 1.85435993 | 1 |
| SLC2A3 | 0.01248949 | 1.85231805 | 1 |
| OR2T1 | 0.00913074 | 1.85158828 | 1 |
| OR6C74 | 0.04369348 | 1.850089 | 1 |
| GNA15 | 0.00167161 | 1.84930044 | 1 |
| GBP3 | 0.00017871 | 1.84885762 | 1 |

| | | | |
|---|---|---|---|
| ZNF570 | 0.00556171 | 1.84845189 | 1 |
| ANKRD30A | 0.03191766 | 1.84791003 | 1 |
| C4orf36 | 0.03191766 | 1.84528632 | 1 |
| PATL2 | 0.03191766 | 1.84249391 | 1 |
| PLIN1 | 0.03191766 | 1.84118074 | 1 |
| KISS1 | 0.04369348 | 1.84019399 | 1 |
| OR10C1 | 0.01707465 | 1.83748547 | 1 |
| HOXB1 | 0.01707465 | 1.83715344 | 1 |
| CYP39A1 | 0.01912588 | 1.83194937 | 1 |
| APCS | 0.02334027 | 1.83190014 | 1 |
| SLITRK5 | 0.01707465 | 1.83030793 | 1 |
| IFNE | 0.02334027 | 1.82719383 | 1 |
| CST8 | 0.03191766 | 1.82677702 | 1 |
| MSS51 | 0.0055612 | 1.82580958 | 1 |
| TRIM63 | 0.01707465 | 1.82469525 | 1 |
| ZFR2 | 0.03191766 | 1.82413171 | 1 |
| FAM104A | 0.00780312 | 1.82195746 | 1 |
| IL15 | 0.00318624 | 1.81960264 | 1 |
| OR9I1 | 0.04369348 | 1.81930397 | 1 |
| ZSWIM2 | 0.03191766 | 1.81897574 | 1 |
| CSRP2 | 0.00025504 | 1.81681188 | 1 |
| BPI | 0.04369348 | 1.81676191 | 1 |
| SLC8A3 | 0.04369348 | 1.81453419 | 1 |
| BLID | 0.00913074 | 1.81304504 | 1 |
| EFHB | 0.01011012 | 1.81278107 | 1 |
| CECR2 | 0.00104983 | 1.81258502 | 1 |
| GCG | 0.01707465 | 1.80981742 | 1 |
| MOS | 0.01248949 | 1.80859135 | 1 |
| SYT5 | 0.02334027 | 1.80668561 | 1 |
| CUZD1 | 0.03191766 | 1.80552032 | 1 |
| KCNJ8 | 0.00605973 | 1.80438715 | 1 |
| OLAH | 0.02334027 | 1.80302494 | 1 |
| EPO | 0.02334027 | 1.80154426 | 1 |
| CD33 | 0.01251904 | 1.80004592 | 1 |
| MLANA | 0.02334027 | 1.79704386 | 1 |
| MTNR1B | 0.01707465 | 1.79621104 | 1 |
| PPP1R17 | 0.03191766 | 1.79495835 | 1 |
| VGLL2 | 0.03191766 | 1.79246996 | 1 |
| KBTBD3 | 0.0003939 | 1.79087699 | 1 |
| KLF1 | 0.03191766 | 1.79044737 | 1 |
| HAPLN4 | 0.01707465 | 1.78964832 | 1 |
| SIRPD | 0.04369348 | 1.78841993 | 1 |
| CCL7 | 0.03191766 | 1.78816211 | 1 |
| IHH | 0.02334027 | 1.78630892 | 1 |
| CENPI | 0.01890495 | 1.78617837 | 1 |
| VSX2 | 0.04369348 | 1.78514119 | 1 |
| ACSM5 | 0.00049689 | 1.78341362 | 1 |
| CC2D2B | 0.02334027 | 1.77927826 | 1 |
| RBFOX1 | 0.01826651 | 1.77869241 | 1 |
| DCAF4L2 | 0.03191766 | 1.77809225 | 1 |
| AL845331.2 | 0.02334027 | 1.77786163 | 1 |
| MMP3 | 0.03436443 | 1.7757645 | 1 |
| PADI4 | 0.03191766 | 1.77519 | 1 |

| | | | |
|---|---|---|---|
| ST8SIA5 | 0.02334027 | 1.77382997 | 1 |
| HIST3H2A | 0.03648149 | 1.76703002 | 1 |
| PLCD4 | 0.00421295 | 1.76147276 | 1 |
| AC233723.1 | 0.02334027 | 1.76100288 | 1 |
| WDR12 | 6.80E-05 | 1.76086516 | 1 |
| DEF8 | 0.00128682 | 1.75776855 | 1 |
| PPP1R42 | 0.03191766 | 1.75682933 | 1 |
| GLDN | 0.0078049 | 1.7536313 | 1 |
| ZNF891 | 0.00318624 | 1.75222034 | 1 |
| ZNF699 | 0.01017189 | 1.74842198 | 1 |
| JPH2 | 0.02334027 | 1.74397508 | 1 |
| ACOT4 | 0.03049774 | 1.74323027 | 1 |
| NTN3 | 0.04369348 | 1.74216168 | 1 |
| MAMLD1 | 0.00019298 | 1.74154655 | 1 |
| PSTPIP2 | 0.00842844 | 1.73837954 | 1 |
| WDHD1 | 0.0078049 | 1.73793399 | 1 |
| MAB21L3 | 0.03191766 | 1.73763165 | 1 |
| KRT80 | 0.01707465 | 1.73734513 | 1 |
| NMU | 0.02334027 | 1.7357264 | 1 |
| PFN3 | 0.04369348 | 1.73505175 | 1 |
| RPP14 | 0.01707465 | 1.73349076 | 1 |
| MAK | 0.01248949 | 1.73272186 | 1 |
| FBLN7 | 0.02334027 | 1.73218732 | 1 |
| NBPF3 | 0.02663715 | 1.73041231 | 1 |
| NTS | 0.02334027 | 1.72978135 | 1 |
| ZNF10 | 0.00744588 | 1.72932388 | 1 |
| PCSK2 | 0.02334027 | 1.72787293 | 1 |
| CYP4F11 | 0.04369348 | 1.72464023 | 1 |
| LY6L | 0.03191766 | 1.72316776 | 1 |
| IL6 | 0.02334027 | 1.72177267 | 1 |
| MAGEA4 | 0.02334027 | 1.72175207 | 1 |
| MC1R | 0.02623829 | 1.71726445 | 1 |
| CCNJ | 0.04369348 | 1.71250608 | 1 |
| GBGT1 | 0.00047716 | 1.71220592 | 1 |
| IGSF6 | 0.00035842 | 1.71033882 | 1 |
| RBP2 | 0.04369348 | 1.70948551 | 1 |
| CEACAM7 | 0.04369348 | 1.70930572 | 1 |
| ABCE1 | 0.00035842 | 1.70782166 | 1 |
| MED31 | 0.00768354 | 1.70557089 | 1 |
| SYT17 | 0.02469403 | 1.70177368 | 1 |
| XKR3 | 0.02334027 | 1.70140099 | 1 |
| REXO5 | 0.02334027 | 1.70044242 | 1 |
| HP | 0.00805279 | 1.7003799 | 1 |
| LILRB4 | 0.01248949 | 1.69922595 | 1 |
| TPTE | 0.01707465 | 1.69902225 | 1 |
| SLC25A14 | 0.0348836 | 1.69833147 | 1 |
| FAM71D | 0.02334027 | 1.69682513 | 1 |
| SINHCAF | 0.01026902 | 1.69556135 | 1 |
| CIB3 | 0.04369348 | 1.69551802 | 1 |
| IL25 | 0.01707465 | 1.69426188 | 1 |
| GPR132 | 0.01031501 | 1.69349682 | 1 |
| B4GALNT2 | 0.04369348 | 1.69318268 | 1 |
| TEC | 0.00583111 | 1.6931009 | 1 |

| | | | |
|---|---|---|---|
| PEG3 | 0.00044289 | 1.6905385 | 1 |
| PROC | 0.00061525 | 1.68975983 | 1 |
| DAB1 | 0.01707465 | 1.68788523 | 1 |
| ADAMDEC1 | 0.0023293 | 1.68785855 | 1 |
| CHRM5 | 0.03191766 | 1.68660721 | 1 |
| ARL14 | 0.02334027 | 1.68037811 | 1 |
| DYRK4 | 0.00024256 | 1.68000897 | 1 |
| MAJIN | 0.04369348 | 1.67958187 | 1 |
| MED27 | 0.00297009 | 1.67845288 | 1 |
| MIP | 0.04369348 | 1.67705147 | 1 |
| OR13C8 | 0.03191766 | 1.67677027 | 1 |
| CENPH | 0.02334027 | 1.67599273 | 1 |
| APC2 | 0.04369348 | 1.67427685 | 1 |
| NUP210L | 0.01707465 | 1.67330544 | 1 |
| RNF17 | 0.04917111 | 1.67128594 | 1 |
| SERPINE3 | 0.04369348 | 1.67082032 | 1 |
| POU3F1 | 0.01248949 | 1.6706229 | 1 |
| IFNA14 | 0.01707465 | 1.66870614 | 1 |
| NIBAN3 | 0.04369348 | 1.66767929 | 1 |
| CLEC4A | 9.61E-05 | 1.66750164 | 1 |
| ARHGAP11B | 0.04369348 | 1.66725249 | 1 |
| GPR33 | 0.01248949 | 1.66672856 | 1 |
| CTU2 | 0.00224313 | 1.66435089 | 1 |
| CERS3 | 0.02334027 | 1.66174208 | 1 |
| FPR2 | 0.00428046 | 1.66005186 | 1 |
| KRTAP5-9 | 0.04369348 | 1.65758432 | 1 |
| SH3BGR | 0.04369348 | 1.65697618 | 1 |
| EPHA1 | 0.00125665 | 1.65455694 | 1 |
| CCR6 | 0.01435417 | 1.65446785 | 1 |
| AIDA | 6.39E-05 | 1.65423945 | 1 |
| CCDC39 | 0.02334027 | 1.65420013 | 1 |
| FILIP1 | 0.00010744 | 1.65139064 | 1 |
| TJP3 | 0.00573986 | 1.65137645 | 1 |
| CPA3 | 0.04369348 | 1.65109005 | 1 |
| SP9 | 0.03191766 | 1.65063664 | 1 |
| THBS4 | 0.04369348 | 1.64958781 | 1 |
| ZNF48 | 0.00050178 | 1.64418218 | 1 |
| IGHM | 0.03291924 | 1.6439685 | 1 |
| CPXM2 | 0.00019477 | 1.6438996 | 1 |
| GINS4 | 0.04845233 | 1.64358201 | 1 |
| CCDC24 | 0.00084618 | 1.64347083 | 1 |
| ADM2 | 0.000338 | 1.64174538 | 1 |
| BCO1 | 0.02704124 | 1.64043454 | 1 |
| SRL | 0.04369348 | 1.63958841 | 1 |
| ADGRD2 | 0.03191766 | 1.63901166 | 1 |
| LYPD1 | 0.01075913 | 1.63674818 | 1 |
| BIRC3 | 0.00089425 | 1.63577178 | 1 |
| CADM2 | 0.02334027 | 1.6352585 | 1 |
| RFX8 | 0.04566483 | 1.63519023 | 1 |
| SPRY3 | 0.0078049 | 1.63471328 | 1 |
| RHBG | 7.24E-05 | 1.63112511 | 1 |
| EPHB2 | 0.00431983 | 1.63077264 | 1 |
| ADAM8 | 0.00428046 | 1.63013605 | 1 |

| | | | |
|---|---|---|---|
| OR52N1 | 0.03191766 | 1.62710347 | 1 |
| HNRNPCL2 | 0.03191766 | 1.62339051 | 1 |
| NEFM | 0.03191766 | 1.62294456 | 1 |
| HSDL1 | 0.00012815 | 1.6224658 | 1 |
| FBXL13 | 0.01707465 | 1.61866506 | 1 |
| IL23R | 0.04369348 | 1.61820807 | 1 |
| TM4SF19 | 0.02507252 | 1.61708665 | 1 |
| OR2AG2 | 0.04369348 | 1.61686316 | 1 |
| IQANK1 | 0.03334586 | 1.61574067 | 1 |
| ZCCHC18 | 0.02663715 | 1.61490694 | 1 |
| NSUN3 | 0.00022494 | 1.61474868 | 1 |
| SPRED3 | 0.01707465 | 1.61413389 | 1 |
| FOCAD | 0.00622448 | 1.6138811 | 1 |
| MBOAT4 | 0.04369348 | 1.61328842 | 1 |
| XKR4 | 0.00573986 | 1.61241546 | 1 |
| BIN2 | 0.00019607 | 1.61196037 | 1 |
| C16orf91 | 0.0023061 | 1.61184054 | 1 |
| DYTN | 0.04369348 | 1.61170589 | 1 |
| SYNCRIP | 0.02334027 | 1.61150685 | 1 |
| GRM4 | 0.04369348 | 1.6101132 | 1 |
| C10orf71 | 0.04369348 | 1.60992135 | 1 |
| ZBED6CL | 0.00842844 | 1.60946334 | 1 |
| RAG1 | 0.00169897 | 1.60646194 | 1 |
| INA | 0.03191766 | 1.6057514 | 1 |
| TTYH2 | 0.00127132 | 1.60555406 | 1 |
| KLHL34 | 0.04369348 | 1.60549666 | 1 |
| RAB39A | 0.00125665 | 1.60403142 | 1 |
| ZBP1 | 0.02623829 | 1.60169159 | 1 |
| PAX6 | 0.03191766 | 1.60120424 | 1 |
| TMEM272 | 0.04369348 | 1.59986852 | 1 |
| CBLC | 0.00068951 | 1.59819761 | 1 |
| GLRA1 | 0.04369348 | 1.59801809 | 1 |
| GPR61 | 0.03191766 | 1.59675997 | 1 |
| TMEM247 | 0.03191766 | 1.59318262 | 1 |
| PRTFDC1 | 0.01435417 | 1.59262104 | 1 |
| CBLN4 | 0.03191766 | 1.5925976 | 1 |
| OR52E5 | 0.02334027 | 1.59144469 | 1 |
| TRPC7 | 0.02334027 | 1.59130548 | 1 |
| C3orf20 | 0.04369348 | 1.59082761 | 1 |
| COL24A1 | 0.04369348 | 1.58765823 | 1 |
| ATAD2 | 0.00592368 | 1.58290532 | 1 |
| CDCA7L | 0.00123348 | 1.58022704 | 1 |
| CRABP2 | 0.0148021 | 1.57997277 | 1 |
| MAIP1 | 9.72E-05 | 1.57982185 | 1 |
| EDDM3B | 0.03191766 | 1.57750059 | 1 |
| GALNT3 | 0.01391824 | 1.57668642 | 1 |
| RIPK3 | 0.00556171 | 1.57639528 | 1 |
| HOXB8 | 0.00036981 | 1.57582268 | 1 |
| ELOVL3 | 0.04369348 | 1.57537139 | 1 |
| CNMD | 0.04369348 | 1.57442624 | 1 |
| MED26 | 0.00033317 | 1.57400799 | 1 |
| LRRC31 | 0.03191766 | 1.57315769 | 1 |
| NCAPH | 0.04369348 | 1.57141053 | 1 |

| | | | |
|---|---|---|---|
| FLRT1 | 0.04369348 | 1.5704355 | 1 |
| CLGN | 0.01707465 | 1.57023725 | 1 |
| BEST3 | 0.03206424 | 1.5688988 | 1 |
| BZW2 | 0.00051833 | 1.56853362 | 1 |
| TMEM81 | 0.02320012 | 1.56808706 | 1 |
| RASGRP1 | 0.00428046 | 1.5677093 | 1 |
| CYP11A1 | 0.04369348 | 1.56494148 | 1 |
| ATP10B | 0.02334027 | 1.5627491 | 1 |
| PRXL2C | 0.00025374 | 1.56144204 | 1 |
| HIST1H2AE | 0.00172674 | 1.5611743 | 1 |
| LRRTM3 | 0.03191766 | 1.56005357 | 1 |
| PRSS53 | 0.00249636 | 1.55937744 | 1 |
| SLC1A7 | 0.03191766 | 1.55905721 | 1 |
| BDNF | 0.04369348 | 1.55845017 | 1 |
| C12orf74 | 0.04774239 | 1.55807834 | 1 |
| BMP3 | 0.03191766 | 1.55775867 | 1 |
| WDR54 | 0.00601695 | 1.5554876 | 1 |
| OR2B3 | 0.04369348 | 1.55504945 | 1 |
| ONECUT1 | 0.04369348 | 1.55461485 | 1 |
| LEXM | 0.04845233 | 1.55119248 | 1 |
| B3GNT4 | 0.04774239 | 1.55041507 | 1 |
| TRIB3 | 0.00172674 | 1.54972079 | 1 |
| TBC1D2 | 0.00085749 | 1.54844975 | 1 |
| DDX51 | 0.00050961 | 1.54814881 | 1 |
| REEP4 | 0.00323553 | 1.54749862 | 1 |
| TPH2 | 0.03191766 | 1.54559535 | 1 |
| MAP3K15 | 0.01391824 | 1.54463093 | 1 |
| CLDN23 | 0.00366424 | 1.5438206 | 1 |
| PCDHB8 | 0.04369348 | 1.54359582 | 1 |
| ENPP1 | 0.00016844 | 1.53949075 | 1 |
| SHC3 | 0.02334027 | 1.53931902 | 1 |
| SSC4D | 0.00592368 | 1.53901887 | 1 |
| FDXACB1 | 0.03191766 | 1.53807078 | 1 |
| INTS2 | 0.00030577 | 1.53349577 | 1 |
| BHLHE23 | 0.02704124 | 1.53235009 | 1 |
| SRRM4 | 0.04369348 | 1.53031625 | 1 |
| PAQR9 | 0.04369348 | 1.52886546 | 1 |
| SMOC2 | 0.0487174 | 1.52703779 | 1 |
| TFAP4 | 0.0009123 | 1.52679316 | 1 |
| LIPJ | 0.04369348 | 1.52638998 | 1 |
| ZNF286A | 0.00792798 | 1.52545585 | 1 |
| BAIAP2L2 | 0.00034324 | 1.52286883 | 1 |
| NKX2-8 | 0.04369348 | 1.52249356 | 1 |
| C1QTNF4 | 0.00614654 | 1.52109476 | 1 |
| KRBA1 | 0.00043612 | 1.51738505 | 1 |
| ALDH1A3 | 0.00119871 | 1.51565447 | 1 |
| OR2S2 | 0.04369348 | 1.5156392 | 1 |
| C2orf66 | 0.04369348 | 1.51498747 | 1 |
| B3GNTL1 | 0.02545603 | 1.5134904 | 1 |
| P2RY8 | 0.04369348 | 1.5129923 | 1 |
| OR3A3 | 0.04369348 | 1.50815444 | 1 |
| ULBP1 | 0.04369348 | 1.50502084 | 1 |
| PINX1 | 0.01379228 | 1.50461439 | 1 |

| | | | |
|---|---|---|---|
| IGHG1 | 0.0368644 | 1.50207147 | 1 |
| OR8K3 | 0.02334027 | 1.50188207 | 1 |
| TMEM59L | 0.03191766 | 1.50125091 | 1 |
| KIF21B | 0.00236709 | 1.49721213 | 1 |
| TRAT1 | 0.02334027 | 1.48270053 | 1 |
| CHRNB4 | 0.03540946 | 1.48124321 | 1 |
| APOBEC1 | 0.03191766 | 1.48109764 | 1 |
| F7 | 0.03191766 | 1.4810292 | 1 |
| MPHOSPH6 | 0.00153096 | 1.47731207 | 1 |
| HARBI1 | 0.03191766 | 1.47704899 | 1 |
| ACTL9 | 0.04369348 | 1.47649019 | 1 |
| IGFBPL1 | 0.04369348 | 1.47536525 | 1 |
| TNFSF4 | 0.03540946 | 1.47525853 | 1 |
| CCL4 | 0.00614654 | 1.47395741 | 1 |
| OR10A3 | 0.03191766 | 1.47387698 | 1 |
| CD38 | 0.00062255 | 1.47349299 | 1 |
| BCL2L10 | 0.04774239 | 1.473123 | 1 |
| USP50 | 0.02623829 | 1.47169187 | 1 |
| GSTCD | 9.00E-05 | 1.47038073 | 1 |
| IL18 | 0.00101182 | 1.46925263 | 1 |
| BCL2A1 | 0.00079934 | 1.46900042 | 1 |
| SLC28A2 | 0.04369348 | 1.46756735 | 1 |
| ZNF681 | 0.03191766 | 1.46652614 | 1 |
| TLR1 | 0.00138744 | 1.46580906 | 1 |
| CNTNAP3B | 0.00044289 | 1.46446518 | 1 |
| TMEM179 | 0.01457662 | 1.46422325 | 1 |
| TMEM178B | 0.00601756 | 1.4636708 | 1 |
| AIRE | 0.04369348 | 1.46052749 | 1 |
| MAGIX | 0.0032831 | 1.46008052 | 1 |
| INSL3 | 0.04369348 | 1.45970838 | 1 |
| AMBN | 0.04369348 | 1.45793636 | 1 |
| FCHO2 | 0.00046721 | 1.45740395 | 1 |
| ERCC8 | 0.00202038 | 1.45200757 | 1 |
| NDP | 0.04369348 | 1.45156992 | 1 |
| KCNN4 | 0.01092706 | 1.45056658 | 1 |
| NELFE | 0.00010165 | 1.44983365 | 1 |
| GP2 | 0.01457662 | 1.4466134 | 1 |
| PRR5L | 0.00028494 | 1.44596399 | 1 |
| CD1A | 0.04369348 | 1.44231627 | 1 |
| ZNF442 | 0.01092706 | 1.4410424 | 1 |
| OR1L3 | 0.04369348 | 1.43983466 | 1 |
| SULT1E1 | 0.0258446 | 1.43901795 | 1 |
| OR6B1 | 0.04917111 | 1.43856585 | 1 |
| C17orf100 | 0.02663715 | 1.4354852 | 1 |
| C16orf96 | 0.03191766 | 1.43289168 | 1 |
| WDR4 | 0.01007733 | 1.43188489 | 1 |
| ZNF385D | 0.01942008 | 1.43118108 | 1 |
| VWA5B1 | 0.00034372 | 1.43059799 | 1 |
| PHETA1 | 0.00023396 | 1.42567451 | 1 |
| C20orf27 | 0.00441844 | 1.42452331 | 1 |
| PIGC | 0.00062759 | 1.42306454 | 1 |
| TNFRSF10D | 0.00013223 | 1.42302205 | 1 |
| ZNF77 | 0.00456428 | 1.42233635 | 1 |

| | | | |
|---|---|---|---|
| SLC6A2 | 0.04369348 | 1.42007892 | 1 |
| GVQW3 | 0.00817935 | 1.41780477 | 1 |
| RTN4RL1 | 0.0143745 | 1.41776675 | 1 |
| KCNK7 | 0.04704121 | 1.41718489 | 1 |
| ZMYND19 | 0.00019477 | 1.41531472 | 1 |
| MTNR1A | 0.03191766 | 1.41468052 | 1 |
| GPR146 | 0.0023293 | 1.41353758 | 1 |
| C10orf143 | 0.00805279 | 1.41325628 | 1 |
| OR6C70 | 0.04369348 | 1.41303932 | 1 |
| GKAP1 | 0.02514278 | 1.41290973 | 1 |
| KLHL23 | 0.00071101 | 1.41223228 | 1 |
| FKBP1B | 0.03191766 | 1.41181352 | 1 |
| CYP26C1 | 0.02704124 | 1.40834978 | 1 |
| FNDC5 | 0.04369348 | 1.40818156 | 1 |
| DEPDC1B | 0.04774239 | 1.40784262 | 1 |
| ANO9 | 0.00549079 | 1.40445519 | 1 |
| SLAMF7 | 0.01379392 | 1.40432765 | 1 |
| GRID2 | 0.02334027 | 1.40374275 | 1 |
| KRTAP19-6 | 0.04369348 | 1.40285022 | 1 |
| ATP6V1G2 | 0.04566483 | 1.40246728 | 1 |
| SOCS2 | 9.85E-05 | 1.40089251 | 1 |
| TTLL13P | 0.04369348 | 1.40069955 | 1 |
| CFHR2 | 0.03191766 | 1.40020334 | 1 |
| DDIT4L | 0.00013005 | 1.39795763 | 1 |
| ZNF596 | 0.01075913 | 1.39730857 | 1 |
| CBX8 | 0.01778465 | 1.39558287 | 1 |
| LCA5L | 0.0148021 | 1.39333368 | 1 |
| TRGC2 | 0.03191766 | 1.39286196 | 1 |
| PLIN4 | 0.02288935 | 1.39277285 | 1 |
| RGS9BP | 0.01995308 | 1.39264516 | 1 |
| SLC1A3 | 0.00114355 | 1.3911736 | 1 |
| GCK | 0.03648149 | 1.39093452 | 1 |
| SLC7A2 | 0.01045993 | 1.38904237 | 1 |
| ADA | 0.00025924 | 1.3875258 | 1 |
| FCRLB | 0.04369348 | 1.38717283 | 1 |
| ASGR2 | 0.03191766 | 1.38632571 | 1 |
| FCER2 | 0.04369348 | 1.38331785 | 1 |
| LRRN4 | 0.00858802 | 1.382512 | 1 |
| CASS4 | 0.00395465 | 1.38225262 | 1 |
| HOXA1 | 0.04917111 | 1.38210831 | 1 |
| FCGR2B | 0.03191766 | 1.38094833 | 1 |
| PFN4 | 0.03594207 | 1.3798661 | 1 |
| ZNF66 | 0.04369348 | 1.37904103 | 1 |
| TENM4 | 0.02334027 | 1.37816239 | 1 |
| TTLL11 | 0.00097673 | 1.37716803 | 1 |
| CDK3 | 0.00778465 | 1.37602222 | 1 |
| FEZF1 | 0.01942008 | 1.37493103 | 1 |
| MEFV | 0.00323754 | 1.37323118 | 1 |
| CLRN3 | 0.00020934 | 1.372925 | 1 |
| ZNF787 | 0.00066064 | 1.3721432 | 1 |
| DPAGT1 | 0.00025374 | 1.37177084 | 1 |
| CHRNA4 | 0.00814001 | 1.37044636 | 1 |
| ARSJ | 0.00508832 | 1.36616233 | 1 |

| | | | |
|---|---|---|---|
| AFG1L | 0.00054212 | 1.36577424 | 1 |
| IGSF21 | 0.00323754 | 1.36448706 | 1 |
| RAB30 | 0.00456428 | 1.36318907 | 1 |
| ZNF404 | 0.04369348 | 1.36269265 | 1 |
| LRRC3 | 0.04528727 | 1.35894891 | 1 |
| SLC25A47 | 0.04917111 | 1.35874753 | 1 |
| LRGUK | 0.02623829 | 1.35438889 | 1 |
| ZNF232 | 0.00288972 | 1.35409053 | 1 |
| LILRA5 | 9.32E-05 | 1.3506543 | 1 |
| UGT3A1 | 0.0002017 | 1.34926877 | 1 |
| C1orf146 | 0.04369348 | 1.34743473 | 1 |
| CHEK2 | 0.00825516 | 1.34653295 | 1 |
| SOWAHD | 0.04917111 | 1.34637875 | 1 |
| MARCO | 0.00064979 | 1.34500801 | 1 |
| LDB3 | 0.04021005 | 1.3441535 | 1 |
| IKZF2 | 0.00588943 | 1.34376421 | 1 |
| MAD1L1 | 0.00017963 | 1.34096584 | 1 |
| PLA2G4F | 0.00047153 | 1.34033185 | 1 |
| FAM131A | 0.00012261 | 1.33954455 | 1 |
| S1PR5 | 0.03648149 | 1.33850301 | 1 |
| CXCL1 | 0.01942008 | 1.3373118 | 1 |
| RXRG | 0.04369348 | 1.33560672 | 1 |
| EVI2A | 0.00062365 | 1.33362378 | 1 |
| DPEP2 | 0.00125665 | 1.33287788 | 1 |
| ASTN2 | 0.00181161 | 1.33225921 | 1 |
| LOXL2 | 9.72E-05 | 1.33075749 | 1 |
| LAG3 | 0.02663715 | 1.32890877 | 1 |
| HIST1H2BN | 0.03191766 | 1.32454996 | 1 |
| MYO1G | 0.00218175 | 1.32345193 | 1 |
| KCNIP3 | 0.0148021 | 1.3230609 | 1 |
| RELN | 0.00449906 | 1.32278244 | 1 |
| C2CD4B | 0.0342393 | 1.32222836 | 1 |
| FMO3 | 0.01815076 | 1.32202729 | 1 |
| DACT1 | 0.00068924 | 1.3198262 | 1 |
| CXCL11 | 0.02663715 | 1.31898983 | 1 |
| ZKSCAN7 | 0.00323754 | 1.31792038 | 1 |
| HPSE | 0.00297009 | 1.31467335 | 1 |
| TUBA3E | 0.03191766 | 1.31351554 | 1 |
| PLCXD3 | 0.00039798 | 1.31348904 | 1 |
| TMEM184A | 0.00173214 | 1.31260562 | 1 |
| CHRM1 | 0.04774239 | 1.31015461 | 1 |
| COL8A2 | 0.03648149 | 1.30653262 | 1 |
| PLEKHG6 | 0.01916236 | 1.3059965 | 1 |
| RND1 | 0.00236709 | 1.30346813 | 1 |
| EEF1AKMT4 | 0.02704124 | 1.30221567 | 1 |
| GEMIN4 | 0.00318624 | 1.30182382 | 1 |
| OASL | 0.00696261 | 1.30140164 | 1 |
| NUP210 | 0.00034838 | 1.30013073 | 1 |
| RASA4B | 0.03191766 | 1.29917535 | 1 |
| IKZF4 | 0.0007198 | 1.29688576 | 1 |
| HOXA2 | 0.00017689 | 1.29376493 | 1 |
| CALML5 | 0.04369348 | 1.29291512 | 1 |
| DUSP8 | 0.02002038 | 1.29085844 | 1 |

| | | | |
|---|---|---|---|
| TRIM68 | 0.00082322 | 1.2904676 | 1 |
| FKBP14 | 0.01045993 | 1.28954875 | 1 |
| PIAS4 | 0.0001435 | 1.2895059 | 1 |
| SVEP1 | 0.00079393 | 1.28530516 | 1 |
| SEMA3A | 0.00441844 | 1.28334634 | 1 |
| NEXN | 0.00108316 | 1.2831597 | 1 |
| ZNF296 | 0.02616585 | 1.28306707 | 1 |
| HNF4G | 0.00947681 | 1.28249522 | 1 |
| HSPBAP1 | 0.00018733 | 1.28236391 | 1 |
| PIGA | 0.01815076 | 1.27876674 | 1 |
| CTLA4 | 0.04369348 | 1.27867904 | 1 |
| ALDOC | 0.00486304 | 1.27656445 | 1 |
| BLOC1S2 | 0.00097806 | 1.27476186 | 1 |
| SLC22A4 | 0.01075913 | 1.27429325 | 1 |
| OR5M9 | 0.03594207 | 1.27337529 | 1 |
| TSHR | 0.0148021 | 1.27296943 | 1 |
| TENT5C | 0.00029903 | 1.27228422 | 1 |
| LGR5 | 0.04369348 | 1.27188666 | 1 |
| AIFM2 | 0.00010784 | 1.27150056 | 1 |
| PI3 | 0.04369348 | 1.26957269 | 1 |
| IQUB | 0.00463036 | 1.26950315 | 1 |
| LILRB5 | 0.03049774 | 1.26934168 | 1 |
| RUBCNL | 0.0009123 | 1.26815045 | 1 |
| PSTPIP1 | 0.00456428 | 1.26812809 | 1 |
| AC104389.5 | 0.02032656 | 1.26481023 | 1 |
| CLEC4C | 0.04369348 | 1.2636724 | 1 |
| P2RY11 | 0.00087399 | 1.26273544 | 1 |
| AAR2 | 0.0007198 | 1.26249411 | 1 |
| BBS5 | 0.00402732 | 1.26228103 | 1 |
| ATP5PO | 0.04191158 | 1.25901238 | 1 |
| PDRG1 | 0.00196488 | 1.25663325 | 1 |
| B3GAT2 | 0.01092706 | 1.25562763 | 1 |
| KNCN | 0.03191766 | 1.25458199 | 1 |
| SRD5A3 | 0.0028329 | 1.25282096 | 1 |
| CCL8 | 0.02704124 | 1.25245314 | 1 |
| SLAMF8 | 0.00013994 | 1.25146648 | 1 |
| CCR1 | 0.00026143 | 1.24918722 | 1 |
| ZGLP1 | 0.00323754 | 1.24884619 | 1 |
| CA13 | 0.01075913 | 1.24858571 | 1 |
| SLC35G2 | 0.0032831 | 1.24665549 | 1 |
| KITLG | 0.00066934 | 1.24531317 | 1 |
| C5orf49 | 0.04917111 | 1.24201968 | 1 |
| NANOGP8 | 0.04369348 | 1.23819001 | 1 |
| FATE1 | 0.04369348 | 1.23713878 | 1 |
| MMD | 0.00127378 | 1.23709683 | 1 |
| OR2L3 | 0.04369348 | 1.23696524 | 1 |
| C15orf41 | 0.00025131 | 1.23582904 | 1 |
| SIRPB2 | 0.00040606 | 1.23543332 | 1 |
| CYP4F3 | 9.06E-05 | 1.23244257 | 1 |
| POP4 | 0.0005732 | 1.23174662 | 1 |
| ABHD8 | 0.00347981 | 1.23163324 | 1 |
| KRBOX4 | 0.00231882 | 1.23098462 | 1 |
| LRRC42 | 7.32E-05 | 1.229699 | 1 |

| | | | |
|---|---|---|---|
| NUDT13 | 0.0139851 | 1.22902071 | 1 |
| FN1 | 0.0405248 | 1.22607914 | 1 |
| CABP1 | 0.04369348 | 1.22553428 | 1 |
| RARG | 0.00037166 | 1.2250356 | 1 |
| ZNF677 | 6.34E-05 | 1.22475696 | 1 |
| KIAA0825 | 0.00035757 | 1.22407687 | 1 |
| ADRA1D | 0.03191766 | 1.22167718 | 1 |
| MANSC4 | 0.04774239 | 1.22022827 | 1 |
| PPP1R32 | 0.02623829 | 1.21879718 | 1 |
| TRMT9B | 0.02941866 | 1.21667409 | 1 |
| B3GNT3 | 0.04369348 | 1.21539021 | 1 |
| RHOF | 0.00139133 | 1.21536107 | 1 |
| TMEM218 | 0.00049928 | 1.21497037 | 1 |
| C12orf49 | 0.00010225 | 1.21448876 | 1 |
| TENT5B | 0.01092706 | 1.21375194 | 1 |
| CCL24 | 0.03648149 | 1.21371531 | 1 |
| TMEM216 | 0.00070948 | 1.21107835 | 1 |
| PLK1 | 0.00592368 | 1.20807201 | 1 |
| CYCS | 0.00322362 | 1.20793279 | 1 |
| CCNJL | 0.04369348 | 1.20774806 | 1 |
| NFATC2 | 0.00047716 | 1.20721068 | 1 |
| SLC22A18 | 7.37E-05 | 1.20480972 | 1 |
| ZNF74 | 0.02086917 | 1.20296203 | 1 |
| ZNF583 | 0.00109797 | 1.20225594 | 1 |
| IQGAP3 | 0.02704124 | 1.20102836 | 1 |
| GPR135 | 0.00076251 | 1.19562853 | 1 |
| LANCL3 | 0.04369348 | 1.19381029 | 1 |
| CPN2 | 0.00441844 | 1.19364342 | 1 |
| DENND1C | 0.00592368 | 1.19344195 | 1 |
| KIAA1211L | 0.02448013 | 1.19263201 | 1 |
| TCEANC2 | 0.02871767 | 1.19022431 | 1 |
| UBXN8 | 0.00424653 | 1.18997067 | 1 |
| OTOF | 0.04917111 | 1.18990289 | 1 |
| GLMN | 0.00227442 | 1.1896675 | 1 |
| CCDC30 | 0.00198771 | 1.188168 | 1 |
| SHH | 0.00588943 | 1.18415056 | 1 |
| CTRB1 | 0.04369348 | 1.18400366 | 1 |
| MYO19 | 0.00019939 | 1.18366115 | 1 |
| ERG28 | 0.00014476 | 1.18352502 | 1 |
| BNIPL | 0.02704124 | 1.18340794 | 1 |
| SCLY | 0.00342965 | 1.18124842 | 1 |
| KCTD6 | 0.00047716 | 1.18039172 | 1 |
| C19orf57 | 0.04369348 | 1.17904213 | 1 |
| TMTC2 | 0.00042259 | 1.17869316 | 1 |
| SVIP | 0.00323754 | 1.17864735 | 1 |
| GAPVD1 | 0.0070283 | 1.17393284 | 1 |
| BBS10 | 9.44E-05 | 1.1738045 | 1 |
| BZW1 | 0.00055353 | 1.17259471 | 1 |
| CWC22 | 0.00035633 | 1.17159574 | 1 |
| NEGR1 | 0.01037103 | 1.17118593 | 1 |
| OR5T3 | 0.04369348 | 1.17106495 | 1 |
| HIST1H1A | 0.00186603 | 1.16916056 | 1 |
| RAB40B | 0.00032591 | 1.1686867 | 1 |

| | | | |
|---|---|---|---|
| CARD16 | 0.02663715 | 1.16693463 | 1 |
| SCNN1B | 0.00233819 | 1.16674123 | 1 |
| ZNF35 | 0.00020787 | 1.16485215 | 1 |
| NEFH | 0.04369348 | 1.16420621 | 1 |
| DUSP2 | 9.03E-05 | 1.16283841 | 1 |
| LINS1 | 0.00016122 | 1.16252193 | 1 |
| TTC21A | 0.0024239 | 1.16249091 | 1 |
| IGFLR1 | 0.00016493 | 1.16232258 | 1 |
| LIX1 | 0.02032656 | 1.15661832 | 1 |
| PIFO | 0.01497654 | 1.15477996 | 1 |
| POPDC2 | 0.00011847 | 1.15470977 | 1 |
| RRP1 | 0.00026624 | 1.15340035 | 1 |
| CEP95 | 7.11E-05 | 1.15082208 | 1 |
| GDPD3 | 0.00028009 | 1.15073318 | 1 |
| ISM1 | 0.0002024 | 1.15070062 | 1 |
| CAPN8 | 0.03648149 | 1.14896926 | 1 |
| ARHGAP27 | 0.00012679 | 1.14790796 | 1 |
| GXYLT1 | 0.00028945 | 1.1476212 | 1 |
| DNAJB6 | 9.23E-05 | 1.14647473 | 1 |
| LEF1 | 0.02480954 | 1.14555674 | 1 |
| CYP4X1 | 0.00025791 | 1.14543033 | 1 |
| ETV1 | 0.01890495 | 1.14370125 | 1 |
| IRF6 | 0.00106926 | 1.14210356 | 1 |
| PLEKHA3 | 0.00014958 | 1.14171533 | 1 |
| CYB5RL | 0.00183852 | 1.14168294 | 1 |
| IKBIP | 0.00072354 | 1.13961544 | 1 |
| ACSS3 | 6.54E-05 | 1.13894465 | 1 |
| PPP2R3C | 9.92E-05 | 1.13696417 | 1 |
| RIOX1 | 0.00250483 | 1.13280901 | 1 |
| CCDC141 | 0.03594207 | 1.13160502 | 1 |
| SLC17A9 | 0.00172674 | 1.13142097 | 1 |
| PCGF1 | 0.00011388 | 1.13117508 | 1 |
| EVPL | 0.00602061 | 1.1304866 | 1 |
| PIK3R5 | 0.00026149 | 1.12961364 | 1 |
| TMX3 | 0.0001333 | 1.12923924 | 1 |
| EBPL | 0.00250483 | 1.12828243 | 1 |
| KCNK13 | 0.00323754 | 1.1278361 | 1 |
| RAD18 | 0.01109732 | 1.12686457 | 1 |
| ESRP2 | 0.00099094 | 1.12682232 | 1 |
| NRBF2 | 0.00072285 | 1.12666999 | 1 |
| NTRK3 | 0.00035757 | 1.12580071 | 1 |
| BEGAIN | 0.00125665 | 1.12557767 | 1 |
| NRG2 | 0.02663715 | 1.12142585 | 1 |
| PAX9 | 0.04369348 | 1.12090648 | 1 |
| PTAFR | 0.00283377 | 1.11824847 | 1 |
| ZBTB40 | 0.0001435 | 1.11805011 | 1 |
| ATP7A | 0.00019144 | 1.11550637 | 1 |
| WNK3 | 0.02002038 | 1.11513906 | 1 |
| NLGN1 | 0.04587159 | 1.11483257 | 1 |
| CYP2B6 | 0.01301192 | 1.11273731 | 1 |
| TUBA1A | 0.00125665 | 1.11260223 | 1 |
| SIRPB1 | 0.00601756 | 1.11251018 | 1 |
| AP4M1 | 0.00167 | 1.10972133 | 1 |

| | | | |
|---|---|---|---|
| TMEM150B | 0.03648149 | 1.10958734 | 1 |
| NUGGC | 0.00601756 | 1.10936628 | 1 |
| FDXR | 0.00802629 | 1.10851239 | 1 |
| RECQL5 | 0.00105615 | 1.10762877 | 1 |
| REP15 | 0.01123533 | 1.10745452 | 1 |
| ZNF888 | 0.01092706 | 1.10196871 | 1 |
| ZC3H12D | 0.00758553 | 1.10189512 | 1 |
| TTF1 | 0.00064778 | 1.10149391 | 1 |
| ACKR3 | 0.00029403 | 1.10057669 | 1 |
| SARM1 | 0.00071261 | 1.10038342 | 1 |
| ASGR1 | 0.04845233 | 1.09921239 | 1 |
| IL15RA | 0.00098794 | 1.09908237 | 1 |
| GSKIP | 0.00175057 | 1.09809082 | 1 |
| TSEN34 | 0.00123693 | 1.09595917 | 1 |
| MEAK7 | 0.00101106 | 1.09564714 | 1 |
| ACTA2 | 8.12E-05 | 1.09340976 | 1 |
| TAS2R38 | 0.04369348 | 1.09122489 | 1 |
| GLIPR1 | 0.00046552 | 1.0904738 | 1 |
| VWA3A | 0.02022291 | 1.08935821 | 1 |
| TMEM273 | 0.01995308 | 1.08779765 | 1 |
| MAGI1 | 6.94E-05 | 1.08492205 | 1 |
| SPARCL1 | 0.00031091 | 1.08276103 | 1 |
| IL18R1 | 0.00017599 | 1.08264876 | 1 |
| ZNF669 | 0.00441844 | 1.0806149 | 1 |
| RNASEH2B | 0.00186603 | 1.07846876 | 1 |
| MTMR14 | 0.00051341 | 1.07819617 | 1 |
| FRMD8 | 0.00119087 | 1.0777 | 1 |
| TRIM17 | 0.0023884 | 1.07758428 | 1 |
| ESRP1 | 0.00125665 | 1.07707908 | 1 |
| CHTF8 | 0.00704585 | 1.076848 | 1 |
| TFAP2B | 0.00101919 | 1.07575289 | 1 |
| FAM210A | 0.00017152 | 1.07546226 | 1 |
| CCDC171 | 0.00087171 | 1.07362347 | 1 |
| ZNF420 | 0.0314304 | 1.07296488 | 1 |
| HS1BP3 | 0.01475864 | 1.07204696 | 1 |
| MYH11 | 0.00018744 | 1.07114468 | 1 |
| CAPN13 | 0.00463036 | 1.06920676 | 1 |
| EML5 | 0.01109732 | 1.06868763 | 1 |
| ARG1 | 0.04845233 | 1.06716304 | 1 |
| SLC41A2 | 7.76E-05 | 1.06679214 | 1 |
| PCDHGA12 | 9.44E-05 | 1.06629497 | 1 |
| ZSCAN20 | 0.00253332 | 1.0654613 | 1 |
| PACC1 | 0.00059921 | 1.06476107 | 1 |
| DNAJC12 | 0.0011749 | 1.06436363 | 1 |
| PMAIP1 | 0.0009123 | 1.06350331 | 1 |
| GTF2B | 8.10E-05 | 1.06309225 | 1 |
| RSPO1 | 0.02616585 | 1.06301918 | 1 |
| FAM131C | 0.02745061 | 1.06009441 | 1 |
| CEP152 | 0.0009123 | 1.05993 | 1 |
| PTTG2 | 0.00601756 | 1.05980017 | 1 |
| TFDP1 | 0.00027028 | 1.05972056 | 1 |
| SLC7A1 | 0.00104983 | 1.05847102 | 1 |
| GBP2 | 0.00026083 | 1.05705709 | 1 |

| | | | |
|---|---|---|---|
| COL26A1 | 0.02547989 | 1.05563676 | 1 |
| ARL14EPL | 0.04917111 | 1.054785 | 1 |
| TST | 0.00011975 | 1.05440871 | 1 |
| BMP8B | 0.02032656 | 1.05421851 | 1 |
| ANG | 0.00100597 | 1.05384068 | 1 |
| TBX21 | 0.00817935 | 1.05263539 | 1 |
| ANGPT1 | 6.23E-05 | 1.05099038 | 1 |
| C19orf48 | 0.02621771 | 1.05021683 | 1 |
| GZMB | 0.0342393 | 1.04934646 | 1 |
| GJB1 | 0.00039452 | 1.04885176 | 1 |
| SAYSD1 | 7.41E-05 | 1.0468096 | 1 |
| TRIM34 | 0.00246553 | 1.04588205 | 1 |
| RBM20 | 0.0148021 | 1.04354392 | 1 |
| ASB9 | 0.000457 | 1.04315466 | 1 |
| USP42 | 0.00118859 | 1.04196786 | 1 |
| ZBTB43 | 0.00066064 | 1.04130387 | 1 |
| TBXAS1 | 0.02032656 | 1.04000869 | 1 |
| EXOSC5 | 0.00250483 | 1.03988524 | 1 |
| GNGT2 | 0.00454701 | 1.03941821 | 1 |
| GJD3 | 0.00184845 | 1.03846839 | 1 |
| RITA1 | 0.00102888 | 1.03776993 | 1 |
| LYRM1 | 6.31E-05 | 1.03641118 | 1 |
| EDNRA | 0.00014204 | 1.03631082 | 1 |
| PTPRU | 0.00172674 | 1.0362049 | 1 |
| DPY19L1 | 0.00024109 | 1.0360525 | 1 |
| SELL | 0.00426735 | 1.0350836 | 1 |
| MID1 | 0.00021643 | 1.03506756 | 1 |
| RASD1 | 0.00088778 | 1.0340953 | 1 |
| DSEL | 0.00021102 | 1.03392505 | 1 |
| GFER | 0.00012128 | 1.03211726 | 1 |
| CDH6 | 0.00023889 | 1.03006969 | 1 |
| CD72 | 0.00038076 | 1.02996326 | 1 |
| SIGLEC10 | 0.00601756 | 1.02946013 | 1 |
| GPR26 | 0.04369348 | 1.02908483 | 1 |
| CTDSPL2 | 7.11E-05 | 1.02782808 | 1 |
| UBE2C | 0.02221215 | 1.02718875 | 1 |
| FGFRL1 | 0.00117633 | 1.02660803 | 1 |
| RPUSD3 | 0.00094851 | 1.02623444 | 1 |
| EFCAB10 | 0.04917111 | 1.02597277 | 1 |
| ANKRD24 | 0.02663715 | 1.0249889 | 1 |
| TTC38 | 6.85E-05 | 1.02169025 | 1 |
| RHPN2 | 0.00054325 | 1.02149808 | 1 |
| TM6SF1 | 0.01317796 | 1.02116874 | 1 |
| PDE3B | 0.0019222 | 1.0201787 | 1 |
| SHLD2 | 7.13E-05 | 1.02011384 | 1 |
| ZNF202 | 0.00038754 | 1.02003278 | 1 |
| FSTL3 | 0.00175404 | 1.01986168 | 1 |
| NR2C2AP | 0.00323754 | 1.01681768 | 1 |
| FOLH1 | 0.04576652 | 1.01666368 | 1 |
| POLR3K | 0.00040506 | 1.01549355 | 1 |
| PSD4 | 0.00052059 | 1.01289081 | 1 |
| IGKC | 0.0020217 | 1.0119312 | 1 |
| ZMYND12 | 0.01105788 | 1.01100726 | 1 |

| | | | |
|---|---|---|---|
| ENTPD2 | 0.01109732 | 1.00962926 | 1 |
| EEFSEC | 0.00119533 | 1.00629258 | 1 |
| B3GALT6 | 7.97E-05 | 1.00378452 | 1 |
| KRT19 | 0.00013744 | 1.00289891 | 1 |
| TLCD5 | 0.00236709 | 1.0028142 | 1 |
| ATP8A2 | 0.02332712 | 1.00243367 | 1 |
| BDH1 | 0.00017885 | 1.00188017 | 1 |
| RIIAD1 | 0.03648149 | 1.00186239 | 1 |
| RGS1 | 0.04917111 | 1.00147603 | 1 |
| TNFRSF10A | 0.01192848 | 1.00142313 | 1 |
| KLHL6 | 0.00941941 | 1.00094276 | 1 |
| TRMT13 | 9.21E-05 | 1.00056821 | 1 |
| RIPK4 | 0.00644706 | 0.99932342 | 1 |
| ZNF614 | 0.00599547 | 0.99883043 | 1 |
| RXFP4 | 0.04917111 | 0.99762513 | 1 |
| ADRB2 | 0.00338015 | 0.99570579 | 1 |
| PINLYP | 0.00286147 | 0.99550393 | 1 |
| TOM1 | 0.00013732 | 0.99478022 | 1 |
| AQP9 | 0.00143361 | 0.99448731 | 1 |
| HCN3 | 0.03615466 | 0.99380744 | 1 |
| SIGLEC12 | 0.02002038 | 0.99370958 | 1 |
| KIF27 | 0.01733605 | 0.99362848 | 1 |
| RNF4 | 0.00053788 | 0.99350873 | 1 |
| EIF3I | 0.00448664 | 0.99349694 | 1 |
| ZNF576 | 0.00015761 | 0.99303472 | 1 |
| CDIP1 | 0.00013791 | 0.99251026 | 1 |
| GALNT1 | 6.71E-05 | 0.99247485 | 1 |
| TOMM22 | 0.00019173 | 0.99111076 | 1 |
| RMND1 | 0.00034808 | 0.9886998 | 1 |
| TMEM132C | 0.03836235 | 0.9871286 | 1 |
| LINC02693 | 0.00742781 | 0.98660684 | 1 |
| AZGP1 | 8.36E-05 | 0.98631768 | 1 |
| HOXC5 | 0.0093112 | 0.98566046 | 1 |
| KIF17 | 0.00825516 | 0.9842983 | 1 |
| ATG4A | 0.00398241 | 0.98379803 | 1 |
| ESRRB | 0.00326774 | 0.98322867 | 1 |
| DTWD2 | 0.01334587 | 0.98179791 | 1 |
| NR2E3 | 0.04845233 | 0.9801206 | 1 |
| ELOVL7 | 0.00040388 | 0.97966138 | 1 |
| STAC2 | 0.00817935 | 0.97941387 | 1 |
| ITGA4 | 7.51E-05 | 0.97926598 | 1 |
| RWDD2B | 0.00312842 | 0.97921473 | 1 |
| B4GALNT3 | 0.04357182 | 0.97912469 | 1 |
| SACS | 0.00089715 | 0.97902374 | 1 |
| SLC35G1 | 0.01105788 | 0.97834942 | 1 |
| CCDC163 | 0.02745061 | 0.97801291 | 1 |
| RFC4 | 0.01995308 | 0.97769758 | 1 |
| APOA1 | 0.01916236 | 0.97714086 | 1 |
| UBN2 | 0.00010256 | 0.9764037 | 1 |
| MOCS3 | 0.00096149 | 0.97611416 | 1 |
| ZNF438 | 0.00041538 | 0.97472859 | 1 |
| ZRSR2 | 0.01251002 | 0.97446282 | 1 |
| TEX30 | 0.02651478 | 0.97427751 | 1 |

| | | | |
|---|---|---|---|
| IQCB1 | 0.00047669 | 0.97373067 | 1 |
| NECAP1 | 5.74E-05 | 0.97338222 | 1 |
| ABCG2 | 0.0258209 | 0.97338217 | 1 |
| ARMC12 | 0.03648149 | 0.9729573 | 1 |
| LACC1 | 0.00023088 | 0.97251936 | 1 |
| VPS16 | 0.00014387 | 0.97231355 | 1 |
| POLR3C | 6.85E-05 | 0.97176065 | 1 |
| C10orf95 | 0.03702779 | 0.97175761 | 1 |
| ACD | 0.00150175 | 0.97070681 | 1 |
| DDR2 | 0.00022127 | 0.97013614 | 1 |
| MSANTD3 | 0.00040388 | 0.97006854 | 1 |
| DGKA | 0.00013536 | 0.96965983 | 1 |
| NRCAM | 0.03291963 | 0.96929033 | 1 |
| CEP350 | 0.00031509 | 0.96719466 | 1 |
| KLHL13 | 0.01475864 | 0.96405952 | 1 |
| ARHGEF35 | 0.00012477 | 0.96340358 | 1 |
| SRPX2 | 0.00019197 | 0.96338009 | 1 |
| DHX33 | 0.00017963 | 0.9623277 | 1 |
| HOXB9 | 0.0280311 | 0.96073471 | 1 |
| PARD6A | 0.03974119 | 0.95977801 | 1 |
| CES3 | 0.00091636 | 0.95901293 | 1 |
| SLC25A29 | 0.00050657 | 0.95881641 | 1 |
| TMEM242 | 7.67E-05 | 0.95805625 | 1 |
| UNC5D | 0.00033309 | 0.95771484 | 1 |
| LGALSL | 8.79E-05 | 0.9559931 | 1 |
| ADRA2B | 0.00041168 | 0.95443949 | 1 |
| ZNF236 | 0.00011826 | 0.95436466 | 1 |
| CAV1 | 0.00027126 | 0.9537054 | 1 |
| TNFRSF11A | 0.01023885 | 0.95330295 | 1 |
| VPS9D1 | 5.81E-05 | 0.95177394 | 1 |
| PICK1 | 0.00020821 | 0.95020668 | 1 |
| C15orf48 | 0.00805279 | 0.94818099 | 1 |
| AKR1C3 | 0.00086663 | 0.94686785 | 1 |
| OSCAR | 0.00449906 | 0.94665331 | 1 |
| FAM20A | 0.02985804 | 0.94436345 | 1 |
| TSSK4 | 0.00323754 | 0.9429747 | 1 |
| MTFMT | 0.00787452 | 0.94213194 | 1 |
| RALGPS2 | 0.00442701 | 0.94089535 | 1 |
| VSTM4 | 0.00020455 | 0.93922068 | 1 |
| EPB41L4B | 0.00588943 | 0.93872249 | 1 |
| P2RX1 | 0.01503064 | 0.93795631 | 1 |
| RANBP6 | 0.00143331 | 0.93790125 | 1 |
| TMEM88B | 0.00260876 | 0.93755115 | 1 |
| PARP11 | 0.00535527 | 0.93654424 | 1 |
| UBE2E1 | 6.89E-05 | 0.93566392 | 1 |
| C6orf132 | 0.01497654 | 0.93542717 | 1 |
| ALG9 | 0.00019431 | 0.93362908 | 1 |
| ACACA | 0.00192265 | 0.93232227 | 1 |
| DDX20 | 0.01535855 | 0.93227489 | 1 |
| ROGDI | 0.0001627 | 0.93181933 | 1 |
| DMBT1 | 0.00323754 | 0.9313161 | 1 |
| ROR1 | 0.00108005 | 0.93089407 | 1 |
| PUS7L | 0.00101718 | 0.93059377 | 1 |

| | | | |
|---|---|---|---|
| SLC6A1 | 0.04413743 | 0.93031225 | 1 |
| C19orf54 | 0.00426735 | 0.92962847 | 1 |
| DAO | 0.00031558 | 0.92741704 | 1 |
| SERPINB8 | 0.00051341 | 0.92661117 | 1 |
| PDLIM1 | 0.00022352 | 0.92650669 | 1 |
| SLC13A2 | 0.00017177 | 0.9264622 | 1 |
| FYB1 | 0.00323553 | 0.92573956 | 1 |
| SHROOM1 | 0.00314213 | 0.92522556 | 1 |
| ATXN7L2 | 0.01136809 | 0.92497541 | 1 |
| APOC1 | 0.0175649 | 0.92464738 | 1 |
| NEB | 0.04917111 | 0.92428858 | 1 |
| CD274 | 0.00148122 | 0.92363159 | 1 |
| NDUFA6 | 0.00519312 | 0.92242494 | 1 |
| TACR1 | 0.00817935 | 0.92194258 | 1 |
| FLAD1 | 0.00486304 | 0.9203899 | 1 |
| MX2 | 6.07E-05 | 0.91990994 | 1 |
| SLC25A48 | 0.00074442 | 0.91968032 | 1 |
| SMIM14 | 0.00013045 | 0.91967782 | 1 |
| PLSCR1 | 8.61E-05 | 0.9191277 | 1 |
| NLRP6 | 0.03702779 | 0.91851762 | 1 |
| DNAH7 | 0.00787946 | 0.9176255 | 1 |
| NUMB | 0.00014326 | 0.91706845 | 1 |
| ZKSCAN5 | 0.00421372 | 0.91688551 | 1 |
| MRC1 | 0.0001028 | 0.91633353 | 1 |
| NGEF | 0.030393 | 0.91623118 | 1 |
| FKBP3 | 0.00290428 | 0.91608898 | 1 |
| UFL1 | 0.00029285 | 0.91582727 | 1 |
| PRKCB | 0.01779674 | 0.91419799 | 1 |
| TRERF1 | 0.0015185 | 0.91271619 | 1 |
| QRFPR | 0.02745061 | 0.91160887 | 1 |
| GYPC | 7.54E-05 | 0.91133349 | 1 |
| FAM214B | 0.0003998 | 0.91096327 | 1 |
| WDR53 | 0.00080724 | 0.91092455 | 1 |
| TRAF1 | 0.00154277 | 0.91050535 | 1 |
| MRM2 | 0.0019771 | 0.91018601 | 1 |
| ENO2 | 0.01330446 | 0.91001893 | 1 |
| UCP3 | 0.00621715 | 0.90956233 | 1 |
| GDF7 | 0.00097767 | 0.90956067 | 1 |
| SPHK1 | 0.00055055 | 0.90889459 | 1 |
| LRMP | 0.00102888 | 0.90694074 | 1 |
| DENND1B | 0.00042203 | 0.90604654 | 1 |
| PAK1IP1 | 0.02412932 | 0.90566956 | 1 |
| BX255925.3 | 0.00115828 | 0.90538479 | 1 |
| C2CD3 | 0.00081892 | 0.90512282 | 1 |
| ADAM33 | 0.00347981 | 0.90353258 | 1 |
| SLC4A4 | 7.87E-05 | 0.9032118 | 1 |
| OR2T10 | 0.00763188 | 0.90218461 | 1 |
| FGD3 | 0.00078551 | 0.90215013 | 1 |
| SYT1 | 0.03702779 | 0.90212174 | 1 |
| NDOR1 | 0.00151331 | 0.9018348 | 1 |
| NDUFA5 | 0.00078997 | 0.9016162 | 1 |
| TNFAIP8L2 | 0.00074442 | 0.90125725 | 1 |
| KCNH6 | 0.00183701 | 0.90068069 | 1 |

| | | | |
|---|---|---|---|
| GPT | 0.00299754 | 0.8991623 | 1 |
| CLEC5A | 0.00323754 | 0.89891568 | 1 |
| AFM | 7.40E-05 | 0.89880163 | 1 |
| VAMP4 | 0.00315924 | 0.89748998 | 1 |
| RNASEH1 | 0.00191207 | 0.8964116 | 1 |
| SPX | 0.00237149 | 0.8961311 | 1 |
| FANCI | 0.01092706 | 0.89594348 | 1 |
| ACP1 | 0.00029462 | 0.8954803 | 1 |
| TCF19 | 0.00125665 | 0.89514974 | 1 |
| VTN | 0.00476511 | 0.89514045 | 1 |
| CAPZA1 | 0.03468912 | 0.89464636 | 1 |
| PIH1D2 | 0.00186603 | 0.89387736 | 1 |
| MZF1 | 0.01037103 | 0.89373265 | 1 |
| DUSP18 | 0.00137063 | 0.88912924 | 1 |
| DACT3 | 0.01970305 | 0.88724995 | 1 |
| C11orf74 | 0.00036931 | 0.8871068 | 1 |
| OPA3 | 0.00077829 | 0.88694479 | 1 |
| POLA1 | 0.00058257 | 0.88602818 | 1 |
| SRPX | 0.01009255 | 0.88575623 | 1 |
| AKAP6 | 0.00293284 | 0.88480089 | 1 |
| COL13A1 | 0.00848985 | 0.883586 | 1 |
| ADSSL1 | 8.67E-05 | 0.88068731 | 1 |
| KLHL26 | 0.03123586 | 0.88066236 | 1 |
| RORC | 0.00242775 | 0.88041882 | 1 |
| SGTB | 0.00216565 | 0.88034843 | 1 |
| VIPR1 | 0.00012007 | 0.88024181 | 1 |
| EP400 | 0.0014403 | 0.87829551 | 1 |
| C11orf45 | 0.02745061 | 0.87647996 | 1 |
| GFOD1 | 0.00011103 | 0.87612871 | 1 |
| SLC25A20 | 0.00978621 | 0.87273392 | 1 |
| NUP155 | 0.00013363 | 0.87146444 | 1 |
| HAUS1 | 0.01368738 | 0.87133037 | 1 |
| WNT7B | 0.04917111 | 0.8707918 | 1 |
| HEATR5A | 0.00549079 | 0.86980623 | 1 |
| DHX58 | 0.00030868 | 0.8689291 | 1 |
| AMD1 | 0.00284258 | 0.8672644 | 1 |
| PHF8 | 0.00012354 | 0.8671293 | 1 |
| GSTA2 | 0.00091204 | 0.86664612 | 1 |
| NUDT7 | 0.00014608 | 0.86650557 | 1 |
| RNF2 | 0.00311843 | 0.86628956 | 1 |
| ARHGEF33 | 0.00597404 | 0.8660078 | 1 |
| CBLN1 | 0.04989882 | 0.86577395 | 1 |
| NIF3L1 | 0.0003865 | 0.86561535 | 1 |
| INTS7 | 0.00063696 | 0.86527656 | 1 |
| ELOA2 | 0.04917111 | 0.86494959 | 1 |
| PRDM1 | 0.00400816 | 0.86490644 | 1 |
| C18orf21 | 0.0128101 | 0.86488742 | 1 |
| ZNF595 | 0.00037538 | 0.86484345 | 1 |
| KY | 0.02686771 | 0.86483354 | 1 |
| ITGA10 | 0.00173733 | 0.86420129 | 1 |
| C11orf21 | 0.02032656 | 0.86236622 | 1 |
| COQ8A | 0.00012307 | 0.86014912 | 1 |
| BCL2L11 | 0.00064183 | 0.85987151 | 1 |

| | | | |
|---|---|---|---|
| MUC15 | 0.01109732 | 0.85740332 | 1 |
| HELLS | 0.00040061 | 0.85737749 | 1 |
| ZNF776 | 0.00016896 | 0.85672085 | 1 |
| CNPPD1 | 0.00042422 | 0.85560197 | 1 |
| CLEC1A | 0.00395954 | 0.85477793 | 1 |
| PRR15L | 0.00080724 | 0.85470388 | 1 |
| ENPP2 | 8.37E-05 | 0.85420379 | 1 |
| CELF2 | 0.00036248 | 0.8534964 | 1 |
| INSM1 | 0.02032656 | 0.85348747 | 1 |
| GPR35 | 0.00323754 | 0.85318006 | 1 |
| ALDH8A1 | 0.00131398 | 0.85308335 | 1 |
| PBX3 | 0.00021238 | 0.85302344 | 1 |
| SYNC | 0.04021005 | 0.85222268 | 1 |
| TRMT12 | 0.00331241 | 0.85204391 | 1 |
| NF1 | 7.89E-05 | 0.85179672 | 1 |
| SMIM6 | 0.00323754 | 0.85163909 | 1 |
| MBIP | 0.00042832 | 0.85016953 | 1 |
| TMEM14B | 0.00026879 | 0.84968139 | 1 |
| ANKRD6 | 0.00642018 | 0.84860606 | 1 |
| ADAMTS6 | 0.02636977 | 0.84829472 | 1 |
| TRMT10A | 0.0013237 | 0.84800269 | 1 |
| IL17RB | 0.00012255 | 0.84782288 | 1 |
| NUBPL | 0.00036053 | 0.84763826 | 1 |
| TSPAN11 | 0.03648149 | 0.84735187 | 1 |
| ELF1 | 5.83E-05 | 0.84727138 | 1 |
| STAC3 | 0.02002038 | 0.84719181 | 1 |
| AADAT | 0.02871767 | 0.8467577 | 1 |
| AURKB | 0.03702779 | 0.84555913 | 1 |
| ZNF627 | 0.00092304 | 0.84509926 | 1 |
| SLC26A1 | 0.00098794 | 0.84501001 | 1 |
| MT1H | 0.0003158 | 0.84481913 | 1 |
| CBLL1 | 0.00037946 | 0.84449501 | 1 |
| BCCIP | 0.00115403 | 0.84437033 | 1 |
| RAB3A | 0.03648149 | 0.84374185 | 1 |
| EFCAB2 | 0.00065757 | 0.84323547 | 1 |
| RHBDD1 | 0.00152781 | 0.84178025 | 1 |
| FOXRED2 | 0.00441844 | 0.84107327 | 1 |
| LYG1 | 0.0148021 | 0.83985022 | 1 |
| PNPLA8 | 0.0001757 | 0.83922403 | 1 |
| IFI27L1 | 0.02305375 | 0.83856121 | 1 |
| C1orf35 | 0.00017373 | 0.83853165 | 1 |
| PTCH1 | 0.00133426 | 0.83844951 | 1 |
| TUFT1 | 0.00059127 | 0.83713016 | 1 |
| ZNF57 | 0.02616585 | 0.83647531 | 1 |
| MFSD8 | 0.00366291 | 0.83601565 | 1 |
| TTC5 | 0.00017699 | 0.83574662 | 1 |
| SMIM8 | 0.00175382 | 0.83465539 | 1 |
| RNF215 | 0.00041001 | 0.83419032 | 1 |
| SLC35F4 | 0.04989882 | 0.83414111 | 1 |
| COPRS | 0.00034144 | 0.8341401 | 1 |
| JAK3 | 8.14E-05 | 0.83178677 | 1 |
| BCKDHB | 0.00277588 | 0.83144425 | 1 |
| TRMT11 | 0.00479609 | 0.83137976 | 1 |

| | | | |
|---|---|---|---|
| CYB561D2 | 0.00012394 | 0.83130384 | 1 |
| CYP2E1 | 0.01207157 | 0.83107156 | 1 |
| COMMD10 | 0.00012493 | 0.83083872 | 1 |
| GCAT | 0.00825083 | 0.83061233 | 1 |
| FADS1 | 0.00343489 | 0.83054919 | 1 |
| SYPL1 | 0.00031196 | 0.83046727 | 1 |
| PLIN2 | 6.26E-05 | 0.82800037 | 1 |
| KLF12 | 0.00016998 | 0.82773178 | 1 |
| ETAA1 | 5.66E-05 | 0.82757412 | 1 |
| TEX9 | 0.00092383 | 0.82753661 | 1 |
| SUPT3H | 0.01611641 | 0.82751654 | 1 |
| ZFAT | 0.00343775 | 0.82749148 | 1 |
| CAMSAP3 | 0.00536823 | 0.82673295 | 1 |
| FLVCR2 | 0.00400528 | 0.82641985 | 1 |
| SCLT1 | 8.23E-05 | 0.82598604 | 1 |
| DONSON | 0.0157187 | 0.82560902 | 1 |
| GRWD1 | 0.00410697 | 0.82513896 | 1 |
| SCIMP | 0.00097767 | 0.82509249 | 1 |
| TERC | 0.01031501 | 0.82499064 | 1 |
| GM2A | 0.00116445 | 0.82483086 | 1 |
| ZC3H12C | 0.00160401 | 0.82369113 | 1 |
| TARSL2 | 0.00042815 | 0.82278052 | 1 |
| HASPIN | 0.03560392 | 0.82270157 | 1 |
| KIAA0513 | 0.00287929 | 0.821141 | 1 |
| TMEM134 | 6.20E-05 | 0.82113664 | 1 |
| CERS5 | 0.00039682 | 0.82093734 | 1 |
| ALDH1L1 | 6.03E-05 | 0.82081721 | 1 |
| PEPD | 0.00043453 | 0.82072152 | 1 |
| UBTD2 | 0.00051297 | 0.82058981 | 1 |
| C5orf30 | 0.00577114 | 0.81852255 | 1 |
| ISG20 | 6.94E-05 | 0.81812682 | 1 |
| ALAS1 | 0.00053176 | 0.81808102 | 1 |
| STK11IP | 0.00923842 | 0.81757672 | 1 |
| MMRN1 | 0.0046973 | 0.81732115 | 1 |
| RIPK1 | 0.0001311 | 0.81607937 | 1 |
| RP9 | 0.00112869 | 0.81595748 | 1 |
| SUCLG1 | 6.15E-05 | 0.81541338 | 1 |
| GPN1 | 0.00016044 | 0.81524133 | 1 |
| CFD | 0.00098794 | 0.8151665 | 1 |
| ZNF501 | 0.00103472 | 0.81415209 | 1 |
| PEX14 | 0.00469116 | 0.81381915 | 1 |
| ABCD3 | 0.00013897 | 0.81258069 | 1 |
| ROCK1 | 8.36E-05 | 0.81208506 | 1 |
| IER5L | 0.00041753 | 0.81158511 | 1 |
| CCL14 | 0.00178653 | 0.81138202 | 1 |
| ZBTB12 | 0.04600568 | 0.81104263 | 1 |
| CAPN5 | 0.00407443 | 0.81052505 | 1 |
| UBAC2 | 0.00024702 | 0.80907803 | 1 |
| LY86 | 0.0180068 | 0.80853483 | 1 |
| CNOT3 | 0.00012403 | 0.80843424 | 1 |
| EZHIP | 0.02032656 | 0.80830943 | 1 |
| AGMAT | 0.00028627 | 0.8068478 | 1 |
| POM121L12 | 0.04989882 | 0.80534977 | 1 |

| | | | |
|---|---|---|---|
| GPATCH8 | 6.08E-05 | 0.80507406 | 1 |
| RNFT1 | 0.00022173 | 0.80500928 | 1 |
| C16orf54 | 0.03514398 | 0.80485892 | 1 |
| TNIP2 | 0.00024804 | 0.80484462 | 1 |
| SOCS1 | 0.0011817 | 0.80461881 | 1 |
| EBI3 | 0.0196864 | 0.80365939 | 1 |
| HIST1H1D | 0.00052171 | 0.80351122 | 1 |
| ZNF326 | 9.47E-05 | 0.80343096 | 1 |
| SUGCT | 8.90E-05 | 0.80262402 | 1 |
| RARB | 0.00025531 | 0.80191629 | 1 |
| CHD8 | 9.84E-05 | 0.80133076 | 1 |
| AASDHPPT | 0.00015434 | 0.80028089 | 1 |
| DZIP1L | 0.00136452 | 0.80011741 | 1 |
| INAFM2 | 0.00319277 | 0.79953559 | 1 |
| GRIPAP1 | 0.00015167 | 0.79945293 | 1 |
| HECW2 | 0.00015318 | 0.79815035 | 1 |
| SLC2A10 | 0.00627065 | 0.79801393 | 1 |
| IRF5 | 0.00901575 | 0.79757373 | 1 |
| SELPLG | 0.01853018 | 0.79639092 | 1 |
| BTN3A2 | 0.0004984 | 0.79616696 | 1 |
| GMPR | 0.00359232 | 0.79612783 | 1 |
| TEFM | 6.67E-05 | 0.79542703 | 1 |
| MMP7 | 0.00070023 | 0.79445603 | 1 |
| GRB14 | 0.00224561 | 0.79431001 | 1 |
| GOLGA5 | 0.00052366 | 0.79424689 | 1 |
| ZNF423 | 0.00011476 | 0.79343678 | 1 |
| RAD1 | 0.00026879 | 0.79263403 | 1 |
| GPRASP1 | 0.0001064 | 0.79260183 | 1 |
| XRCC5 | 0.0008074 | 0.79250205 | 1 |
| KATNBL1 | 0.04917111 | 0.79249312 | 1 |
| FCGR2A | 5.55E-05 | 0.79238877 | 1 |
| OAS3 | 0.00103784 | 0.79217087 | 1 |
| CPTP | 0.00073153 | 0.79186436 | 1 |
| ICK | 0.01218479 | 0.79102596 | 1 |
| ARHGAP1 | 0.00016538 | 0.79073981 | 1 |
| PPAN | 7.65E-05 | 0.79027694 | 1 |
| SLC5A10 | 0.00049061 | 0.79013636 | 1 |
| H2AFZ | 0.00011586 | 0.7900636 | 1 |
| MET | 0.00022412 | 0.78941344 | 1 |
| IBA57 | 0.00080493 | 0.78825452 | 1 |
| KYAT1 | 0.00053011 | 0.78734937 | 1 |
| LRRC19 | 0.00012836 | 0.78697362 | 1 |
| CETN3 | 5.90E-05 | 0.78612187 | 1 |
| GTF3C5 | 0.00012628 | 0.78513812 | 1 |
| GTF2E1 | 0.00138087 | 0.78467519 | 1 |
| ERC2 | 0.02304392 | 0.78411222 | 1 |
| LNP1 | 0.0258209 | 0.78410656 | 1 |
| SETBP1 | 0.00031127 | 0.78391991 | 1 |
| UPRT | 0.00038374 | 0.78343597 | 1 |
| PGM1 | 0.00032991 | 0.78290777 | 1 |
| PLA2G15 | 0.00019363 | 0.78187472 | 1 |
| CNOT1 | 0.00019924 | 0.78061116 | 1 |
| MRI1 | 7.91E-05 | 0.77890451 | 1 |

| | | | |
|---|---|---|---|
| PYCARD | 7.90E-05 | 0.77865941 | 1 |
| SP2 | 0.00014887 | 0.77778506 | 1 |
| RBBP9 | 0.00012413 | 0.77719308 | 1 |
| STOML1 | 0.00430988 | 0.77691076 | 1 |
| FLI1 | 0.00519866 | 0.77618234 | 1 |
| KLK13 | 0.03468912 | 0.77587936 | 1 |
| TOP3B | 0.00042177 | 0.77573836 | 1 |
| NME6 | 0.00044905 | 0.77552065 | 1 |
| CNN1 | 0.00074442 | 0.77498937 | 1 |
| MOSMO | 0.0019222 | 0.77449845 | 1 |
| FAH | 0.00076925 | 0.77442669 | 1 |
| SLC36A2 | 0.00051231 | 0.77430859 | 1 |
| ZNF689 | 0.000632 | 0.77385353 | 1 |
| RBM18 | 0.0002482 | 0.77384204 | 1 |
| UBQLNL | 0.00323754 | 0.7728911 | 1 |
| ZNF212 | 0.00019806 | 0.77197938 | 1 |
| TWF2 | 0.00028575 | 0.77195817 | 1 |
| PDZK1IP1 | 0.00147504 | 0.77173018 | 1 |
| SIGLEC7 | 0.04917111 | 0.77127427 | 1 |
| C6orf136 | 0.00084297 | 0.77061153 | 1 |
| MPZ | 0.03560392 | 0.77045203 | 1 |
| SHKBP1 | 0.00024075 | 0.76883355 | 1 |
| PDE8B | 0.0016154 | 0.76879588 | 1 |
| TRNP1 | 0.00118859 | 0.76875883 | 1 |
| ATG7 | 0.00016229 | 0.76868139 | 1 |
| CIAO3 | 7.64E-05 | 0.76736063 | 1 |
| AQP2 | 0.00030787 | 0.76728721 | 1 |
| AL590560.2 | 0.00022173 | 0.76631503 | 1 |
| C19orf38 | 0.00817935 | 0.76614022 | 1 |
| TOP1MT | 0.00064901 | 0.76594718 | 1 |
| CPOX | 0.00024728 | 0.764477 | 1 |
| OR2K2 | 0.04989882 | 0.76371862 | 1 |
| RASA3 | 0.00018477 | 0.76347978 | 1 |
| XPR1 | 8.47E-05 | 0.76329445 | 1 |
| EIF3H | 7.90E-05 | 0.76242737 | 1 |
| RBM48 | 0.00749198 | 0.76152701 | 1 |
| FBLN5 | 0.00051371 | 0.7611878 | 1 |
| CFI | 0.00185655 | 0.76117177 | 1 |
| MX1 | 0.00306342 | 0.76063117 | 1 |
| TNXB | 0.00853836 | 0.76017425 | 1 |
| MYCBP2 | 0.00010714 | 0.75974225 | 1 |
| CMTR2 | 0.00232649 | 0.75853872 | 1 |
| SLC25A17 | 0.00018733 | 0.75848775 | 1 |
| EML2 | 0.00056102 | 0.75767203 | 1 |
| DAGLA | 0.02884145 | 0.7572917 | 1 |
| TAS2R30 | 0.04470923 | 0.75703192 | 1 |
| ATF5 | 0.00032312 | 0.75686974 | 1 |
| FST | 0.03702779 | 0.75684912 | 1 |
| ENPP7 | 0.0342393 | 0.75644978 | 1 |
| WIPF1 | 6.13E-05 | 0.75644342 | 1 |
| GGT6 | 0.00037418 | 0.75561457 | 1 |
| AMN1 | 0.00064045 | 0.75515472 | 1 |
| ASTL | 0.00817935 | 0.75503973 | 1 |

| | | | |
|---|---|---|---|
| ATP23 | 0.00121496 | 0.75430518 | 1 |
| CYP2J2 | 0.0148021 | 0.75420156 | 1 |
| IL27RA | 7.59E-05 | 0.75391474 | 1 |
| HSPG2 | 0.00016803 | 0.75390364 | 1 |
| UFD1 | 0.0003221 | 0.75382549 | 1 |
| ETV6 | 0.0001489 | 0.75380954 | 1 |
| GRHL2 | 0.01865056 | 0.75347441 | 1 |
| DYNC1H1 | 0.00020915 | 0.753066 | 1 |
| ARSK | 0.01497654 | 0.75305598 | 1 |
| CACNA2D1 | 0.00022424 | 0.75257721 | 1 |
| BRCA1 | 0.00817935 | 0.75247731 | 1 |
| STRN3 | 0.00015786 | 0.75181037 | 1 |
| COX20 | 9.89E-05 | 0.75157039 | 1 |
| UBE2W | 0.00058094 | 0.75150125 | 1 |
| P2RY13 | 0.01334587 | 0.75054301 | 1 |
| RHEX | 0.02388406 | 0.75053155 | 1 |
| FBXL15 | 0.00039311 | 0.75016392 | 1 |
| GRK6 | 0.00116168 | 0.7497206 | 1 |
| PLD2 | 0.00027395 | 0.74934045 | 1 |
| TERF2 | 0.00038176 | 0.74866348 | 1 |
| EFNA4 | 0.03702779 | 0.7484958 | 1 |
| ARL15 | 8.39E-05 | 0.74824168 | 1 |
| CYGB | 0.00634938 | 0.74794945 | 1 |
| YY1AP1 | 8.78E-05 | 0.74758657 | 1 |
| C14orf28 | 0.00469116 | 0.74755327 | 1 |
| MARK4 | 0.0011011 | 0.74719265 | 1 |
| PROZ | 0.01246409 | 0.74604219 | 1 |
| NUP35 | 0.00236709 | 0.74579982 | 1 |
| PCDHGA7 | 0.00073354 | 0.74566178 | 1 |
| SEPTIN1 | 0.01647013 | 0.7454151 | 1 |
| MMP17 | 0.03702779 | 0.74500051 | 1 |
| INIP | 0.00347981 | 0.74470212 | 1 |
| GNRH1 | 0.00075587 | 0.74444367 | 1 |
| UPF2 | 0.00047959 | 0.74387769 | 1 |
| ADRB1 | 0.01916236 | 0.74332313 | 1 |
| COG5 | 0.00011788 | 0.74330201 | 1 |
| COL6A3 | 0.00072378 | 0.74307235 | 1 |
| ATF3 | 0.00060974 | 0.743004 | 1 |
| TMEM161B | 0.00322764 | 0.741839 | 1 |
| PARP12 | 0.00016518 | 0.74133398 | 1 |
| CSNK2A1 | 0.00036763 | 0.74118704 | 1 |
| FBXL8 | 0.00246553 | 0.74101162 | 1 |
| TMEM69 | 0.00120704 | 0.74068668 | 1 |
| TENM3 | 0.01535443 | 0.74025335 | 1 |
| NLRC3 | 0.03049774 | 0.74011247 | 1 |
| ZNF684 | 0.00016665 | 0.73999949 | 1 |
| MAPK8IP1 | 0.00508831 | 0.73989224 | 1 |
| PRPS2 | 0.00017551 | 0.73985503 | 1 |
| SETD5 | 0.00022316 | 0.73915494 | 1 |
| PCK2 | 0.0015813 | 0.73900279 | 1 |
| SLC35A2 | 0.02361334 | 0.73889168 | 1 |
| HPCAL1 | 9.39E-05 | 0.73805389 | 1 |
| KHDRBS3 | 0.00028099 | 0.7375561 | 1 |

| | | | |
|---|---|---|---|
| KALRN | 0.00067429 | 0.73625538 | 1 |
| SH2D4A | 0.00629931 | 0.73502762 | 1 |
| VPS26A | 0.00067344 | 0.7346345 | 1 |
| SKAP1 | 0.00981363 | 0.73458696 | 1 |
| UVRAG | 0.00013678 | 0.73361362 | 1 |
| THOC2 | 0.00065224 | 0.73334595 | 1 |
| IFI44 | 0.00837312 | 0.73222861 | 1 |
| ZNF687 | 0.00032008 | 0.73180344 | 1 |
| GBP1 | 0.00054981 | 0.73115355 | 1 |
| ZC3H18 | 0.00021716 | 0.73082548 | 1 |
| TBC1D31 | 0.01109732 | 0.73044232 | 1 |
| TP53I13 | 0.00250483 | 0.72947676 | 1 |
| TAF1 | 0.00010991 | 0.72922627 | 1 |
| SLC26A5 | 0.03702779 | 0.72895565 | 1 |
| DDHD1 | 0.00971248 | 0.72876178 | 1 |
| MYH3 | 0.04917111 | 0.72875618 | 1 |
| PGF | 0.00035237 | 0.72819253 | 1 |
| SVBP | 0.00048688 | 0.72770262 | 1 |
| CCL2 | 0.00360153 | 0.72740902 | 1 |
| CDR2L | 0.00078869 | 0.72641308 | 1 |
| HIRIP3 | 7.98E-05 | 0.72623108 | 1 |
| RNF149 | 0.00015973 | 0.72453542 | 1 |
| C5AR2 | 0.00483381 | 0.72418489 | 1 |
| OTUD4 | 0.00104343 | 0.7240253 | 1 |
| CBFA2T3 | 0.00463036 | 0.72381151 | 1 |
| LRRC32 | 0.00011557 | 0.72360715 | 1 |
| SLC25A28 | 7.54E-05 | 0.72349638 | 1 |
| DHX30 | 8.29E-05 | 0.72342347 | 1 |
| CCL15 | 0.00442701 | 0.72334373 | 1 |
| APOC2 | 0.01915439 | 0.72264395 | 1 |
| TNFAIP2 | 7.84E-05 | 0.72228263 | 1 |
| PCDH10 | 7.07E-05 | 0.72218814 | 1 |
| ELF5 | 0.01670954 | 0.72169271 | 1 |
| B9D2 | 0.0004721 | 0.72124586 | 1 |
| IYD | 0.00110179 | 0.7208937 | 1 |
| APOBEC3D | 0.00253332 | 0.72050737 | 1 |
| GGCX | 0.00061964 | 0.72025455 | 1 |
| ARL3 | 0.00038954 | 0.71918644 | 1 |
| LRTOMT | 0.00265572 | 0.71874167 | 1 |
| SMG9 | 0.00114289 | 0.71855941 | 1 |
| FBXO41 | 0.00434895 | 0.71845844 | 1 |
| POLR2G | 0.00014945 | 0.71777193 | 1 |
| FANCM | 0.00038199 | 0.7176759 | 1 |
| USP53 | 0.0001351 | 0.71619917 | 1 |
| CEP170 | 0.00092769 | 0.71618187 | 1 |
| CSF2RB | 0.0009549 | 0.71594847 | 1 |
| MYADM | 0.00011399 | 0.71592249 | 1 |
| FAM204A | 0.00392548 | 0.71554606 | 1 |
| TANC2 | 0.00025305 | 0.71543303 | 1 |
| QSOX2 | 0.00432159 | 0.71491512 | 1 |
| RDH13 | 0.00100828 | 0.71412343 | 1 |
| CLEC3B | 0.00014054 | 0.71380633 | 1 |
| DCTN4 | 0.00034913 | 0.71335726 | 1 |

| | | | |
|---|---|---|---|
| HAUS4 | 0.00010343 | 0.71318277 | 1 |
| EOGT | 0.00012903 | 0.71305865 | 1 |
| TMEM128 | 0.00021385 | 0.71185603 | 1 |
| C11orf54 | 0.00013378 | 0.71156741 | 1 |
| FCAR | 0.0019222 | 0.71154303 | 1 |
| NIBAN1 | 0.02549125 | 0.71101299 | 1 |
| ADA2 | 0.00031859 | 0.71074237 | 1 |
| SAMD9 | 0.00017709 | 0.71057559 | 1 |
| FAM189B | 0.00053531 | 0.71014017 | 1 |
| ASF1B | 0.04646225 | 0.71010662 | 1 |
| PPFIBP2 | 0.00049599 | 0.7092181 | 1 |
| SOCS6 | 0.00011648 | 0.70909098 | 1 |
| GPC4 | 8.31E-05 | 0.70864688 | 1 |
| ZNF260 | 0.00884587 | 0.70823481 | 1 |
| C16orf70 | 7.51E-05 | 0.70806451 | 1 |
| ARHGAP26 | 0.00102648 | 0.70798373 | 1 |
| CNOT4 | 0.00010587 | 0.70770387 | 1 |
| FBXL2 | 0.0250906 | 0.70747834 | 1 |
| BCOR | 7.45E-05 | 0.70740089 | 1 |
| SPOP | 0.00012036 | 0.7072849 | 1 |
| C6orf62 | 7.84E-05 | 0.70705143 | 1 |
| DISC1 | 0.00026542 | 0.70675299 | 1 |
| LACTB | 0.00033012 | 0.70623074 | 1 |
| LY96 | 7.49E-05 | 0.70461008 | 1 |
| ERP27 | 0.0009297 | 0.70344854 | 1 |
| EPSTI1 | 7.84E-05 | 0.70317057 | 1 |
| RBP7 | 0.00024707 | 0.70313407 | 1 |
| SLC26A2 | 0.00039585 | 0.70173601 | 1 |
| EEPD1 | 0.00339627 | 0.70066164 | 1 |
| DFFA | 6.39E-05 | 0.7006021 | 1 |
| IL1B | 0.00236709 | 0.70053371 | 1 |
| OFD1 | 6.17E-05 | 0.70040221 | 1 |
| ZFP1 | 0.00676468 | 0.69985876 | 1 |
| PDK3 | 0.00217034 | 0.6997594 | 1 |
| CSF2RA | 0.00161966 | 0.69944173 | 1 |
| MLLT1 | 0.00055096 | 0.69871698 | 1 |
| ACY1 | 0.00022148 | 0.69863459 | 1 |
| PRELID3B | 0.00066644 | 0.69796802 | 1 |
| MTHFD2 | 0.00011796 | 0.69775937 | 1 |
| C15orf61 | 0.00112869 | 0.69772484 | 1 |
| MNAT1 | 0.00242114 | 0.69757146 | 1 |
| BCAT1 | 0.00233423 | 0.69724 | 1 |
| HFE | 0.00279724 | 0.69709205 | 1 |
| LMBR1 | 0.00037607 | 0.69638129 | 1 |
| ATG2A | 0.007214 | 0.69620406 | 1 |
| SSU72 | 0.00022071 | 0.69597212 | 1 |
| ZNF789 | 0.01457277 | 0.69591021 | 1 |
| LGI2 | 0.03396976 | 0.6956586 | 1 |
| ACO2 | 0.00019355 | 0.69483607 | 1 |
| HAGHL | 0.02260259 | 0.69480694 | 1 |
| ZNF181 | 0.00793595 | 0.69458517 | 1 |
| TOR4A | 0.0008106 | 0.69418972 | 1 |
| ZNF3 | 0.00040961 | 0.69350235 | 1 |

| | | | |
|---|---|---|---|
| CLEC7A | 0.00011905 | 0.69317485 | 1 |
| PLA2G4A | 0.01093176 | 0.69279823 | 1 |
| AFAP1 | 0.00013338 | 0.69271756 | 1 |
| MEX3B | 0.04218912 | 0.69227847 | 1 |
| SEMA5B | 0.00137063 | 0.69208893 | 1 |
| TSC22D3 | 0.00069941 | 0.69167863 | 1 |
| TMEM213 | 0.00060197 | 0.69153612 | 1 |
| PRR5 | 0.00134391 | 0.69075721 | 1 |
| PNPO | 0.00074155 | 0.69074147 | 1 |
| CLIC2 | 0.00709694 | 0.69006089 | 1 |
| TMC4 | 0.00204737 | 0.68887363 | 1 |
| SLC39A13 | 0.00019534 | 0.68857111 | 1 |
| RAB9A | 0.00753841 | 0.68856388 | 1 |
| NRROS | 0.00051042 | 0.68849592 | 1 |
| ZNF696 | 0.00186601 | 0.68763545 | 1 |
| AMZ2 | 0.00454701 | 0.68719193 | 1 |
| GJA3 | 0.00923842 | 0.68659416 | 1 |
| RIBC1 | 0.04528727 | 0.68586293 | 1 |
| FYCO1 | 0.04069948 | 0.68493112 | 1 |
| YDJC | 0.00394614 | 0.68482128 | 1 |
| MT1M | 0.00441844 | 0.6845394 | 1 |
| PROK2 | 0.00430843 | 0.68441824 | 1 |
| TCEAL4 | 6.08E-05 | 0.68372192 | 1 |
| TMEM65 | 0.0003075 | 0.68345186 | 1 |
| LILRB1 | 0.00150723 | 0.6834237 | 1 |
| CD44 | 0.00025146 | 0.68291716 | 1 |
| TGFBI | 0.00014092 | 0.68243498 | 1 |
| FGFR2 | 0.00037206 | 0.6821964 | 1 |
| UPB1 | 0.00014152 | 0.68182314 | 1 |
| ENO1 | 0.00021207 | 0.68134995 | 1 |
| ZNRD1 | 0.00225323 | 0.68134982 | 1 |
| ERI2 | 0.00049724 | 0.68115499 | 1 |
| ZNF234 | 0.00103502 | 0.6806601 | 1 |
| CDC7 | 0.00780312 | 0.68044949 | 1 |
| NEDD4 | 0.00035226 | 0.67892928 | 1 |
| NARS | 6.19E-05 | 0.67886798 | 1 |
| INO80B | 0.00117457 | 0.67882891 | 1 |
| GFOD2 | 0.00092778 | 0.67877113 | 1 |
| RBBP5 | 0.00227175 | 0.67862854 | 1 |
| COX6B1 | 0.00382761 | 0.67840642 | 1 |
| DEPDC5 | 0.00100239 | 0.67786542 | 1 |
| HMGCS1 | 0.00350712 | 0.67785025 | 1 |
| STK17A | 0.00419142 | 0.67777976 | 1 |
| MBNL1 | 5.56E-05 | 0.67745795 | 1 |
| DCAF1 | 0.00205582 | 0.67715406 | 1 |
| FNDC10 | 0.01109732 | 0.67711618 | 1 |
| C1orf116 | 0.01778465 | 0.67593834 | 1 |
| PCP4 | 0.00108005 | 0.67548962 | 1 |
| PPP1R14A | 0.00034425 | 0.67494978 | 1 |
| MFHAS1 | 0.00159923 | 0.67454313 | 1 |
| METTL7A | 0.00016496 | 0.67426585 | 1 |
| SLC16A3 | 0.00120698 | 0.67354575 | 1 |
| YIF1B | 0.00015299 | 0.67325672 | 1 |

| | | | |
|---|---|---|---|
| SLC16A1 | 0.00052698 | 0.67308616 | 1 |
| BIN3 | 0.00068893 | 0.67237884 | 1 |
| ASL | 0.0009444 | 0.6718198 | 1 |
| NCEH1 | 0.00039656 | 0.67169159 | 1 |
| ECHS1 | 0.00040174 | 0.67157203 | 1 |
| CP | 0.04362521 | 0.67129269 | 1 |
| NIPA1 | 0.01294181 | 0.67079255 | 1 |
| NCKAP5L | 9.29E-05 | 0.67046709 | 1 |
| PGAP3 | 0.00069101 | 0.67045444 | 1 |
| ACAT1 | 0.00047925 | 0.66958054 | 1 |
| STK17B | 0.00016247 | 0.66925925 | 1 |
| UBA3 | 8.30E-05 | 0.66918306 | 1 |
| P2RX7 | 0.00077355 | 0.66906492 | 1 |
| SMTNL2 | 0.00032774 | 0.66890679 | 1 |
| C3orf14 | 0.02049592 | 0.6685877 | 1 |
| RNF34 | 0.01351567 | 0.6682102 | 1 |
| FBRSL1 | 0.00136696 | 0.66820728 | 1 |
| PGBD1 | 0.00515916 | 0.66771692 | 1 |
| CLDN2 | 0.00011535 | 0.66738624 | 1 |
| ACSL1 | 9.07E-05 | 0.66615249 | 1 |
| STAT5A | 0.00138418 | 0.66592751 | 1 |
| ERCC2 | 0.00093125 | 0.66589245 | 1 |
| LIN54 | 0.00033683 | 0.6653699 | 1 |
| IKBKE | 0.00084607 | 0.66510725 | 1 |
| GPBAR1 | 0.01109732 | 0.66489935 | 1 |
| CD109 | 0.0001276 | 0.6644746 | 1 |
| ACPP | 0.00239587 | 0.66447173 | 1 |
| NUTF2 | 0.00132278 | 0.66437397 | 1 |
| PLLP | 0.00218009 | 0.66427761 | 1 |
| ZFYVE26 | 0.00015725 | 0.66337879 | 1 |
| THAP12 | 0.00684337 | 0.66330251 | 1 |
| EML4 | 0.00111861 | 0.66270661 | 1 |
| ALDH1A2 | 6.76E-05 | 0.66239751 | 1 |
| ZNF131 | 0.00501294 | 0.6621547 | 1 |
| EFNB3 | 0.02686771 | 0.6619356 | 1 |
| HSPA4 | 0.00033994 | 0.66175082 | 1 |
| HAX1 | 5.91E-05 | 0.66083826 | 1 |
| PHOSPHO2 | 0.00052825 | 0.6606741 | 1 |
| DUS3L | 0.00456833 | 0.66055724 | 1 |
| RBMS3 | 0.00029442 | 0.66042789 | 1 |
| SH3YL1 | 0.00063037 | 0.66025252 | 1 |
| RUNX2 | 0.01647013 | 0.66004689 | 1 |
| UCHL5 | 0.00097312 | 0.659971 | 1 |
| TAPT1 | 0.00072269 | 0.65959214 | 1 |
| NAALADL2 | 0.00476682 | 0.65913413 | 1 |
| CASTOR1 | 0.00054467 | 0.65908787 | 1 |
| ARHGAP25 | 0.00019967 | 0.6590455 | 1 |
| ERCC5 | 0.00090021 | 0.65869241 | 1 |
| AMH | 0.01090561 | 0.65777861 | 1 |
| XIAP | 0.00889209 | 0.65753911 | 1 |
| DNAL4 | 0.00112875 | 0.65746569 | 1 |
| PTPRC | 0.00268993 | 0.65729004 | 1 |
| NAF1 | 0.01122212 | 0.65677472 | 1 |

| | | | |
|---|---|---|---|
| CCNT2 | 0.00019515 | 0.65648556 | 1 |
| JAK1 | 0.000135 | 0.65631812 | 1 |
| C2 | 0.0069527 | 0.6552151 | 1 |
| NUFIP2 | 0.00222452 | 0.65512629 | 1 |
| ZNF714 | 0.02686771 | 0.65495868 | 1 |
| LSM1 | 0.00013632 | 0.65442204 | 1 |
| YPEL2 | 0.0006979 | 0.65411597 | 1 |
| G6PC | 0.00047976 | 0.65301891 | 1 |
| ERVK3-1 | 0.03468912 | 0.65277861 | 1 |
| WSCD2 | 0.0308649 | 0.65203581 | 1 |
| TLK2 | 0.0002741 | 0.65175037 | 1 |
| KDM6B | 0.00092477 | 0.6514588 | 1 |
| LRCH2 | 0.00242775 | 0.65144886 | 1 |
| SPRTN | 0.00159921 | 0.65047235 | 1 |
| YAF2 | 0.00033781 | 0.65025697 | 1 |
| KLHL9 | 0.00083503 | 0.64985635 | 1 |
| TMEM108 | 0.02686771 | 0.64960745 | 1 |
| BTK | 0.00454488 | 0.64951132 | 1 |
| PHGDH | 0.00745904 | 0.64912787 | 1 |
| EPS15 | 0.00028096 | 0.64889434 | 1 |
| ACACB | 0.00012745 | 0.64877822 | 1 |
| RAB1B | 0.00028274 | 0.64870786 | 1 |
| SCAI | 0.0044881 | 0.64845818 | 1 |
| AGXT2 | 0.00012334 | 0.64793667 | 1 |
| GPR182 | 0.04989882 | 0.647826 | 1 |
| TBCE.1 | 0.0014723 | 0.64775652 | 1 |
| SRPK1 | 9.43E-05 | 0.64771559 | 1 |
| C1orf131 | 0.00335007 | 0.64749622 | 1 |
| SIM2 | 0.01092706 | 0.64703157 | 1 |
| C14orf132 | 0.01053227 | 0.6467172 | 1 |
| LY6G5C | 0.02745061 | 0.64659925 | 1 |
| APOE | 0.00943438 | 0.64657442 | 1 |
| TOMM70 | 0.00125718 | 0.64535145 | 1 |
| TSPAN1 | 0.00020438 | 0.64525493 | 1 |
| UBXN7 | 0.00024503 | 0.64520115 | 1 |
| LRRC58 | 0.0003228 | 0.644607 | 1 |
| SARDH | 0.00433182 | 0.64407434 | 1 |
| HOXA9 | 0.00648819 | 0.64338341 | 1 |
| PPP1R3A | 0.02032656 | 0.64292226 | 1 |
| KLHDC7B | 0.02651478 | 0.64284284 | 1 |
| GBP4 | 8.23E-05 | 0.64271435 | 1 |
| NECAP2 | 0.00029445 | 0.64176923 | 1 |
| EXOSC2 | 0.00638276 | 0.64153316 | 1 |
| PCNT | 0.00019815 | 0.64112773 | 1 |
| RABEP1 | 6.18E-05 | 0.64048071 | 1 |
| MAB21L4 | 0.00038374 | 0.64039187 | 1 |
| INO80C | 0.00011437 | 0.63992804 | 1 |
| MTHFD1L | 0.01131002 | 0.63972279 | 1 |
| APOBEC3A | 0.0175649 | 0.63967646 | 1 |
| ZBTB26 | 0.00031136 | 0.63902876 | 1 |
| FLNB | 0.00033324 | 0.63894516 | 1 |
| TAFA5 | 0.02686771 | 0.63889995 | 1 |
| TSC22D4 | 0.00091947 | 0.63846592 | 1 |

| | | | |
|---|---|---|---|
| FAM110D | 0.02349625 | 0.63822451 | 1 |
| RRP9 | 0.00419142 | 0.63809799 | 1 |
| CDK8 | 0.00875518 | 0.63753538 | 1 |
| ABCA7 | 0.00012477 | 0.63749078 | 1 |
| ACSL5 | 0.00025781 | 0.63712072 | 1 |
| RBM45 | 0.00053788 | 0.63697677 | 1 |
| LIMA1 | 0.00068233 | 0.63650089 | 1 |
| HECA | 0.0001734 | 0.63526506 | 1 |
| UGT1A1 | 0.01474577 | 0.63525928 | 1 |
| FDPS | 0.00013617 | 0.63513723 | 1 |
| LYSMD2 | 0.00050129 | 0.63505129 | 1 |
| ICAM1 | 5.91E-05 | 0.63485291 | 1 |
| ATP5PF | 0.00114006 | 0.63481772 | 1 |
| ALDH6A1 | 0.00192079 | 0.63463871 | 1 |
| DNAH1 | 0.00284644 | 0.63452688 | 1 |
| ZNF317 | 0.00182241 | 0.63413509 | 1 |
| ALDH2 | 0.00064611 | 0.6340799 | 1 |
| ABCB8 | 8.73E-05 | 0.63401559 | 1 |
| LARP1 | 7.31E-05 | 0.63356113 | 1 |
| CLIC4 | 0.0004215 | 0.6333879 | 1 |
| SPATA2L | 0.00117723 | 0.63337799 | 1 |
| FCF1 | 0.00030016 | 0.63323529 | 1 |
| WDR72 | 0.00048766 | 0.63298758 | 1 |
| ERGIC1 | 0.00154424 | 0.63261997 | 1 |
| SURF6 | 0.00042602 | 0.6324027 | 1 |
| ARHGEF5 | 0.00260953 | 0.63233059 | 1 |
| PRAG1 | 0.01118993 | 0.63202774 | 1 |
| CACNA2D4 | 0.02002038 | 0.63147074 | 1 |
| CKB | 0.00051392 | 0.63137908 | 1 |
| SLC9A9 | 0.01384461 | 0.63102729 | 1 |
| STARD4 | 0.00311531 | 0.63101101 | 1 |
| NR6A1 | 0.00817935 | 0.630816 | 1 |
| ZNF136 | 0.00331468 | 0.63071293 | 1 |
| RUNX1T1 | 0.02695679 | 0.63061367 | 1 |
| SSBP4 | 0.00402658 | 0.63044279 | 1 |
| PAFAH1B3 | 0.00186274 | 0.62951996 | 1 |
| SLC26A11 | 0.00018082 | 0.6295083 | 1 |
| STRADB | 0.01109732 | 0.62915584 | 1 |
| NXF3 | 0.00189393 | 0.6291486 | 1 |
| GLUL | 0.00107373 | 0.62846351 | 1 |
| PHYHD1 | 0.0020729 | 0.62822446 | 1 |
| WASHC3 | 0.00177188 | 0.62815416 | 1 |
| KLHL2 | 0.00012541 | 0.62800268 | 1 |
| ITGB7 | 0.02651478 | 0.62783515 | 1 |
| FAM76B | 8.01E-05 | 0.62767955 | 1 |
| DNAJB1 | 0.00011966 | 0.62754584 | 1 |
| CCDC137 | 0.00170306 | 0.62715517 | 1 |
| PCGF3 | 0.00658831 | 0.62708183 | 1 |
| GOLPH3L | 0.00313473 | 0.62688976 | 1 |
| SLC9A3 | 0.00386399 | 0.62683882 | 1 |
| PRKAR2A | 0.00016285 | 0.62645216 | 1 |
| HEYL | 0.01859722 | 0.62631389 | 1 |
| PSMB2 | 0.00010513 | 0.62534451 | 1 |

| | | | |
|---|---|---|---|
| LENG1 | 0.0002296 | 0.62439598 | 1 |
| PPP1R8 | 0.00032323 | 0.62422344 | 1 |
| TMEM18 | 7.19E-05 | 0.62410878 | 1 |
| ULK3 | 0.00016001 | 0.62337024 | 1 |
| CENPP | 0.0009126 | 0.62331137 | 1 |
| XPO5 | 0.00054833 | 0.62215185 | 1 |
| MAP3K14 | 0.00111108 | 0.62146885 | 1 |
| ADRA2C | 0.00335764 | 0.6207337 | 1 |
| ICAM4 | 0.02745061 | 0.62042652 | 1 |
| CCDC28A | 0.00050294 | 0.62039856 | 1 |
| CMPK2 | 0.00093897 | 0.62009046 | 1 |
| ATP5F1C | 0.00011962 | 0.61983041 | 1 |
| FAM177A1 | 0.00065832 | 0.6196353 | 1 |
| MNDA | 0.00074367 | 0.61954194 | 1 |
| C16orf46 | 0.04262532 | 0.61951131 | 1 |
| SLFN11 | 0.00020151 | 0.61875861 | 1 |
| TSNARE1 | 0.00079474 | 0.61854621 | 1 |
| ZNF782 | 0.00300584 | 0.61854113 | 1 |
| CFDP1 | 6.89E-05 | 0.61838932 | 1 |
| BOP1 | 0.01031448 | 0.61813104 | 1 |
| ZXDC | 0.00023816 | 0.61808268 | 1 |
| ASXL1 | 0.00024653 | 0.61774591 | 1 |
| FAM162B | 0.00033571 | 0.61762195 | 1 |
| LITAF | 0.00013506 | 0.61761632 | 1 |
| MYBPH | 0.00479589 | 0.61737137 | 1 |
| GLYAT | 0.00164443 | 0.61697223 | 1 |
| RBMXL1 | 0.01853018 | 0.61661164 | 1 |
| GNB5 | 0.00034658 | 0.61648019 | 1 |
| AKIRIN2 | 0.00014833 | 0.61607143 | 1 |
| TBC1D16 | 0.00187021 | 0.61595302 | 1 |
| COQ10B | 0.00015663 | 0.61594275 | 1 |
| PDE6B | 0.01908676 | 0.61580161 | 1 |
| PARM1 | 0.00013182 | 0.61541519 | 1 |
| BPHL | 0.00027486 | 0.61457831 | 1 |
| ABAT | 0.00065283 | 0.61426818 | 1 |
| RBM43 | 9.82E-05 | 0.61412964 | 1 |
| SLC11A2 | 0.00022075 | 0.61411411 | 1 |
| AFF1 | 8.21E-05 | 0.61379683 | 1 |
| ZNRF1 | 0.00056673 | 0.61302344 | 1 |
| MMUT | 0.00011282 | 0.61222228 | 1 |
| SH2B3 | 0.00019899 | 0.61094095 | 1 |
| PSPN | 0.04989882 | 0.61086552 | 1 |
| CPNE6 | 0.01037103 | 0.61050587 | 1 |
| TNRC6B | 0.0001447 | 0.61037708 | 1 |
| TPCN1 | 0.00012694 | 0.61001864 | 1 |
| CRH | 0.01503064 | 0.60907386 | 1 |
| IQCD | 0.00441844 | 0.60847849 | 1 |
| CCDC86 | 0.03239992 | 0.60824742 | 1 |
| ANK3 | 0.00016022 | 0.60804352 | 1 |
| LIG4 | 0.00095608 | 0.60788328 | 1 |
| MMS22L | 0.00817935 | 0.60764836 | 1 |
| PPP4R3A | 0.00042388 | 0.60703873 | 1 |
| RHOJ | 0.01373372 | 0.60657783 | 1 |

| | | | |
|---|---|---|---|
| PLEKHG4 | 0.00589811 | 0.60652159 | 1 |
| MTPN | 0.00011018 | 0.6062141 | 1 |
| CASP4 | 0.00091522 | 0.60606934 | 1 |
| RHBDF2 | 0.0024428 | 0.60581375 | 1 |
| TSTA3 | 0.00146024 | 0.60458774 | 1 |
| TTF2 | 0.00268228 | 0.60454003 | 1 |
| CGRRF1 | 0.00582679 | 0.60423193 | 1 |
| MAP4 | 0.00064857 | 0.60380169 | 1 |
| GLUD1 | 8.12E-05 | 0.60332469 | 1 |
| SDHB | 8.13E-05 | 0.60301225 | 1 |
| COQ10A | 0.01482194 | 0.60293742 | 1 |
| SURF2 | 0.00662818 | 0.60211798 | 1 |
| TRIB1 | 0.00121687 | 0.60210282 | 1 |
| ATP5PB | 0.00012982 | 0.60170732 | 1 |
| PCTP | 0.00030069 | 0.60163468 | 1 |
| PACS2 | 6.10E-05 | 0.60125987 | 1 |
| CXCR6 | 0.02745061 | 0.60083256 | 1 |
| THOC1 | 0.02295934 | 0.60078166 | 1 |
| NID1 | 0.00015246 | 0.59976586 | 1 |
| ZSCAN16 | 0.03237176 | 0.59922189 | 1 |
| DISP1 | 0.00209001 | 0.59914877 | 1 |
| SPHK2 | 9.01E-05 | 0.59891882 | 1 |
| SYDE1 | 0.00111844 | 0.59871135 | 1 |
| BLCAP | 0.00010728 | 0.59851163 | 1 |
| MILR1 | 0.02555541 | 0.59820009 | 1 |
| CFAP70 | 0.0059161 | 0.59806959 | 1 |
| NME5 | 0.01101704 | 0.59803507 | 1 |
| PARL | 0.00027637 | 0.59788641 | 1 |
| SLC25A25 | 0.00016284 | 0.59782963 | 1 |
| ZFHX3 | 0.0001732 | 0.59717981 | 1 |
| ADO | 0.00126132 | 0.59675295 | 1 |
| ZNF485 | 0.02704124 | 0.59526319 | 1 |
| UBE4B | 0.0023053 | 0.59524235 | 1 |
| PAX8 | 0.00211845 | 0.5947077 | 1 |
| ATPAF2 | 0.00601756 | 0.59461045 | 1 |
| NTMT1 | 8.10E-05 | 0.59347493 | 1 |
| OSBPL11 | 5.73E-05 | 0.59332049 | 1 |
| TMEM140 | 0.00025088 | 0.59293726 | 1 |
| ZNF334 | 0.00105047 | 0.59260788 | 1 |
| SAMD9L | 0.00019198 | 0.59256519 | 1 |
| WDR55 | 0.00018311 | 0.59212166 | 1 |
| SIGLEC11 | 0.00236709 | 0.59168538 | 1 |
| HECTD2 | 0.00211665 | 0.59165753 | 1 |
| HAUS2 | 0.00322764 | 0.5915182 | 1 |
| CXCL14 | 0.0007897 | 0.59142926 | 1 |
| STC1 | 0.00011636 | 0.59132304 | 1 |
| CAB39L | 0.0017039 | 0.5911584 | 1 |
| ADORA2A | 0.00460811 | 0.59064371 | 1 |
| CHST10 | 0.01995308 | 0.59016087 | 1 |
| RNF144A | 0.00015029 | 0.58967191 | 1 |
| PROCR | 0.00614654 | 0.58908921 | 1 |
| HNRNPM | 0.00016382 | 0.58884151 | 1 |
| TMUB1 | 0.00448037 | 0.58838145 | 1 |

| | | | |
|---|---|---|---|
| PROM2 | 0.00121304 | 0.58797302 | 1 |
| TDRD10 | 0.02022291 | 0.58794204 | 1 |
| VCAM1 | 0.00471561 | 0.58770247 | 1 |
| SDE2 | 0.00010691 | 0.58767075 | 1 |
| TSTD2 | 0.0146075 | 0.58631167 | 1 |
| ISCU | 0.00047384 | 0.58626869 | 1 |
| POLR1A | 0.0266209 | 0.58613205 | 1 |
| RNF24 | 0.00302308 | 0.58610766 | 1 |
| SLC49A4 | 0.00026879 | 0.58597711 | 1 |
| EGLN1 | 0.00080461 | 0.58517038 | 1 |
| ZBTB33 | 0.00017728 | 0.5851207 | 1 |
| TDRD5 | 0.03803087 | 0.58508984 | 1 |
| PCDH20 | 0.04989882 | 0.58503569 | 1 |
| UFM1 | 0.00024957 | 0.58456241 | 1 |
| PRR22 | 0.03161063 | 0.5844188 | 1 |
| CDC42SE1 | 0.00027494 | 0.58437542 | 1 |
| ARL5A | 0.00032572 | 0.5842683 | 1 |
| TMPO | 0.0009295 | 0.58401808 | 1 |
| ZFP14 | 0.00060633 | 0.58334414 | 1 |
| LSR | 0.00086809 | 0.58317395 | 1 |
| SLC37A2 | 0.00504059 | 0.58316281 | 1 |
| UGT2A3 | 0.00607805 | 0.58275433 | 1 |
| GRK3 | 0.00058238 | 0.58255164 | 1 |
| IFRD1 | 0.00090568 | 0.5813339 | 1 |
| ILF2 | 9.84E-05 | 0.58112001 | 1 |
| FUT4 | 0.00427453 | 0.58087153 | 1 |
| FES | 0.00115828 | 0.58008806 | 1 |
| TAF5 | 0.00601756 | 0.57952479 | 1 |
| MED28 | 0.00019886 | 0.57901428 | 1 |
| BAIAP2L1 | 0.00150723 | 0.57881964 | 1 |
| ABHD12 | 0.0002437 | 0.57871881 | 1 |
| CRACR2B | 0.00787452 | 0.57861657 | 1 |
| MANBA | 0.00465713 | 0.57851067 | 1 |
| SORBS2 | 0.00071932 | 0.57833477 | 1 |
| CTCF | 7.44E-05 | 0.57813345 | 1 |
| EEF2K | 7.53E-05 | 0.57793896 | 1 |
| GUCD1 | 0.0001886 | 0.57791306 | 1 |
| ARNT2 | 0.00090755 | 0.57764464 | 1 |
| OR2G6 | 0.02032656 | 0.57756334 | 1 |
| NBDY | 0.01977919 | 0.5763309 | 1 |
| EEF1AKMT3 | 0.00088557 | 0.57629155 | 1 |
| PRRT1 | 0.01136809 | 0.57597689 | 1 |
| PLA1A | 0.00025203 | 0.57594018 | 1 |
| SOD1 | 0.00186207 | 0.57588756 | 1 |
| ALDOB | 0.00254907 | 0.57547114 | 1 |
| WHRN | 0.00067549 | 0.57450518 | 1 |
| RBKS | 0.00037564 | 0.57447576 | 1 |
| PILRB | 0.00236788 | 0.5742803 | 1 |
| TMA16 | 0.00028144 | 0.57427265 | 1 |
| GSR | 0.00020898 | 0.57419335 | 1 |
| KPNA5 | 0.01230969 | 0.57405395 | 1 |
| PML | 0.00092428 | 0.57373181 | 1 |
| PDCD6 | 0.00013128 | 0.57290615 | 1 |

| | | | |
|---|---|---|---|
| ADNP2 | 0.00041658 | 0.57255232 | 1 |
| ZNF736 | 0.0186879 | 0.5724682 | 1 |
| SART1 | 0.00254674 | 0.57224378 | 1 |
| POLE2 | 0.03468912 | 0.57171281 | 1 |
| FOXE3 | 0.04021005 | 0.57143058 | 1 |
| CWF19L1 | 0.00367824 | 0.57111401 | 1 |
| DGKG | 0.03980533 | 0.57086734 | 1 |
| TIAF1 | 0.00075162 | 0.56968863 | 1 |
| DCTN3 | 0.00063016 | 0.56938182 | 1 |
| EVI5 | 0.00014926 | 0.56917317 | 1 |
| HOMEZ | 0.00122775 | 0.56888459 | 1 |
| ATXN10 | 0.0001998 | 0.56845964 | 1 |
| ZNF628 | 0.0406836 | 0.56829596 | 1 |
| PANK1 | 0.0004054 | 0.56797499 | 1 |
| PEX11B | 0.00190304 | 0.56769346 | 1 |
| MUC1 | 0.00103678 | 0.56759135 | 1 |
| CCDC125 | 0.00020649 | 0.56751557 | 1 |
| PDE10A | 0.00123728 | 0.56740559 | 1 |
| PDGFRA | 0.0015813 | 0.56705484 | 1 |
| C16orf74 | 0.04551531 | 0.56657407 | 1 |
| UTP6 | 0.00047518 | 0.56656791 | 1 |
| ZNF512B | 0.0025381 | 0.56652339 | 1 |
| ATG16L2 | 0.00413289 | 0.56608512 | 1 |
| TALDO1 | 0.00129317 | 0.56575141 | 1 |
| STAT1 | 0.00591755 | 0.56549001 | 1 |
| ZNF883 | 0.03702779 | 0.56544679 | 1 |
| ZNF641 | 0.00136109 | 0.56491333 | 1 |
| GHSR | 0.04989882 | 0.56487172 | 1 |
| CATSPERG | 0.01327217 | 0.56475791 | 1 |
| FMN2 | 0.02666836 | 0.56465573 | 1 |
| PID1 | 0.01550942 | 0.56448632 | 1 |
| NOTCH2 | 0.00064196 | 0.56424794 | 1 |
| GALNT7 | 0.00825083 | 0.5641299 | 1 |
| TOGARAM1 | 0.00228705 | 0.56408275 | 1 |
| NXPE3 | 0.0008638 | 0.56402533 | 1 |
| PAH | 0.00016884 | 0.56358184 | 1 |
| USP19 | 0.00018136 | 0.56339287 | 1 |
| WDR5 | 0.00015274 | 0.56310061 | 1 |
| OAS2 | 0.01849221 | 0.56290694 | 1 |
| ECSCR | 0.00035015 | 0.56283353 | 1 |
| NDRG1 | 0.00140939 | 0.56250978 | 1 |
| ID2 | 0.00023494 | 0.5625085 | 1 |
| ARHGAP4 | 0.00076107 | 0.56241682 | 1 |
| NOC2L | 7.56E-05 | 0.56213541 | 1 |
| TMEM208 | 8.10E-05 | 0.56186145 | 1 |
| CCNA2 | 0.02993241 | 0.56184807 | 1 |
| OTUD7B | 0.00135171 | 0.56140938 | 1 |
| PPP3CC | 0.00303384 | 0.56140323 | 1 |
| CC2D1B | 0.00118598 | 0.56086518 | 1 |
| ACAD9 | 0.00098174 | 0.56062128 | 1 |
| EPHA7 | 0.00086585 | 0.56042145 | 1 |
| MTMR4 | 0.00147567 | 0.55991536 | 1 |
| MMAA | 0.04315549 | 0.55947499 | 1 |

| | | | |
|---|---|---|---|
| FN3KRP | 0.00072557 | 0.55942836 | 1 |
| PGAM1 | 0.00012589 | 0.55941965 | 1 |
| AKAP7 | 0.00152781 | 0.55924639 | 1 |
| CNEP1R1 | 0.00087965 | 0.55893581 | 1 |
| GRK5 | 9.12E-05 | 0.55893256 | 1 |
| ERFE | 0.02032656 | 0.55871618 | 1 |
| COX8A | 0.00055021 | 0.55863228 | 1 |
| EPGN | 0.04989882 | 0.55857528 | 1 |
| GRIK2 | 0.03881848 | 0.5584813 | 1 |
| PSMG1 | 0.00315681 | 0.55784763 | 1 |
| TMEM204 | 0.00082877 | 0.55771991 | 1 |
| HEXD | 0.00054947 | 0.55716839 | 1 |
| APIP | 0.00167392 | 0.55650649 | 1 |
| ENPP5 | 0.03511061 | 0.55647944 | 1 |
| SLC25A1 | 0.00020064 | 0.55564911 | 1 |
| ZNF836 | 0.01439361 | 0.55505183 | 1 |
| PTGFRN | 0.00313325 | 0.55413358 | 1 |
| C15orf40 | 0.00041951 | 0.55307246 | 1 |
| CDKN2D | 6.19E-05 | 0.55266882 | 1 |
| MAPRE2 | 0.00032566 | 0.55262234 | 1 |
| FAM151A | 0.00099737 | 0.55246328 | 1 |
| SUMO3 | 0.00291251 | 0.55242011 | 1 |
| WDTC1 | 8.92E-05 | 0.55237674 | 1 |
| ZNF688 | 0.00491827 | 0.55198958 | 1 |
| ZNF264 | 0.00052933 | 0.55151621 | 1 |
| DTD1 | 0.00013327 | 0.55140949 | 1 |
| FLT1 | 0.00727843 | 0.55134441 | 1 |
| H2AFX | 0.0017039 | 0.55108465 | 1 |
| ZC3H7A | 0.00044318 | 0.55100415 | 1 |
| EVI2B | 0.00402732 | 0.55098278 | 1 |
| FOSB | 0.01421411 | 0.55094296 | 1 |
| HIVEP3 | 0.00027007 | 0.55082747 | 1 |
| FNBP4 | 0.00022085 | 0.55044948 | 1 |
| KRT7 | 0.00036094 | 0.54907214 | 1 |
| TIRAP | 0.00119144 | 0.54902154 | 1 |
| TMEM170A | 0.00611897 | 0.54827404 | 1 |
| PA2G4 | 0.00086384 | 0.54810495 | 1 |
| ZNF350 | 0.00396758 | 0.54738342 | 1 |
| FOXRED1 | 0.01666285 | 0.54736395 | 1 |
| EAF1 | 0.00061195 | 0.54729876 | 1 |
| METTL26 | 0.00491815 | 0.54725329 | 1 |
| CEACAM1 | 0.00165613 | 0.54717811 | 1 |
| UTP20 | 0.00576103 | 0.54715472 | 1 |
| TRMT10C | 0.0005666 | 0.54713639 | 1 |
| PRSS22 | 0.00817935 | 0.54609992 | 1 |
| COL6A2 | 0.00019094 | 0.54595871 | 1 |
| HERC5 | 0.00076944 | 0.54594364 | 1 |
| ZFYVE21 | 0.00049892 | 0.54590467 | 1 |
| PIGO | 0.00191487 | 0.54584652 | 1 |
| BCL11B | 0.04551531 | 0.5450665 | 1 |
| SCAMP5 | 0.04983694 | 0.54494128 | 1 |
| IL3RA | 0.00186274 | 0.54473665 | 1 |
| CCDC106 | 0.00015367 | 0.54458256 | 1 |

| | | | |
|---|---|---|---|
| RSRC2 | 0.00155686 | 0.54436869 | 1 |
| GLRX5 | 0.0003475 | 0.54435083 | 1 |
| LTBR | 0.00074593 | 0.54433429 | 1 |
| SLC22A8 | 0.00139067 | 0.54408772 | 1 |
| MFSD11 | 0.00017629 | 0.54366869 | 1 |
| DUSP12 | 0.00268749 | 0.54296039 | 1 |
| NFKBIA | 0.00051356 | 0.54289071 | 1 |
| MAST3 | 0.0047322 | 0.54197371 | 1 |
| MALT1 | 0.00060146 | 0.54185531 | 1 |
| ME2 | 0.00047141 | 0.54184866 | 1 |
| WDFY2 | 0.00016194 | 0.54161472 | 1 |
| GRAMD1B | 0.00070263 | 0.54125186 | 1 |
| PHC2 | 0.00111022 | 0.54099597 | 1 |
| ANKMY2 | 0.01281719 | 0.54077668 | 1 |
| CCNB1IP1 | 0.00490243 | 0.54008026 | 1 |
| YAP1 | 7.10E-05 | 0.53974104 | 1 |
| CREG1 | 0.00094107 | 0.53918753 | 1 |
| COX7B | 0.00027946 | 0.53822005 | 1 |
| LMOD1 | 0.01606816 | 0.53729382 | 1 |
| TAF6 | 0.00045359 | 0.53719274 | 1 |
| SAMD4B | 0.00085084 | 0.5371804 | 1 |
| ACLY | 0.00379482 | 0.53699345 | 1 |
| DDB2 | 0.00094851 | 0.53597873 | 1 |
| RBP4 | 0.01053804 | 0.53595246 | 1 |
| CLEC2D | 0.00031318 | 0.53582046 | 1 |
| FUT8 | 0.00100141 | 0.53525671 | 1 |
| C17orf58 | 0.00346343 | 0.53480607 | 1 |
| NAA38 | 0.00083693 | 0.53455638 | 1 |
| TFB1M | 0.0051622 | 0.53455061 | 1 |
| PMS1 | 0.00038633 | 0.53442514 | 1 |
| STARD8 | 0.00087566 | 0.53429288 | 1 |
| MBNL3 | 0.00050761 | 0.53423456 | 1 |
| SMCR8 | 0.00287244 | 0.53401882 | 1 |
| NRXN2 | 0.00047337 | 0.53383746 | 1 |
| PYCR3 | 0.00204955 | 0.53383539 | 1 |
| MMAB | 0.00012981 | 0.53353631 | 1 |
| BCR | 0.00065841 | 0.53336051 | 1 |
| SPR | 0.00083083 | 0.53321633 | 1 |
| PPARG | 8.95E-05 | 0.53320027 | 1 |
| FRAT1 | 0.00814465 | 0.53295651 | 1 |
| MGMT | 0.00041064 | 0.53293104 | 1 |
| MAP2K4 | 0.00062213 | 0.53289893 | 1 |
| RNF25 | 0.00086585 | 0.53281705 | 1 |
| SERPINF2 | 0.00492585 | 0.53235683 | 1 |
| NHS | 0.00015274 | 0.53206383 | 1 |
| NR4A3 | 0.03123586 | 0.53165431 | 1 |
| PSENEN | 9.08E-05 | 0.53109066 | 1 |
| ZNF652 | 5.56E-05 | 0.53108735 | 1 |
| TACC1 | 0.00094965 | 0.53103164 | 1 |
| COX4I2 | 0.00817935 | 0.53096615 | 1 |
| STAP2 | 0.00444882 | 0.53051248 | 1 |
| ELK1 | 0.00246311 | 0.53040713 | 1 |
| SERPINA5 | 0.00104195 | 0.52988785 | 1 |

| | | | |
|---|---|---|---|
| MCM5 | 0.00277842 | 0.52972605 | 1 |
| ADGRL2 | 0.00095057 | 0.52961081 | 1 |
| ORMDL2 | 0.00022629 | 0.52930089 | 1 |
| DCAF13 | 0.00317018 | 0.52877869 | 1 |
| USP13 | 0.00485113 | 0.52875653 | 1 |
| NOS1AP | 0.00152321 | 0.52863726 | 1 |
| UBR4 | 0.00047505 | 0.52764263 | 1 |
| CIPC | 0.00517292 | 0.52733309 | 1 |
| HOXB6 | 0.00316934 | 0.52731474 | 1 |
| CHD4 | 0.00169617 | 0.52728499 | 1 |
| CBX4 | 6.93E-05 | 0.52727455 | 1 |
| CPNE2 | 0.00247567 | 0.52698259 | 1 |
| CRTC3 | 0.00189547 | 0.5269196 | 1 |
| SLC46A1 | 0.02449036 | 0.5267684 | 1 |
| YWHAH | 0.00034092 | 0.52622401 | 1 |
| APH1A | 0.00014802 | 0.52621092 | 1 |
| SAPCD1 | 0.01647013 | 0.52584277 | 1 |
| PBRM1 | 0.00017415 | 0.52579376 | 1 |
| NUB1 | 0.00023231 | 0.52572223 | 1 |
| TLK1 | 0.00088012 | 0.52543869 | 1 |
| LURAP1 | 0.03324655 | 0.52509907 | 1 |
| PHF13 | 0.00146635 | 0.52475002 | 1 |
| FGFR1OP | 0.00233423 | 0.52469265 | 1 |
| MED15 | 0.00032925 | 0.52467002 | 1 |
| TNFSF13B | 0.01424723 | 0.52416339 | 1 |
| GPR151 | 0.02745061 | 0.5240589 | 1 |
| BSG | 0.00813256 | 0.52337952 | 1 |
| CYP2U1 | 0.00062083 | 0.52321575 | 1 |
| TCP1 | 0.00052048 | 0.52304259 | 1 |
| FUCA1 | 8.12E-05 | 0.52301299 | 1 |
| SAMD14 | 0.01114307 | 0.52242397 | 1 |
| PPP1CA | 0.00252025 | 0.5223471 | 1 |
| GNB2 | 0.00178391 | 0.52173166 | 1 |
| ALKBH4 | 0.00303806 | 0.52171557 | 1 |
| POLA2 | 0.04306052 | 0.52163237 | 1 |
| PKNOX1 | 0.00408519 | 0.52147071 | 1 |
| TINAGL1 | 6.91E-05 | 0.5211813 | 1 |
| PPA1 | 0.00057334 | 0.52097308 | 1 |
| DDX19B | 0.00209027 | 0.52080547 | 1 |
| LSM4 | 0.00201214 | 0.52026739 | 1 |
| PANK4 | 0.000462 | 0.51999967 | 1 |
| FAM8A1 | 6.31E-05 | 0.51992144 | 1 |
| TBC1D13 | 0.00022669 | 0.51969503 | 1 |
| DCAF12L2 | 0.00817935 | 0.51936354 | 1 |
| CCDC59 | 0.01247587 | 0.51904028 | 1 |
| MDM4 | 0.00134285 | 0.51800139 | 1 |
| GMPR2 | 7.80E-05 | 0.5173936 | 1 |
| SMU1 | 0.00087653 | 0.51719056 | 1 |
| TNNC2 | 0.00323754 | 0.51703055 | 1 |
| CMTM3 | 6.36E-05 | 0.51702639 | 1 |
| ATOH8 | 0.00124926 | 0.51698429 | 1 |
| MAP3K20 | 0.00023026 | 0.51649296 | 1 |
| COLEC11 | 0.00578349 | 0.51640882 | 1 |

| | | | |
|---|---|---|---|
| GPAT4 | 0.00043722 | 0.51616377 | 1 |
| PTPN1 | 0.00027876 | 0.51577752 | 1 |
| HSD17B11 | 0.00398965 | 0.5155271 | 1 |
| KLF4 | 0.00391142 | 0.51548427 | 1 |
| NDUFA4L2 | 0.00108171 | 0.51542276 | 1 |
| PFDN1 | 0.00018397 | 0.51520457 | 1 |
| TIPIN | 0.02032656 | 0.5151889 | 1 |
| RGS5 | 0.0205515 | 0.51508392 | 1 |
| PREX1 | 0.00172066 | 0.51445007 | 1 |
| NFIL3 | 0.00037406 | 0.51411195 | 1 |
| HHLA3 | 0.00076749 | 0.51365468 | 1 |
| KIF16B | 0.00033059 | 0.51356644 | 1 |
| HMGN5 | 0.0115686 | 0.51327765 | 1 |
| BTF3L4 | 0.00123558 | 0.51269817 | 1 |
| BTG3 | 0.00093125 | 0.512306 | 1 |
| MYL6B | 0.00220114 | 0.51202143 | 1 |
| TAGLN2 | 0.00023723 | 0.51159107 | 1 |
| PXDN | 0.00211436 | 0.51153371 | 1 |
| DDA1 | 0.0044326 | 0.51127833 | 1 |
| PRPSAP2 | 0.00029784 | 0.51046105 | 1 |
| TAGAP | 0.04213269 | 0.51009817 | 1 |
| CFB | 0.00114328 | 0.50909208 | 1 |
| PRKAG1 | 0.00195982 | 0.50898232 | 1 |
| FAM189A2 | 0.00642808 | 0.50884454 | 1 |
| ACADSB | 0.00020334 | 0.50883939 | 1 |
| SLC22A3 | 0.01503064 | 0.50879255 | 1 |
| DDT | 0.00080846 | 0.50878982 | 1 |
| PAPOLG | 0.00613597 | 0.50878506 | 1 |
| ZFAND4 | 0.00186109 | 0.50845117 | 1 |
| ERCC3 | 0.0045244 | 0.50797507 | 1 |
| TOMM40L | 0.01806597 | 0.50763212 | 1 |
| RIC8A | 0.00096578 | 0.50755278 | 1 |
| TBC1D10C | 0.01178689 | 0.50737186 | 1 |
| AP2S1 | 0.00010617 | 0.50706671 | 1 |
| TMEM99 | 0.01457512 | 0.50645625 | 1 |
| MTOR | 0.00264626 | 0.50622704 | 1 |
| MBLAC2 | 0.00249114 | 0.5060858 | 1 |
| RIN2 | 0.00295831 | 0.50575119 | 1 |
| U2AF2 | 0.00110464 | 0.50563802 | 1 |
| COL7A1 | 0.00127996 | 0.50554448 | 1 |
| ACAP2 | 0.00047795 | 0.50547699 | 1 |
| KCTD10 | 0.00123078 | 0.50498247 | 1 |
| CACFD1 | 0.00246191 | 0.50492187 | 1 |
| SIGLEC9 | 0.00044754 | 0.50471253 | 1 |
| ANP32B | 0.00433667 | 0.50461889 | 1 |
| MIDN | 0.00227761 | 0.50458086 | 1 |
| STK10 | 8.14E-05 | 0.5041635 | 1 |
| PDSS2 | 0.00024204 | 0.50414541 | 1 |
| GORAB | 0.00923842 | 0.50394124 | 1 |
| DNMT1 | 0.00022256 | 0.50323669 | 1 |
| SLC2A8 | 0.00695954 | 0.50286686 | 1 |
| NIP7 | 0.00031713 | 0.50235317 | 1 |
| ACP2 | 0.00174221 | 0.50150938 | 1 |

| | | | |
|---|---|---|---|
| TACC2 | 0.00081913 | 0.50149926 | 1 |
| YEATS2 | 0.00026406 | 0.50138936 | 1 |
| KLF3 | 0.00019793 | 0.50110694 | 1 |
| CDC27 | 0.00591593 | 0.50110014 | 1 |
| CTH | 0.00143657 | 0.5010189 | 1 |
| UBE2R2 | 0.00031446 | 0.50089655 | 1 |
| LAMA4 | 0.0011985 | 0.50069992 | 1 |
| ZNF554 | 0.00612426 | 0.50067925 | 1 |
| ANKDD1A | 0.00343091 | 0.50061624 | 1 |
| COX18 | 0.00104343 | 0.50010885 | 1 |
| LAMA3 | 0.02364826 | -0.5004654 | 1 |
| CCND1 | 0.01561796 | -0.5008284 | 1 |
| UBOX5 | 0.0279543 | -0.5014334 | 1 |
| REPS2 | 0.0289865 | -0.5016854 | 1 |
| MT-ND1 | 0.03886932 | -0.5017584 | 1 |
| ZNF587 | 0.03461672 | -0.5019365 | 1 |
| TMEM185B | 0.00366633 | -0.5033901 | 1 |
| PPIP5K2 | 0.02500088 | -0.503484 | 1 |
| PKP4 | 0.00917439 | -0.5043417 | 1 |
| TSSC4 | 0.01709042 | -0.5044094 | 1 |
| PIK3CG | 0.04502936 | -0.5045534 | 1 |
| CHD1 | 0.04991283 | -0.505151 | 1 |
| PLA2G4B | 0.01218068 | -0.5052093 | 1 |
| FAHD1 | 0.00580245 | -0.5060813 | 1 |
| SYNPO | 0.00267745 | -0.5063078 | 1 |
| ZNF436 | 0.00446144 | -0.507181 | 1 |
| RRAGC | 0.02071208 | -0.5079145 | 1 |
| MAT2B | 0.04301446 | -0.5080668 | 1 |
| SNTG2 | 0.04989882 | -0.5084936 | 1 |
| DDX10 | 0.04505191 | -0.5099569 | 1 |
| DSC2 | 0.04559283 | -0.5109363 | 1 |
| GZF1 | 0.00975424 | -0.5114501 | 1 |
| SLC25A26 | 0.0315008 | -0.5119272 | 1 |
| NUMBL | 0.01838144 | -0.5121991 | 1 |
| SPA17 | 0.00022493 | -0.5131645 | 1 |
| DDX19A | 0.03552946 | -0.5137664 | 1 |
| MAGED2 | 0.04356072 | -0.5143451 | 1 |
| SLC16A10 | 0.00637097 | -0.5144869 | 1 |
| CNOT6 | 0.03794573 | -0.5147045 | 1 |
| JAKMIP2 | 0.03819166 | -0.5159679 | 1 |
| NUP205 | 0.04919168 | -0.5175848 | 1 |
| TNRC6C | 0.04627931 | -0.5176188 | 1 |
| HYKK | 0.02576888 | -0.5176191 | 1 |
| AHI1 | 0.04892635 | -0.5176274 | 1 |
| ANKRD44 | 0.02944754 | -0.5188485 | 1 |
| NEMP1 | 0.04823223 | -0.5189661 | 1 |
| ARAP2 | 0.0185046 | -0.5194958 | 1 |
| LRRC8D | 0.01147749 | -0.5195754 | 1 |
| ZNF584 | 0.01560068 | -0.5211434 | 1 |
| HOXA10 | 0.00134248 | -0.5216165 | 1 |
| GCGR | 0.00612426 | -0.5219289 | 1 |
| HEATR1 | 0.03806818 | -0.5232025 | 1 |
| ZNF585B | 0.01508786 | -0.5236089 | 1 |

| | | | |
|---|---|---|---|
| WDR76 | 0.02032656 | -0.523952 | 1 |
| EML1 | 0.00446144 | -0.5256712 | 1 |
| PEA15 | 0.03157331 | -0.5267978 | 1 |
| SLC52A2 | 0.03510289 | -0.5277255 | 1 |
| TCEAL1 | 0.02653812 | -0.5277791 | 1 |
| SLC12A3 | 0.04881459 | -0.5280247 | 1 |
| LDAH | 0.02887598 | -0.5283529 | 1 |
| ASPSCR1 | 0.03860626 | -0.5300051 | 1 |
| TMED10 | 0.01694121 | -0.5307037 | 1 |
| ZNF500 | 0.01901828 | -0.5310591 | 1 |
| ECRG4 | 0.00589748 | -0.5318068 | 1 |
| SGMS1 | 0.0429967 | -0.5325571 | 1 |
| RIPOR1 | 0.00149783 | -0.5331562 | 1 |
| SKP2 | 0.04897279 | -0.5343369 | 1 |
| RIPK2 | 0.03462156 | -0.5347776 | 1 |
| BAK1 | 0.02786678 | -0.5350917 | 1 |
| CIR1 | 0.03607556 | -0.535429 | 1 |
| POLE | 0.02890653 | -0.537261 | 1 |
| TMEM220 | 0.01757794 | -0.5386816 | 1 |
| ZNF19 | 0.0416936 | -0.539026 | 1 |
| TADA2A | 0.04525088 | -0.5401963 | 1 |
| SPATA5 | 0.01751274 | -0.5408331 | 1 |
| TMEM63B | 0.01442531 | -0.5423135 | 1 |
| COMMD4 | 0.0135698 | -0.5430489 | 1 |
| COL18A1 | 0.00255353 | -0.5447988 | 1 |
| STRN | 0.0378567 | -0.5453492 | 1 |
| NUAK2 | 0.04934484 | -0.5459171 | 1 |
| PLD1 | 0.02826819 | -0.5463986 | 1 |
| MYOZ1 | 0.04563231 | -0.5465793 | 1 |
| SSNA1 | 0.00099908 | -0.5469003 | 1 |
| DZANK1 | 0.01294181 | -0.5502189 | 1 |
| COL4A1 | 0.00085284 | -0.5504404 | 1 |
| ATL2 | 0.0170851 | -0.5509658 | 1 |
| NEMP2 | 0.01391114 | -0.5509793 | 1 |
| ANXA1 | 0.00619148 | -0.5515214 | 1 |
| PLK2 | 0.03859691 | -0.55309 | 1 |
| ZKSCAN3 | 0.01308059 | -0.5533717 | 1 |
| CASP3 | 0.0220863 | -0.5545743 | 1 |
| CARF | 0.0193345 | -0.5547823 | 1 |
| TRIM23 | 0.02323801 | -0.5579171 | 1 |
| LRR1 | 0.04312282 | -0.5586249 | 1 |
| CDC42EP5 | 0.02206463 | -0.5616445 | 1 |
| PTS | 0.02967589 | -0.5620402 | 1 |
| ZNF132 | 0.01757794 | -0.5635471 | 1 |
| RBFA | 0.01604703 | -0.5639162 | 1 |
| RECQL4 | 0.0476628 | -0.5664371 | 1 |
| HDDC2 | 0.02055092 | -0.5666636 | 1 |
| AGER | 0.03912036 | -0.5668477 | 1 |
| CHPF2 | 0.01374688 | -0.5686554 | 1 |
| ADAL | 0.0068977 | -0.5694356 | 1 |
| RPP21 | 0.01516512 | -0.5697156 | 1 |
| ARSD | 0.01892266 | -0.5721626 | 1 |
| GRIP2 | 0.0113244 | -0.5730526 | 1 |

| | | | |
|---|---|---|---|
| POLB | 0.01680638 | -0.5747093 | 1 |
| NTNG2 | 0.03606899 | -0.574726 | 1 |
| CD151 | 0.03192891 | -0.5751745 | 1 |
| BST2 | 0.00016791 | -0.5758228 | 1 |
| ZNF512 | 0.02489288 | -0.578131 | 1 |
| PKD1 | 0.00630058 | -0.5783463 | 1 |
| MTHFSD | 0.03582631 | -0.5789045 | 1 |
| LRRC29 | 0.04748519 | -0.580009 | 1 |
| ALDH18A1 | 0.01922363 | -0.5809699 | 1 |
| BEND3 | 0.04956432 | -0.5815516 | 1 |
| CRB2 | 0.04065358 | -0.58216 | 1 |
| RAMP3 | 0.01626237 | -0.5829369 | 1 |
| FPGS | 0.00263532 | -0.5845606 | 1 |
| PCDHGA10 | 0.03482255 | -0.5862579 | 1 |
| ANO3 | 0.03915636 | -0.5883589 | 1 |
| PRSS27 | 0.04989882 | -0.5884614 | 1 |
| SLC35B4 | 0.03144508 | -0.58924 | 1 |
| ADRA1B | 0.0335424 | -0.5896947 | 1 |
| HOMER1 | 0.01944073 | -0.5898714 | 1 |
| SLC25A40 | 0.03822309 | -0.5906301 | 1 |
| TRAF6 | 0.04313972 | -0.5914835 | 1 |
| CNIH4 | 0.04726226 | -0.5924748 | 1 |
| PDLIM2 | 0.01537619 | -0.5926745 | 1 |
| SELENOW | 0.0164545 | -0.594832 | 1 |
| ZDBF2 | 0.04517873 | -0.5959242 | 1 |
| BAG4 | 0.03161125 | -0.5961878 | 1 |
| FOXI1 | 0.02255953 | -0.5962833 | 1 |
| PIAS3 | 0.01955707 | -0.5963542 | 1 |
| RBMS1 | 0.04750158 | -0.5964726 | 1 |
| MSMO1 | 0.02358964 | -0.5974403 | 1 |
| METTL5 | 0.01181507 | -0.5989151 | 1 |
| CACUL1 | 0.01903339 | -0.6038807 | 1 |
| FAM210B | 0.00629338 | -0.6042875 | 1 |
| NR1D1 | 0.02561154 | -0.6045106 | 1 |
| SAMD12 | 0.01120312 | -0.6057415 | 1 |
| AVL9 | 0.02643944 | -0.6064726 | 1 |
| ABCC9 | 0.03413079 | -0.6073503 | 1 |
| ITGAV | 0.00045532 | -0.6079025 | 1 |
| KIT | 0.03529232 | -0.6085836 | 1 |
| UNK | 0.01084617 | -0.6108562 | 1 |
| TERF1 | 0.04758406 | -0.6115808 | 1 |
| CLASP1 | 0.01931839 | -0.6119101 | 1 |
| BOK | 0.04337573 | -0.6123819 | 1 |
| HCFC1 | 0.02436768 | -0.6123976 | 1 |
| TUSC2 | 0.01316858 | -0.6125654 | 1 |
| SUPT7L | 0.0429648 | -0.6125955 | 1 |
| PRIM2 | 0.0137489 | -0.613131 | 1 |
| ADAMTS19 | 0.0103081 | -0.6143543 | 1 |
| LSG1 | 0.00762311 | -0.6149345 | 1 |
| PPP1CC | 0.03345413 | -0.615913 | 1 |
| PKIB | 0.03653922 | -0.6185985 | 1 |
| ARMC6 | 0.02010155 | -0.6191295 | 1 |
| CHMP1A | 0.04144159 | -0.6192183 | 1 |

| | | | |
|---|---|---|---|
| FHIT | 0.03582631 | -0.6201028 | 1 |
| NHSL2 | 0.03817282 | -0.6204302 | 1 |
| ZNF329 | 0.03292503 | -0.6210807 | 1 |
| DMAC2L | 0.02824305 | -0.6258588 | 1 |
| TBCD | 0.04372628 | -0.6261893 | 1 |
| SUFU | 0.00907374 | -0.6290391 | 1 |
| MPPE1 | 0.00722611 | -0.6311368 | 1 |
| FRS2 | 0.03499935 | -0.6354118 | 1 |
| VIPAS39 | 0.01080006 | -0.6355219 | 1 |
| NPHS1 | 0.00048959 | -0.646919 | 1 |
| A1CF | 0.04758922 | -0.6486703 | 1 |
| CDK2 | 0.04707792 | -0.648837 | 1 |
| MAPK6 | 0.02408588 | -0.6491448 | 1 |
| UPP2 | 0.04976882 | -0.6538126 | 1 |
| FKRP | 0.01443693 | -0.6542185 | 1 |
| NRXN1 | 0.03434359 | -0.6556768 | 1 |
| CAMKK1 | 0.02418665 | -0.6567277 | 1 |
| RRAGB | 0.0241426 | -0.658275 | 1 |
| BTRC | 0.00933326 | -0.6592089 | 1 |
| GPATCH4 | 0.01309718 | -0.6599554 | 1 |
| FAT1 | 0.04128969 | -0.6656181 | 1 |
| ZNF521 | 0.04925899 | -0.6665082 | 1 |
| ALS2CL | 0.00552984 | -0.6718199 | 1 |
| HMGA1 | 0.04956432 | -0.6735446 | 1 |
| HIST3H2BB | 0.03653922 | -0.6811408 | 1 |
| ZSCAN9 | 0.01471944 | -0.6846655 | 1 |
| PPIL1 | 0.02403515 | -0.6851638 | 1 |
| PRMT3 | 0.01513975 | -0.685352 | 1 |
| KIAA1841 | 0.01622918 | -0.6876178 | 1 |
| GIN1 | 0.02822204 | -0.6887172 | 1 |
| PTPRQ | 0.00498007 | -0.6919581 | 1 |
| TMEM71 | 0.011686 | -0.692777 | 1 |
| GLT8D1 | 0.00335698 | -0.6938093 | 1 |
| ZBTB48 | 0.00149293 | -0.694446 | 1 |
| MCU | 0.01507584 | -0.6951848 | 1 |
| TAL1 | 0.01360535 | -0.6976514 | 1 |
| ATL1 | 0.02255953 | -0.7011849 | 1 |
| SDC2 | 0.03280181 | -0.7021196 | 1 |
| PRAMEF18 | 0.03702779 | -0.7029022 | 1 |
| VTI1A | 0.01417007 | -0.7062683 | 1 |
| CNTROB | 0.03580701 | -0.7072917 | 1 |
| TOE1 | 0.00501247 | -0.7114671 | 1 |
| ZUP1 | 0.02353735 | -0.7155681 | 1 |
| MPP7 | 0.04339053 | -0.7175362 | 1 |
| BBC3 | 0.02917611 | -0.7178745 | 1 |
| MGAT5 | 0.02498836 | -0.7181178 | 1 |
| ABCA2 | 0.04957046 | -0.7211268 | 1 |
| CORO7 | 0.00285451 | -0.7228192 | 1 |
| ST7L | 0.04925899 | -0.7251556 | 1 |
| EIF4EBP1 | 0.04863075 | -0.7340744 | 1 |
| TCEAL2 | 0.02871218 | -0.7343141 | 1 |
| C1QTNF7 | 0.00045413 | -0.7349876 | 1 |
| ATXN7L3 | 0.00718947 | -0.7355565 | 1 |

| | | | |
|---|---|---|---|
| TFIP11 | 0.04778814 | -0.7356741 | 1 |
| NUDT8 | 0.02278001 | -0.7401157 | 1 |
| MB21D2 | 0.00155554 | -0.7426023 | 1 |
| PRORP | 0.02285454 | -0.7426738 | 1 |
| SFT2D3 | 0.04418466 | -0.749926 | 1 |
| POU5F1B | 0.04989882 | -0.7519501 | 1 |
| FICD | 0.00637482 | -0.7537126 | 1 |
| BST1 | 0.01754994 | -0.7541234 | 1 |
| FAM221A | 0.02780227 | -0.7546978 | 1 |
| TGFB3 | 0.03015039 | -0.7554762 | 1 |
| DDX39B | 0.02363907 | -0.7569277 | 1 |
| MOK | 0.04180234 | -0.7571839 | 1 |
| KDM5B | 0.03968907 | -0.7587977 | 1 |
| KLHL7 | 0.01641844 | -0.7620479 | 1 |
| MXRA8 | 0.00045242 | -0.7626025 | 1 |
| LNX1 | 0.04296744 | -0.7650365 | 1 |
| AFAP1L2 | 0.01266732 | -0.7663511 | 1 |
| CCRL2 | 0.04246389 | -0.768144 | 1 |
| WDR49 | 0.02153028 | -0.7695507 | 1 |
| CIAPIN1 | 0.01029655 | -0.7699642 | 1 |
| HSPB11 | 0.0442802 | -0.7739116 | 1 |
| HCCS | 0.02338709 | -0.7742292 | 1 |
| EFNB2 | 0.00080782 | -0.7793612 | 1 |
| RPAP3 | 0.01622771 | -0.7806382 | 1 |
| KCNK6 | 0.04444242 | -0.7809429 | 1 |
| ARIH2OS | 0.03921745 | -0.7817257 | 1 |
| CA4 | 0.00313093 | -0.7861504 | 1 |
| ZNF221 | 0.01503064 | -0.7878616 | 1 |
| INSIG2 | 0.03946803 | -0.7898922 | 1 |
| DNAJC18 | 0.01493504 | -0.7924729 | 1 |
| ZNF92 | 0.011686 | -0.7947866 | 1 |
| FMO4 | 0.04524933 | -0.795718 | 1 |
| DRICH1 | 0.03434359 | -0.800405 | 1 |
| HS6ST1 | 0.02886666 | -0.8009749 | 1 |
| HDHD2 | 0.04991283 | -0.8039759 | 1 |
| TGS1 | 0.00491239 | -0.8044928 | 1 |
| KLHL32 | 0.04989882 | -0.8087738 | 1 |
| DZIP1 | 0.03268222 | -0.8130184 | 1 |
| ECSIT | 0.03811774 | -0.8155578 | 1 |
| C18orf32 | 0.04839963 | -0.8157525 | 1 |
| NFASC | 0.00013844 | -0.8168238 | 1 |
| FOXJ2 | 0.00869411 | -0.8194957 | 1 |
| VPS29 | 0.01837361 | -0.8199224 | 1 |
| PRPF4 | 0.04608074 | -0.8319537 | 1 |
| KCNK3 | 0.03034474 | -0.8351186 | 1 |
| CENPV | 0.02473564 | -0.8355211 | 1 |
| PLAGL2 | 0.0151318 | -0.8362584 | 1 |
| ARHGAP33 | 0.04253634 | -0.8383992 | 1 |
| LONRF3 | 0.02612737 | -0.8416843 | 1 |
| LAS1L | 0.01499827 | -0.8428535 | 1 |
| FAM160A1 | 0.01977919 | -0.848756 | 1 |
| TMEM42 | 0.02564158 | -0.8540702 | 1 |
| HOXA13 | 0.0250906 | -0.8553395 | 1 |

| | | | |
|---|---|---|---|
| PPP2R3B | 0.01353254 | -0.8590431 | 1 |
| SEMA3B | 0.00740212 | -0.8672579 | 1 |
| EFCAB6 | 0.02418665 | -0.8711069 | 1 |
| FEZF2 | 0.04989882 | -0.8722957 | 1 |
| C12orf29 | 0.02123407 | -0.8751534 | 1 |
| SLX4 | 0.01011005 | -0.8759703 | 1 |
| TFPT | 0.00804058 | -0.8770082 | 1 |
| WARS2 | 0.04767375 | -0.8796405 | 1 |
| KIRREL1 | 9.00E-05 | -0.8802849 | 1 |
| CENPT | 0.029759 | -0.8824256 | 1 |
| PTRH2 | 0.04404061 | -0.8872252 | 1 |
| ALG6 | 0.01304572 | -0.899897 | 1 |
| C2orf69 | 0.04748519 | -0.9001119 | 1 |
| SCG5 | 0.04827278 | -0.9076173 | 1 |
| HES6 | 0.02510038 | -0.9100309 | 1 |
| BBIP1 | 0.0474021 | -0.912082 | 1 |
| LRIG3 | 0.01835229 | -0.9137191 | 1 |
| XRRA1 | 0.04564603 | -0.9186635 | 1 |
| ATPAF1 | 0.00982907 | -0.9221582 | 1 |
| MAGI2 | 0.00550303 | -0.9255686 | 1 |
| ST3GAL4 | 0.03996999 | -0.932906 | 1 |
| RCAN3 | 0.03369872 | -0.9346033 | 1 |
| THY1 | 0.04760208 | -0.9401793 | 1 |
| AREL1 | 0.00353285 | -0.9496224 | 1 |
| ZCWPW1 | 0.04989882 | -0.9499953 | 1 |
| NDUFAF2 | 0.04587211 | -0.9554189 | 1 |
| ZNF514 | 0.03671546 | -0.9575313 | 1 |
| RECQL | 0.03007671 | -0.9589628 | 1 |
| CLDN5 | 0.00129987 | -0.9592794 | 1 |
| PAM | 6.04E-05 | -0.9647252 | 1 |
| ZNF619 | 0.03194034 | -0.9680824 | 1 |
| NR2C1 | 0.03576438 | -0.9687869 | 1 |
| NCAPD2 | 0.00637097 | -0.9698504 | 1 |
| FAM47E | 0.01044258 | -0.9740796 | 1 |
| TPMT | 0.04670618 | -0.9743822 | 1 |
| POMGNT2 | 0.03606501 | -0.9766122 | 1 |
| FUT1 | 0.01278853 | -0.9799317 | 1 |
| FANCF | 0.03764547 | -0.9843107 | 1 |
| NAPB | 0.02364826 | -0.9869295 | 1 |
| ARHGAP39 | 0.03696971 | -0.9910356 | 1 |
| C1orf198 | 0.00109314 | -0.998073 | 1 |
| FADS3 | 0.01498138 | -1.0008803 | 1 |
| ST6GALNAC | 0.00033789 | -1.0084413 | 1 |
| ZNF771 | 0.0106725 | -1.0091652 | 1 |
| MRC2 | 0.00350068 | -1.0117248 | 1 |
| SNCA | 0.00204949 | -1.0200933 | 1 |
| PSMD5 | 0.04344199 | -1.0251485 | 1 |
| GCFC2 | 0.04087652 | -1.0400519 | 1 |
| ZNF214 | 0.03253755 | -1.048737 | 1 |
| SHBG | 0.01560068 | -1.0494524 | 1 |
| ZNF808 | 0.0188744 | -1.0657613 | 1 |
| LBH | 0.00421348 | -1.0695136 | 1 |
| CHPF | 0.04563524 | -1.0723631 | 1 |

| | | | |
|---|---|---|---|
| PPIP5K1 | 0.04366913 | -1.0794667 | 1 |
| TTC13 | 0.02501888 | -1.0906226 | 1 |
| ARL6IP6 | 0.03580701 | -1.1027504 | 1 |
| IPCEF1 | 0.02473564 | -1.1044967 | 1 |
| ITIH5 | 0.00264262 | -1.1056231 | 1 |
| THNSL1 | 0.00663777 | -1.114664 | 1 |
| GRPEL2 | 0.03678252 | -1.1283232 | 1 |
| WDR78 | 0.02197151 | -1.1288263 | 1 |
| ZNF439 | 0.03516586 | -1.1408312 | 1 |
| ZNF835 | 0.01532147 | -1.1453635 | 1 |
| CENPN | 0.02437644 | -1.1694221 | 1 |
| STOX1 | 0.03702779 | -1.1755659 | 1 |
| FGF9 | 0.0091381 | -1.1826814 | 1 |
| NXPH2 | 0.00873049 | -1.1837142 | 1 |
| LMX1B | 0.0329035 | -1.2216767 | 1 |
| MAFF | 0.01440805 | -1.2321918 | 1 |
| SNX25 | 0.00079999 | -1.2331643 | 1 |
| PMEPA1 | 0.00922888 | -1.2334062 | 1 |
| RCN1 | 0.01280578 | -1.2453566 | 1 |
| CDH13 | 0.02533139 | -1.2538898 | 1 |
| GCA | 0.00478481 | -1.3357003 | 1 |
| PLTP | 0.00104187 | -1.3498229 | 1 |
| SMAGP | 0.03263825 | -1.3532449 | 1 |
| MTMR11 | 0.02533153 | -1.3782708 | 1 |
| LONRF2 | 0.03354287 | -1.381759 | 1 |
| TSPAN2 | 0.00020576 | -1.3823581 | 1 |
| TMEM200C | 0.01347902 | -1.3852186 | 1 |
| SH3BGRL2 | 6.20E-05 | -1.4033628 | 1 |
| NBL1 | 0.01560577 | -1.4113528 | 1 |
| RHBDL2 | 0.03702779 | -1.5504182 | 1 |
| AEN | 0.01846008 | -1.6123032 | 1 |
| ENC1 | 0.00495771 | -1.7817356 | 1 |
| GEM | 0.00613053 | -1.7863196 | 1 |
| BMP2 | 0.00038803 | -1.8155744 | 1 |
| ADCYAP1 | 0.04333605 | -2.1689951 | 1 |
| CTSV | 0.00076021 | -2.2472848 | 1 |
| REN | 0.0065392 | -2.289998 | 1 |
| RGS2 | 0.04128292 | -2.4321457 | 1 |
| MMP15 | 0.04386713 | -2.5425897 | 1 |
| TPSD1 | 0.04411388 | -2.564934 | 1 |
| BHMG1 | 0.01332395 | -2.9440393 | 1 |
| DPEP2NB | 0.02542802 | -3.1758698 | 1 |
| FGB | 0.04742713 | -3.1938359 | 1 |
| FCER1A | 0.02542802 | -3.3348383 | 1 |
| POF1B | 0.02297356 | -3.3750721 | 1 |
| GPX5 | 0.04411388 | -3.3797387 | 1 |
| IFNK | 0.04411388 | -3.4574009 | 1 |
| SLC10A1 | 0.04411388 | -3.4900112 | 1 |
| LHFPL1 | 0.02297356 | -3.647631 | 1 |
| OR5K3 | 0.01332395 | -3.6789636 | 1 |
| RIT2 | 0.04411388 | -3.9041431 | 1 |
| SMR3B | 0.00414207 | -4.1995091 | 1 |
| GABRG2 | 0.01332395 | -4.6633035 | 1 |

### Supplemental Table S4

**Human Monocyte Derived Macrophage C5a Stimulation**

DESeq2 differential gene expression analysis comparing 24h C5a stimulation to control with patient and condition included in the model. Monocyte derived macrophages from N=3 independent healthy donors. Significantly differentially expressed genes defined as those with log2FC>|1|, padj <0.05.

| | log2FoldChange | pvalue | padj |
|---|---|---|---|
| NT5E | 7.316222134 | 2.77E-26 | 6.31E-25 |
| PLPP4 | 7.277964026 | 7.87E-20 | 1.21E-18 |
| MMP1 | 7.164766002 | 1.96E-26 | 4.50E-25 |
| PTPRD | 6.936096599 | 5.15E-07 | 2.56E-06 |
| IL1R2 | 6.840810278 | 2.30E-69 | 3.40E-67 |
| APCDD1L | 6.18990807 | 5.39E-08 | 3.01E-07 |
| CXCL5 | 5.884232658 | 1.59E-37 | 6.53E-36 |
| MMP3 | 5.75310413 | 8.75E-13 | 7.85E-12 |
| MMP8 | 5.509843126 | 7.84E-21 | 1.30E-19 |
| MMP10 | 5.504242541 | 8.39E-24 | 1.67E-22 |
| PAQR5 | 5.359799046 | 6.48E-26 | 1.45E-24 |
| TNFSF18 | 5.308939637 | 1.72E-05 | 6.92E-05 |
| GLDN | 5.1765409 | 2.37E-05 | 9.30E-05 |
| LRRC73 | 5.158159443 | 0.00071661 | 0.00217509 |
| LINC01989 | 5.135718218 | 0.00456301 | 0.01168528 |
| STEAP1 | 5.123896132 | 1.55E-44 | 8.83E-43 |
| TRIM9 | 5.108719459 | 1.71E-29 | 4.70E-28 |
| SERPINE1 | 5.011165171 | 1.57E-66 | 2.06E-64 |
| MEPE | 4.968021097 | 0.0005418 | 0.00168567 |
| CCL7 | 4.925256041 | 4.91E-45 | 2.90E-43 |
| CA12 | 4.910730646 | 3.71E-58 | 3.50E-56 |
| LPP-AS1 | 4.873629684 | 0.00361223 | 0.0094361 |
| LOC1019273 | 4.790308922 | 0.00459368 | 0.01174745 |
| EGR1 | 4.784246365 | 3.52E-14 | 3.56E-13 |
| MMP12 | 4.74505055 | 8.38E-22 | 1.46E-20 |
| IL1RN | 4.725599565 | 3.51E-28 | 8.90E-27 |
| INSYN2A | 4.711312519 | 0.00489002 | 0.01241693 |
| FABP4 | 4.689844382 | 1.41E-24 | 2.95E-23 |
| SYNPO2L | 4.657239183 | 0.00021209 | 0.00071314 |
| IL36RN | 4.642159261 | 1.68E-07 | 8.88E-07 |
| ALDH1A2 | 4.609280428 | 1.23E-15 | 1.40E-14 |
| IL1RL2 | 4.607636829 | 0.00014107 | 0.00048905 |

| | | | |
|---|---|---|---|
| RNF152 | 4.590056242 | 0.00268806 | 0.00723098 |
| FLT1 | 4.538508408 | 3.30E-21 | 5.61E-20 |
| NOTCH3 | 4.528323242 | 5.97E-191 | 8.98E-188 |
| REM1 | 4.455477244 | 1.02E-26 | 2.41E-25 |
| NDP | 4.434800054 | 8.85E-06 | 3.71E-05 |
| IL36B | 4.410867857 | 1.86E-17 | 2.45E-16 |
| VGF | 4.391198315 | 0.00025297 | 0.00083713 |
| ARNT2 | 4.360801662 | 1.08E-10 | 7.92E-10 |
| RGS16 | 4.331927255 | 5.35E-21 | 8.96E-20 |
| CCL20 | 4.31987758 | 5.74E-18 | 7.87E-17 |
| TMEM121 | 4.25579031 | 0.015043 | 0.0339948 |
| TCN1 | 4.226986649 | 3.50E-23 | 6.69E-22 |
| CCDC85A | 4.207351656 | 0.01437995 | 0.03267859 |
| MLXIPL | 4.145161014 | 5.27E-14 | 5.26E-13 |
| PRR16 | 4.126301721 | 9.17E-13 | 8.20E-12 |
| LINC02185 | 4.108467448 | 2.35E-05 | 9.25E-05 |
| CCL24 | 4.098506518 | 2.33E-54 | 1.99E-52 |
| TFAP2A | 4.081351678 | 1.22E-10 | 8.93E-10 |
| STEAP2 | 4.073596707 | 9.90E-08 | 5.38E-07 |
| LRTM2 | 4.072827012 | 0.01957635 | 0.04287605 |
| MT1JP | 4.050226595 | 2.54E-30 | 7.35E-29 |
| KCNJ15 | 4.035728873 | 3.56E-33 | 1.16E-31 |
| SPRED3 | 4.002047587 | 3.18E-08 | 1.82E-07 |
| FERMT1 | 3.987475098 | 1.82E-14 | 1.89E-13 |
| P4HA3 | 3.96514934 | 8.23E-26 | 1.84E-24 |
| GREM1 | 3.951586882 | 3.17E-11 | 2.45E-10 |
| BICDL2 | 3.870744561 | 0.00344828 | 0.00906077 |
| CPNE6 | 3.841763195 | 0.00355173 | 0.00929567 |
| TNC | 3.792528469 | 1.54E-17 | 2.05E-16 |
| LINC01629 | 3.78271374 | 4.81E-06 | 2.10E-05 |
| SLAMF9 | 3.774704383 | 1.55E-81 | 3.19E-79 |
| BTBD11 | 3.760115897 | 2.13E-23 | 4.12E-22 |
| SPP1 | 3.728805492 | 0.01356223 | 0.03100019 |
| CXCL8 | 3.725338379 | 9.14E-14 | 8.96E-13 |
| F5 | 3.720633675 | 4.99E-50 | 3.58E-48 |
| LINC02201 | 3.697945284 | 8.51E-41 | 4.15E-39 |
| MT1A | 3.697062461 | 1.69E-33 | 5.57E-32 |
| ADAMTS8 | 3.600438157 | 4.34E-06 | 1.90E-05 |
| GOLGA7B | 3.595533778 | 9.15E-41 | 4.45E-39 |
| VCAN | 3.569725817 | 1.61E-17 | 2.13E-16 |
| TIMP4 | 3.568666565 | 3.96E-11 | 3.03E-10 |
| AQP9 | 3.529859187 | 5.70E-34 | 1.92E-32 |
| EGR2 | 3.519420405 | 1.83E-18 | 2.61E-17 |

| | | | |
|---|---|---|---|
| CST6 | 3.515580642 | 0.00257955 | 0.00696289 |
| SHROOM4 | 3.500584326 | 5.90E-14 | 5.87E-13 |
| FLRT2 | 3.478388519 | 7.94E-61 | 8.20E-59 |
| LINC01258 | 3.455471608 | 0.01640065 | 0.03674146 |
| DKK2 | 3.442288246 | 5.59E-15 | 6.00E-14 |
| CACNA1G | 3.416009102 | 4.97E-19 | 7.31E-18 |
| SLC16A10 | 3.391590253 | 1.03E-21 | 1.80E-20 |
| CD1B | 3.359702997 | 1.84E-07 | 9.66E-07 |
| ADM2 | 3.349921442 | 5.18E-09 | 3.22E-08 |
| LOC1053780 | 3.328073562 | 0.00355029 | 0.00929337 |
| LBH | 3.314904639 | 1.63E-68 | 2.36E-66 |
| LINC01614 | 3.311207619 | 1.88E-06 | 8.64E-06 |
| EDN1 | 3.309202172 | 3.48E-06 | 1.54E-05 |
| PPBP | 3.307910659 | 7.18E-22 | 1.26E-20 |
| SRPX2 | 3.300833744 | 4.97E-41 | 2.47E-39 |
| FOXD1 | 3.29232552 | 0.01692474 | 0.03774676 |
| AQP3 | 3.290622745 | 1.93E-14 | 2.00E-13 |
| PHLDA1 | 3.28886221 | 7.20E-119 | 3.84E-116 |
| SLC28A3 | 3.269710618 | 6.43E-13 | 5.85E-12 |
| LINC02015 | 3.262241953 | 0.00176192 | 0.00492057 |
| SLC24A3 | 3.251703435 | 1.02E-11 | 8.29E-11 |
| CEMIP | 3.223589525 | 6.00E-37 | 2.38E-35 |
| ARMH1 | 3.223158358 | 2.70E-16 | 3.25E-15 |
| SLC39A8 | 3.211300391 | 5.21E-231 | 1.44E-227 |
| NBL1 | 3.205226136 | 1.12E-09 | 7.52E-09 |
| CLEC5A | 3.194404013 | 3.48E-27 | 8.37E-26 |
| CD1E | 3.192161631 | 3.78E-08 | 2.15E-07 |
| FN1 | 3.167821446 | 1.03E-43 | 5.59E-42 |
| MMP19 | 3.151150432 | 1.25E-141 | 1.14E-138 |
| HS3ST3A1 | 3.133791975 | 0.00859368 | 0.02061511 |
| HMGA2 | 3.122064172 | 1.05E-05 | 4.37E-05 |
| ZMIZ1-AS1 | 3.113545012 | 2.38E-67 | 3.33E-65 |
| RFX8 | 3.107275553 | 1.69E-16 | 2.07E-15 |
| LPL | 3.084638078 | 7.28E-23 | 1.36E-21 |
| SLC7A5 | 3.044479898 | 7.20E-87 | 1.73E-84 |
| ICAM5 | 3.043749512 | 4.21E-66 | 5.44E-64 |
| CH25H | 3.026769193 | 2.26E-09 | 1.46E-08 |
| LOC1019281 | 3.019585743 | 1.79E-17 | 2.37E-16 |
| CHAC1 | 2.989743097 | 2.60E-13 | 2.44E-12 |
| STX1A | 2.936824741 | 1.61E-61 | 1.72E-59 |
| MUCL1 | 2.927312587 | 6.49E-10 | 4.43E-09 |
| APOC1P1 | 2.924586847 | 5.62E-11 | 4.22E-10 |
| SLC7A11 | 2.919143833 | 1.98E-247 | 6.53E-244 |

| | | | |
|---|---|---|---|
| CXCL3 | 2.892646677 | 2.72E-16 | 3.26E-15 |
| COL1A1 | 2.888812704 | 0.00809572 | 0.019514 |
| CLDN14 | 2.867240319 | 1.68E-05 | 6.76E-05 |
| LAMA1 | 2.854844047 | 0.00062089 | 0.00190839 |
| IL1B | 2.83946635 | 2.48E-95 | 7.75E-93 |
| NCS1 | 2.824623698 | 1.45E-117 | 7.50E-115 |
| S1PR3 | 2.822285401 | 0.00067988 | 0.00207159 |
| MATK | 2.820411161 | 1.87E-125 | 1.19E-122 |
| HOMER1 | 2.817427184 | 1.36E-24 | 2.86E-23 |
| GPRACR | 2.808534538 | 0.01542135 | 0.03478324 |
| TLE1 | 2.794485651 | 2.48E-44 | 1.40E-42 |
| HSD11B1 | 2.792508919 | 7.33E-06 | 3.11E-05 |
| SGCG | 2.767368156 | 2.92E-05 | 0.00011265 |
| PSAT1 | 2.747341549 | 3.21E-35 | 1.14E-33 |
| EGR3 | 2.742409327 | 7.24E-08 | 3.99E-07 |
| APOC4-APO( | 2.736172018 | 1.92E-05 | 7.66E-05 |
| OLR1 | 2.72236349 | 2.81E-83 | 6.12E-81 |
| GLIS3 | 2.721453442 | 1.21E-106 | 4.99E-104 |
| NANOS3 | 2.720287853 | 0.02165996 | 0.04688454 |
| SPOCD1 | 2.695676434 | 9.99E-19 | 1.44E-17 |
| CXCL6 | 2.682016151 | 0.00016005 | 0.0005498 |
| SPINK1 | 2.675953732 | 0.00030279 | 0.00098816 |
| DUSP5 | 2.672237583 | 3.72E-36 | 1.41E-34 |
| UCN2 | 2.648536883 | 0.01526498 | 0.03445875 |
| PLS3 | 2.647878367 | 1.91E-34 | 6.56E-33 |
| TSPAN6 | 2.640477644 | 0.00213795 | 0.00586961 |
| MAMLD1 | 2.63205893 | 6.28E-32 | 1.96E-30 |
| LAMC2 | 2.631670997 | 0.00788949 | 0.01908927 |
| EREG | 2.629769134 | 1.82E-33 | 5.98E-32 |
| EBF1 | 2.608146174 | 0.00382363 | 0.00992871 |
| LINC01127 | 2.592996374 | 8.95E-30 | 2.50E-28 |
| MYOSLID | 2.592724891 | 5.72E-09 | 3.54E-08 |
| MT1L | 2.588988704 | 7.72E-54 | 6.38E-52 |
| BCL2A1 | 2.585093789 | 1.87E-21 | 3.21E-20 |
| TNFRSF12A | 2.575112096 | 6.71E-108 | 2.85E-105 |
| ZFYVE9 | 2.564835183 | 4.05E-14 | 4.07E-13 |
| C2orf72 | 2.557744203 | 0.00029098 | 0.00095299 |
| CXXC5-AS1 | 2.545728584 | 0.00092034 | 0.0027264 |
| TPST1 | 2.544202959 | 4.13E-36 | 1.57E-34 |
| SPINK6 | 2.540639412 | 8.45E-05 | 0.00030306 |
| TAFA3 | 2.51257223 | 0.00115462 | 0.00334666 |
| BMP6 | 2.507132294 | 0.00835003 | 0.02008306 |
| MT1M | 2.49002346 | 2.03E-17 | 2.66E-16 |

| | | | |
|---|---|---|---|
| HEY1 | 2.487663771 | 0.00192384 | 0.00533672 |
| PTPRF | 2.482814953 | 2.16E-05 | 8.56E-05 |
| SCN7A | 2.47865386 | 0.00721198 | 0.01760195 |
| PCDHGC3 | 2.475306334 | 3.30E-68 | 4.71E-66 |
| TFPI2 | 2.470391022 | 0.00825554 | 0.01987316 |
| MAS1 | 2.455241047 | 7.76E-05 | 0.0002799 |
| PPP1R3C | 2.452283184 | 0.00662411 | 0.01631894 |
| ST20 | 2.425531793 | 9.07E-11 | 6.69E-10 |
| CAVIN1 | 2.41724799 | 7.20E-23 | 1.35E-21 |
| TMEM45A | 2.412715672 | 4.07E-15 | 4.41E-14 |
| APOC1 | 2.404043386 | 5.04E-121 | 2.87E-118 |
| MT2A | 2.403210758 | 5.34E-12 | 4.48E-11 |
| RAI14 | 2.387146831 | 9.70E-14 | 9.48E-13 |
| CCL4L2 | 2.385238359 | 7.55E-11 | 5.60E-10 |
| RGMA | 2.378070427 | 0.00196074 | 0.0054245 |
| CDKN2B | 2.374146746 | 6.50E-34 | 2.17E-32 |
| PPP1R14C | 2.371139105 | 3.50E-10 | 2.45E-09 |
| MELTF | 2.368486425 | 2.34E-158 | 2.42E-155 |
| GEM | 2.362563965 | 1.44E-06 | 6.76E-06 |
| KANK1 | 2.344830978 | 8.70E-15 | 9.23E-14 |
| STXBP5-AS1 | 2.344737837 | 2.98E-25 | 6.50E-24 |
| NRIP3 | 2.344556317 | 1.65E-79 | 3.29E-77 |
| SDC1 | 2.332545838 | 7.70E-28 | 1.92E-26 |
| ADAM12 | 2.330588282 | 1.60E-49 | 1.12E-47 |
| PTX3 | 2.319833904 | 2.44E-77 | 4.38E-75 |
| SLC1A2 | 2.305350852 | 0.00112757 | 0.00327628 |
| ENO2 | 2.303719207 | 1.59E-10 | 1.15E-09 |
| C6orf132 | 2.279968106 | 0.00037352 | 0.0011984 |
| KCNN4 | 2.274652024 | 6.13E-08 | 3.41E-07 |
| SYN2 | 2.274039431 | 0.00679233 | 0.0166762 |
| FOSL1 | 2.27274996 | 4.61E-32 | 1.45E-30 |
| FBP1 | 2.262776848 | 1.19E-53 | 9.78E-52 |
| SLAMF1 | 2.259931132 | 4.68E-08 | 2.64E-07 |
| F3 | 2.239874456 | 3.81E-14 | 3.84E-13 |
| MYOZ1 | 2.23606259 | 3.69E-05 | 0.00014028 |
| MITF | 2.230216088 | 4.47E-95 | 1.37E-92 |
| IL7R | 2.228526342 | 4.58E-54 | 3.85E-52 |
| SLC7A11-AS | 2.218130395 | 0.00784378 | 0.01899534 |
| TNFAIP6 | 2.214763402 | 2.47E-31 | 7.49E-30 |
| SEMA3C | 2.214717838 | 4.52E-101 | 1.70E-98 |
| PYCR1 | 2.21366144 | 3.60E-13 | 3.33E-12 |
| ENPP2 | 2.210209669 | 6.14E-08 | 3.41E-07 |
| CAVIN4 | 2.209608635 | 0.00062374 | 0.00191535 |

| | | | |
|---|---|---|---|
| LGI2 | 2.208393189 | 1.25E-62 | 1.41E-60 |
| MPP6 | 2.206794755 | 2.95E-26 | 6.70E-25 |
| CXCL1 | 2.201972664 | 1.95E-07 | 1.02E-06 |
| SAMSN1-AS1 | 2.201950237 | 0.00750996 | 0.01825914 |
| MME | 2.200614075 | 1.12E-14 | 1.18E-13 |
| CNIH3 | 2.185776018 | 7.28E-58 | 6.80E-56 |
| LAT | 2.183240761 | 8.61E-70 | 1.29E-67 |
| TPRG1 | 2.180863302 | 2.61E-27 | 6.32E-26 |
| FERMT2 | 2.180558937 | 6.17E-12 | 5.15E-11 |
| SLC38A5 | 2.175800385 | 1.40E-19 | 2.12E-18 |
| LPAR3 | 2.175174 | 1.54E-07 | 8.15E-07 |
| PHLDA2 | 2.170603477 | 1.74E-35 | 6.34E-34 |
| MET | 2.169607675 | 2.46E-36 | 9.45E-35 |
| SGIP1 | 2.166350955 | 4.83E-06 | 2.10E-05 |
| ACOD1 | 2.160960574 | 0.00032062 | 0.00104286 |
| MGAM | 2.15579411 | 1.11E-46 | 7.00E-45 |
| ETV4 | 2.146799523 | 0.00034939 | 0.0011271 |
| CSPG4 | 2.146408221 | 2.57E-22 | 4.62E-21 |
| ENAH | 2.145749456 | 4.21E-33 | 1.36E-31 |
| TGM2 | 2.14259613 | 1.80E-51 | 1.36E-49 |
| DNER | 2.135431227 | 0.00038421 | 0.00122985 |
| TMEM52B | 2.134505682 | 4.97E-35 | 1.76E-33 |
| FABP5 | 2.126699399 | 9.42E-62 | 1.02E-59 |
| ITGB7 | 2.125825431 | 5.56E-34 | 1.88E-32 |
| MT1G | 2.124585915 | 6.76E-11 | 5.04E-10 |
| TMEM145 | 2.124227994 | 0.0103278 | 0.02431636 |
| FCRLA | 2.101171397 | 1.49E-11 | 1.19E-10 |
| ANKRD22 | 2.089767578 | 5.07E-09 | 3.16E-08 |
| HMGN2P46 | 2.085116154 | 4.72E-06 | 2.06E-05 |
| CDK14 | 2.058038242 | 3.93E-91 | 1.08E-88 |
| NRCAM | 2.057921485 | 0.00248221 | 0.00672541 |
| ANPEP | 2.055352734 | 6.32E-120 | 3.49E-117 |
| NTSR1 | 2.049445271 | 5.77E-17 | 7.30E-16 |
| PLOD2 | 2.049137745 | 6.72E-29 | 1.79E-27 |
| TREM1 | 2.044187753 | 5.48E-54 | 4.58E-52 |
| IGLON5 | 2.037172397 | 7.72E-07 | 3.75E-06 |
| SPSB1 | 2.032375327 | 3.49E-28 | 8.86E-27 |
| PDGFA | 2.025374923 | 4.31E-10 | 3.00E-09 |
| DCSTAMP | 2.011420177 | 3.02E-07 | 1.55E-06 |
| ASPHD1 | 2.00892311 | 3.43E-06 | 1.52E-05 |
| DPP4 | 2.000802782 | 1.57E-55 | 1.38E-53 |
| NAB2 | 1.998476661 | 2.93E-99 | 1.05E-96 |
| SYNJ2 | 1.995038002 | 3.96E-32 | 1.25E-30 |

| | | | |
|---|---|---|---|
| TMEM132A | 1.990898033 | 3.83E-31 | 1.15E-29 |
| TMEM198 | 1.979430742 | 0.01529578 | 0.03452357 |
| MT1E | 1.977785716 | 1.14E-09 | 7.59E-09 |
| CGREF1 | 1.975735897 | 0.00167829 | 0.00470689 |
| PIF1 | 1.971081447 | 9.15E-14 | 8.97E-13 |
| CLCF1 | 1.967956682 | 6.21E-09 | 3.83E-08 |
| RAC3 | 1.962909565 | 1.87E-15 | 2.09E-14 |
| IL1R1 | 1.958457914 | 1.09E-58 | 1.06E-56 |
| MT1X | 1.956691015 | 1.30E-07 | 6.98E-07 |
| DOCK3 | 1.953802747 | 3.90E-12 | 3.31E-11 |
| MYOF | 1.950949673 | 5.38E-86 | 1.27E-83 |
| FGFRL1 | 1.939250812 | 9.44E-45 | 5.51E-43 |
| DPYSL4 | 1.935919794 | 2.60E-11 | 2.02E-10 |
| ARAP3 | 1.929120545 | 7.00E-24 | 1.40E-22 |
| PROCR | 1.922208777 | 8.07E-37 | 3.16E-35 |
| C4orf47 | 1.921089564 | 0.01579871 | 0.03553258 |
| SCIN | 1.91872484 | 5.23E-22 | 9.25E-21 |
| RNF165 | 1.916136787 | 0.00041953 | 0.00133412 |
| LUCAT1 | 1.910526455 | 5.95E-25 | 1.28E-23 |
| CYP27B1 | 1.899949342 | 5.74E-10 | 3.94E-09 |
| ZNF462 | 1.898102428 | 2.17E-24 | 4.50E-23 |
| TUBB3 | 1.892515737 | 1.69E-07 | 8.95E-07 |
| CLEC12B | 1.889909374 | 0.00127226 | 0.00365941 |
| HPDL | 1.885007992 | 0.00034489 | 0.001115 |
| CHST6 | 1.884775657 | 0.01421173 | 0.03235506 |
| FJX1 | 1.876661584 | 6.62E-37 | 2.61E-35 |
| CFAP58-DT | 1.875504766 | 2.97E-17 | 3.87E-16 |
| DCLK2 | 1.874520448 | 0.00488651 | 0.0124114 |
| RND3 | 1.873466153 | 2.12E-20 | 3.40E-19 |
| SDC2 | 1.866269732 | 6.03E-84 | 1.35E-81 |
| LINC00346 | 1.854308542 | 6.44E-12 | 5.36E-11 |
| OLIG2 | 1.854113978 | 0.00028041 | 0.00092003 |
| MYO1B | 1.852761048 | 7.62E-39 | 3.33E-37 |
| RBFOX2 | 1.852405835 | 4.22E-12 | 3.58E-11 |
| PTPRN | 1.837812031 | 0.01548995 | 0.03491891 |
| LINC00937 | 1.834380778 | 1.38E-05 | 5.62E-05 |
| MMP7 | 1.833730371 | 1.62E-06 | 7.52E-06 |
| IRAK2 | 1.82743326 | 5.81E-38 | 2.42E-36 |
| MSC | 1.823431183 | 1.89E-25 | 4.19E-24 |
| LOC1019284 | 1.821969563 | 1.14E-11 | 9.25E-11 |
| BHLHE41 | 1.818544232 | 2.98E-25 | 6.50E-24 |
| G0S2 | 1.81842771 | 1.04E-08 | 6.30E-08 |
| EHD2 | 1.816660805 | 8.31E-17 | 1.04E-15 |

| | | | |
|---|---|---|---|
| ALCAM | 1.816483426 | 3.35E-113 | 1.58E-110 |
| ARG2 | 1.809215743 | 1.27E-16 | 1.57E-15 |
| ATP13A3 | 1.8076732 | 4.63E-70 | 7.09E-68 |
| SLC7A1 | 1.798625684 | 5.36E-67 | 7.21E-65 |
| DLGAP1-AS2 | 1.7948568 | 1.13E-06 | 5.38E-06 |
| A4GALT | 1.790014421 | 0.00524111 | 0.01322517 |
| TNFRSF11A | 1.786232229 | 2.30E-64 | 2.77E-62 |
| ST6GALNAC4 | 1.780729883 | 8.38E-31 | 2.46E-29 |
| NIM1K | 1.777596142 | 5.07E-05 | 0.00018857 |
| ADGRE2 | 1.773438257 | 2.47E-88 | 6.20E-86 |
| CLEC6A | 1.772413916 | 6.10E-05 | 0.00022359 |
| TEAD4 | 1.771445109 | 0.00598584 | 0.0149042 |
| LRP12 | 1.768338243 | 5.45E-58 | 5.12E-56 |
| MT1H | 1.76676448 | 1.39E-09 | 9.17E-09 |
| METTL1 | 1.764816145 | 4.36E-43 | 2.31E-41 |
| PLXNB3 | 1.760884452 | 5.29E-19 | 7.76E-18 |
| GLB1L2 | 1.759341172 | 4.65E-05 | 0.00017414 |
| LINC00884 | 1.757468398 | 6.07E-11 | 4.54E-10 |
| CNKSR3 | 1.753711803 | 1.01E-07 | 5.48E-07 |
| CDS1 | 1.752409684 | 5.51E-23 | 1.04E-21 |
| RAPGEFL1 | 1.752186581 | 2.78E-13 | 2.59E-12 |
| MREG | 1.750963754 | 6.54E-66 | 8.38E-64 |
| MICALL2 | 1.740177601 | 1.83E-79 | 3.60E-77 |
| RAB7B | 1.735865962 | 6.84E-98 | 2.31E-95 |
| FYN | 1.733423474 | 3.72E-25 | 8.07E-24 |
| PPM1J | 1.732185823 | 0.00138188 | 0.0039418 |
| STC2 | 1.725830912 | 4.53E-05 | 0.00017014 |
| CD109 | 1.71837609 | 2.92E-12 | 2.50E-11 |
| LOC1053783 | 1.717043185 | 2.32E-17 | 3.04E-16 |
| SOCS3 | 1.713725677 | 1.28E-06 | 6.02E-06 |
| LRP6 | 1.710693743 | 5.94E-11 | 4.45E-10 |
| AK4 | 1.71051755 | 1.82E-08 | 1.08E-07 |
| TMEM45B | 1.708095304 | 6.84E-09 | 4.21E-08 |
| C1orf21 | 1.705811075 | 4.40E-148 | 4.28E-145 |
| PDE2A | 1.695801929 | 1.55E-13 | 1.48E-12 |
| LINC01605 | 1.693204631 | 4.71E-05 | 0.00017584 |
| CCL3L1 | 1.6877898 | 1.36E-06 | 6.42E-06 |
| LINC01010 | 1.685823965 | 4.54E-07 | 2.28E-06 |
| RFLNB | 1.677026252 | 2.55E-19 | 3.80E-18 |
| TFR2 | 1.664795071 | 1.82E-08 | 1.08E-07 |
| PITPNM2 | 1.663244319 | 4.45E-11 | 3.38E-10 |
| TNFAIP8L3 | 1.650820816 | 2.57E-06 | 1.16E-05 |
| SLCO4A1 | 1.648096341 | 5.07E-16 | 5.97E-15 |

| | | | |
|---|---|---|---|
| CHST2 | 1.646551755 | 2.11E-101 | 8.13E-99 |
| PLAUR | 1.646043254 | 4.27E-38 | 1.79E-36 |
| MT1F | 1.644865765 | 1.06E-41 | 5.40E-40 |
| VAV3-AS1 | 1.638094861 | 0.00094795 | 0.00280216 |
| SLC12A8 | 1.637176862 | 2.70E-09 | 1.74E-08 |
| IL1A | 1.636830696 | 4.51E-05 | 0.00016918 |
| C1orf122 | 1.635750581 | 1.47E-94 | 4.42E-92 |
| TGM3 | 1.635685091 | 0.01472553 | 0.0333627 |
| CXCL2 | 1.632259336 | 2.20E-24 | 4.55E-23 |
| ANKRD13B | 1.624208047 | 0.00033128 | 0.00107436 |
| RGCC | 1.620978271 | 3.93E-24 | 8.06E-23 |
| CKAP4 | 1.613164577 | 6.38E-105 | 2.57E-102 |
| LINC00607 | 1.607284457 | 0.00566151 | 0.01417629 |
| SPHK1 | 1.607110296 | 1.65E-29 | 4.54E-28 |
| DYRK3 | 1.601339552 | 2.83E-14 | 2.88E-13 |
| SHOX2 | 1.59833136 | 0.00075096 | 0.00226643 |
| TRAF3IP2 | 1.595597476 | 9.16E-25 | 1.93E-23 |
| CHPF | 1.592386347 | 2.65E-47 | 1.72E-45 |
| TNIP3 | 1.589336871 | 1.10E-35 | 4.02E-34 |
| UNC5A | 1.584761243 | 4.31E-06 | 1.89E-05 |
| KALRN | 1.570304126 | 2.15E-07 | 1.12E-06 |
| PCDHGA12 | 1.569702123 | 1.59E-05 | 6.43E-05 |
| LOC1001280 | 1.566414861 | 0.00013705 | 0.00047603 |
| DNAJC6 | 1.564195767 | 1.43E-06 | 6.70E-06 |
| ITGB8 | 1.563979354 | 6.87E-40 | 3.20E-38 |
| NKD1 | 1.563052697 | 1.49E-06 | 6.96E-06 |
| IGFBP6 | 1.558097038 | 0.00502705 | 0.01273166 |
| CYTIP | 1.558067584 | 2.61E-25 | 5.73E-24 |
| CSF1 | 1.554927614 | 0.00855413 | 0.02053812 |
| LINC01181 | 1.547599876 | 5.05E-17 | 6.43E-16 |
| LINC01679 | 1.540909981 | 0.00177698 | 0.0049593 |
| NME1 | 1.540652417 | 6.81E-66 | 8.66E-64 |
| TM4SF19 | 1.539472258 | 2.91E-08 | 1.68E-07 |
| COL7A1 | 1.526688661 | 0.00056634 | 0.00175605 |
| ARHGAP20 | 1.523698423 | 5.85E-09 | 3.61E-08 |
| ATP2B1 | 1.521124396 | 6.26E-56 | 5.63E-54 |
| PLEK2 | 1.518223532 | 5.43E-13 | 4.97E-12 |
| C13orf46 | 1.516414339 | 2.41E-06 | 1.10E-05 |
| LINC02345 | 1.508716873 | 0.00396069 | 0.01026365 |
| CLEC4E | 1.508457302 | 1.57E-14 | 1.63E-13 |
| SYDE1 | 1.496763113 | 0.01122218 | 0.0261503 |
| PHLDB1 | 1.494932853 | 3.58E-52 | 2.78E-50 |
| CCDC103 | 1.49260636 | 2.09E-09 | 1.36E-08 |

| | | | |
|---|---|---|---|
| LAMA3 | 1.481937345 | 1.01E-11 | 8.21E-11 |
| MSANTD3 | 1.48105963 | 1.04E-50 | 7.72E-49 |
| DPYSL3 | 1.481006271 | 1.24E-05 | 5.11E-05 |
| TDO2 | 1.476837054 | 2.16E-05 | 8.54E-05 |
| FAM20C | 1.472947804 | 6.63E-37 | 2.61E-35 |
| PPARG | 1.471180644 | 7.01E-16 | 8.16E-15 |
| PTPN7 | 1.470385511 | 1.36E-33 | 4.49E-32 |
| FGD5 | 1.467317127 | 9.57E-19 | 1.38E-17 |
| SCGB1A1 | 1.465643588 | 0.01696747 | 0.03781146 |
| MANEAL | 1.464285556 | 2.04E-33 | 6.67E-32 |
| LIF | 1.462811253 | 1.87E-11 | 1.47E-10 |
| ELOVL7 | 1.459633438 | 5.25E-07 | 2.61E-06 |
| GLUD1P3 | 1.45466421 | 1.13E-08 | 6.79E-08 |
| ADAMTS14 | 1.454494739 | 1.04E-13 | 1.01E-12 |
| WDR66 | 1.454405605 | 3.38E-06 | 1.50E-05 |
| CYP19A1 | 1.453308871 | 0.00473784 | 0.01206944 |
| TNKS1BP1 | 1.449732658 | 2.60E-37 | 1.05E-35 |
| LTBP2 | 1.447689403 | 1.98E-46 | 1.25E-44 |
| ZDHHC9 | 1.446105075 | 1.33E-51 | 1.02E-49 |
| PLXNB1 | 1.444942238 | 0.00078585 | 0.00236142 |
| NETO2 | 1.442508845 | 3.24E-26 | 7.33E-25 |
| VMO1 | 1.440093463 | 2.70E-12 | 2.32E-11 |
| LOC399975 | 1.438493667 | 4.34E-05 | 0.00016341 |
| VWA1 | 1.430194072 | 1.61E-11 | 1.27E-10 |
| PHLDA3 | 1.424572433 | 2.27E-37 | 9.19E-36 |
| TJP1 | 1.424334974 | 8.00E-07 | 3.87E-06 |
| AOX1 | 1.423880298 | 0.00813272 | 0.01959747 |
| USP2 | 1.423855415 | 0.00027956 | 0.00091754 |
| IPO4 | 1.423009998 | 8.55E-99 | 3.01E-96 |
| DNAJB5 | 1.422676215 | 3.15E-42 | 1.63E-40 |
| MIR17HG | 1.414948983 | 0.00014853 | 0.00051256 |
| GLIPR2 | 1.413901714 | 4.68E-07 | 2.34E-06 |
| PAPSS2 | 1.412617402 | 1.73E-23 | 3.37E-22 |
| DIXDC1 | 1.411866443 | 1.48E-06 | 6.93E-06 |
| KIF26A | 1.410109437 | 0.00148654 | 0.004212 |
| SLC27A3 | 1.4028184 | 1.95E-26 | 4.49E-25 |
| KAZN | 1.402740465 | 0.001853 | 0.00515318 |
| RTN4RL2 | 1.401929695 | 1.51E-06 | 7.06E-06 |
| APBB2 | 1.401689089 | 2.79E-09 | 1.78E-08 |
| TUBB6 | 1.398408013 | 3.57E-59 | 3.50E-57 |
| CD300LB | 1.398025474 | 3.48E-10 | 2.44E-09 |
| ZC3HAV1L | 1.396510781 | 4.77E-17 | 6.11E-16 |
| LOC1053733 | 1.391895151 | 2.85E-05 | 0.00011028 |

| | | | |
|---|---|---|---|
| LINC00520 | 1.391226902 | 0.00063964 | 0.00196019 |
| SLC35F2 | 1.389962843 | 6.29E-16 | 7.35E-15 |
| TCTEX1D4 | 1.388004528 | 0.00061187 | 0.00188347 |
| CD44 | 1.384119128 | 1.33E-44 | 7.63E-43 |
| LOC1002894 | 1.381586118 | 0.01293338 | 0.02967355 |
| PCOLCE2 | 1.380043288 | 0.00850482 | 0.02042268 |
| GGN | 1.37935342 | 0.01464519 | 0.03321384 |
| FGR | 1.374115947 | 1.33E-41 | 6.71E-40 |
| SH3BGRL3 | 1.370816877 | 7.95E-114 | 3.87E-111 |
| PVT1 | 1.369593849 | 1.11E-15 | 1.26E-14 |
| AK1 | 1.36684459 | 3.79E-14 | 3.82E-13 |
| YRDC | 1.365572374 | 5.41E-28 | 1.36E-26 |
| PALM2AKAP | 1.36275691 | 1.86E-40 | 8.90E-39 |
| CD276 | 1.359735643 | 8.91E-45 | 5.24E-43 |
| IL3RA | 1.356639239 | 1.85E-05 | 7.39E-05 |
| MTCL1 | 1.350577681 | 0.00171689 | 0.00480456 |
| B4GALT2 | 1.349663328 | 3.29E-37 | 1.32E-35 |
| PAICS | 1.349533121 | 1.93E-47 | 1.26E-45 |
| SEMA5A | 1.347870252 | 0.00056856 | 0.0017613 |
| SNORD139 | 1.347029509 | 0.00588156 | 0.01467328 |
| ZSWIM4 | 1.34448863 | 9.24E-30 | 2.58E-28 |
| WHRN | 1.344102345 | 1.63E-09 | 1.07E-08 |
| MAFF | 1.34343005 | 8.16E-13 | 7.34E-12 |
| BNIP3 | 1.34254408 | 2.43E-31 | 7.37E-30 |
| CLBA1 | 1.340311018 | 9.29E-17 | 1.16E-15 |
| CCN3 | 1.332840803 | 3.35E-11 | 2.57E-10 |
| ARFGEF3 | 1.331773312 | 6.67E-08 | 3.69E-07 |
| LOC388813 | 1.330627757 | 4.08E-08 | 2.31E-07 |
| NIBAN2 | 1.329332127 | 9.15E-45 | 5.37E-43 |
| E2F5 | 1.328432557 | 9.78E-05 | 0.00034809 |
| SGMS2 | 1.326439028 | 3.60E-39 | 1.61E-37 |
| C15orf48 | 1.324822959 | 1.74E-39 | 7.94E-38 |
| SRM | 1.323786954 | 5.53E-91 | 1.50E-88 |
| CTSL | 1.323658092 | 5.60E-50 | 4.00E-48 |
| PNP | 1.318687504 | 3.40E-27 | 8.19E-26 |
| APOE | 1.318041368 | 6.17E-23 | 1.16E-21 |
| MMP2-AS1 | 1.316414455 | 6.50E-10 | 4.44E-09 |
| PRKAR1B-AS | 1.314909862 | 2.40E-13 | 2.25E-12 |
| SSTR2 | 1.314868347 | 0.00167711 | 0.0047044 |
| CHST15 | 1.311276835 | 2.99E-08 | 1.72E-07 |
| GXYLT2 | 1.310649069 | 0.01157096 | 0.0268495 |
| FOS | 1.309168359 | 4.94E-20 | 7.74E-19 |
| PDLIM7 | 1.308810734 | 2.78E-54 | 2.36E-52 |

| | | | |
|---|---|---|---|
| GPR3 | 1.308671429 | 0.0003622 | 0.00116526 |
| THBD | 1.307722624 | 3.47E-06 | 1.54E-05 |
| GSN | 1.306225937 | 1.23E-83 | 2.71E-81 |
| NOP16 | 1.305113646 | 6.95E-38 | 2.87E-36 |
| ADAMTS2 | 1.304228009 | 5.00E-05 | 0.00018627 |
| PPFIA4 | 1.30221168 | 1.41E-08 | 8.41E-08 |
| TMEM38B | 1.300606216 | 4.49E-27 | 1.08E-25 |
| MYEOV | 1.298101024 | 1.60E-05 | 6.47E-05 |
| SNORD17 | 1.297792889 | 0.00019325 | 0.00065456 |
| MYC | 1.295804804 | 9.50E-62 | 1.02E-59 |
| CHRNA5 | 1.293689582 | 0.00290425 | 0.00775201 |
| NEK10 | 1.290980719 | 0.00368257 | 0.00960617 |
| NFKBID | 1.290648284 | 1.08E-38 | 4.69E-37 |
| SMPDL3A | 1.289834063 | 1.79E-40 | 8.61E-39 |
| ZNF239 | 1.289547804 | 0.01574523 | 0.03543993 |
| MARCKSL1 | 1.285192951 | 2.10E-11 | 1.65E-10 |
| CDK20 | 1.281548017 | 6.55E-20 | 1.02E-18 |
| GFOD1 | 1.279508174 | 1.77E-26 | 4.11E-25 |
| WDR11 | 1.278910979 | 2.66E-87 | 6.47E-85 |
| GDF15 | 1.275214832 | 9.42E-16 | 1.08E-14 |
| SHANK3 | 1.274837446 | 0.01013251 | 0.02391725 |
| HMGA1 | 1.274492023 | 2.09E-73 | 3.52E-71 |
| CCDC102B | 1.273750844 | 3.80E-06 | 1.68E-05 |
| BCAR1 | 1.271980379 | 2.56E-18 | 3.61E-17 |
| FAM83G | 1.270461684 | 1.76E-24 | 3.66E-23 |
| GK | 1.269903241 | 6.86E-18 | 9.33E-17 |
| FAM124B | 1.266336195 | 1.76E-07 | 9.28E-07 |
| FAM216A | 1.26530446 | 4.20E-18 | 5.81E-17 |
| SHC3 | 1.2641177 | 0.00161952 | 0.00455874 |
| SH3PXD2B | 1.264091221 | 1.97E-56 | 1.79E-54 |
| CTPS1 | 1.263296541 | 1.78E-30 | 5.18E-29 |
| FAM124A | 1.261511472 | 0.00010335 | 0.00036574 |
| EFR3B | 1.257714969 | 0.00036998 | 0.00118845 |
| ANKRD28 | 1.253214581 | 1.77E-35 | 6.40E-34 |
| MTHFD1L | 1.2512545 | 1.47E-53 | 1.20E-51 |
| MLLT11 | 1.251182916 | 2.17E-07 | 1.13E-06 |
| NRARP | 1.248547138 | 0.01858206 | 0.04098962 |
| LINC02709 | 1.248095837 | 0.00941423 | 0.02238214 |
| DUSP4 | 1.248044451 | 8.00E-09 | 4.90E-08 |
| LOC1053744 | 1.247794445 | 4.83E-07 | 2.41E-06 |
| KHDRBS3 | 1.247296508 | 5.68E-06 | 2.45E-05 |
| CXXC5 | 1.246756265 | 6.58E-29 | 1.75E-27 |
| SLC4A7 | 1.246648827 | 8.12E-80 | 1.64E-77 |

| | | | |
|---|---|---|---|
| EMILIN1 | 1.24630898 | 6.35E-48 | 4.23E-46 |
| MAP4K3-DT | 1.24460321 | 4.71E-18 | 6.49E-17 |
| ARHGAP39 | 1.244364189 | 0.00158765 | 0.00447318 |
| DDX21 | 1.243341687 | 2.36E-49 | 1.65E-47 |
| ZNF697 | 1.243002481 | 2.71E-20 | 4.33E-19 |
| TNFRSF18 | 1.242071025 | 0.00164039 | 0.00460938 |
| SLC20A1 | 1.24039972 | 4.21E-31 | 1.25E-29 |
| APLN | 1.239248232 | 2.09E-07 | 1.09E-06 |
| SLC25A22 | 1.23287359 | 4.96E-39 | 2.20E-37 |
| UPP1 | 1.231122836 | 1.65E-28 | 4.29E-27 |
| TWNK | 1.224267588 | 2.28E-37 | 9.22E-36 |
| TOMM40 | 1.223360343 | 3.18E-36 | 1.22E-34 |
| GPR68 | 1.221949236 | 1.74E-26 | 4.05E-25 |
| TMEM158 | 1.220997418 | 0.00014814 | 0.00051133 |
| ULBP1 | 1.220485893 | 0.01737667 | 0.03860885 |
| ZMYND15 | 1.218402849 | 1.18E-11 | 9.52E-11 |
| ME3 | 1.217805338 | 9.65E-10 | 6.50E-09 |
| LINC01599 | 1.21776567 | 0.00032025 | 0.00104184 |
| CRNDE | 1.216502099 | 0.00373432 | 0.00972429 |
| TMED6 | 1.213660501 | 0.01051784 | 0.02469699 |
| MRTO4 | 1.21249722 | 2.12E-50 | 1.55E-48 |
| LRRC8E | 1.210737848 | 0.00170978 | 0.00478952 |
| PYCR3 | 1.209578356 | 6.53E-18 | 8.92E-17 |
| TTC39B | 1.20948675 | 8.36E-14 | 8.22E-13 |
| IARS1 | 1.206870126 | 9.76E-65 | 1.21E-62 |
| LOC1001289 | 1.204435341 | 0.00024793 | 0.0008221 |
| CD36 | 1.203427003 | 3.08E-08 | 1.77E-07 |
| RAB13 | 1.202513194 | 9.01E-23 | 1.67E-21 |
| IGF2BP1 | 1.202252234 | 0.01033652 | 0.02433343 |
| TPRG1-AS1 | 1.199727023 | 2.22E-10 | 1.58E-09 |
| COL5A3 | 1.196062818 | 0.00626838 | 0.01553751 |
| MRAS | 1.195837582 | 9.06E-44 | 4.96E-42 |
| SLC11A1 | 1.194550838 | 2.31E-05 | 9.07E-05 |
| FCER2 | 1.19426872 | 0.0027016 | 0.00726388 |
| TGFBI | 1.193100976 | 2.27E-08 | 1.33E-07 |
| PPP1R14B | 1.190108058 | 1.97E-89 | 5.16E-87 |
| SEMA7A | 1.189844637 | 7.37E-31 | 2.17E-29 |
| PITPNC1 | 1.187692344 | 4.98E-22 | 8.83E-21 |
| ASPH | 1.186507018 | 4.08E-44 | 2.28E-42 |
| ST18 | 1.184116665 | 1.41E-11 | 1.12E-10 |
| CD82 | 1.181430679 | 8.86E-31 | 2.59E-29 |
| NR4A1 | 1.179856724 | 0.01068711 | 0.02506597 |
| LAMA5 | 1.178052327 | 0.00184276 | 0.00512555 |

| | | | |
|---|---|---|---|
| SLC2A5 | 1.175961763 | 5.59E-24 | 1.13E-22 |
| FLNB | 1.174556169 | 5.00E-42 | 2.57E-40 |
| PHETA1 | 1.165716293 | 4.18E-18 | 5.78E-17 |
| ARRDC4 | 1.162796282 | 2.55E-17 | 3.34E-16 |
| RUVBL1 | 1.162189117 | 2.87E-48 | 1.95E-46 |
| FAM86DP | 1.161700942 | 6.04E-30 | 1.71E-28 |
| ST3GAL6-AS | 1.161538126 | 7.96E-06 | 3.36E-05 |
| CXCL16 | 1.161176862 | 1.08E-35 | 3.96E-34 |
| LETM2 | 1.160174659 | 6.65E-10 | 4.54E-09 |
| LINC01588 | 1.158432367 | 1.89E-08 | 1.11E-07 |
| NOS1AP | 1.157334542 | 0.00294659 | 0.00786034 |
| KIF21A | 1.156001239 | 2.86E-05 | 0.00011057 |
| IFRD1 | 1.153844657 | 6.83E-22 | 1.20E-20 |
| BCL6 | 1.153736274 | 7.66E-20 | 1.18E-18 |
| TFRC | 1.150069214 | 1.59E-17 | 2.10E-16 |
| DDX11L2 | 1.149321311 | 0.01698744 | 0.03784602 |
| TNFSF14 | 1.147574352 | 2.18E-05 | 8.61E-05 |
| TXLNB | 1.14554978 | 5.58E-39 | 2.46E-37 |
| RRP12 | 1.138677894 | 8.65E-37 | 3.38E-35 |
| C1orf226 | 1.13838609 | 0.00188963 | 0.00524975 |
| MRPL17 | 1.13725626 | 3.22E-61 | 3.37E-59 |
| PTGS1 | 1.13640785 | 1.10E-87 | 2.71E-85 |
| OSM | 1.135500255 | 6.21E-09 | 3.83E-08 |
| PTGS2 | 1.133430123 | 1.85E-26 | 4.28E-25 |
| WDR4 | 1.129886092 | 2.05E-19 | 3.08E-18 |
| LDHA | 1.128339693 | 3.42E-36 | 1.30E-34 |
| LAPTM4B | 1.12746226 | 9.61E-12 | 7.85E-11 |
| TRAF4 | 1.127061286 | 5.50E-11 | 4.13E-10 |
| FOSL2 | 1.125050187 | 2.88E-16 | 3.45E-15 |
| PDSS1 | 1.119701291 | 3.80E-24 | 7.79E-23 |
| LPCAT2 | 1.119528097 | 5.42E-40 | 2.53E-38 |
| PRG2 | 1.116381895 | 0.00491481 | 0.01247798 |
| MIR155HG | 1.114306468 | 2.07E-07 | 1.08E-06 |
| GNPNAT1 | 1.11414255 | 3.80E-39 | 1.70E-37 |
| RFX2 | 1.112126288 | 1.16E-17 | 1.56E-16 |
| BOLA2-SMG1 | 1.111668896 | 7.15E-09 | 4.39E-08 |
| SLC41A2 | 1.108684101 | 8.58E-19 | 1.24E-17 |
| RAC2 | 1.108389049 | 1.43E-57 | 1.33E-55 |
| CLEC4D | 1.106142869 | 0.00965408 | 0.02289312 |
| CEP170B | 1.100409666 | 3.06E-28 | 7.86E-27 |
| DLEU1 | 1.099659683 | 5.96E-08 | 3.32E-07 |
| DYNC2H1 | 1.098291757 | 1.86E-17 | 2.45E-16 |
| PCSK6 | 1.097547556 | 2.72E-12 | 2.34E-11 |

| | | | |
|---|---|---|---|
| NFIL3 | 1.097523136 | 1.13E-12 | 1.01E-11 |
| TCAF2 | 1.096444766 | 1.76E-08 | 1.04E-07 |
| PANX1 | 1.096267244 | 5.23E-24 | 1.06E-22 |
| CMSS1 | 1.09499465 | 6.70E-17 | 8.46E-16 |
| ACOX2 | 1.093486534 | 2.33E-10 | 1.65E-09 |
| LIMK1 | 1.093398084 | 8.29E-39 | 3.62E-37 |
| STEAP3 | 1.091782894 | 1.44E-31 | 4.42E-30 |
| MTF1 | 1.088731731 | 3.62E-27 | 8.68E-26 |
| PGAM1 | 1.088361482 | 9.10E-66 | 1.15E-63 |
| FARSB | 1.087198527 | 6.35E-36 | 2.36E-34 |
| PHGDH | 1.086659783 | 1.24E-09 | 8.25E-09 |
| FADS3 | 1.085530602 | 1.45E-25 | 3.24E-24 |
| CRPPA | 1.085328608 | 0.00104094 | 0.0030476 |
| PLK3 | 1.084933372 | 8.05E-16 | 9.31E-15 |
| BOLA3 | 1.083770901 | 7.19E-36 | 2.67E-34 |
| P2RX7 | 1.083385509 | 1.58E-20 | 2.56E-19 |
| SLC35E4 | 1.083264904 | 1.02E-13 | 9.93E-13 |
| ADSS2 | 1.083023639 | 5.62E-25 | 1.21E-23 |
| TM4SF19-AS | 1.080388145 | 3.38E-06 | 1.51E-05 |
| SLC1A5 | 1.077900419 | 1.01E-39 | 4.65E-38 |
| IFRD2 | 1.076817048 | 7.72E-38 | 3.18E-36 |
| TTC25 | 1.074148422 | 0.00454643 | 0.01164825 |
| EIF5AL1 | 1.073517142 | 2.16E-09 | 1.40E-08 |
| CDC42EP1 | 1.072794919 | 2.13E-12 | 1.85E-11 |
| NPM3 | 1.069247646 | 9.73E-20 | 1.49E-18 |
| PDK1 | 1.068740974 | 8.55E-16 | 9.86E-15 |
| ATP2B1-AS1 | 1.06848461 | 0.00013771 | 0.00047811 |
| SCN9A | 1.06823034 | 2.58E-05 | 0.00010077 |
| PFKP | 1.067473302 | 1.40E-39 | 6.42E-38 |
| DUSP14 | 1.06281717 | 1.65E-15 | 1.86E-14 |
| WDR74 | 1.062196132 | 8.58E-34 | 2.84E-32 |
| ALAS1 | 1.061982981 | 7.88E-14 | 7.77E-13 |
| C1QBP | 1.059248943 | 2.48E-53 | 2.00E-51 |
| MSC-AS1 | 1.056456518 | 3.04E-16 | 3.64E-15 |
| TCTEX1D1 | 1.056148833 | 7.91E-06 | 3.34E-05 |
| ABCB4 | 1.053027297 | 6.92E-13 | 6.26E-12 |
| UBE2J1 | 1.052354964 | 7.25E-06 | 3.08E-05 |
| TLCD3A | 1.050533475 | 5.01E-09 | 3.13E-08 |
| GGT8P | 1.050196273 | 0.00760025 | 0.01845423 |
| SNORD104 | 1.047882862 | 0.00368671 | 0.00961394 |
| GYS1 | 1.046744086 | 9.55E-44 | 5.21E-42 |
| CFAP36 | 1.046680392 | 2.11E-29 | 5.77E-28 |
| WDR34 | 1.045909398 | 1.20E-14 | 1.26E-13 |

| | | | |
|---|---|---|---|
| SEH1L | 1.045819976 | 5.40E-43 | 2.85E-41 |
| GSN-AS1 | 1.043789914 | 0.00084931 | 0.00253597 |
| ITPRIP | 1.043710666 | 6.94E-21 | 1.15E-19 |
| MBOAT7 | 1.043651039 | 4.29E-32 | 1.35E-30 |
| SDC4 | 1.042095954 | 4.57E-21 | 7.67E-20 |
| ZBTB17 | 1.041289826 | 2.10E-18 | 2.98E-17 |
| ABHD2 | 1.039120314 | 1.80E-26 | 4.17E-25 |
| GPATCH4 | 1.038886846 | 6.60E-27 | 1.57E-25 |
| ANGPTL6 | 1.038417235 | 8.14E-08 | 4.46E-07 |
| THBS2 | 1.037936229 | 2.39E-07 | 1.24E-06 |
| ACSL4 | 1.037852258 | 1.78E-33 | 5.87E-32 |
| GPT2 | 1.037434152 | 3.77E-13 | 3.49E-12 |
| CHRNE | 1.037052822 | 0.00888919 | 0.02125306 |
| IL6 | 1.036210232 | 0.00584401 | 0.01459281 |
| EBNA1BP2 | 1.035905904 | 1.30E-38 | 5.58E-37 |
| PUS7 | 1.034909179 | 5.88E-29 | 1.57E-27 |
| DDIT4L | 1.03483904 | 0.00248397 | 0.00672847 |
| TBL2 | 1.034788114 | 1.72E-18 | 2.45E-17 |
| POLR3G | 1.032104707 | 5.29E-14 | 5.28E-13 |
| ARL10 | 1.032028033 | 9.88E-24 | 1.95E-22 |
| LINC00869 | 1.031782675 | 5.51E-05 | 0.00020357 |
| LINC00623 | 1.029157905 | 3.35E-11 | 2.57E-10 |
| IER3 | 1.028648171 | 2.67E-05 | 0.00010391 |
| LINC01004 | 1.025915303 | 2.36E-06 | 1.07E-05 |
| CCDC57 | 1.025556908 | 1.49E-26 | 3.49E-25 |
| TGFBR3L | 1.025356438 | 0.01231793 | 0.02842718 |
| NOP2 | 1.024286224 | 2.41E-40 | 1.15E-38 |
| FAIM | 1.023738923 | 5.84E-12 | 4.89E-11 |
| WDR63 | 1.021854009 | 4.77E-06 | 2.08E-05 |
| SLC22A4 | 1.020213435 | 1.94E-07 | 1.02E-06 |
| ABCE1 | 1.019114945 | 7.34E-41 | 3.60E-39 |
| VEGFA | 1.017946019 | 5.70E-17 | 7.22E-16 |
| PTPRM | 1.017019837 | 6.29E-16 | 7.35E-15 |
| FAM198B-AS | 1.01645861 | 1.70E-14 | 1.77E-13 |
| RET | 1.015522911 | 1.86E-05 | 7.44E-05 |
| BCL11A | 1.011611315 | 1.64E-06 | 7.62E-06 |
| FDX2 | 1.011539811 | 1.50E-16 | 1.84E-15 |
| SLC2A1 | 1.009762153 | 0.0027884 | 0.00747538 |
| BTBD19 | 1.006547279 | 2.72E-09 | 1.74E-08 |
| CD1C | 1.006026103 | 0.00026875 | 0.0008853 |
| MAP2K3 | 1.004288946 | 6.16E-21 | 1.02E-19 |
| PRMT5 | 1.002616295 | 9.57E-35 | 3.35E-33 |
| BCYRN1 | 1.002139605 | 0.01545948 | 0.03486449 |

| | | | |
|---|---|---|---|
| PLEKHF1 | 1.001501069 | 1.78E-06 | 8.25E-06 |
| TRAF1 | 1.000588929 | 2.91E-14 | 2.96E-13 |
| COLGALT2 | 1.000515092 | 1.65E-12 | 1.44E-11 |
| CALCOCO1 | -1.000250994 | 2.31E-50 | 1.66E-48 |
| EPB41 | -1.001575977 | 2.62E-38 | 1.12E-36 |
| PDE6B | -1.00358454 | 1.09E-16 | 1.36E-15 |
| OPTN | -1.005278423 | 2.83E-52 | 2.20E-50 |
| ULK2 | -1.006337308 | 5.68E-30 | 1.61E-28 |
| HAAO | -1.006692583 | 1.64E-10 | 1.18E-09 |
| RABGAP1L | -1.006757404 | 2.27E-28 | 5.90E-27 |
| INKA1 | -1.007747915 | 0.00085183 | 0.00254257 |
| ADRB2 | -1.007892239 | 3.44E-17 | 4.45E-16 |
| TES | -1.008475905 | 7.59E-36 | 2.80E-34 |
| MICA-AS1 | -1.010248544 | 7.95E-07 | 3.85E-06 |
| RSAD2 | -1.010387104 | 0.01754176 | 0.0389495 |
| FCHSD2 | -1.011562353 | 8.83E-16 | 1.02E-14 |
| TLR10 | -1.011824751 | 0.01286785 | 0.02953548 |
| TSPAN15 | -1.012255444 | 7.97E-20 | 1.23E-18 |
| PARP8 | -1.012453475 | 3.22E-28 | 8.22E-27 |
| RHOU | -1.014689716 | 4.81E-53 | 3.84E-51 |
| CHDH | -1.014727228 | 0.00056539 | 0.00175378 |
| GHRL | -1.017151126 | 0.00873794 | 0.02092474 |
| GABBR1 | -1.020743782 | 6.47E-20 | 1.00E-18 |
| TMEM176B | -1.022118811 | 1.19E-38 | 5.12E-37 |
| SOWAHD | -1.022794189 | 1.24E-32 | 3.96E-31 |
| MPO | -1.024367937 | 1.17E-08 | 7.06E-08 |
| ABTB1 | -1.024502465 | 1.26E-32 | 4.03E-31 |
| RNASE1 | -1.024752962 | 2.25E-79 | 4.38E-77 |
| TMEM144 | -1.024866436 | 5.62E-48 | 3.78E-46 |
| MAPK12 | -1.025080609 | 7.26E-09 | 4.46E-08 |
| TESK2 | -1.027356752 | 7.36E-12 | 6.10E-11 |
| SETBP1 | -1.031133153 | 0.00047277 | 0.00148853 |
| RAB3IL1 | -1.031242129 | 2.48E-55 | 2.16E-53 |
| GIMAP8 | -1.032244136 | 1.99E-55 | 1.74E-53 |
| ZNF774 | -1.032914939 | 7.73E-07 | 3.75E-06 |
| RTP4 | -1.033135206 | 3.67E-07 | 1.86E-06 |
| HEXA | -1.034006004 | 2.51E-65 | 3.14E-63 |
| PAQR7 | -1.03468784 | 6.35E-19 | 9.28E-18 |
| ZNF610 | -1.035239563 | 0.01094084 | 0.02558486 |
| APOL3 | -1.035859345 | 6.84E-16 | 7.97E-15 |
| ABHD12B | -1.036270565 | 0.00470051 | 0.01198727 |
| ZNF185 | -1.036560444 | 2.60E-61 | 2.73E-59 |
| HLA-F-AS1 | -1.037008156 | 5.93E-06 | 2.54E-05 |

| | | | |
|---|---|---|---|
| TSSK3 | -1.039489971 | 0.00236417 | 0.00643723 |
| STAB1 | -1.039575139 | 1.30E-42 | 6.82E-41 |
| C1QA | -1.039767755 | 9.91E-27 | 2.34E-25 |
| SNORA44 | -1.04130923 | 0.00991386 | 0.02345198 |
| PTPRB | -1.041355104 | 0.00761619 | 0.01848889 |
| TRIM22 | -1.041834833 | 1.71E-13 | 1.63E-12 |
| KCNJ2 | -1.042883201 | 9.24E-13 | 8.25E-12 |
| EPB41L1 | -1.043097 | 1.50E-15 | 1.70E-14 |
| PDCD4-AS1 | -1.043561934 | 4.19E-05 | 0.00015813 |
| STARD10 | -1.045461857 | 2.25E-09 | 1.46E-08 |
| AMDHD2 | -1.047349973 | 5.93E-41 | 2.93E-39 |
| NISCH | -1.048734617 | 4.46E-52 | 3.41E-50 |
| RAB11FIP1 | -1.051575263 | 4.69E-60 | 4.70E-58 |
| TAGAP | -1.052446338 | 3.24E-54 | 2.73E-52 |
| TNFRSF14 | -1.053034082 | 2.79E-39 | 1.26E-37 |
| GPR18 | -1.053266224 | 0.00847949 | 0.02037072 |
| AHRR | -1.053901808 | 8.43E-21 | 1.39E-19 |
| LILRB5 | -1.054662658 | 3.13E-29 | 8.47E-28 |
| PDE4B | -1.054851611 | 1.47E-16 | 1.80E-15 |
| COL18A1 | -1.055317564 | 5.72E-10 | 3.93E-09 |
| WEE2-AS1 | -1.055473302 | 2.27E-05 | 8.94E-05 |
| CES4A | -1.055572643 | 5.53E-10 | 3.80E-09 |
| EMP2 | -1.05703718 | 0.00016496 | 0.00056467 |
| NAPRT | -1.057851972 | 3.83E-21 | 6.45E-20 |
| HLA-DPA1 | -1.057955515 | 5.07E-51 | 3.77E-49 |
| S1PR1 | -1.05839308 | 4.73E-09 | 2.96E-08 |
| EPAS1 | -1.058471497 | 2.62E-25 | 5.74E-24 |
| ISG20 | -1.058872576 | 0.00238634 | 0.00648985 |
| PRKAR2A-AS | -1.05922987 | 4.33E-06 | 1.90E-05 |
| FBXO6 | -1.059505691 | 1.09E-26 | 2.57E-25 |
| PPP1R21 | -1.060079934 | 5.90E-36 | 2.21E-34 |
| PFKFB2 | -1.060399022 | 1.05E-41 | 5.36E-40 |
| GNGT2 | -1.061954511 | 1.76E-19 | 2.65E-18 |
| GSTM4 | -1.062154158 | 5.19E-22 | 9.19E-21 |
| CAPS2 | -1.062785819 | 7.75E-05 | 0.00027974 |
| PLEKHA5 | -1.063606197 | 4.13E-11 | 3.14E-10 |
| PDZD7 | -1.064687885 | 0.0009569 | 0.00282507 |
| DENND11 | -1.067186239 | 6.01E-31 | 1.77E-29 |
| LOC1019270 | -1.069703957 | 5.06E-11 | 3.82E-10 |
| NALT1 | -1.06990572 | 0.00761676 | 0.01848889 |
| MIR635 | -1.070125972 | 0.00548168 | 0.01377544 |
| KIAA1211L | -1.070188285 | 7.70E-10 | 5.22E-09 |
| CASP1 | -1.070302338 | 1.48E-47 | 9.75E-46 |

| | | | |
|---|---|---|---|
| ANO9 | -1.073073824 | 0.01163081 | 0.02696567 |
| AKR1B1 | -1.073352863 | 1.46E-48 | 9.91E-47 |
| LINC01678 | -1.074672733 | 7.56E-08 | 4.16E-07 |
| PEX11G | -1.075392244 | 2.27E-09 | 1.47E-08 |
| LINC00968 | -1.076456833 | 0.00136989 | 0.00390893 |
| LINC01637 | -1.076870048 | 2.79E-07 | 1.43E-06 |
| LY75 | -1.07703317 | 2.52E-23 | 4.87E-22 |
| CCDC9B | -1.077773632 | 2.05E-25 | 4.51E-24 |
| JAK3 | -1.079103952 | 6.99E-25 | 1.49E-23 |
| BCAS3 | -1.079159015 | 5.96E-27 | 1.42E-25 |
| MTMR8 | -1.079920369 | 0.00098492 | 0.00290159 |
| PRAME | -1.080834923 | 0.00709538 | 0.01736096 |
| L1CAM | -1.083935073 | 0.00604813 | 0.01504119 |
| LOC1005075 | -1.084928942 | 0.00011361 | 0.00039963 |
| IRF1-AS1 | -1.085036406 | 0.00107979 | 0.00314799 |
| EIF1B-AS1 | -1.085633915 | 0.00230465 | 0.00628656 |
| HDHD3 | -1.086246934 | 2.31E-21 | 3.95E-20 |
| RGS18 | -1.087894537 | 6.20E-12 | 5.17E-11 |
| KRBA1 | -1.087923903 | 4.82E-26 | 1.08E-24 |
| OAZ3 | -1.088101001 | 4.56E-08 | 2.57E-07 |
| DENND2C | -1.088166519 | 0.00177042 | 0.00494182 |
| TNFSF13B | -1.088787131 | 3.52E-25 | 7.66E-24 |
| MYLK4 | -1.089793693 | 0.00102173 | 0.00299454 |
| SLC44A2 | -1.091015959 | 8.20E-24 | 1.63E-22 |
| IL11RA | -1.091250897 | 1.02E-06 | 4.87E-06 |
| IFITM2 | -1.091319156 | 6.55E-13 | 5.95E-12 |
| RHPN1 | -1.091358845 | 3.10E-07 | 1.58E-06 |
| AVPR2 | -1.091596303 | 0.00363965 | 0.00950171 |
| BACE2 | -1.091764963 | 9.59E-08 | 5.22E-07 |
| LOC1019280 | -1.092042219 | 0.01691157 | 0.03772247 |
| LPAR6 | -1.094564638 | 1.10E-16 | 1.36E-15 |
| DDX43 | -1.094749776 | 2.26E-05 | 8.92E-05 |
| FCGRT | -1.095292923 | 6.51E-56 | 5.82E-54 |
| M1AP | -1.095722951 | 1.08E-11 | 8.78E-11 |
| ZNF114 | -1.096676893 | 0.00070992 | 0.00215677 |
| APOL1 | -1.09784868 | 1.14E-22 | 2.10E-21 |
| ADA | -1.098177544 | 2.66E-15 | 2.93E-14 |
| PCBP1-AS1 | -1.098227076 | 1.02E-07 | 5.51E-07 |
| LOC1019283 | -1.101304678 | 0.02195539 | 0.04744947 |
| KSR1 | -1.102975061 | 7.58E-21 | 1.25E-19 |
| DLEC1 | -1.103159227 | 2.30E-05 | 9.05E-05 |
| LOC1002881 | -1.103810826 | 4.00E-05 | 0.00015127 |
| CTSV | -1.104919845 | 1.08E-05 | 4.47E-05 |

| | | | |
|---|---|---|---|
| KLRF1 | -1.105715084 | 0.01576662 | 0.0354749 |
| SIRPB2 | -1.106340799 | 2.22E-78 | 4.16E-76 |
| PRR5 | -1.106677134 | 3.75E-23 | 7.14E-22 |
| GIMAP4 | -1.106721306 | 6.69E-46 | 4.16E-44 |
| RCN3 | -1.110001382 | 1.17E-52 | 9.24E-51 |
| FUT7 | -1.111408536 | 5.38E-10 | 3.71E-09 |
| GARNL3 | -1.113595269 | 0.00011291 | 0.00039753 |
| BTN2A2 | -1.115107001 | 3.02E-31 | 9.08E-30 |
| ATP8B4 | -1.115765435 | 1.79E-35 | 6.47E-34 |
| LINC02021 | -1.116103242 | 0.01644168 | 0.03679848 |
| CECR2 | -1.11616639 | 0.00959587 | 0.02277144 |
| ATP9A | -1.116430156 | 0.00041405 | 0.0013177 |
| GIMAP5 | -1.118411473 | 1.99E-15 | 2.22E-14 |
| SLCO4C1 | -1.118728159 | 0.00129972 | 0.00373254 |
| METTL7A | -1.121670196 | 3.11E-23 | 5.97E-22 |
| FAM13A | -1.122406873 | 0.00075155 | 0.00226676 |
| MAP7 | -1.122919216 | 2.38E-19 | 3.57E-18 |
| LOC1027237 | -1.124111623 | 3.45E-05 | 0.00013185 |
| C2orf15 | -1.124842552 | 0.01682534 | 0.0375656 |
| TOB1-AS1 | -1.125742681 | 0.01611967 | 0.03617572 |
| RALGPS1 | -1.126718891 | 6.58E-13 | 5.97E-12 |
| TCN2 | -1.126958324 | 2.99E-28 | 7.70E-27 |
| MMEL1 | -1.128058063 | 0.00389621 | 0.01010448 |
| LOC1005072 | -1.128130289 | 0.00740409 | 0.01802029 |
| ABHD12 | -1.128374111 | 2.97E-62 | 3.25E-60 |
| AKR1C3 | -1.128379224 | 0.02233416 | 0.04817989 |
| C5orf63 | -1.128428265 | 0.00775132 | 0.01879346 |
| ADCY10P1 | -1.130117122 | 1.17E-06 | 5.56E-06 |
| HORMAD1 | -1.130326047 | 0.00237625 | 0.00646478 |
| ZSCAN31 | -1.131301485 | 0.00011116 | 0.00039177 |
| AAMDC | -1.131391473 | 4.53E-18 | 6.25E-17 |
| FOLR3 | -1.131742134 | 0.02183362 | 0.04721717 |
| PNMA8B | -1.131963948 | 0.00618353 | 0.01534651 |
| ZNF540 | -1.131975056 | 0.00915757 | 0.02182215 |
| TMEM176A | -1.132049708 | 1.17E-50 | 8.63E-49 |
| FAM117A | -1.133648019 | 1.41E-17 | 1.88E-16 |
| PIK3CD-AS1 | -1.133657258 | 0.00168965 | 0.00473635 |
| ACRBP | -1.134160991 | 4.10E-24 | 8.37E-23 |
| HLA-DPB1 | -1.134572137 | 8.01E-53 | 6.37E-51 |
| PNMA6A | -1.135064249 | 0.00272409 | 0.00731483 |
| FEZ1 | -1.136088682 | 0.00369142 | 0.00962318 |
| SSPO | -1.137988741 | 1.08E-05 | 4.48E-05 |
| C1QC | -1.138638932 | 2.03E-35 | 7.28E-34 |

| | | | |
|---|---|---|---|
| SDC3 | -1.139351985 | 5.06E-05 | 0.00018826 |
| KLHL24 | -1.140572508 | 1.87E-34 | 6.43E-33 |
| SUSD3 | -1.14116912 | 0.00194614 | 0.00539134 |
| FCGBP | -1.145330553 | 0.01229278 | 0.02837707 |
| TIFAB | -1.146684382 | 0.00314862 | 0.00834903 |
| CCDC69 | -1.146976826 | 4.60E-47 | 2.94E-45 |
| SLC22A23 | -1.14779242 | 1.89E-16 | 2.30E-15 |
| KIAA1522 | -1.147839879 | 1.17E-22 | 2.14E-21 |
| ZNF233 | -1.148257715 | 0.00494882 | 0.01255276 |
| ADAMTS10 | -1.150559918 | 5.27E-05 | 0.00019551 |
| NBEA | -1.150892777 | 2.54E-11 | 1.97E-10 |
| TNFRSF14-A: | -1.150997406 | 1.57E-24 | 3.27E-23 |
| STAC3 | -1.151686596 | 2.29E-54 | 1.97E-52 |
| COL9A2 | -1.151932204 | 6.32E-06 | 2.70E-05 |
| MROCKI | -1.152688914 | 0.00075859 | 0.00228573 |
| ILDR2 | -1.152844116 | 0.01603931 | 0.03601981 |
| CD320 | -1.153272384 | 0.00242641 | 0.00659042 |
| MPEG1 | -1.153581438 | 3.54E-64 | 4.21E-62 |
| NCOA4 | -1.154257796 | 8.84E-128 | 5.84E-125 |
| RNASE2 | -1.154462644 | 3.58E-17 | 4.61E-16 |
| STAP1 | -1.15503074 | 0.00245686 | 0.00666656 |
| HVCN1 | -1.155038239 | 1.05E-43 | 5.71E-42 |
| FES | -1.155751439 | 3.28E-67 | 4.52E-65 |
| C1QB | -1.158658351 | 4.39E-25 | 9.49E-24 |
| MXD4 | -1.159793261 | 1.40E-48 | 9.56E-47 |
| ABAT | -1.159806046 | 3.09E-30 | 8.88E-29 |
| TSPAN9 | -1.159833495 | 1.10E-05 | 4.57E-05 |
| PNMA2 | -1.160585093 | 0.01342921 | 0.03072161 |
| LINC00886 | -1.163203023 | 0.00538877 | 0.01356257 |
| CBLN3 | -1.164708988 | 2.67E-09 | 1.72E-08 |
| DLL1 | -1.165255018 | 3.39E-24 | 6.99E-23 |
| LY86 | -1.167497711 | 5.53E-61 | 5.75E-59 |
| TMEM104 | -1.168063375 | 4.66E-49 | 3.24E-47 |
| GAMT | -1.171550536 | 1.52E-13 | 1.45E-12 |
| CYP2T1P | -1.171571079 | 0.00047556 | 0.00149619 |
| CYB561A3 | -1.172031493 | 1.22E-27 | 3.03E-26 |
| MIR1249 | -1.1748141 | 0.01426335 | 0.03245025 |
| LGMN | -1.176638268 | 3.67E-31 | 1.10E-29 |
| STK32B | -1.179127196 | 5.63E-24 | 1.14E-22 |
| FBXL8 | -1.179157331 | 3.06E-09 | 1.94E-08 |
| FZD9 | -1.179919165 | 0.0003351 | 0.00108545 |
| PCSK4 | -1.181136883 | 4.82E-12 | 4.07E-11 |
| CSF3R | -1.182691283 | 2.19E-58 | 2.09E-56 |

| | | | |
|---|---|---|---|
| PDCD1LG2 | -1.182960235 | 6.82E-41 | 3.36E-39 |
| CDC14B | -1.184184552 | 8.45E-05 | 0.00030306 |
| CTF1 | -1.18609948 | 0.00175508 | 0.00490312 |
| PLCXD2 | -1.186719166 | 0.01296045 | 0.02972327 |
| GM2A | -1.187267345 | 1.06E-49 | 7.50E-48 |
| FAM171B | -1.188553129 | 0.00025006 | 0.00082802 |
| MTMR9LP | -1.189379813 | 2.69E-05 | 0.00010467 |
| CD302 | -1.191573718 | 1.03E-19 | 1.56E-18 |
| LINC01907 | -1.197450733 | 8.83E-05 | 0.00031622 |
| IL32 | -1.197708641 | 3.60E-06 | 1.59E-05 |
| ABCA5 | -1.198051947 | 4.23E-24 | 8.63E-23 |
| NINJ2 | -1.201032175 | 1.97E-34 | 6.76E-33 |
| SH2D3C | -1.201499042 | 8.40E-34 | 2.79E-32 |
| FAM157A | -1.202875166 | 0.01291021 | 0.0296286 |
| LRRC8C-DT | -1.202944808 | 0.00301121 | 0.00802326 |
| CACNB1 | -1.203380237 | 3.46E-09 | 2.19E-08 |
| COL23A1 | -1.204533711 | 7.05E-40 | 3.27E-38 |
| PGA5 | -1.20682012 | 0.00182188 | 0.00507262 |
| HESX1 | -1.208100001 | 2.32E-05 | 9.12E-05 |
| P2RY6 | -1.209365786 | 1.47E-62 | 1.64E-60 |
| RNF144A | -1.209399536 | 4.73E-06 | 2.07E-05 |
| AIFM3 | -1.209407288 | 2.08E-14 | 2.15E-13 |
| RELL1 | -1.209552856 | 1.40E-23 | 2.75E-22 |
| SLC46A3 | -1.209804705 | 4.61E-45 | 2.74E-43 |
| NOXA1 | -1.20984798 | 5.65E-36 | 2.12E-34 |
| FKBP1B | -1.210318691 | 8.43E-12 | 6.92E-11 |
| GRIN3A | -1.210631839 | 7.59E-34 | 2.54E-32 |
| HLA-DQA1 | -1.211621681 | 1.05E-45 | 6.44E-44 |
| GRAMD1B | -1.211838777 | 1.06E-31 | 3.27E-30 |
| SPINT2 | -1.211857395 | 2.63E-73 | 4.39E-71 |
| NAIP | -1.212461229 | 6.79E-60 | 6.72E-58 |
| HFE | -1.213234698 | 1.33E-35 | 4.87E-34 |
| ZNF192P1 | -1.216499688 | 0.01650243 | 0.03691444 |
| ID3 | -1.217158907 | 3.62E-11 | 2.77E-10 |
| XYLT1 | -1.217634308 | 2.46E-32 | 7.83E-31 |
| LINC00653 | -1.217871072 | 0.00618483 | 0.01534651 |
| TRPM2 | -1.218038653 | 1.99E-54 | 1.71E-52 |
| IDH1-AS1 | -1.218927662 | 0.01251864 | 0.02882198 |
| GPR141 | -1.220933526 | 5.38E-71 | 8.56E-69 |
| SH3RF3 | -1.221176429 | 6.04E-12 | 5.05E-11 |
| RAB32 | -1.221689051 | 2.21E-50 | 1.60E-48 |
| ADGRG6 | -1.226202595 | 5.16E-12 | 4.34E-11 |
| SH3RF3-AS1 | -1.228329924 | 0.00349268 | 0.00916578 |

| | | | |
|---|---|---|---|
| MICAL1 | -1.234836172 | 1.08E-38 | 4.69E-37 |
| N4BP2L1 | -1.236717797 | 4.20E-29 | 1.13E-27 |
| PPP1R1A | -1.238172366 | 0.02280118 | 0.04905932 |
| ITGA7 | -1.238445857 | 4.88E-06 | 2.12E-05 |
| FRY | -1.238872223 | 5.04E-16 | 5.94E-15 |
| TMEM169 | -1.239246302 | 4.04E-05 | 0.00015282 |
| CCDC80 | -1.239457587 | 0.00238664 | 0.00648985 |
| PRKN | -1.239704438 | 0.00707529 | 0.01731948 |
| LINC01002 | -1.241124295 | 0.00313563 | 0.00831591 |
| SULT1A1 | -1.241647123 | 2.76E-67 | 3.83E-65 |
| TMEM86A | -1.244802894 | 3.71E-38 | 1.57E-36 |
| ECHDC2 | -1.246860158 | 4.47E-07 | 2.25E-06 |
| SEC31B | -1.24941419 | 1.49E-16 | 1.82E-15 |
| NID1 | -1.251146871 | 3.95E-20 | 6.25E-19 |
| LINC02246 | -1.252467063 | 1.02E-06 | 4.89E-06 |
| RNF157 | -1.254310806 | 0.00321992 | 0.00851759 |
| TSPAN4 | -1.255979953 | 1.64E-60 | 1.67E-58 |
| AASS | -1.256310082 | 2.25E-05 | 8.87E-05 |
| PBLD | -1.256504204 | 2.40E-18 | 3.40E-17 |
| HLA-DQB1 | -1.259851724 | 1.48E-42 | 7.77E-41 |
| LINC02542 | -1.262310929 | 4.35E-05 | 0.00016353 |
| DACT1 | -1.265476033 | 0.00261355 | 0.00704776 |
| SHF | -1.265520493 | 0.00173954 | 0.0048622 |
| SH3BP4 | -1.265679357 | 2.00E-06 | 9.19E-06 |
| LOC1003107 | -1.268399898 | 6.24E-10 | 4.27E-09 |
| NREP | -1.270347204 | 6.24E-25 | 1.34E-23 |
| THBS1 | -1.272675367 | 1.48E-13 | 1.42E-12 |
| CCDC146 | -1.274054622 | 3.85E-07 | 1.95E-06 |
| CHADL | -1.274375182 | 0.00033458 | 0.00108398 |
| APBA1 | -1.274881688 | 1.43E-19 | 2.16E-18 |
| MPP7 | -1.275234682 | 3.57E-08 | 2.04E-07 |
| USP30-AS1 | -1.276052191 | 0.00246442 | 0.00668268 |
| ARHGAP27P | -1.276717411 | 5.02E-39 | 2.22E-37 |
| AMIGO2 | -1.277802272 | 9.30E-06 | 3.89E-05 |
| STARD13 | -1.278727338 | 7.70E-75 | 1.33E-72 |
| MEGF6 | -1.281550468 | 1.07E-26 | 2.53E-25 |
| GSDMD | -1.282436364 | 2.94E-44 | 1.65E-42 |
| EPSTI1 | -1.283018969 | 3.11E-07 | 1.59E-06 |
| CMAHP | -1.283173481 | 1.80E-20 | 2.90E-19 |
| EPOR | -1.284185161 | 1.10E-24 | 2.31E-23 |
| C12orf75 | -1.290949646 | 3.16E-05 | 0.00012139 |
| TTC9 | -1.29174073 | 5.86E-12 | 4.91E-11 |
| KLRG1 | -1.29179858 | 2.46E-08 | 1.43E-07 |

| | | | |
|---|---|---|---|
| ABCB5 | -1.291927232 | 0.00049592 | 0.00155521 |
| PLXNC1 | -1.292289143 | 1.80E-36 | 6.97E-35 |
| CMKLR1 | -1.292392301 | 1.28E-56 | 1.18E-54 |
| EMB | -1.292915347 | 1.17E-28 | 3.09E-27 |
| ABLIM2 | -1.298547896 | 0.00063448 | 0.0019458 |
| TMC4 | -1.299410775 | 0.00012703 | 0.00044335 |
| GRID2IP | -1.300201649 | 0.00676225 | 0.01660976 |
| FCGR2A | -1.300734614 | 1.73E-58 | 1.66E-56 |
| WAKMAR2 | -1.301444582 | 0.02035574 | 0.04433426 |
| GVINP1 | -1.302040745 | 5.99E-34 | 2.01E-32 |
| CBX7 | -1.304749485 | 7.81E-24 | 1.56E-22 |
| SELL | -1.30691608 | 5.04E-23 | 9.54E-22 |
| FHIT | -1.309416036 | 3.93E-12 | 3.34E-11 |
| CTSF | -1.311336436 | 1.46E-38 | 6.26E-37 |
| LOC1001290 | -1.311779343 | 0.01643426 | 0.03678685 |
| ZBTB46 | -1.312780252 | 6.89E-15 | 7.35E-14 |
| ABLIM3 | -1.313088018 | 3.96E-26 | 8.93E-25 |
| MEST | -1.313930685 | 1.90E-08 | 1.12E-07 |
| TSKS | -1.31614194 | 0.00244041 | 0.00662299 |
| CLIC2 | -1.316752845 | 4.40E-47 | 2.82E-45 |
| TRPM2-AS | -1.319466132 | 0.00013291 | 0.00046252 |
| CLEC3B | -1.320413621 | 3.92E-14 | 3.96E-13 |
| CMPK2 | -1.322096482 | 9.25E-06 | 3.87E-05 |
| ATP6V0D2 | -1.32524188 | 2.65E-41 | 1.33E-39 |
| TRPV4 | -1.328258534 | 1.28E-24 | 2.68E-23 |
| RAB37 | -1.331416211 | 0.00195432 | 0.00541219 |
| SECTM1 | -1.331879079 | 2.01E-45 | 1.22E-43 |
| YPEL3 | -1.332180842 | 1.42E-27 | 3.48E-26 |
| GPR155 | -1.333774647 | 2.22E-27 | 5.39E-26 |
| FAM43A | -1.333782838 | 1.07E-07 | 5.78E-07 |
| SLC25A45 | -1.334928296 | 1.35E-23 | 2.65E-22 |
| SIRPB1 | -1.33692544 | 1.84E-37 | 7.50E-36 |
| SGMS1 | -1.337105538 | 1.83E-92 | 5.13E-90 |
| MRVI1 | -1.339623139 | 4.20E-20 | 6.62E-19 |
| SIRPD | -1.339892258 | 5.53E-10 | 3.80E-09 |
| TNFSF12 | -1.341642103 | 4.51E-37 | 1.80E-35 |
| PGGHG | -1.344102756 | 1.36E-29 | 3.75E-28 |
| CDC42EP3 | -1.344986414 | 5.15E-33 | 1.66E-31 |
| SHE | -1.34502244 | 0.00784208 | 0.01899402 |
| NCF4 | -1.345226246 | 5.00E-112 | 2.30E-109 |
| MFSD6L | -1.345845135 | 0.00354977 | 0.00929337 |
| MFAP3L | -1.347515509 | 0.00078626 | 0.0023622 |
| DAB2 | -1.351287185 | 5.87E-48 | 3.93E-46 |

| | | | |
|---|---|---|---|
| ELFN2 | -1.352894774 | 0.00154443 | 0.00436108 |
| ZNF33B | -1.353480548 | 2.82E-46 | 1.77E-44 |
| DNASE2 | -1.354753526 | 5.94E-63 | 6.73E-61 |
| LINC00528 | -1.360591344 | 1.01E-06 | 4.81E-06 |
| ACSS1 | -1.360904974 | 2.63E-31 | 7.96E-30 |
| LGR4 | -1.36170573 | 5.11E-30 | 1.45E-28 |
| NATD1 | -1.361823445 | 6.66E-24 | 1.34E-22 |
| CREBRF | -1.361946877 | 2.43E-33 | 7.93E-32 |
| ZNF491 | -1.362071129 | 4.20E-06 | 1.84E-05 |
| LINC02035 | -1.363312121 | 5.22E-21 | 8.74E-20 |
| FMO5 | -1.364465627 | 6.54E-16 | 7.64E-15 |
| FHDC1 | -1.365114978 | 0.00960486 | 0.02278948 |
| TSGA10 | -1.36571771 | 1.38E-08 | 8.25E-08 |
| OCIAD2 | -1.366895779 | 0.00027549 | 0.00090532 |
| TP53INP1 | -1.367503231 | 4.94E-60 | 4.92E-58 |
| PTH2R | -1.368316983 | 0.00369862 | 0.00963893 |
| CFP | -1.369606713 | 5.56E-21 | 9.29E-20 |
| SEC1P | -1.370429228 | 0.00029833 | 0.00097514 |
| PER3 | -1.370902901 | 7.94E-73 | 1.30E-70 |
| GRAP | -1.371390816 | 0.00012627 | 0.00044107 |
| LINC00926 | -1.371534572 | 8.32E-07 | 4.02E-06 |
| SDHAP3 | -1.371703954 | 3.56E-10 | 2.49E-09 |
| ADAMTS13 | -1.372290395 | 0.00183317 | 0.00510233 |
| KIF17 | -1.372833337 | 1.15E-05 | 4.76E-05 |
| STARD4-AS1 | -1.376185842 | 1.45E-07 | 7.73E-07 |
| PLEKHG3 | -1.378302972 | 4.44E-40 | 2.09E-38 |
| A2M-AS1 | -1.378515355 | 2.31E-15 | 2.56E-14 |
| OSBPL10 | -1.380006592 | 0.00025244 | 0.00083555 |
| RASL10A | -1.380310716 | 0.0040531 | 0.01047194 |
| NUDT7 | -1.381657072 | 1.84E-11 | 1.46E-10 |
| AMY2B | -1.382892684 | 5.24E-18 | 7.20E-17 |
| LRMP | -1.382957231 | 9.59E-19 | 1.38E-17 |
| FARP1 | -1.384590187 | 3.75E-21 | 6.32E-20 |
| OXER1 | -1.384905546 | 1.07E-11 | 8.67E-11 |
| PLAC8 | -1.386233723 | 6.39E-06 | 2.73E-05 |
| CPVL | -1.386234256 | 3.78E-52 | 2.92E-50 |
| SAT1 | -1.386293237 | 5.98E-24 | 1.21E-22 |
| WLS | -1.386366138 | 1.66E-39 | 7.60E-38 |
| SCARB1 | -1.387787786 | 1.28E-31 | 3.94E-30 |
| TRIM14 | -1.388926326 | 6.24E-38 | 2.59E-36 |
| SLC16A7 | -1.390069088 | 5.04E-17 | 6.42E-16 |
| CASTOR3 | -1.39037523 | 2.24E-12 | 1.94E-11 |
| CCDC88B | -1.390812511 | 3.01E-40 | 1.42E-38 |

| | | | |
|---|---|---|---|
| HCP5 | -1.392337538 | 3.31E-20 | 5.26E-19 |
| KCNQ1 | -1.394463476 | 1.99E-62 | 2.19E-60 |
| MAF | -1.394736505 | 1.31E-60 | 1.35E-58 |
| GBP5 | -1.394836901 | 4.15E-33 | 1.35E-31 |
| NTNG2 | -1.398136113 | 5.40E-06 | 2.33E-05 |
| CYBRD1 | -1.400703764 | 6.50E-75 | 1.13E-72 |
| FPR1 | -1.400921399 | 1.46E-07 | 7.79E-07 |
| LOC1001343 | -1.401738052 | 0.00107685 | 0.00313996 |
| SEMA6B | -1.403203349 | 5.41E-10 | 3.73E-09 |
| IGF2BP3 | -1.405989502 | 1.03E-12 | 9.22E-12 |
| MYOM1 | -1.406982343 | 1.17E-05 | 4.84E-05 |
| BIRC7 | -1.407187798 | 2.35E-07 | 1.22E-06 |
| KLF2 | -1.408061364 | 1.42E-15 | 1.61E-14 |
| CALCRL | -1.408324011 | 2.08E-20 | 3.34E-19 |
| MROH6 | -1.408397802 | 1.48E-14 | 1.55E-13 |
| TRIM2 | -1.411492676 | 5.69E-06 | 2.45E-05 |
| CDKN1C | -1.411677229 | 0.00011542 | 0.00040559 |
| ZNF812P | -1.411703596 | 0.02323694 | 0.04989951 |
| FAM227B | -1.411711687 | 0.00094138 | 0.00278472 |
| REPS2 | -1.413094189 | 3.42E-35 | 1.22E-33 |
| RALGPS2 | -1.41431013 | 3.02E-28 | 7.76E-27 |
| IQCN | -1.414839208 | 0.00087566 | 0.00260429 |
| SIGLEC10 | -1.415931718 | 2.33E-30 | 6.77E-29 |
| PARD3B | -1.420356977 | 0.01750734 | 0.03888351 |
| JAG2 | -1.42150824 | 3.89E-06 | 1.71E-05 |
| TRIM25 | -1.421550545 | 1.48E-77 | 2.68E-75 |
| PKD1L3 | -1.422840998 | 9.90E-08 | 5.38E-07 |
| AIG1 | -1.42359194 | 5.24E-42 | 2.69E-40 |
| SERPINB9P1 | -1.426030612 | 6.08E-15 | 6.51E-14 |
| PAQR8 | -1.428443366 | 7.70E-08 | 4.23E-07 |
| H2AC6 | -1.429608172 | 0.00472293 | 0.01203887 |
| ITM2B | -1.432744774 | 6.14E-78 | 1.14E-75 |
| CARD11 | -1.433014218 | 3.04E-06 | 1.36E-05 |
| SIGLEC1 | -1.433185008 | 5.02E-44 | 2.78E-42 |
| PCBP3 | -1.434439804 | 2.47E-05 | 9.68E-05 |
| TMEM71 | -1.434674821 | 1.11E-10 | 8.14E-10 |
| LOC1019295 | -1.435435297 | 1.41E-31 | 4.35E-30 |
| NAALAD2 | -1.436530804 | 0.01942771 | 0.04258794 |
| NFE2 | -1.438811252 | 0.01229176 | 0.02837707 |
| APPL2 | -1.442101208 | 1.59E-50 | 1.17E-48 |
| KLF4 | -1.442511367 | 1.16E-14 | 1.22E-13 |
| CCNG2 | -1.443011555 | 1.12E-21 | 1.95E-20 |
| DMTN | -1.443876638 | 0.00618392 | 0.01534651 |

| | | | |
|---|---|---|---|
| ALOX15B | -1.446129952 | 4.48E-18 | 6.19E-17 |
| TRAF3IP3 | -1.450635856 | 5.40E-31 | 1.60E-29 |
| GTSF1 | -1.451354859 | 1.98E-07 | 1.04E-06 |
| KCNAB1 | -1.453815067 | 1.53E-05 | 6.21E-05 |
| PBXIP1 | -1.456296532 | 2.06E-102 | 8.13E-100 |
| TXNDC16 | -1.457094531 | 1.17E-34 | 4.05E-33 |
| IGSF6 | -1.462502949 | 2.34E-44 | 1.33E-42 |
| CCSER1 | -1.465102113 | 2.04E-08 | 1.20E-07 |
| UNC80 | -1.465405222 | 5.32E-05 | 0.00019725 |
| IGSF22 | -1.466611903 | 0.00067396 | 0.00205545 |
| GSDMA | -1.467620934 | 1.22E-07 | 6.55E-07 |
| F2RL1 | -1.469849915 | 0.01866316 | 0.04114656 |
| SPATC1 | -1.470200149 | 3.91E-10 | 2.73E-09 |
| RHOB | -1.470997061 | 1.13E-49 | 7.94E-48 |
| DYNC1I1 | -1.473112318 | 0.00375955 | 0.00978074 |
| ZCCHC24 | -1.473118675 | 7.53E-78 | 1.38E-75 |
| RCSD1 | -1.476415153 | 3.73E-108 | 1.62E-105 |
| NFIA | -1.477829628 | 9.20E-16 | 1.06E-14 |
| PECAM1 | -1.480469172 | 5.24E-171 | 6.66E-168 |
| ACTR3C | -1.481769785 | 0.00068227 | 0.00207849 |
| GIMAP7 | -1.482002342 | 7.31E-76 | 1.29E-73 |
| THNSL2 | -1.48259396 | 4.09E-09 | 2.57E-08 |
| STING1 | -1.4834028 | 2.08E-64 | 2.53E-62 |
| SRPX | -1.484668288 | 2.90E-05 | 0.00011221 |
| PRUNE2 | -1.48494057 | 0.00015553 | 0.00053513 |
| COLQ | -1.485777309 | 1.45E-11 | 1.16E-10 |
| GUCY2D | -1.485790067 | 0.01115859 | 0.02602046 |
| CCDC153 | -1.486499539 | 2.06E-07 | 1.07E-06 |
| LMNTD2 | -1.489664343 | 5.99E-23 | 1.13E-21 |
| KCNA3 | -1.49005782 | 9.35E-09 | 5.67E-08 |
| LOC400499 | -1.490194144 | 1.46E-05 | 5.95E-05 |
| HTR2B | -1.492448714 | 6.00E-08 | 3.34E-07 |
| LINC00278 | -1.494261169 | 9.19E-10 | 6.20E-09 |
| KLHDC8B | -1.494782385 | 1.86E-61 | 1.97E-59 |
| LINC00865 | -1.497466881 | 9.47E-10 | 6.39E-09 |
| CCDC88C | -1.497636128 | 2.69E-43 | 1.44E-41 |
| PCED1B | -1.500201654 | 9.74E-39 | 4.24E-37 |
| EBI3 | -1.503349917 | 9.42E-32 | 2.92E-30 |
| ZNF781 | -1.504059155 | 0.01153174 | 0.02677575 |
| PLA2R1 | -1.51056709 | 0.01810884 | 0.04009017 |
| MARVELD3 | -1.511337042 | 0.00297596 | 0.00793444 |
| EPHX1 | -1.5116778 | 5.73E-64 | 6.72E-62 |
| CC2D2B | -1.512683237 | 2.84E-05 | 0.00011007 |

| | | | |
|---|---|---|---|
| DNM3 | -1.512946447 | 5.20E-14 | 5.20E-13 |
| HES1 | -1.513076226 | 2.16E-05 | 8.56E-05 |
| LINC01506 | -1.513398296 | 0.0010728 | 0.00312925 |
| TRPC2 | -1.513976788 | 4.82E-05 | 0.00017983 |
| ACCS | -1.515025824 | 9.73E-19 | 1.40E-17 |
| CMBL | -1.517968483 | 2.82E-43 | 1.51E-41 |
| FMO4 | -1.522942507 | 1.39E-32 | 4.43E-31 |
| LOC1033525 | -1.52394818 | 0.02202469 | 0.04757437 |
| GPR85 | -1.524226707 | 2.19E-05 | 8.65E-05 |
| CELF6 | -1.524244417 | 3.32E-27 | 8.00E-26 |
| PIPOX | -1.525142746 | 1.54E-16 | 1.88E-15 |
| OLIG1 | -1.525557663 | 1.08E-10 | 7.90E-10 |
| ABHD11-AS1 | -1.525762705 | 0.02025323 | 0.04415173 |
| MYCL | -1.526938463 | 2.86E-30 | 8.23E-29 |
| TMEM37 | -1.528056385 | 2.33E-72 | 3.77E-70 |
| CDA | -1.529019538 | 8.75E-62 | 9.52E-60 |
| SCNN1A | -1.530161627 | 0.02267336 | 0.04881608 |
| CARD16 | -1.532145721 | 4.44E-37 | 1.78E-35 |
| PELI2 | -1.532479291 | 3.62E-21 | 6.12E-20 |
| SLIT1 | -1.537109154 | 7.45E-13 | 6.73E-12 |
| PIK3R3 | -1.540987362 | 7.41E-08 | 4.08E-07 |
| SEPTIN3 | -1.543477938 | 2.70E-17 | 3.51E-16 |
| CFH | -1.545238272 | 1.40E-10 | 1.01E-09 |
| PRKAR2B | -1.545284848 | 2.05E-24 | 4.27E-23 |
| CES3 | -1.550160846 | 1.57E-15 | 1.77E-14 |
| YPEL2 | -1.551947765 | 6.58E-24 | 1.32E-22 |
| WWP1 | -1.554506275 | 8.51E-75 | 1.45E-72 |
| SLC45A4 | -1.555081051 | 7.10E-54 | 5.90E-52 |
| ETS2 | -1.555222778 | 2.26E-69 | 3.36E-67 |
| PTPRN2 | -1.555562053 | 1.55E-18 | 2.22E-17 |
| PLSCR4 | -1.559969534 | 0.00573573 | 0.01434193 |
| SLC5A9 | -1.562920859 | 0.01126703 | 0.02623261 |
| FBLIM1 | -1.563662242 | 9.91E-07 | 4.74E-06 |
| RBP7 | -1.566865987 | 4.86E-15 | 5.23E-14 |
| LOC1027241 | -1.570705147 | 0.01419952 | 0.03233729 |
| PHACTR1 | -1.571093289 | 1.86E-37 | 7.59E-36 |
| ADSS1 | -1.571748276 | 4.51E-23 | 8.58E-22 |
| LINC00954 | -1.571893527 | 0.00505086 | 0.01278608 |
| SLC39A10 | -1.573408311 | 1.55E-31 | 4.74E-30 |
| SKOR1 | -1.581127673 | 0.01850288 | 0.04084765 |
| AZIN2 | -1.582102528 | 0.02073964 | 0.04506353 |
| CLEC10A | -1.583064457 | 3.26E-08 | 1.87E-07 |
| ALDH2 | -1.58596121 | 1.66E-92 | 4.74E-90 |

| | | | |
|---|---|---|---|
| CXCL9 | -1.587233092 | 3.81E-09 | 2.41E-08 |
| SNX7 | -1.590575656 | 2.51E-05 | 9.79E-05 |
| CNTNAP1 | -1.591645072 | 5.54E-14 | 5.52E-13 |
| RNASE6 | -1.592655396 | 3.03E-84 | 6.87E-82 |
| PYHIN1 | -1.593830357 | 4.12E-07 | 2.08E-06 |
| NMNAT3 | -1.594760455 | 6.65E-12 | 5.53E-11 |
| RAG1 | -1.595033753 | 1.52E-10 | 1.10E-09 |
| CD22 | -1.59550776 | 6.83E-24 | 1.37E-22 |
| ANKRD33B | -1.59883594 | 0.00287039 | 0.00767774 |
| ZDHHC23 | -1.599715639 | 3.35E-16 | 3.98E-15 |
| SLC9A9 | -1.599797786 | 1.01E-28 | 2.66E-27 |
| ZBTB16 | -1.599986412 | 2.29E-07 | 1.19E-06 |
| MMP25-AS1 | -1.601031317 | 5.87E-07 | 2.90E-06 |
| LURAP1 | -1.601260215 | 4.12E-08 | 2.34E-07 |
| SLC47A1 | -1.609585906 | 3.56E-38 | 1.51E-36 |
| VTN | -1.615522735 | 3.03E-05 | 0.00011647 |
| DMWD | -1.617663817 | 1.18E-17 | 1.58E-16 |
| CGN | -1.622532563 | 0.01322659 | 0.03028324 |
| LSR | -1.62254909 | 5.82E-16 | 6.82E-15 |
| TMEM163 | -1.623331146 | 1.40E-05 | 5.72E-05 |
| ACSS3 | -1.62692285 | 0.00615334 | 0.01527755 |
| SYNC | -1.629821876 | 0.00493281 | 0.01251791 |
| MYO1G | -1.641996124 | 3.93E-52 | 3.02E-50 |
| COLEC12 | -1.645535167 | 1.53E-36 | 5.97E-35 |
| MAP3K9 | -1.64594063 | 2.80E-10 | 1.98E-09 |
| FOLR2 | -1.647135172 | 1.81E-84 | 4.16E-82 |
| LOC1019277 | -1.651553117 | 0.00980974 | 0.02323226 |
| LOC1019293 | -1.65310222 | 1.31E-29 | 3.63E-28 |
| TNFSF12-TNI | -1.655762129 | 0.0214183 | 0.04639183 |
| CD72 | -1.657284761 | 1.54E-20 | 2.50E-19 |
| SULT1C2 | -1.657801127 | 4.83E-09 | 3.02E-08 |
| SPTLC3 | -1.660033402 | 1.91E-12 | 1.66E-11 |
| TMEM220-AS | -1.667606228 | 0.00310815 | 0.00825362 |
| MTARC2 | -1.667645661 | 3.36E-07 | 1.71E-06 |
| P3H4 | -1.668832556 | 1.31E-11 | 1.05E-10 |
| LMNTD2-AS1 | -1.669517584 | 9.46E-07 | 4.54E-06 |
| VNN1 | -1.670368673 | 4.49E-33 | 1.45E-31 |
| CYP2U1 | -1.671945387 | 3.12E-32 | 9.88E-31 |
| CD101 | -1.67452779 | 5.11E-35 | 1.81E-33 |
| SLC6A16 | -1.675683951 | 7.52E-05 | 0.00027208 |
| SMOC1 | -1.675917318 | 0.00036009 | 0.00115958 |
| FFAR4 | -1.681297755 | 1.04E-10 | 7.60E-10 |
| ELL3 | -1.685220824 | 4.21E-05 | 0.00015858 |

| | | | |
|---|---|---|---|
| C4orf36 | -1.685841709 | 0.00046297 | 0.00145936 |
| NHLRC4 | -1.686313796 | 1.81E-12 | 1.57E-11 |
| SPRY1 | -1.687077899 | 1.31E-07 | 7.04E-07 |
| LOC1019300 | -1.690344949 | 0.00116102 | 0.00336343 |
| HPSE | -1.692826155 | 2.10E-60 | 2.13E-58 |
| NCF4-AS1 | -1.696542652 | 0.01105366 | 0.02581582 |
| MX1 | -1.697483173 | 2.91E-06 | 1.31E-05 |
| ABCA7 | -1.697977063 | 1.00E-109 | 4.47E-107 |
| ACSM5 | -1.699357908 | 1.29E-05 | 5.27E-05 |
| TOX2 | -1.699556039 | 0.00198533 | 0.00548736 |
| TNFAIP2 | -1.700996104 | 1.93E-123 | 1.14E-120 |
| PPARGC1A | -1.704672126 | 9.33E-06 | 3.90E-05 |
| IL12RB1 | -1.704731557 | 1.60E-35 | 5.83E-34 |
| MCC | -1.70532341 | 4.58E-23 | 8.70E-22 |
| CAVIN2 | -1.707840258 | 0.0003806 | 0.00121918 |
| SIGLEC12 | -1.708541304 | 9.56E-07 | 4.59E-06 |
| ARRDC5 | -1.710711067 | 0.00335767 | 0.00884376 |
| IL18BP | -1.713619893 | 3.27E-48 | 2.21E-46 |
| RARRES1 | -1.713948879 | 7.59E-55 | 6.57E-53 |
| FABP3 | -1.714868513 | 1.13E-21 | 1.96E-20 |
| ANGPTL2 | -1.716723885 | 0.02021954 | 0.04410735 |
| CIITA | -1.718621337 | 3.81E-67 | 5.20E-65 |
| SLC22A16 | -1.730692723 | 9.63E-08 | 5.24E-07 |
| RASL11A | -1.732265783 | 8.34E-09 | 5.09E-08 |
| TNFSF13 | -1.738101313 | 1.74E-47 | 1.14E-45 |
| TRAPPC6A | -1.739374123 | 7.26E-50 | 5.17E-48 |
| FN3K | -1.740054933 | 2.69E-16 | 3.23E-15 |
| KCNMB4 | -1.740223671 | 7.39E-12 | 6.12E-11 |
| QPRT | -1.740606921 | 2.06E-53 | 1.67E-51 |
| RTN4RL1 | -1.741619117 | 0.00056871 | 0.00176143 |
| LDHD | -1.742250217 | 2.17E-38 | 9.28E-37 |
| CFAP61 | -1.744622437 | 9.85E-08 | 5.36E-07 |
| SLC6A11 | -1.745612784 | 1.65E-06 | 7.64E-06 |
| GNG2 | -1.74614418 | 3.27E-44 | 1.83E-42 |
| SEMA4C | -1.749971928 | 2.33E-45 | 1.40E-43 |
| ADORA3 | -1.750805181 | 1.76E-14 | 1.83E-13 |
| HDAC9 | -1.750840104 | 2.68E-15 | 2.95E-14 |
| FBXO32 | -1.755517889 | 3.26E-12 | 2.79E-11 |
| FAAH | -1.755768899 | 1.13E-21 | 1.96E-20 |
| LMTK3 | -1.75855459 | 3.07E-08 | 1.77E-07 |
| ARHGAP42 | -1.759764162 | 0.00249705 | 0.00676005 |
| CBFA2T3 | -1.769653824 | 1.17E-14 | 1.22E-13 |
| SLC18B1 | -1.770962258 | 4.47E-64 | 5.27E-62 |

| | | | |
|---|---|---|---|
| CEBPD | -1.772028144 | 1.58E-64 | 1.93E-62 |
| VNN2 | -1.772589019 | 2.93E-47 | 1.89E-45 |
| AFF3 | -1.774572312 | 0.00751316 | 0.01826424 |
| NCF1 | -1.775995264 | 2.31E-12 | 2.00E-11 |
| ABCG1 | -1.778448933 | 9.75E-23 | 1.81E-21 |
| GFRA2 | -1.78320106 | 4.83E-11 | 3.65E-10 |
| ATP8A1 | -1.78752565 | 1.89E-35 | 6.81E-34 |
| CABLES1 | -1.789252501 | 1.50E-44 | 8.60E-43 |
| SORL1 | -1.790054195 | 4.57E-30 | 1.31E-28 |
| SLC2A9 | -1.791803749 | 1.31E-88 | 3.34E-86 |
| RGS11 | -1.792360131 | 0.00653686 | 0.01613284 |
| CYSLTR1 | -1.795599301 | 2.17E-50 | 1.58E-48 |
| GPR160 | -1.796199318 | 4.36E-69 | 6.38E-67 |
| ITM2C | -1.798209706 | 1.59E-26 | 3.70E-25 |
| SULF2 | -1.802525762 | 1.30E-05 | 5.33E-05 |
| MAP4K1 | -1.80380389 | 4.07E-47 | 2.62E-45 |
| SLC24A4 | -1.806467485 | 3.67E-24 | 7.54E-23 |
| CORO2A | -1.807193795 | 1.89E-51 | 1.42E-49 |
| PLS1 | -1.808681401 | 2.35E-08 | 1.37E-07 |
| FCGR2B | -1.8107186 | 8.57E-93 | 2.48E-90 |
| SLC34A3 | -1.812679419 | 0.01012619 | 0.02390696 |
| TLR3 | -1.816926141 | 1.09E-13 | 1.06E-12 |
| AZU1 | -1.818419617 | 2.41E-18 | 3.40E-17 |
| SIDT1 | -1.818674922 | 7.50E-13 | 6.76E-12 |
| ASB2 | -1.818775072 | 0.00868049 | 0.02080524 |
| NLRP1 | -1.821114422 | 2.01E-45 | 1.22E-43 |
| LAMP5 | -1.823994552 | 0.0107076 | 0.02511045 |
| GRIN2C | -1.824218281 | 0.00575538 | 0.01438672 |
| CLCN4 | -1.830136416 | 1.41E-22 | 2.57E-21 |
| IPCEF1 | -1.830765857 | 7.00E-25 | 1.49E-23 |
| LINC02724 | -1.835724426 | 6.57E-07 | 3.23E-06 |
| EDNRB | -1.835808883 | 2.16E-15 | 2.40E-14 |
| PLEKHA6 | -1.836516913 | 0.0002157 | 0.00072396 |
| SMARCD3 | -1.838785277 | 1.10E-63 | 1.28E-61 |
| HCN2 | -1.840412797 | 6.15E-16 | 7.20E-15 |
| CROCC2 | -1.842505624 | 2.79E-09 | 1.78E-08 |
| LRG1 | -1.847877765 | 1.94E-07 | 1.02E-06 |
| CYTL1 | -1.853915964 | 3.26E-13 | 3.03E-12 |
| MAP2K6 | -1.855082176 | 1.11E-29 | 3.07E-28 |
| LOC728392 | -1.85653908 | 9.89E-07 | 4.74E-06 |
| MGC16275 | -1.856636475 | 3.87E-11 | 2.96E-10 |
| H3C6 | -1.85970979 | 0.01043776 | 0.0245508 |
| SMIM1 | -1.859986895 | 0.00016732 | 0.00057214 |

| | | | |
|---|---|---|---|
| TUBB4A | -1.864118443 | 0.00109123 | 0.00317798 |
| PDE6G | -1.865577162 | 2.27E-11 | 1.78E-10 |
| CC2D2A | -1.866840879 | 4.20E-28 | 1.06E-26 |
| LINGO3 | -1.867431489 | 7.15E-35 | 2.51E-33 |
| LOC200772 | -1.86950165 | 1.10E-20 | 1.81E-19 |
| SEMA6C | -1.870863457 | 7.65E-13 | 6.90E-12 |
| NUPR1 | -1.878238499 | 7.73E-15 | 8.22E-14 |
| SOCS2 | -1.879420316 | 5.10E-05 | 0.00018937 |
| LOC1019292 | -1.879691298 | 0.0006664 | 0.00203464 |
| SERPINF2 | -1.880404052 | 1.20E-11 | 9.66E-11 |
| SLC17A7 | -1.883865957 | 1.45E-14 | 1.52E-13 |
| C9orf139 | -1.887679078 | 7.50E-28 | 1.88E-26 |
| C21orf62-AS1 | -1.887913955 | 0.00042591 | 0.00135363 |
| LINC02352 | -1.888420368 | 0.00267339 | 0.00719505 |
| FCN1 | -1.889385663 | 4.19E-44 | 2.33E-42 |
| C17orf113 | -1.891142932 | 4.88E-07 | 2.43E-06 |
| SERPINF1 | -1.894979761 | 4.86E-79 | 9.34E-77 |
| BANK1 | -1.895391571 | 3.26E-11 | 2.51E-10 |
| DBP | -1.896270811 | 9.93E-49 | 6.84E-47 |
| PLCL1 | -1.899004422 | 1.51E-26 | 3.52E-25 |
| RIPOR2 | -1.900608115 | 3.17E-11 | 2.45E-10 |
| TENT5B | -1.901298156 | 0.00094633 | 0.00279788 |
| PLAAT4 | -1.9031506 | 1.17E-14 | 1.23E-13 |
| IL6R-AS1 | -1.903333372 | 1.93E-10 | 1.38E-09 |
| RASGRP2 | -1.905729961 | 6.95E-19 | 1.01E-17 |
| GASK1A | -1.910613552 | 3.36E-09 | 2.13E-08 |
| YPEL1 | -1.91187254 | 2.19E-10 | 1.56E-09 |
| TMEM38A | -1.913218417 | 3.42E-05 | 0.0001306 |
| HLA-DMA | -1.921011716 | 7.12E-162 | 8.41E-159 |
| LYPD3 | -1.9224029 | 1.11E-07 | 6.02E-07 |
| ADGRE3 | -1.922601456 | 0.00289559 | 0.00773388 |
| TMEM63C | -1.923069366 | 1.98E-35 | 7.11E-34 |
| CX3CR1 | -1.926785317 | 2.09E-05 | 8.30E-05 |
| FGL2 | -1.938886724 | 1.61E-123 | 9.88E-121 |
| HSD17B14 | -1.945251703 | 3.42E-129 | 2.36E-126 |
| CERS4 | -1.945409157 | 2.00E-41 | 1.01E-39 |
| ENTPD1 | -1.946148098 | 5.44E-70 | 8.26E-68 |
| C16orf74 | -1.948933559 | 1.49E-06 | 6.95E-06 |
| SULT1B1 | -1.956612343 | 6.65E-23 | 1.25E-21 |
| ETV1 | -1.961277076 | 4.00E-10 | 2.79E-09 |
| CTTNBP2 | -1.965732153 | 2.32E-12 | 2.01E-11 |
| FRMD3 | -1.969872182 | 1.68E-23 | 3.28E-22 |
| PALD1 | -1.975686332 | 1.05E-12 | 9.35E-12 |

| | | | |
|---|---|---|---|
| FGD2 | -1.976787507 | 4.64E-96 | 1.50E-93 |
| HCG26 | -1.982277996 | 1.06E-06 | 5.08E-06 |
| TRPC1 | -1.982854029 | 0.01555287 | 0.03502732 |
| FXYD6 | -1.988637841 | 4.86E-73 | 8.04E-71 |
| CRHBP | -1.993704386 | 7.87E-16 | 9.10E-15 |
| CACNA2D3 | -2.002669026 | 1.67E-56 | 1.52E-54 |
| ASIP | -2.008173735 | 0.00361536 | 0.00944278 |
| SGPP2 | -2.009418398 | 7.40E-39 | 3.25E-37 |
| SNN | -2.014529125 | 1.41E-280 | 7.77E-277 |
| RBP1 | -2.015741306 | 4.33E-14 | 4.35E-13 |
| SIX5 | -2.016647695 | 1.29E-12 | 1.14E-11 |
| CEACAM4 | -2.019385087 | 7.61E-17 | 9.56E-16 |
| AMOT | -2.032412402 | 5.13E-06 | 2.22E-05 |
| MATN2 | -2.033505212 | 0.0023886 | 0.00649306 |
| GATM | -2.03577812 | 3.63E-93 | 1.07E-90 |
| GALNT3 | -2.046687129 | 1.79E-16 | 2.18E-15 |
| TRIM58 | -2.050530274 | 1.92E-22 | 3.46E-21 |
| TMEM229B | -2.052660654 | 1.26E-15 | 1.43E-14 |
| PIK3IP1 | -2.053308987 | 1.58E-59 | 1.56E-57 |
| CLUL1 | -2.053339669 | 0.01271744 | 0.02921866 |
| SPATA41 | -2.056882151 | 0.01021341 | 0.02408819 |
| NINL | -2.056972171 | 5.04E-10 | 3.48E-09 |
| NR3C2 | -2.058830307 | 1.72E-11 | 1.37E-10 |
| CALCR | -2.065662168 | 0.00110374 | 0.0032127 |
| OR2W3 | -2.067143344 | 5.76E-05 | 0.00021207 |
| JPH4 | -2.070526251 | 6.25E-36 | 2.34E-34 |
| HACD4 | -2.071563945 | 1.76E-70 | 2.75E-68 |
| DPEP2 | -2.078787121 | 1.92E-51 | 1.43E-49 |
| NCF1C | -2.086173706 | 2.29E-36 | 8.85E-35 |
| DMPK | -2.087020861 | 2.92E-08 | 1.69E-07 |
| NCF1B | -2.089217814 | 2.14E-40 | 1.02E-38 |
| TGFB2 | -2.090575305 | 6.15E-09 | 3.79E-08 |
| TESC | -2.092016645 | 3.76E-41 | 1.87E-39 |
| SLC30A4 | -2.092574982 | 2.37E-36 | 9.13E-35 |
| FAM157C | -2.093215656 | 0.01269437 | 0.02918187 |
| CHN2 | -2.094603449 | 3.09E-70 | 4.78E-68 |
| HGF | -2.098787336 | 1.88E-76 | 3.35E-74 |
| RPGRIP1 | -2.100327231 | 2.10E-14 | 2.16E-13 |
| SDS | -2.105385835 | 1.82E-52 | 1.43E-50 |
| RAVER2 | -2.114019245 | 6.68E-12 | 5.55E-11 |
| CCDC152 | -2.115633997 | 3.57E-18 | 4.97E-17 |
| GRIP1 | -2.116848008 | 1.06E-09 | 7.12E-09 |
| ASAP3 | -2.119285061 | 5.58E-05 | 0.00020596 |

| | | | |
|---|---|---|---|
| LINC01356 | -2.12325557 | 0.00060109 | 0.00185406 |
| TSPAN10 | -2.126980428 | 0.00144582 | 0.00410154 |
| LTC4S | -2.128156398 | 2.36E-12 | 2.04E-11 |
| SULT1A2 | -2.129746785 | 7.86E-05 | 0.00028366 |
| DEPTOR | -2.13923909 | 1.28E-44 | 7.37E-43 |
| ETV7 | -2.146800864 | 0.00106859 | 0.00311972 |
| TLR7 | -2.148729032 | 1.90E-98 | 6.54E-96 |
| ZG16B | -2.149175792 | 3.76E-22 | 6.72E-21 |
| CEACAM3 | -2.153169004 | 6.21E-12 | 5.17E-11 |
| IFI44L | -2.154638237 | 9.15E-05 | 0.0003263 |
| GJB2 | -2.161318949 | 6.93E-19 | 1.01E-17 |
| LINC00612 | -2.162395152 | 0.0198947 | 0.04350782 |
| IGFBP4 | -2.165389108 | 1.92E-62 | 2.13E-60 |
| STON1 | -2.171268979 | 0.00049169 | 0.00154282 |
| GBP4 | -2.176271946 | 2.79E-31 | 8.44E-30 |
| PHOSPHO1 | -2.177342756 | 2.21E-20 | 3.53E-19 |
| SV2C | -2.17900595 | 0.00037345 | 0.0011984 |
| WNT9B | -2.185395922 | 0.00705973 | 0.01728651 |
| HSF4 | -2.186176596 | 1.08E-66 | 1.44E-64 |
| PDGFB | -2.188780998 | 3.53E-15 | 3.84E-14 |
| LINC02384 | -2.193412071 | 0.00466066 | 0.01189665 |
| PRR5L | -2.199939589 | 1.14E-58 | 1.10E-56 |
| SCN1B | -2.203019619 | 5.18E-68 | 7.32E-66 |
| GPRC5C | -2.205394046 | 4.21E-14 | 4.23E-13 |
| HEYL | -2.20714311 | 5.86E-12 | 4.90E-11 |
| FLJ44635 | -2.207473878 | 4.33E-10 | 3.01E-09 |
| ZBP1 | -2.219881816 | 4.66E-07 | 2.33E-06 |
| CPNE9 | -2.22097072 | 5.30E-18 | 7.28E-17 |
| LILRA2 | -2.227348758 | 2.47E-68 | 3.55E-66 |
| TGFA | -2.228096538 | 1.75E-13 | 1.66E-12 |
| FAXDC2 | -2.231874851 | 1.51E-34 | 5.20E-33 |
| TPPP | -2.232393247 | 1.87E-08 | 1.10E-07 |
| KLF12 | -2.232760719 | 9.15E-24 | 1.81E-22 |
| LOC1053692 | -2.234789984 | 0.01700158 | 0.03787216 |
| CD200R1 | -2.235960116 | 6.94E-06 | 2.95E-05 |
| B3GNT7 | -2.243308282 | 5.92E-63 | 6.73E-61 |
| LINC01480 | -2.243485046 | 7.49E-06 | 3.18E-05 |
| VNN3 | -2.244205444 | 0.00123581 | 0.00356199 |
| SMPD3 | -2.251541422 | 8.37E-13 | 7.52E-12 |
| ACSM4 | -2.254450976 | 2.52E-15 | 2.78E-14 |
| MPPED2 | -2.266193584 | 1.38E-13 | 1.32E-12 |
| DYSF | -2.267357841 | 3.12E-261 | 1.29E-257 |
| KCNMA1 | -2.270534865 | 6.93E-71 | 1.09E-68 |

| | | | |
|---|---|---|---|
| GSTM2 | -2.271054532 | 7.59E-20 | 1.17E-18 |
| ISY1-RAB43 | -2.271431536 | 0.00517639 | 0.01308583 |
| NME8 | -2.273824318 | 1.30E-09 | 8.64E-09 |
| CPA6 | -2.280889331 | 0.00787833 | 0.01907064 |
| GLYATL1 | -2.281406818 | 0.0001834 | 0.00062327 |
| C10orf105 | -2.28617852 | 3.18E-13 | 2.95E-12 |
| TNFSF10 | -2.289487921 | 1.79E-40 | 8.62E-39 |
| HSPA7 | -2.291319736 | 2.23E-12 | 1.94E-11 |
| FCGR2C | -2.301856945 | 5.24E-12 | 4.41E-11 |
| KRT36 | -2.309660835 | 0.00023637 | 0.00078677 |
| GPBAR1 | -2.313908389 | 3.60E-14 | 3.64E-13 |
| LOC1005070 | -2.320879822 | 2.41E-22 | 4.34E-21 |
| LINC00324 | -2.325097847 | 3.33E-16 | 3.96E-15 |
| HHIPL1 | -2.325159897 | 1.18E-05 | 4.85E-05 |
| HLA-DQB1-A: | -2.327773519 | 1.04E-07 | 5.64E-07 |
| LINC00996 | -2.329056398 | 1.71E-09 | 1.12E-08 |
| AOAH | -2.329899213 | 4.46E-97 | 1.48E-94 |
| TBX1 | -2.334320074 | 2.33E-07 | 1.21E-06 |
| PGM5 | -2.360239285 | 3.92E-09 | 2.47E-08 |
| CHRNA6 | -2.365499925 | 1.04E-11 | 8.46E-11 |
| SLC26A8 | -2.369063563 | 0.00372396 | 0.00970191 |
| LINC00677 | -2.374605397 | 8.12E-11 | 6.01E-10 |
| LINC01960 | -2.374863416 | 0.00593885 | 0.01480281 |
| ADHFE1 | -2.377875859 | 3.24E-15 | 3.54E-14 |
| SESN3 | -2.38897888 | 1.75E-14 | 1.82E-13 |
| LEP | -2.390392582 | 1.84E-15 | 2.07E-14 |
| C8orf37-AS1 | -2.398156161 | 0.00114051 | 0.00331042 |
| PATJ | -2.408041048 | 1.98E-05 | 7.86E-05 |
| ADGRE1 | -2.411697218 | 1.31E-99 | 4.80E-97 |
| INPP4B | -2.418064252 | 5.87E-21 | 9.79E-20 |
| OPRD1 | -2.420135475 | 9.13E-06 | 3.83E-05 |
| VSTM1 | -2.422774718 | 8.08E-13 | 7.27E-12 |
| PEAK3 | -2.424685996 | 8.33E-36 | 3.07E-34 |
| RPL31P11 | -2.427392479 | 0.00651754 | 0.01608995 |
| SPOCK1 | -2.431658826 | 6.26E-10 | 4.28E-09 |
| ADRB1 | -2.436028508 | 1.36E-10 | 9.90E-10 |
| MAMDC2 | -2.444279502 | 0.00221821 | 0.00607481 |
| LOC1019277 | -2.444924008 | 0.01434912 | 0.03262743 |
| NLRP6 | -2.465027909 | 1.02E-06 | 4.89E-06 |
| PRICKLE1 | -2.465971833 | 3.55E-09 | 2.25E-08 |
| TMEM255A | -2.46744256 | 1.02E-37 | 4.21E-36 |
| SMAD3 | -2.468297823 | 1.44E-18 | 2.06E-17 |
| TMEM154 | -2.481579339 | 6.61E-36 | 2.46E-34 |

| | | | |
|---|---|---|---|
| C11orf21 | -2.493382271 | 1.50E-22 | 2.73E-21 |
| LINC01857 | -2.508680313 | 1.19E-64 | 1.47E-62 |
| SLC35F3 | -2.508845978 | 4.93E-12 | 4.15E-11 |
| CACNB4 | -2.508922608 | 7.62E-05 | 0.00027541 |
| RAB39B | -2.512387561 | 0.00529757 | 0.01335532 |
| CR1 | -2.513176644 | 1.88E-201 | 3.45E-198 |
| CXCR2 | -2.514159041 | 6.90E-13 | 6.25E-12 |
| COL9A3 | -2.545390167 | 1.42E-09 | 9.40E-09 |
| AATBC | -2.552390081 | 4.63E-83 | 9.93E-81 |
| ALDH1A1 | -2.554712211 | 4.98E-212 | 1.03E-208 |
| STON2 | -2.557661149 | 4.65E-12 | 3.92E-11 |
| RETN | -2.563704415 | 2.60E-05 | 0.00010126 |
| HAMP | -2.564866573 | 1.74E-11 | 1.38E-10 |
| LRRC4 | -2.574724118 | 5.84E-59 | 5.68E-57 |
| FGF13 | -2.575793122 | 0.00578168 | 0.01445028 |
| ST6GALNAC: | -2.582751001 | 1.10E-14 | 1.16E-13 |
| MLC1 | -2.584109058 | 2.42E-29 | 6.58E-28 |
| NYNRIN | -2.58693112 | 4.29E-40 | 2.02E-38 |
| TSPAN32 | -2.59076211 | 2.13E-29 | 5.80E-28 |
| HLA-DOA | -2.600134815 | 7.31E-44 | 4.02E-42 |
| MVB12B | -2.602220539 | 2.25E-89 | 5.82E-87 |
| OLFML3 | -2.604935071 | 5.84E-31 | 1.73E-29 |
| CNR2 | -2.605782039 | 5.92E-05 | 0.00021746 |
| RHPN2 | -2.617736785 | 0.0003735 | 0.0011984 |
| ALK | -2.618647264 | 3.64E-141 | 3.17E-138 |
| ABCD2 | -2.626437008 | 0.00042688 | 0.00135567 |
| GPA33 | -2.633863184 | 0.00051615 | 0.00161437 |
| PHYHD1 | -2.644094806 | 1.21E-13 | 1.17E-12 |
| DPF3 | -2.661660014 | 5.73E-06 | 2.47E-05 |
| MMP25 | -2.676267385 | 1.82E-132 | 1.37E-129 |
| ALDH3A1 | -2.683156759 | 0.00060125 | 0.00185421 |
| ZNF385C | -2.698674143 | 2.44E-07 | 1.26E-06 |
| LOC1019281 | -2.699432255 | 0.0102393 | 0.02413894 |
| METTL7B | -2.70432821 | 7.14E-40 | 3.31E-38 |
| MEFV | -2.704801296 | 1.64E-51 | 1.24E-49 |
| HLA-DMB | -2.708812068 | 0 | 0 |
| LINC01451 | -2.710298845 | 1.14E-05 | 4.72E-05 |
| SCTR | -2.710715784 | 2.20E-05 | 8.70E-05 |
| AOC1 | -2.717142293 | 9.96E-23 | 1.84E-21 |
| PHYHIP | -2.717886832 | 0.02278831 | 0.04903802 |
| IL2RA | -2.718388793 | 5.43E-16 | 6.37E-15 |
| PKP2 | -2.743619613 | 5.87E-06 | 2.52E-05 |
| CXCR2P1 | -2.744242304 | 2.96E-12 | 2.54E-11 |

| | | | |
|---|---|---|---|
| NEURL3 | -2.749453808 | 1.69E-14 | 1.76E-13 |
| PART1 | -2.755266158 | 0.00976187 | 0.02312552 |
| JAML | -2.756625936 | 3.56E-219 | 8.41E-216 |
| CXCL12 | -2.768210171 | 1.63E-10 | 1.18E-09 |
| SCRT2 | -2.774563168 | 2.95E-07 | 1.51E-06 |
| COL6A4P2 | -2.785791431 | 0.01334066 | 0.03053172 |
| P2RY8 | -2.792256248 | 3.18E-136 | 2.63E-133 |
| FUCA1 | -2.796014279 | 3.03E-34 | 1.03E-32 |
| C1orf127 | -2.796866558 | 2.01E-41 | 1.01E-39 |
| ITLN1 | -2.802192562 | 0.0004602 | 0.00145118 |
| RERE-AS1 | -2.809645076 | 0.0151372 | 0.03419366 |
| CT75 | -2.813693314 | 1.02E-20 | 1.67E-19 |
| DTX1 | -2.821813949 | 4.01E-21 | 6.75E-20 |
| TMEM266 | -2.824500715 | 8.44E-23 | 1.57E-21 |
| CHRNA3 | -2.834201043 | 0.00279452 | 0.00749057 |
| ZNF704 | -2.836712852 | 1.07E-49 | 7.55E-48 |
| NFATC2 | -2.838654479 | 8.69E-82 | 1.82E-79 |
| BMF | -2.854254244 | 2.59E-161 | 2.86E-158 |
| JUP | -2.858056294 | 4.14E-17 | 5.32E-16 |
| MYO1A | -2.859860778 | 9.77E-22 | 1.70E-20 |
| OTOAP1 | -2.867166579 | 3.04E-90 | 8.10E-88 |
| MS4A6A | -2.876405955 | 0 | 0 |
| AXL | -2.879657141 | 1.05E-08 | 6.35E-08 |
| SULT2B1 | -2.884966121 | 1.51E-11 | 1.20E-10 |
| SELENOP | -2.906918188 | 2.23E-132 | 1.60E-129 |
| SMPD5 | -2.917745076 | 0.00086775 | 0.00258497 |
| LINC02688 | -2.940387592 | 0.00614572 | 0.01526321 |
| SEZ6L | -2.953639718 | 4.54E-11 | 3.44E-10 |
| GAS6 | -2.970836048 | 7.26E-194 | 1.20E-190 |
| P2RY12 | -3.000655517 | 0.01916487 | 0.04210669 |
| C16orf96 | -3.011979162 | 9.65E-06 | 4.02E-05 |
| PVALB | -3.018140829 | 4.02E-12 | 3.42E-11 |
| GNG7 | -3.046524593 | 1.85E-66 | 2.41E-64 |
| LOC1019283 | -3.047740718 | 6.91E-05 | 0.00025167 |
| CD163L1 | -3.048064221 | 5.68E-23 | 1.07E-21 |
| GPR55 | -3.055907735 | 4.05E-10 | 2.82E-09 |
| LINC02528 | -3.063790816 | 5.98E-06 | 2.57E-05 |
| ZFY-AS1 | -3.063796982 | 1.46E-05 | 5.95E-05 |
| IRF4 | -3.080101471 | 6.35E-37 | 2.51E-35 |
| PDK4 | -3.081844269 | 1.84E-22 | 3.33E-21 |
| VIPR1 | -3.090979402 | 2.92E-05 | 0.0001127 |
| LYPD5 | -3.095141454 | 1.21E-08 | 7.26E-08 |
| CD244 | -3.114470745 | 2.63E-47 | 1.71E-45 |

| | | | |
|---|---|---|---|
| LOC729296 | -3.114622378 | 0.00373174 | 0.00971909 |
| PDGFC | -3.120672734 | 1.86E-95 | 5.92E-93 |
| SPNS3 | -3.121164264 | 0.00317418 | 0.00841274 |
| NHSL2 | -3.12533387 | 3.15E-46 | 1.97E-44 |
| PKD2L1 | -3.176216931 | 3.36E-85 | 7.83E-83 |
| LINC00639 | -3.186016498 | 9.98E-09 | 6.04E-08 |
| APOL4 | -3.205120801 | 1.84E-08 | 1.09E-07 |
| CCL19 | -3.207173149 | 0.00076108 | 0.00229279 |
| GUCY1B2 | -3.251360158 | 0.02081496 | 0.0451975 |
| GCSAM | -3.272723579 | 1.23E-28 | 3.24E-27 |
| CPAMD8 | -3.273194238 | 2.08E-78 | 3.96E-76 |
| ASGR2 | -3.279810187 | 4.74E-23 | 8.97E-22 |
| S100Z | -3.30662647 | 7.11E-13 | 6.43E-12 |
| ITGA9 | -3.363663786 | 5.25E-57 | 4.85E-55 |
| SCAMP5 | -3.376431032 | 3.05E-15 | 3.35E-14 |
| LINC00671 | -3.393596322 | 1.96E-09 | 1.28E-08 |
| GAS6-DT | -3.408434566 | 1.27E-07 | 6.83E-07 |
| UNC5C | -3.414663375 | 0.00553318 | 0.01389431 |
| PNOC | -3.432812366 | 0.00427177 | 0.01099737 |
| PNPLA7 | -3.461559214 | 5.37E-82 | 1.14E-79 |
| CD2 | -3.498546457 | 0.00063058 | 0.00193492 |
| GPR33 | -3.516459294 | 0.00278732 | 0.00747369 |
| GAPT | -3.518302846 | 4.80E-116 | 2.40E-113 |
| MPZL2 | -3.558904923 | 1.26E-11 | 1.01E-10 |
| PCDH12 | -3.565369128 | 2.27E-63 | 2.61E-61 |
| KCNJ5 | -3.607948183 | 2.07E-33 | 6.76E-32 |
| PLA2G2D | -3.630969697 | 4.68E-08 | 2.64E-07 |
| PLXDC1 | -3.640340071 | 1.74E-13 | 1.65E-12 |
| METTL27 | -3.651182378 | 1.42E-09 | 9.37E-09 |
| GGTA1P | -3.66345632 | 1.02E-44 | 5.91E-43 |
| NRIR | -3.706831438 | 1.58E-05 | 6.38E-05 |
| PLPPR3 | -3.729256071 | 5.22E-11 | 3.93E-10 |
| KLRC4 | -3.729451603 | 8.90E-08 | 4.86E-07 |
| SRGAP3 | -3.73047784 | 7.40E-24 | 1.48E-22 |
| F13A1 | -3.741421229 | 3.24E-60 | 3.27E-58 |
| LOC1001302 | -3.745071458 | 8.15E-05 | 0.00029287 |
| LINC02073 | -3.747940123 | 1.93E-24 | 4.01E-23 |
| SELENBP1 | -3.754430066 | 0.00100553 | 0.00295388 |
| TMPRSS11D | -3.754716385 | 0.01940809 | 0.04255623 |
| ITIH1 | -3.786077381 | 0.01551113 | 0.03496191 |
| SPIC | -3.801034262 | 1.60E-10 | 1.16E-09 |
| MS4A4E | -3.813602828 | 1.02E-13 | 9.90E-13 |
| PADI2 | -3.829731804 | 2.66E-39 | 1.20E-37 |

| | | | |
|---|---|---|---|
| DAB2IP | -3.832406598 | 3.58E-15 | 3.89E-14 |
| HPN | -3.841654621 | 2.93E-16 | 3.50E-15 |
| TRIM50 | -3.859596034 | 9.45E-11 | 6.95E-10 |
| KCNJ10 | -3.8682961 | 3.59E-45 | 2.14E-43 |
| FAM3D | -3.888251499 | 0.00280521 | 0.00751783 |
| ITGAD | -3.9329064 | 1.40E-19 | 2.11E-18 |
| AQP12A | -3.975973886 | 0.00143693 | 0.00408051 |
| LOC1005063 | -3.993497357 | 6.68E-12 | 5.55E-11 |
| LOC646030 | -4.023406503 | 2.35E-09 | 1.51E-08 |
| PRLR | -4.025361348 | 5.53E-21 | 9.24E-20 |
| P2RY13 | -4.074267516 | 1.62E-43 | 8.71E-42 |
| MKX | -4.075003603 | 0.00105484 | 0.00308448 |
| HS3ST2 | -4.092921857 | 5.35E-67 | 7.21E-65 |
| NPM2 | -4.168942962 | 6.53E-05 | 0.00023844 |
| ATRNL1 | -4.170611046 | 0.0095499 | 0.0226721 |
| VCAM1 | -4.176681432 | 6.07E-06 | 2.60E-05 |
| SLC40A1 | -4.214099078 | 5.69E-136 | 4.48E-133 |
| IGF1 | -4.224975301 | 3.88E-188 | 5.35E-185 |
| TPPP3 | -4.267462724 | 0.01685766 | 0.03762004 |
| KLHL13 | -4.279335729 | 6.04E-37 | 2.40E-35 |
| PTCHD4 | -4.310233214 | 0.01143484 | 0.02657467 |
| ADGB | -4.338239994 | 0.00564459 | 0.01414184 |
| OLFML2A | -4.354891715 | 0.00013578 | 0.0004719 |
| LOC1019288 | -4.40932333 | 0.01107822 | 0.02586223 |
| KLRC4-KLRK | -4.442644396 | 9.97E-06 | 4.15E-05 |
| ABCG4 | -4.474333787 | 0.00030513 | 0.00099521 |
| SFRP4 | -4.527832603 | 0.0002796 | 0.00091754 |
| CCDC26 | -4.589377364 | 2.92E-08 | 1.69E-07 |
| SPATA13-AS | -4.657564757 | 0.00141012 | 0.00401334 |
| P2RY14 | -4.681593284 | 1.55E-10 | 1.12E-09 |
| NAT8B | -4.772842506 | 0.00105794 | 0.00309246 |
| STEAP4 | -4.862238273 | 7.38E-05 | 0.0002673 |
| DOC2A | -4.876636303 | 3.58E-13 | 3.31E-12 |
| XKR3 | -4.980492143 | 1.57E-05 | 6.37E-05 |
| CLDN4 | -5.046392503 | 0.00134317 | 0.00384066 |
| NECAB2 | -5.096006817 | 0.00289983 | 0.00774272 |
| WNT5B | -5.125599948 | 8.83E-06 | 3.71E-05 |
| SIGLEC8 | -5.169746302 | 0.0001743 | 0.00059417 |
| CYP4F12 | -5.258665395 | 0.0005847 | 0.00180556 |
| ROR2 | -5.334537647 | 4.67E-13 | 4.29E-12 |
| ARL5C | -5.456698295 | 0.00060312 | 0.00185895 |
| MAT1A | -5.806684959 | 0.00014755 | 0.00050949 |
| LINC01736 | -5.813518631 | 7.23E-05 | 0.00026259 |

| | | | |
|---|---|---|---|
| CEACAM1 | -5.919793902 | 1.57E-07 | 8.31E-07 |
| GPRC5B | -5.954110415 | 8.08E-18 | 1.10E-16 |
| LINC00173 | -6.287453313 | 1.21E-08 | 7.27E-08 |
| F2RL3 | -6.966045532 | 3.41E-64 | 4.08E-62 |
